# An *ab initio* information-theoretic approach to proteins and protein-ligand interactions

**DOI:** 10.1101/2024.03.06.583646

**Authors:** Deep Nath, Rajdeep Kaur Grewal, Devrani Mitra, Soumen Roy

## Abstract

Differing conformational structure of a protein, associated with two distinct signaling states or between ligand-free and ligand-bound states, leads to differing inter-residue interactions and consequently different biological function. We propose a fresh first-principles information-theoretic approach for studying such proteins and their interactions. A *de novo* measure called protein residue information (PRI), which incorporates details of interactions between all pairs of atoms within and across all residues of the protein, is introduced herein. We formulate a method to calculate the intrastate and inter-state entropy of every residue, needed to determine PRI across any two states of a protein. The intra-state entropy can be determined for every state of a protein possessing one or more states. The inter-state entropy can be calculated pairwise for proteins possessing more than one state. We analyze twenty eight distinct pairs of protein structures from ten different classes. PRI successfully identifies important residues displaying significant conformational changes bearing influence with respect to itself and all other residues. Furthermore, it also successfully identifies important residues displaying rather subtle conformational changes. The identified residues exhibit influential roles in diverse performative features of proteins like stability, allostery, signaling, etc. PRI successfully recovers known experimental results from literature and predicts important roles for many hitherto unstudied residues.

## I. INTRODUCTION

Understanding the relation between the structure and function of a protein is a grand aim of researchers in various disciplines. These are known to intimately influence each other. Generally, a protein’s structure is rather flexible and undergoes conformational changes leading to diverse functional outcomes. Conformational changes are triggered in the presence of stimuli, ligand binding and other environmental factors. Further, protein folding/misfolding and mutations [1] can also induce conformational fluctuations in a protein. During conformational changes, bonds between the atoms of amino acid sequences remain unchanged and the dihedral angles of the main chain and the orientation of side chains may get altered.

Even seemingly insignificant conformational changes could lead to large or remarkable changes in protein function. Examples range from alteration of catalytic constants of enzymes to negative or positive cooperativity of receptors or even functional alterations in the entire organisms such as swimming patterns of bacteria [2]. Residues can play a crucial role in allostery without undergoing conformational changes [3, 4]. Herein, we formulate an *ab initio* information-theoretic approach to study the consequence of changes in positions of atoms of all residues in a protein. These changes at the atomic-level decide the conformational changes in residues to a varying degree. Such changes, whether small or large, can significantly influence the function of a protein.

Molecular dynamics (MD) simulations provide robust insights into conformational changes of a protein [5, 6]. They deal with the varied dynamical and conformational structures of a protein, to predict associated functional properties. However, they are computationally expensive. It is highly challenging to capture delayed conformational changes on timescales outside the scope of MD simulations [7]. Furthermore, they invoke mathematical approximations to mitigate computational requirements, which in principle, can influence the outcome. Apart from MD, the transitions between various conformational structures could be understood using a protein contact network (PCN) [8, 9]. PCNs help in studying various conformational states of a protein [10]. They find its application in unraveling details of varied phenomena like protein folding, allostery [11], dimer stability [12], residue fluctuations, dynamic structure fluctuations, etc. However, the construction of these network models is nontrivially affected by the choice of the cutoff [8]. Further, these network models often overlook the specific nature of non-covalent interactions (van der Waals interactions or hydrogen bonding) and other complex aspects of the structure of a protein. Further, MD and PCN mostly identify the functional importance of residues, which have undergone significant conformational changes.

The information-theoretic approach proposed herein is plagued neither by demanding computational expense nor by the vagaries of cutoff distances. Moreover, the approach successfully identifies residues that display rather subtle conformational differences between two different states but have significant biological importance. Consider any two given states of the same protein. These states could be experimentally determined structures readily available in the Protein Data Bank (PDB), or obtained through approaches like homology modeling [13] or deep learning, viz. AlphaFold [14]. These structures can represent the protein: (a) in two distinct signaling states, (b) before and after ligand binding, or, (c) in two different conformations under various other stimuli. In response to external stimuli, the position of atoms of all residues in a protein could change leading to a change in position and/or conformation of the residues. However, it must also be remembered that when such changes occur, the surrounding residues often also rearrange themselves to avoid steric clashes so as to make the overall conformation stable. Consequently, the position and/or conformation of every residue with respect to its neighbouring and indeed all other residues also get altered. Both these changes the deviation of a residue with respect to itself and that of a residue with respect to every other residue across two structures of a protein, are of similar importance. As elaborated later, the information-theoretic approach presented herein incorporates both these kinds of changes.

Entropy, an outcome of randomness and disorder, imprints its signature across all areas of life. It is extremely useful in understanding many biological phenomena such as protein folding/unfolding, protein-protein interactions [15] and protein-ligand interactions [16]. Various methods have been introduced to calculate the entropy of proteins. Among these, configuration entropy (CE) is used to study interactions and functions of proteins [16, 17]. CE also finds application in MD simulations to study the folding and unfolding of a protein. Additionally, recent developments have enabled the study of conformational changes in protein folding with the help of conformational entropy [18]. There also exist other forms of entropy, which are distance-based. For example, the mathematical expression of entropy in PCN is strongly influenced by the basic criterion of distance-based cutoff during the very construction of PCN. The interaction between residues is effectively a step function and the choice of cutoff dictates the interaction pattern.

We start with experimentally determined macromolecular structures from PDB. Our formulation proposes that the entropy of a protein can be accounted across its states and is independent of any arbitrary cutoff. We can calculate these contributions separately: (a) in any given state of a protein (*“intra-state entropy”* ), and (b) across any two given states of a protein (*“inter-state entropy”* ). Generally, proteins display conformational change(s) across two given states due to events like signaling or ligandbinding. Due to such changes, the structure of proteins, and the associated entropy will likely differ. Broadly speaking, intra-state entropy can be associated with a particular conformation, while inter-state entropy can be associated with changes in conformation. Intra-state entropy of a residue in a given state is decided by interactions of every atom in that residue with every atom of every other residue. The significance of interactions between residues towards protein-folding and protein stability is well-studied in literature [19, 20]. These interactions are mainly responsible for the conformation of side chains of residues of a protein [20]. On the other hand, inter-state entropy aims to distinguish between the entropy of a residue in a given pair of states. From, these two forms of entropy, we can propose an expression and calculate the *protein residue information* (PRI) of every residue in the protein. By analysing twenty eight distinct pairs of protein structures drawn across ten different classes, we discuss extensively about how each of the residues possessing high PRI essay protein function. Indeed, we provide structure-based mechanistic insights for all residues with higher values of PRI. For well-studied systems, nearly all our findings agree quite well with reported experimental evidence. For the lesser studied systems, we predict hitherto unknown roles of residues, which is amenable to experimental verification.

Our formulation of a distance-dependent form of entropy is inspired by the extremely well-known Lennard-Jones (LJ) potential. LJ potential is very efficient in capturing the nature of interatomic interactions across all sciences. In biology, LJ potential finds its application in soft-docking approach, molecular dynamics simulations and others diverse problems. The use of the LJ potential limits the interaction between any two atoms in an amino acid sequence. This immediately presents the advantage of eliminating arbitrary cut-offs to portray interactions between any two atoms. Unlike MD simulations, herein we deal only with structures of a protein in two different states. Therefore, this technique is also rather computationally inexpensive.

## II. MATERIALS AND METHODS

Let *𝒫* and *𝒬* denote two *diferent states* of a protein. *{ℛ}* and *{𝒜}* respectively denotes the *set of residues* and the *set of all atoms in all residues*, which are simultaneously present in both *𝒫* and *𝒬*. Atoms, *i* and *j*, in residues, *k* and *l*, respectively are separated by the Euclidean distance, 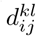, where *k, l ∈ {ℛ}*, and *i, j ∈ {𝒜}*. The strength of these interatomic interactions between atoms in neighboring residues would expectedly be higher than that between distant residues. Naturally, interatomic interactions would decrease with the increase of 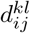

Inspired by well-known Lenard-Jones potential, herein we propose,

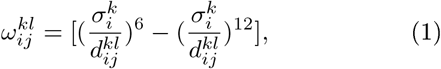

where, 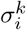 is a constant for a given atom in a residue. 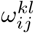 is dimensionless. The value of 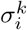 of atom, *i ∈ {𝒜}*, belonging to residue, *k ∈ {ℛ}*, is 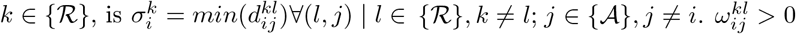 and its value depends only on 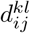 Between two states, all atoms in all residues might undergo conformational changes. Hence, we consider all possible pairwise combinations of atoms, i.e., both intra-residue and inter-residue, to calculate 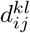. Details about interactions between atoms within and among residues and the calculation of 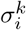 are depicted in Fig. 1.

**FIG. 1.**
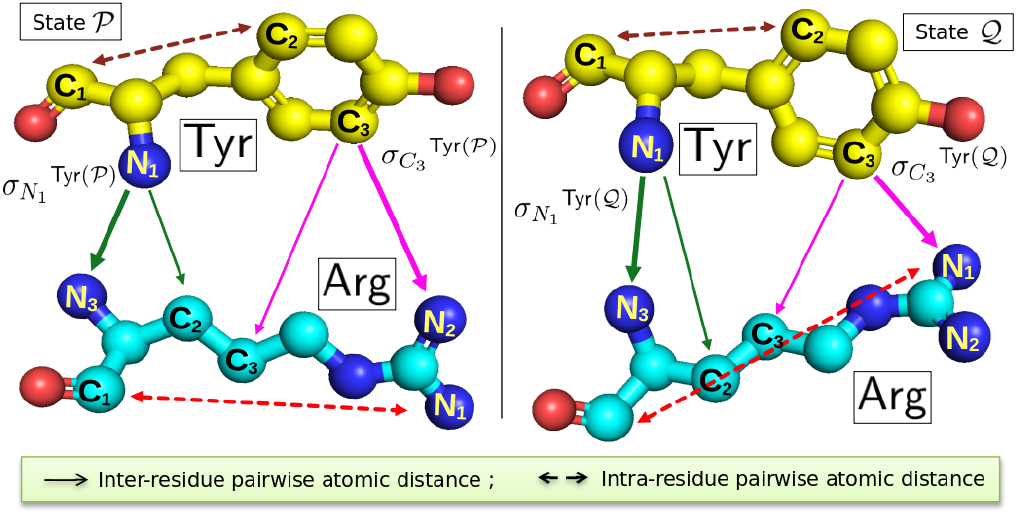
Illustrative example of two residues of a protein, tyrosine, Tyr, (highlighted in yellow) and arginine, Arg, (highlighted in cyan), undergoing conformational change across the states, 𝒫 and𝒬. Here, we observe that the intra-residue pairwise atomicdistances between *C*_1_ and *C*_2_ of Tyr (highlighted in brown dashed line), and, *C*_1_ and *N*_1_ of Arg (highlighted in red dashed line) change between 𝒫 and𝒬. The inter-residue distances of atoms, *N*_1_ and *C*_3_, of Tyr with two different atoms of Arg, depicted in green and magenta respectively, also change across 𝒫 and𝒬. ln𝒫, the minimum of all these respective distances of *N*_1_ and *C*_3_ of Tyr with respect to every atom of Arg equals 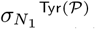 and 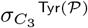 respectively. Similarly, 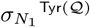 and 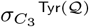 can be determined analogously for 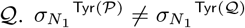 and 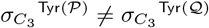. Gen-erally, 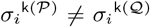, for the *i*^*th*^ atom of the *k*^*th*^ residue of any protein.

Fig. 2 exhibits the conformational change of various residues in two distinct states (holo and apo states) of streptavidin, a biotin-binding protein. As observed in Figs. 2 (a) and 2 (b), the distance between *O*_1_ in Glu51 and *N*_1_ in Arg84 changes from 5.2Å to 5.4Å, implying an increase of only 3.8% across the two states. On the other hand, we observe in Figs. 2 (c) and 2(d), that the distance between *O*_2_ in Glu44 and *N*_1_ in Arg53 across the two states decreases by 61.6% from 7.3Å to 2.8Å. These changes smaller or larger - in the respective distance between all pairs of atoms across two given states are important to assess the overall conformational changes and its effects, as discussed in Results. Therefore, in our formulation, we consider the most general version - that entropy depends on the distance between all pairs of atoms within and between all residues in a given state and the changes in them across any two states.

**FIG. 2.**
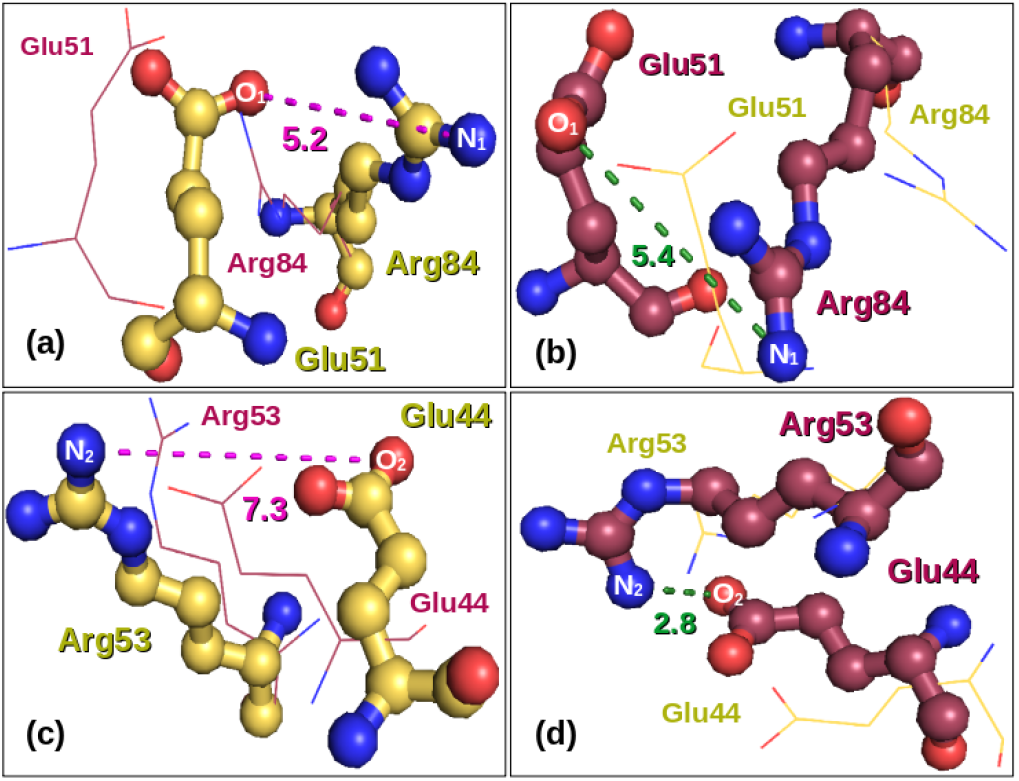
1SWE [(a) and (c)] and 1SWA [(b) and (d)] represent the holo (highlighted in golden) and apo (highlighted in maroon) states of streptavidin respectively. The conformation of Glu51 and Arg84 is depicted in (a) and (b), while that of Glu44 and Arg53 is depicted in (c) and (d).The numbers below dotted lines indicate distance in Å.

### A. Entropy

The entropy of a random event, *x*_*i*_ *∈ 𝒳*, is related to the amount of information or uncertainty associated with that event. Here 𝒳 denotes a complete set of disjoint random events, such that ⋃_*i*_ *x*_*i*_ = Ω, and *x*_*i*_ *⋂x*_*j*_ = ∅. If *p*(*x*_*i*_) is the probability of occurrence of event *x*_*i*_, then the entropy, 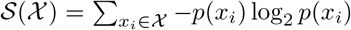. Similarly, if *p*(*x*_*i*_, *y*_*i*_) represents the joint probability of occurrence of two events *x*_*i*_ *∈ 𝒳*, and *y*_*i*_ *∈ 𝒴*, then the joint entropy, 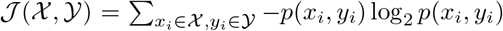. We outline below the procedure for calculation of both intrastate entropy and inter-state entropy.

### B. Intra-state entropy

Herein, we evaluate the entropy associated with each individual residue of a protein in a given state. Henceforth, we refer to 𝒮_*k*_ as the normalised *“intra-state entropy”* of the *k*^*th*^ residue in a given protein. 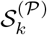 and 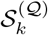 denote the same for two different states, *𝒫* and *𝒬*, respectively of the protein. 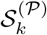 and 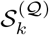 is decided by the underlying structure of that particular state and can be captured adequately through 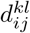 and 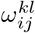.

Considering the most general formulation possible, we posit that the probability, 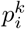, of conformational change of the *i*^*th*^ atom of the *k*^*th*^ residue is influenced in varying measure by all other atoms in every residue excluding the *k*^*th*^ residue. Therefore,

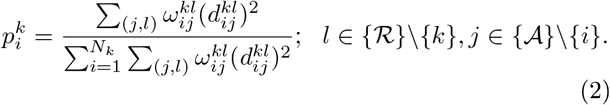

The choice of the functional form, 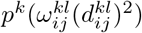, is guided by overwhelming experimental evidence and is elaborated in the S.I.

The corresponding intra-state entropy of the *k*^*th*^ residue equals 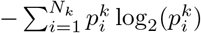. Its normalised intra-state entropy therefore is,

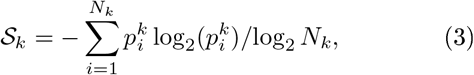

where, *N*_*k*_, is the total number of atoms present in the *k*^*th*^ residue of the protein. A detailed discussion on the physical significance of *𝒮*_*k*_ with concrete examples can be found in the S.I.

### C. Inter-state Entropy

Let 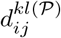and 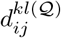denote the pairwise atomic distances in two different states 𝒫 and 𝒬 respectively of a protein. The normalised “inter-state entropy”, 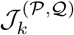, of the *k*^*th*^ residue is calculated *across the states 𝒫 and 𝒬*, and depends on 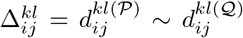 with *k, l ∈ { ℛ }* and *i, j ∈ { 𝒜 }*.

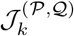 is calculated from the structures of two different states of a protein, *and does not arise from direct interactions between atoms of a protein in any given state*. Therefore, physical interactions manifested through 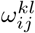and 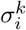 in the case of *𝒮*_*k*_, are not relevant for the determination of 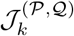. Similar to the expression of 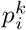 used to determine *𝒮*_*k*_, we posit the probability,

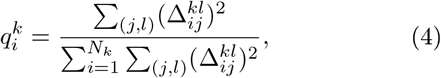

for the determination of inter-state entropy. Thence, we can easily compute the normalised inter-state entropy of the *k*^*th*^ residue,

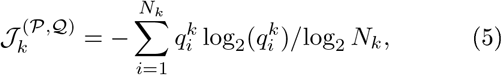

across states, 𝒫 and 𝒬, of the protein. If a given protein has more than two different states, 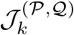can be computed *pairwise ∀{*𝒫, 𝒬*}*, 𝒫 *≠* 𝒬. Further details can be found in the S.I.

### D. Protein Residue Information

Intra-state entropy is associated with the possible interactions experienced by a residue due to other residues. Inter-state entropy incorporates the shift in the position of a residue between two states. To identify functionally important residues, we need to consider the conformation of a residue as well as the interactions it participates in. Therefore, protein residue information (PRI) of residue, *k*, across two different states, *𝒫* and *𝒬*, is expressed as,

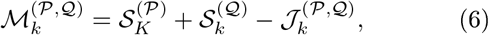

If a given protein has more than two different states, 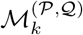can be computed *pairwise∀{*𝒫, 𝒬*}*, 𝒫 *≠* 𝒬.. As aforementioned, the underlying thought behind *PRI* is influenced by mutual information (MI). Notionally, MI helps in quantifying the amount of information between

two variables, events, or systems. Like MI, 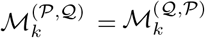. However, an interesting difference exists between MI and PRI. While various reports of negative MI in quantum information and entanglement exist [21, 22], it is well-known to be positive in classical physics. *ℳ*_*k*_ can be negative on occasion, as mentioned in the S.I. This is not unexpected owing to the difference in the physical origin and functional dependence of intra-state and interstate entropy. To reiterate, *𝒮*_*k*_ *d*epends on the Euclidean distance between pairs of atoms and is calculated *for any given state*. In contrast, *𝒥*_*k*_ depends on the *diference* in Euclidean distance between atomic pairs and is calculated *across a pair of states*.

## III. RESULTS AND DISCUSSION

We analyze twenty eight distinct pairs of protein structures from ten different classes. Each pair represents two distinct states in PDB, 𝒫 and 𝒬. Table I displays all residues possessing 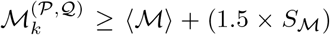. The value of 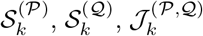, and 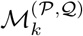 of every residue for each of these twenty eight pairs of protein structures is provided in pp. 10-47 of the Supporting Information (S.I.). 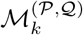 denotes the protein residue information of the *k*^*th*^ residue. *⟨ ℳ ⟩* and *𝒮*_*ℳ*_ respectively denote the mean and standard deviation of the distribution of 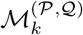 drawn over all residues of a given protein possessing states 𝒫 and 𝒬. We find that not all residues with high PRI values undergo large conformational changes as elaborated in the S.I. This is important because solvent exposed residues with larger side chain could undergo significant conformational changes but bear no structural or functional relevance. On the other hand, some residues often maintain their interactions with the neighbouring residues and are biologically significant. For example, Trp83 in MutY retains its interactions in both states and plays a crucial role in the stability and functionality of MutY, as elaborated later. Our analysis helps in identifying residues that are structurally and functionally important for a protein. Apart from the functional significance of residues with higher values of PRI, we also discuss their conformational changes in great detail in the S.I.

**TABLE 1.**
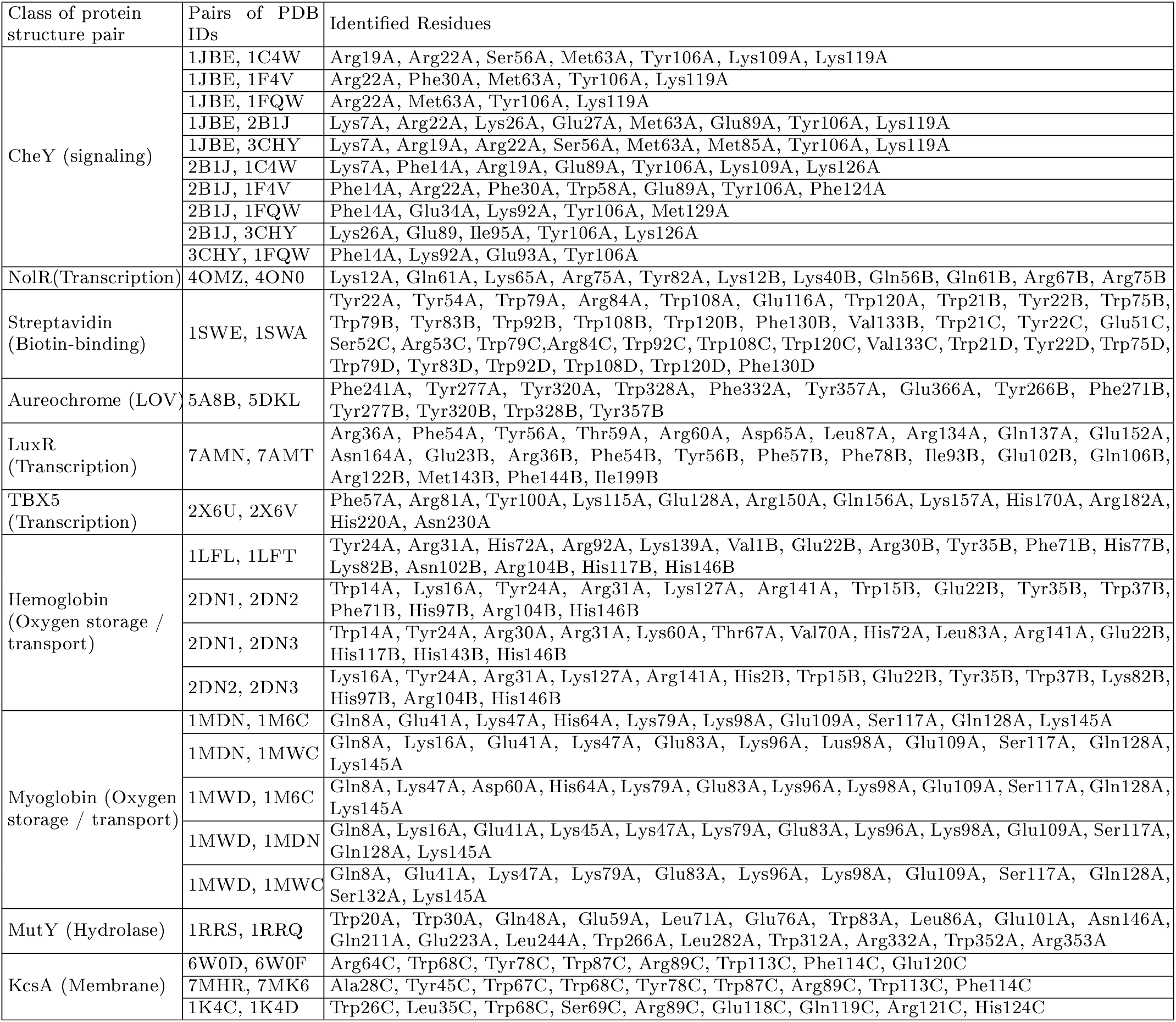
Residues possessing a value greater than 1.5 times the standard deviation from the mean, of the distribution of protein residue information for the respective pair of protein structures. Functional associations of each of these are provided in the text. The roles of the vast majority of these residues have already been verified experimentally.

### A. CheY

We first investigate CheY, which is a 14KDa response regulator protein well known for chemotaxis and thoroughly studied for its allosteric properties. As plenty of experimental structures are available in PDB, we investigate six different PDB structures in ten different combinations. The structures are: 1JBE (apo CheY in meta-active state - antecedent to fully activated state), 1C4W (D57C mutant), 1F4V (activated, phosphorylated CheY bound to FLiM), 1FQW (activated CheY), 2B1J (unphosphorylated CheY bound to FLiM) and 3CHY (apo-CheY). Three distinct molecular events characterise CheY activation: phosphorylation, conformational changes associated with *β*4 *− α*4 loop, and rotational restriction of Tyr106 side chain [23]. While the number and nature of identified amino acids vary in different combinations for obvious reasons, we find an interesting trend. Tyr106 possesses a high value of PRI in all ten combinations of CheY structures as reflected in Table I. Indeed, it is known to play a very crucial role in the activation of CheY [23-25]. It stabilizes the activated configuration of *β*4 *− α*4 loop and is present in most of the identified allosteric paths [26]. Experiments designate Thr87-Tyr106 coupling as the main reason behind allosteric effects in CheY [26]. Glu89 appears in all comparisons that contain 2B1J, except one. Indeed in FLiM bound state, Glu89 forms a hydrogen bond with Tyr106, and thus stabilizes the activated state of CheY [23]. Glu89 lies in several predicted allosteric pathways in CheY [25, 26]. Phosphorylation-induced conformational changes of *β*4 *− α*4 loop along with four residues Glu89, Trp58, Met85, and Tyr106 results in a buried conformation from solvent-exposed form [27]. Moreover, synchronised movement of both Glu89 and Tyr106 seems extremely relevant to FLiM binding to avoid steric clash with the latter. For the same reason, Lys119 moves farther away to accommodate the C-terminal end of the peptide. Met63, Lys119 and Arg22 appear whenever 1JBE is present in any pair. Met63 appears to change its rotameric state and comes in close contact with the adjacent helix. Close contact of the Met63 side chain with the Leu68 backbone pulls the loop harbouring the phosphorylated Asp57, which eventually allows space to Thr87 and Tyr106 to facilitate FLiM binding. Incidentally, the change in chemical shift for Met63 is observed to be high due to phosphorylation [28]. Though distally located, Arg22 rotates to the opposite direction upon FLiM binding. Change in hydrogen bonding pattern (from *β* strand-helix contact to intra-helical) is quite distinct in Arg22. This residue is also found to restrict the binding of Mg^2+^ with the structure by influencing the conformation of Asp35 [29]. Further, the synchronised movement of Arg22 and Arg19 (Figure S2(a)) abolishes polar contact between the two in FLiM bound state. Arg19 and Glu34 undergo minor conformational changes between CheY95IV and wild-type structures [30]. With subtle conformational differences, Glu34 seems to stabilise Arg22 in the unliganded state. A combination of experimental results identifies the key role of Ile95 in maintaining the surface complementarity for FLiM binding [30]. In the Ile95Val mutant, besides the Tyr106-Thr87 coupling, Ala88 and Met85 experience significant conformational alterations pertinent to allosteric signaling [27]. *β*4*−α*4 *loop* (Ala88-Lys91) is known to be responsible for allostery as well [31]. The adjacent Lys92 undergoes minor differences in respect to bond angles when compared between CheY structures solved at 1.1 and 1.7 resolution [23]. However, in the case of CheY-N16-FLiM complex, hydrogen bonding interactions between Lys92 and Asn16-FLiM make the complex stable [25]. Lys109 is an active site residue [24], which behaves as a conformational switch [32]. Phe30 and Phe124 sustain conformational changes, which is triggered by Asp57 due to phosphorylation of CheY [32, 33]. The autophospatase activity of CheY is accelerated manifold in the presence of CheZ. However, Lys26 mutation desists the dephosporylation activity of CheZ [34]. Phospho-CheY (CheY-P) plays a crucial role in chemotaxis, which helps E. Coli. to form swarms in agar. *A nonsense mutation* in Lys7 is observed to hinder the swarming ability of E. Coli [35]. In apostate, the side chain of Ser56 is represented as two distinct rotamers [30]. However, it is possible that phosphorylation at adjacent Asp57 perturbs the Ser56 side-chain to move away from Asp57. Glu27 is also observed to have double conformation in the Ile95Val mutant and its sidechain conformation shows noticeable difference with the wild-type structure [30]. Phe14 located in the vicinity of the phosphorylation site gets buried upon FLiM binding. Four Lysine residues - Lys92, Lys109, Lys119, and Lys126 - are confined to the surface where FLiM, CheZ, and CheA bind 36]. Met129 of *Salmonella enterica* CheY is observed to display high root mean square fluctuation (RMSF) in comparison to *Thermotoga maritima* CheY [37]. Glu93 is located at the solvent-accesible CheA binding surface of CheY [38]. In the activated state, the discrete movement of Glu93 lifts off the steric barrier, thus allowing space for FLiM binding.

### B. NolR

NolR is a global transcriptional regulator of genes involved in symbiosis and nodulation, which is necessary for nitrogen fixation. NolR belongs to the winged helixturn-helix (wHTH) superfamily. Barring a few exceptions, the recognition helix generally binds with the major groove of DNA, while wings comprising of *β*-turn-*β* bind with the minor groove. The comparison of 40MZ40N0 enables us to identify critical residues involved in the conformational switching between DNA-bound and unbound forms. Comparative analyses of crystal structures in (un)/liganded forms, thermodynamic analyses, and mutagenesis studies collectively point towards one particular residue, namely Gln56 [39] as the key player in the NolR mechanism. Conformational switching of this particular residue located at the recognition helix is necessary to accommodate variable target DNA sequences responsible for nodulation and symbiosis. Indeed, our method detects Gln56 as a key residue promoting interactions with nonpalindromic DNA sequences. Incidentally, 80% of the identified residues in both the polypeptide chains of the NolR dimer are located either at the recognition helix or the wing region (Figure S2(b)). Interestingly, our recent findings [40] reveal that 92% of the NolR residues - which exhibit the maximum change in eigenvector centrality - are concentrated in the recognition helix and the wing region. Perhaps this is worth mentioning that the eigenvector centrality measure of protein residue information is effective in identifying key amino acid residues involved in the allosteric signaling process [41]. Gln61 undergoes significant changes in its rotameric form to establish three new contacts with DC18, DT19 of one nucleic acid chain, and DT4 of another chain. Gln61 makes additional contact with conserved Ser57, facilitating bridging interactions between two DNA strands 39]. Similarly, Lys65 too suitably reorients itself to make electrostatic contact with DT17s phosphate group. The guanidium group of Arg67, located at the tip of the recognition helix makes a complete shift to interact with DT8 phosphate. Arg75 and Tyr82 from the wing stabilize the wing structure through hydrogen bonding interactions with the loop residues. Lys12 and Lys40 from different polypeptide chains seem to undergo drastic conformational changes between the two states. They likely have a role in the NolR dimer interface.

### C. Streptavidin

Considering its extraordinarily high affinity to biotin (vitamin H), the streptavidin-biotin complex has been hugely used to connect molecules, especially in immunoassays. Herein, we consider the PDB pair 1SWE and 1SWA representing holo-streptavidin and apo-streptavidin respectively to investigate streptavidin-biotin interaction dynamics. In each of the four streptavidin subunits, four Trp residues (at positions 79, 92, 108, and 120) are located at the biotin-binding site. The changes in intrinsic Trp emission fluorescence upon binding with biotin signify positive cooperativity [42]. A comparative study of quenching in Trp emission at 335nm (350nm) regions for hydrophilic (hydrophobic) surroundings identifies the above-mentioned four Trp residues undergoing direct interaction with biotin. Trp92, which is stabilized by stacking interaction from neighbouring Trp75 is one of the critical residues in Trp-tetrad. Replacement of Trp92 by Phe79 in another biotin binding protein Avidin results in altered specificity towards biotin [43]. The higher stability of holo-streptavidin compared to its apo-form has been largely attributed to the movement of Trp120 into the biotin-binding pocket of the neighboring subunit upon binding with biotin. The binding of biotin displaces water molecules from the active site. The eventual burial of biotin takes place through the ordering of the surface loop constituted of residues 45 to 50. Glu51, Ser52 and Arg53 located in subunit C adjacent to the surface loop are instrumental in the apoholo transition. These residues posess high PRI values. Indeed, a closer examination of this portion further reveals significant structural reorientation in this part of the surface loop (Figure S2(c)). All these residues are completely exposed to the solvent. Glu51C forms a salt bridge interaction with Arg84C in the holo-state, but not in the apo-structure. Conversely Arg53 is salt bridged with Glu44 (preceding the surface loop) in the apostate but not in the holo-state. Upon biotin binding the Arg53Glu44 salt bridge breaks and instead Arg53 gets completely exposed to the solvent. Selective destruction of one salt-bridge and construction of another salt-bridge at the other end of this three-residue stretch suggests the possible entry of biotin through the Glu51-Arg84 side, as shown in Figure 3(a). In agreement with the experimental data we also find the differential participation of each streptavidin subunit towards biotin binding. To illustrate, Tyr54 makes sidechain-backbone hydrogen bonding/polar interactions with both Glu51-Arg84 in chain C but only with Glu51N in chain A. In chains A and C, the role of Arg84 is found to be more significant for biotin binding. In chains B and D, the adjacent solvent exposed Tyr83 has enhanced significance. Tyr83 engages in polar interactions with neighboring surface loops, and likely plays a role in biotin entry in the respective subunits. Further, Glu116A (close to Trp120A) interacts with Arg103B (a *β* barrel residue) via a salt bridge interaction in the holo-form. This particular inter-subunit interaction is not observed between other subunits. Movement of Glu116A towards Arg103B leading to salt bridge formation facilitates biotin binding in subunit A. Despite minor conformational changes, larger changes in the selenoprotein binding to streptavidin is witnessed around Glu116 [44]. Lateral hydrogen bonding interaction between solvent-exposed Tyr22 and Thr20 stabilizes contact regions of higher-order biotinylated streptavidin structures [45]. Val133 also likely has a stabilizing role in the contact regions. Phe130 along with the neighbouring Trp21 from the *β* barrel provides the necessary hydrophobicity in the biotin-binding pocket.

**FIG. 3.**
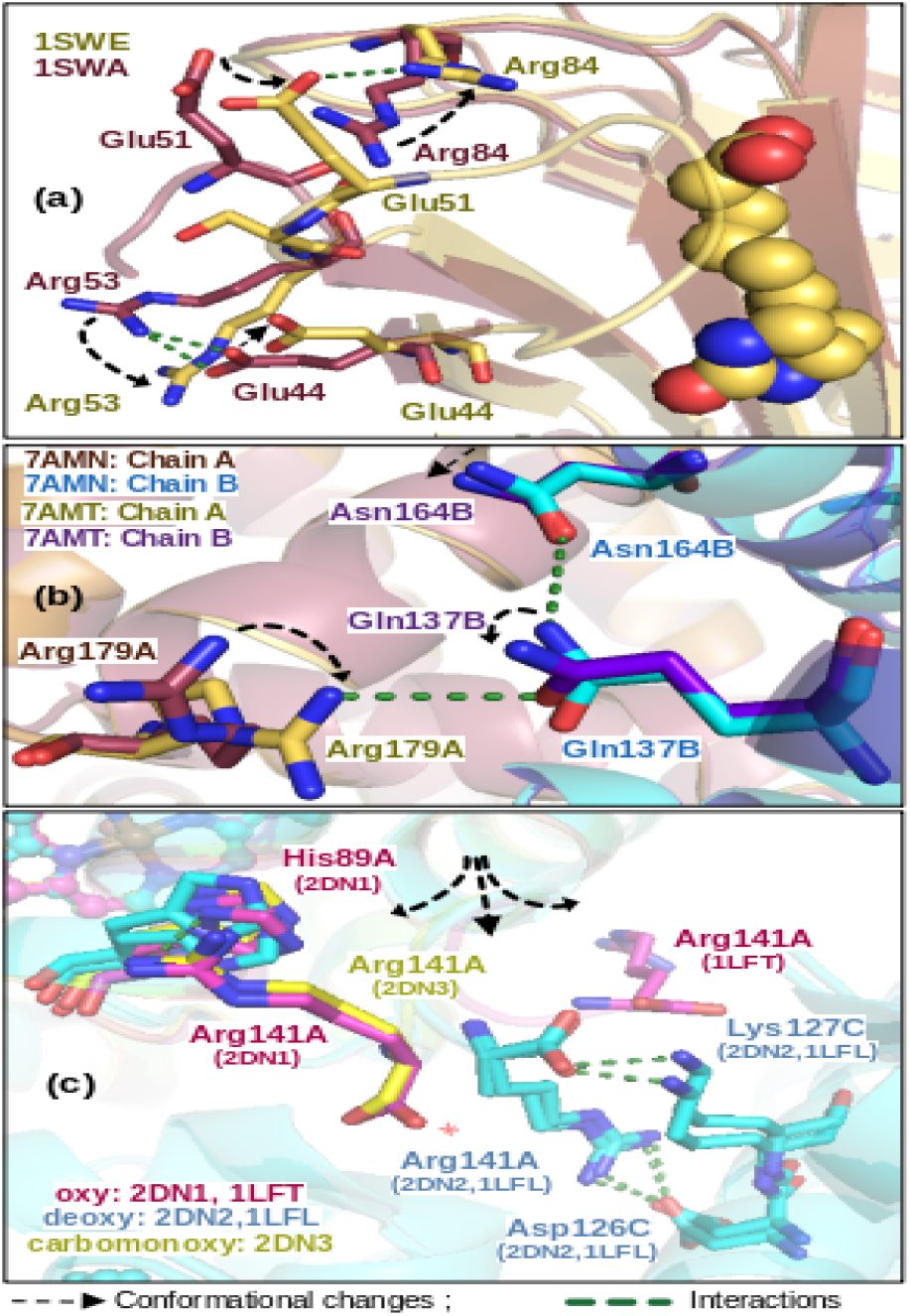
Formation and destruction of various intra-chain, inter-chain, and inter-subunit interactions in distinct states of: (a) streptavidin, (b) LuxR, and (c) haemoglobin due to conformational changes of residues. (a) Salt-bridge interactions exist between Arg53 and Glu44 in 1SWA. The highlighted contacts break down during the holo-apo (1SWA1SWE) transition, forming a novel salt-bridge interaction between Glu51 and Arg84 in 1SWE. (b) In 7AMN, there is an intra-chain hydrogen bonding between Asn164B and Gln137B. However, in 7AMT, that interaction is absent and a new inter-chain hydrogen bonding is developed between Gln137B and Arg179A. (c) Inter-subunit hydrogen bonding interactions mediated by Arg141 break down while transitioning from deoxy to ligand-bound states.

### D. Aureochrome

In the 5A8B-5DKL pair, we investigate the influence of blue light on the conformational dynamics of the signaling aureochrome photoreceptor. Aureochromes [46] are a unique group of blue-light responsive basic leucine zippers (bZIPs), which regulate DNA binding activity upon the reception of light at the light-oxygen-voltage (LOV) domain. Aureo1-LOV from *P. tricornatum* (*Pt*Aureo1 LOV) displays the signature five stranded antiparallel *β* sheet harboring flavin co-factor, flanked and interspersed with *α* helices. Packing/unpacking of both the flanking 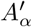 and *J*_*α*_ helices play a critical role in the conformational reorientation of the dimerization interface upon light induction. The simultaneous unfolding of 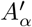, and *J*_*α*_ helices is necessary for light-induced dimerization and transfer of signal to the adjacent bZIP domain for DNA binding activity [47]. The distribution of residues identified from both the polypeptide chains of dimeric *Pt*Aureo1 LOV is shown in Figure S2(d) of SI. Interestingly, none of them is located in the flavinbinding pocket, barring Phe271. Phe271 is proximate to FMN and is fully conserved not just across Aureochromes but throughout the LOV photoreceptor family. Phe46 in *Bacillus sp*. LOV photoreceptor YtvA is homologous to Phe271 in PtAureo1 and undergoes light-induced conformational flip. Mutation of this residue to His significantly accelerates dark recovery kinetics. Conformational flexibility of this residue is known to be important for the dark-light transition [48]. Apart from Phe271, the residues identified herein are dispersed across the LOV structure: (a) featuring mostly on the outer side of *β* sheet lining dimer interface, (b) on the peptide linkers connecting secondary structure motifs, and (c) on both the terminal helices. Recent studies confirm the role of peptide linkers and *C*_*α*_ helix mediating conformational changes associated with light-induced allosteric signaling [49]. Non-polar interactions between the back of 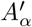 helix

and the *β* core is crucial to maintain the dark state structure. Following illumination by blue light, the unfolding of 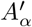 helix exposes the hydrophobic surface of the *β* core, resulting in reorientation of the dimer interface and thereby affecting signal transduction. Phe241 from 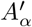 helix, which packs against *β* strands [50], interacts with Phe332 of the *β* core in the native dark state. Upon illumination by blue light, the side chain of Phe332 rotates in the opposite direction, breaks its contact with Phe241 and promotes light-induced conformational changes. The phenol ring of Tyr266 also rotates completely in the opposite direction thereby promoting light-induced reorientation of the dimer interface. Again the Tyr277-Tyr320 pair located at the peptide linkers adjacent to *C*_*α*_ and *G*_*β*_ respectively, reorients subtly to make a new polar contact in the light state. This minor shift in distance from 4.4Å to 3.6Å seemingly contributes towards the compaction of the LOV core following the departure of 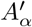 helix in the light state. A similar effect is noticed in Trp328, which is located between *J*_*α*_ and *G*_*β*_. Though the polar contact of Trp328 with Asp355 (at *J*_*α*_) is retained in the light state, the indole ring of Trp328 deviates from planarity and moves farther from *J*_*α*_. This results in a more compact *β* core. It is well-known that Asn319 and Asn329 interact directly with the isoalloxazine ring of the flavin chromophore [51] and are conserved components of the robust hydrogen bonding network needed for LOV photochemistry. Remarkably, our studies find the residues adjacent to Asn319 and Asn329, namely Tyr320 and Trp328 as shown in Figure S2(d) of SI. Due to simultaneous unidirectional shifts in the Tyr357-Gln330 pair, hydrogen bonding is retained in between the N-terminal part of *J*_*α*_ and the *β* core, thereby supporting the partial unfolding of *J*_*α*_ in the light state [51]. Tyr357 further positions itself at the juncture of *J*_*α*_ and 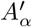 in the light-induced dimer interface due to structural reorientation. Gln366 located at the C-terminal tip of *J*_*α*_ shows a drastic change in conformation between dark-light states. This attests to the unfolding of *J*_*α*_, which induces light-dependent signaling.

### E. LuxR

LuxR is structurally similar to the Tet repressor and belongs to the helix-turn-helix (HTH) group of transcription factors. LuxR is responsible for the activation/repression of hundreds of genes, giving rise to the emergence of quorum sensing in *Vibrio sp*. Therefore, understanding the difference in relative positions of amino acids in LuxR-DNA structures during activation (7AMT) and repression (7AMN), sheds insight into the underlying molecular mechanism leading to such collective behaviorial response. LuxR binds with DNA as a homodimer in both states [52]. A quick PDBePISA [53] analysis interestingly reveals that, while transitioning from activation to repression there is a gross change in proteinprotein and protein-DNA contact surface. While the protein-protein interface area expands in activation, the

protein-DNA interface contracts. The opposite happens during repression. Ile199 is one such residue that makes an additional hydrogen bonding interaction with Met196 only in the activation state but not in the repression state. Interchain hydrogen bonding interactions between Asnl64A and Tyrl93B differ slightly between activation and repression state. Asnl64A-Glnl76 interaction replaces Asnl64A-Tyrl93B interaction in the latter state. In the activation state, Asn164 from chain B makes a single hydrogen bonding contact with Tyr193A, whereas Asn164B makes additional interaction with Glnl76A besides interacting with Tyr193A in the repression state. Interchain salt bridge interactions involving Argl22 and Glull6 is conserved in both activation and repression state. The chain-wise dynamics of Glnl37A (Glnl37B), Asnl64A (Asnl64B) and Argl79B (Argl79A) triad are exciting. From Figure 3(b), we observe that while transitioning from repression to activation state, Gln137A moves subtly to form hydrogen bonding interactions with Asn164A, moving a little farther from Arg179B in the process. However, Gln137B breaks the hydrogen bonding interaction by moving farther from Asn164B, only to create a new hydrogen bonding interaction with Arg179A, as shown in Fig. 3(b). It is indeed interesting to witness antagonistic inter-polypeptide chain interactions as observed in LuxR, which is well-detected by our method. Arg36 and Tyr56 from both A and chain Bs are engaged in hydrogen bonding interactions with both strands of DNA, besides contributing to the protein-protein interaction interface as well. Phe54 demonstrates van der Waals interactions and hydrophobic contacts with Tl4, Al5, and Tl6 of LuxR-DNA repression complex [52]. During repression, the OG1 atom of Thr59 is at a distance amenable to bonding from the OEl of Glu6l in contact with the DNA backbone. However, during the activation state this particular Glu61 shifts rather drastically by 4.2Å from the DNA backbone. Further, Thr59 instead of interacting with Glu51, makes polar contact with Asp62, when both of these are at favorable distance from the DNA backbone. Similarly, Arg60 connects with Glu124 in the activation state, whereas during repression, it moves farther away not only from Glu124 but also from the major DNA binding residue Arg36. In fact, DNA binding to LuxR promotes electrostatic interactions in the amino acid tetrad Arg60-Glul24-Argl22-Glull6. Vhile the role of Phe57B has not yet been ascertained precisely, its location at the end of *α*_3_ helix could be important in the context of the neighboring DNA binding residues and the critical dimer interface. The same applies to Phe78B, located distally towards the end of *α*_4_ helix. Phe78 also seems pertinent for maintaining hydrophobicity in the surrounding region. Hydrophobicity in this region has been helpful in designing future quorum sensing inhibitors 54j. Ile93, Glul02, and Glnl06 are all located in the *α*_5_ helix of the B subunit. In each case, their side chains move in the opposite direction as part of the overall conformational changes associated with interstate transition. The movement of Ile93B occurs in concert with the movement of Metl43B and Phel44B, with both located at the subsequent *α*_6_ helix (Figure S3(a)). Asp65 maintains hydrogen bonding contact with Asn69 in both states. The carbonyl oxygen of Leu87, which is a neighbor to the carbonyl oxygen of Leu85 in the activation model, moves away by *≈* 1Å in the repression state. Arg134 moves closer to interact with Ser131 in the repression state but not in the activation state. The most prominent change of Glu152A is the hydrogen bonding interaction between its side chain oxygen with neighboring His88 imidazole nitrogen in the repression state. Interestingly, in the activation state, the His88 side chain rotates by *≈* 90^*°*^, and extends further to lose interaction with Glul52. Remarkably, His88 in chain B remains largely unaffected during the inter-state transition. Asymmetric conformational changes on both the LuxR subunits seem to be the hallmark of its interstate transition. This is further validated by selective rotameric changes in Glu23B, which is not observed in Glu23A. Further, this observation is remarkable in the sense that to switch between activation and repression, even the most distal amino acid residues from the DNA binding region undergo subtle conformational changes and that too asymmetrically.

### F. Tbx5

Tbx5 belongs to the *T* -box transcription factor family and plays a pivotal role in heart and limb development. Several mutations in Tbx5 have been associated with an autosomal disorder called Halt-Oram syndrome. In the 2X6U-2X6V pair [55], we compare the DNA binding domain (DBD, amino acid residue stretch 58-238) of Tbx5 in DNA bound/unbound states and identify twelve residues spanning across the DBD. Among the residues found important in our study, Arg81 and Asn230 interact with DNA. In the DNA bound state (2X6V), the Arg81 side chain extends to establish contact with the phosphate oxygen of DG l8 (Figure S3(b)). In the DNA-bound state, Asn230 flips by 180^*°*^ not only to place ND2 atom appropriately near the phosphate oxygen of DT l7, but also to place its non-polar backbone at the vicinity of Phe84 from the neighbouring loop. Glnl56 and Lysl57, both parts of nuclear localization signal (NLS) undergo significant conformational changes during transition to the DNA-bound state. Lysll5 exhibits a complete flipping in the DNA-bound state, and belongs to the lid region comprising of *β*-turn-*β*. Being located adjacent to DNA binding site, the structural integrity of the lid is extremely important. Glul28 succeeding the lid interacts with Argll3 from the lid in the DNA-free state. However, this polar contact ceases to exist in the DNA-bound state, indicating necessary structural rearrangement in the Tbx5 structure preceding DNA binding. A similar rearrangement is seen for His220. The imidazole chain of His220 flips completely to the opposite direction only to loose contact with another lid-residue Asnll9 in the DNA bound state. Phe57, Tyr100, Arg150 and Gln180 seem to contribute towards structural stability at the region distal from DNA binding site. Another residue with a high PRI value, His170, is crucial for structural stability, as elaborated in the S.I.

### G. Hemoglobin

In “polyfunctional” hemoglobin (Hb), we choose the most vital function, namely oxygen transport. The existence of multiple signaling pathways leading to Hb’s phenomenal allostery has been confirmed by: (a) multiple ligation intermediates with distinct conformational states and oxygen affinities, and, (b) determination of many structural intermediates through static and dynamic crystallography. In this regard, we compare various PDB pairs 1LFL vs 1LFT (deoxy vs oxy), 2DN1 vs 2DN2 (oxy vs deoxy), 2DN2 vs 2DN3 (deoxy vs carbomonoxy) and 2DN1 vs 2DN3 (oxy vs carbomonoxy). Comparisons of structure between unliganded tensed state *(T* ) with liganded relaxed state *(R)* against low and high oxygen affinity respectively provide molecular insights into the cooperative effects in Hb tetramer. Cooperativity among the four subunits of *α*_2_*β*_2_ tetramer is essential for oxygen uptake and *in vivo* release. Cooperativity is also responsible for shifting the allosteric equilibrium from the *T -*state to *R-*state [56]. Structurefunction studies supported by computational, thermodynamic, and spectroscopic data confirm the presence of multi-state structures besides the classical *T* and *R* endstates, namely *R*2, *RR*2, *R*3 *and RR*3 [56]. As described in the S.I., we identify the residues located across *αβ* subunit of Hb. Among these identified residues, Asn102B is crucial as the Asp94A-Asn102B hydrogen bond in the *R* and *R2* states is absent in the *T* -state. His97B undergoes marked differences in its conformation during the *T → R* transition. In fact, the change in interactions mediated by His97B in both *T* and *R* states is considered as a maJor steric barrier to the quaternary structure change. However, the *R-R*2 screw axis passing through the Trp37B-Arg92A intersection induces significant conformational changes in His97B, which is different from the *T → R* switching movement [57]. In the *T* -state, the C-terminal residue Arg141A interacts electrostatically across α subunits and maintains *T* -state stability. Figure 3(c) shows that in the *R*-state, Arg141A flips completely to lose contact with His89A of E-helix. Arg141α, through inter-subunit hydrogen bonding interactions with Asp126α and Lys127α, stabilises the deoxy state and facilitates oxygen release. [56]. Further in the C-terminal region of the T -state, His146B engages in two inter-, and intrasubunit hydrogen bonding interactions. However, given its high solvation and disorderliness, His146 from both the β subunits move drastically to stack against each other (Figure S3(c)) and establish a new salt bridge connection with Lys82B. Lys82B facilitates the *T → R* transition by contributing to Bohr effect via deoxygenation-linked proton binding. Lys82 being a positively charged residue repels other positively charged residues at the central water cavity and increases the free energy of Hb [56]. Lys82 along with His2, Asn139 and His143 are *β* cleft residues. Interestingly, 2,3 bisphosphoglycerate and its analogues interact with these *β* cleft residues via water-mediated/direct hydrogen bonding interactions, stabilising *T* -state structure with low oxygen affinity [56]. In fact in Hb Hinsdale (a variant of human Hb), Asn139 → Lys mutation introduces additional positive charge in the central water filled cavity, destabilises *T* -state structure due to electrostatic repulsion and increases oxygen affinity. In another Hb variant (Hb Doha), Val1 → Glu substitution in the *β* subunit further promotes the *T* -state structure by preventing the cleavage of the initiator acetylated methionine. This causes structural changes with the formation of intra/inter-subunit contacts especially at the *β*1*β*2 interface, resulting in a stable *T* -state structure [56]. Further, mutations at the *α*1*β*1 *or α*2*β*2 interface can cause significant changes in the oxygen binding properties of Hb. For example,Tyr35 → Phe mutation at the β subunit results in increased oxygen affinity and decreased cooperativity in Hb Philly variant. Again, Phe42 deletion causes abrupt shift in allosteric equilibrium, resulting in reduced oxygen affinity and low cooperativity in ligand binding as found in Hb Bruxelles [58]. Our method detects Thr38 as one of the key residues. Interestingly, Thr38 is located Just one helix turn away from Tyr35 and Phe42, and favourably contacts the vinyl group of heme cofactor via its non-polar end of side chain. Further, mutations in the hinge region (e.g. Tyr35 → Arg) of *α*1*β*2 perturbs the dimer interface, causing variable affinities towards oxygen [56]. Salt-bridge interactions, especially the ones involving the C-terminal residues of Hb (e.g. Asn139, and Arg104 interaction) play a crucial role in the cooperativity mechanism of HbO_2_ [59]. His146 forms two salt bridges in *T* -state. However, in the *R*-state, these salt bridges are removed and His146 gets solvated and disordered [57]. In deoxyhemoglobin, His146 gets delocalised and does not participate in salt-bridge formation [59]. His146 also promotes *T → R* transition. Tyr24A and Trp15B were observed to take part in nitration of human hemoglobin by H_2_O_2_/nitrite [60]. The formation of 4Hydroxybenzyl adduct at Tyr24A in Hb indicates high solvent accessibility [61]. Arg31 from the B-helix engages in interchain hydrogen bonding interactions with Phe122 backbone and Glu27 side chain respectively from G and B-helices. This facilitates the docking of *Staphylococcus aureus* IsdB onto αHb showing the former’s importance in organic iron acquisition [62]. Lys11A-Ala168B contact is the most conserved interaction at Isdb^N1^ : Hb interface [62]. Lys60A also contributes a hydrogen bond at the Isdb^N1^ : Hb interface [62]. Conserved His72A from the EF corner along with the other conserved histidines (89, 97, 103 and 112) are protonated during the transition from R to T -state [63], with distinct role in the Bohr effect. His143B along with His20A, His112A and His77B

are identified to play a role in countering alkaline Bohr effect. His77B is observed to possess different conformations in different structures [64]. Another conserved histidine involved in Bohr effect (Hisll7B), however gets deprotonated while transitioning from *R* to *T* -state [63]. In *R*-state structure, His117B, and Glu22B engage in weak electrostatic interaction. However, in *T* -state, they disengage and drift apart by *≈* 8 Å. Carboxymethylation and hydroimidazolone modification are observed on Arg30B in human Hb l65j. Phe7lB from the E-helix is located at the frontiers between internal cavities of goblin fold and also known as a mechanically sensitive residue [66]. Research on pyrrole based alkaloids is important not just for their structural and conformational similarities with porphyrin moieties but also for the therapeutic purposes like lowering of blood cholesterol. Docking analysis involving Hb and a pyrrole compound, PyS, reveals the significance of Trpl4A in facilitating *π*-*π* interaction between Hb and ligand [67]. T-R structure comparison reveals complete flipping of indole side chain of Trp14A. Sodium dithionite is popularly used as deoxygenation agent of Hb and Mb. Crystal structure reveals the interaction of dithionite molecule with Hb via Hisll6B, Hisll7B and Lysl6A residues, thereby promoting deoxygenated state [68]. Thr67 is located at the channel connecting heme iron binding site of *α* subunit of Hb, and the exterior. Substitution of Thr67 with more obstructing but more mobile Lys suggests similar CO exit path in Hb and Mb [69]. Location of Val70A at the channel connecting the distal site and solvent suggests its role in ligand migration [70]. The heme pocket is generally conserved between CO-Hb and CO-Mb. NMR studies reveal conserved hydrogen bonding interaction between the backbone carbonyl of Leu83A and His87 at the proximal site of CO-Hb. This validates an extremely conserved coordination geometry and similar hydrogen bonding patterns between these two proteins [71].

### H. Myoglobin

Effective discrimination between oxygen and carbon monoxide by Hb and Mb is a prime factor to aerobic respiration [72]. Therefore, besides Hb we also focus on Mb. We primarily compare five experimental structures of Mb: lMDN V68N de-oxy Mb; 1MNO - V68N oxyMb; lM6C - V68N Mb with CO; 1MWD - wild type de-oxy Mb; 1MWC - wild-type Mb with CO (see S.I.). While comparing the aquomet, unliganded, CO and *O*_2_bound Mb structures, differences in electron density are mostly observed at the ligand binding site as well as the hydration network surrounding the solvent side of distal His (His64). Mutation of His64 to Leu forbids the system from binding with *O*_2_ [73]. In the CO-bound form, His64 swings out of the distal heme pocket while the latter gets occupied by two ordered *H*_2_*O* molecules. The movement of His64 is in harmony with that of Lys45, which borders the polar heme surface. Lys45 along with His64 and Val68 borders the entry/exit path of diatomic ligand [74]. Further, the mutation of Lys45 by Arg lowers the inner barrier for CO rebinding - as apparent in our findings. After getting dislodged from the CObinding pocket of Mb, the imidazole side chain of His64 establishes hydrogen bonding/electrostatic contact with heme as well as Asp60. Justifiably, His64 and Asp60 does not feature when wild type Mb with CO bound/unbound forms are compared. Structural superposition of lMDN, lM6C and 1MWC in Figure S3(d) shows insignificant movement of His64 in wild type Mb’s CO-bound structure. This is contrary to that of the CO-bound form in V68N variant. Again, the side chain of Glu83 flips completely to the opposite direction in CO-bound WT Mb forging new interactions with residues from EF corners. This shift does not take place in the V68N variant, where Glu83 maintains polar contacts with residues from F-helix harbouring proximal Histidine. Upon dissociation of CO, the subtle movement of Lys145 at the H helix by 0.6Å ruptures hydrogen bonding interaction with Thr87 from the F helix. Comparison between unliganded and CO-bound Mb reveals shift of FG corner residue Lys98 towards Lys42 from C helix. Gln8 engages with Lys79 at the EF corner and is found to be important for transiting between ligand bound/unbound conformations. An A helix residue, Lys16 drifts its side chain significantly towards GH corner in *O*_2_ bound state (see S.I.). Glu41 from C, Glu83 from EF, Lys96 and Lys98 from FG and Lys145 from H, which exhibit strong dynamic interaction with the heme group in deoxy-Mb are found to be important for unliganded to liganded state. In fact, Lysl45 along with Tyrl46 are central mediators of dynamic interactions [75] at the C-terminal segment of the F to H helix. Glu41 from C helix and Lys47 from CD corner, pair up via a salt bridge interaction. However, upon switching to ligand bound states (*CO/O*_2_), the salt bridge ruptures. Notably, in Mb structures, several residues including Gln8, Glu41, Lys47, Glul09, Serll7 and Glnl28 have double conformers in one/more state(s). One Glu109 conformer salt bridges with His36, which has greater potential to engage in stacking interactions with Phel06 at the ligand (*CO/O*_2_) bound states. Ser117, at the end of G-helix is completely solvent exposed. Therefore, it would have a role in influencing the hydration network of Mb. Gln128 engages in intense hydrogen bonding interactions with residues from G-helix (Gln116 side chain), GH corner (Phel23 backbone) as well as the H-helix (Ser132 side chain). Together, these constitute the G-GH-H hydrogen bonding network. While several residues are common in information transfer from ligand unbound to bound state, subtle change in single amino acid residue apparently activates additional amino acid centres. This is evident not only from our findings but also from altered *CO/O*_2_ binding affinity between WT and the V68N Mb variant.

### I. MutY

Our next target is the adenine DNA glycosylase, MutY. It functions in the base excision repair (BER) pathway and protects our DNA from potentially genotoxic 8-oxoguanine (8-OG). MutY initiates DNA repair by removing adenine base opposite 8-OG and generates an abasic site (see S.I.). Herein, we compare two structures from PDB: 1RRQ (with damaged DNA bearing 8-OG) and 1RRS (with DNA containing an abasic site). This means that a comparison between these two structures should shed insight into the precise enzymatic action of MutY in the first step of BER. As detailed in the S.I., the amide bearing side chain of Gln48, a residue with high PRI value, interacts with the neighbouring 3^′^ base as well as hydrogen bonding interactions with the phosphate backbone [76]. The Watson-Crick face of 8OG is further stabilised by the backbone amide carbonyl of Gln48 as well as the hydroxyl side chain of Thr49, both interconnected via hydrogen bonding interactions. Complete flipping of the Glu223 side chain from the exposed to the buried state happens simultaneously with a major shift in the DNA strand at DC’16, DA’17, adjacent to the abasic site (DA’18). Glu59 side chain rotates by ≈ 90° to be in close contact with *Ile55/0*. The backbone carbonyl of Leu86 further stabilises OG at the minor groove via hydrogen bonding interactions. While the indole nitrogen of Trp30 is close to the amine group of flipped adenine base, the bulky hydrophobic side chain of Trp30 is part of several other hydrophobic amino acids lying in close contact with the adenine base. Trp20, located at the iron-sulfur cluster containing N-terminal catalytic domain seems to be important for structural integrity of the cluster. Through hydrophobic interactions, Trp20 stabilises Val197, located next to Fe-S coordinating Cys198. Maintaining the functional integrity of FeS cluster is extremely important because change in oxidation state between [4*Fe* − 4*S*]^2+^ and [4*Fe −* 4*S*]^3+^ results in the donation of an electron to the damaged site. In absence of FeS cluster, MutY is unable to bind to the DNA, and thus unable to repair OG lesion. [4*Fe ™* 4*S*] cluster is connected to the polypeptide chain via four cysteine residues - 198, 205, 208 and 214. Inversion of *CONH*_2_ side chain of Gln211 is also important to maintain contact with Cys205 in the repaired state. The backbone carbonyl of Gln211 interacts with the backbone amide of Cys214 via hydrogen bonding in both states, thereby signifying its role in the FeS cluster domain. This is in agreement with our formulation that even the residues with subtle to almost no conformational changes during transition plays important role in protein function. Other such residues are Trp83 and Trp266. With no visible conformational changes, Trp83 remains connected to the backbone carbonyl of Gly87 in both states (Figure S4(a)). Maintaining proper conformation in this region is absolutely critical not only to stabilise the Hoogsteen face of OG through Tyr88-OH located next to Gly87 but also to facilitate catalytic process mediated by Tyr88-water-Gln48. While the negatively charged phosphate of OG is stabilised by the backbone amide of Leu262, the hydrophobic side chain of the latter is further stabilised by the non-polar part of the indole of Trp312. Trp266, with its bulky non-polar side chain, pushes Arg332 to move closer to static Tyr342-OH to make additional hydrogen bonding interactions in the repaired state. Loss of contact with the backbone of neighbouring Tyr331 induces the Val327 side chain to rotate in the direction opposite to the repaired state. This results in a subtle widening of the loop. Appearance of Leu282 could be an artefact given the loss of electron density connecting Gly281 and Leu282 in both states. The side chain of Leu71 rotates only to make a new non-polar contact with the carbon atom of imizadole ring in His96, located at the neighbouring helix at the repaired state. Further, the imidazole nitrogen of His96 gets connected to Glu76 in the same state, located in the adjacent helix of Leu71. Non-covalent interactions among Leu71-His96-Glu76 triad that form only in the repaired state, bring neighbouring helices close to each other, resulting in gross conformational changes. Similarly, two beta strands move closer to each other through newly developed interactions with Leu244 in the repaired state. The importance of Glu101 is not readily understandable, especially given the partially unresolved neighbouring Arg105 in the damaged state. In the repaired state, Glu101 makes a salt bridge interaction with Arg105. A shift of the Asn146 side chain in harmony with the formation of an abasic site is quite remarkable. As soon as the abasic site is generated, the amide side chain in Asn146 rotates to stabilise dislocated sugarphosphate backbone of DA 18. Trp352 seems to maintain local hydrophobicity opposite to the DNA binding face especially by non-polar interaction with Leu269 and *π*-*π* interaction with neighbouring Phe268. In contrast, on the DNA facing side, the exposed polar side chain of Arg353 along with Arg350, Glu357 etc. creates necessary polarity for solvent exposure and DNA binding.

### J. KcsA

Potassium ion (*K*^+^) channels facilitate the selective transport of *K*^+^ ions from an intracellular to extracellular environment at an extraordinarily high speed. This is essential for maintaining *K*^+^/*Na*^+^ balance in living systems, thus sustaining homeostasis. Herein, we investigate the bacterial *K*^+^ channel KcsA in two different states: (a) conductive or open state, and, (b) relaxed or closed state. For a better understanding, we compare three PDB pairs, albeit in a single polypeptide format (i.e. chain C vs chain C in PDBs of the two states, due to the lack of tetramer data). The pairs of PDBs considered here are: 6W0F-6W0D, 7MHR-7MK6 and 1K4C1K4D. They all represent S5-S6 segment of KcsA *K*^+^ channel from *Streptomyces lividans*, solved in open and closed state conformations, crystallized under different salt conditions. Incidentally, all of these structure-pairs are solved in an identical space group, thus suitable for structure comparisons. From Table I, we observe that Trp68, Arg89 and Phe114 are a constant presence in all cases, while Tyr78, Trp87, Trp113 are common in the first two pairs. All the residues Arg64, Trp67, Trp68, Ser69 - flank the conserved filter region which is responsible for regulating selective yet high-speed movement of *K*^+^ ions across the membrane (see S.I.). Though not in WT, Trp67 in KcsA E71V mutant structure adopts a different rotamer conformation - this justifies the detection of Trp67 only in 7MHR-7MK6 structure-pair (Figure S4(b)). A 180^*°*^ rotation of the indole moiety of Trp67 in 7MHR-7MK6 abolishes hydrogen bonding interaction with the neighboring Asp80 upon gate opening. Trp87 is located at the N-terminal end of S6 (see S.I.) and shows significant movement in its side chain upon the opening or closing of the KcsA channel. A comparison of structure across the open and closed state KcsA reveals translation and counter-clockwise rotation of the inner helices (S5, and S6) with respect to the centre of symmetry of the channel. While Trp26 and Ala28 are present at the N-terminal end of S5, Leu35 showing signs of conformational mobility is located around the middle of S5. Tyr45 towards the C-terminal end of S5 shows a nearly perpendicular shift in its phenolate ring - which seems to have a role towards interaction between the pore helix and the adjacent voltage sensor related to channel opening. Fully conserved Tyr78 is a part of the signature sequence of the selectivity filter. The backbone carbonyl oxygen of Tyr78 participates in *K*^+^ coordination, whereas the protein core-facing side chain provides the necessary geometric constraints to maintain the geometry of *K*^+^ binding sites. While Trp87 faces the inner side of the protein core, the polar side chain of Arg89 faces the intrasubunit space at the extracellular side. Located in the vicinity of the selectivity filter, Arg89 is connected with the tetramer stability of KcsA [77]. Quite interestingly, the conserved residue Gly99 associated with opening and closing of the gate is the neighbouring residue located at the adjacent inner helix, S6. Considering the direct dependence of *K*+ ion conduction and bending of S6 at Gly99 leading to opening of the gate, contribution from the neigbouring residues appear significant. Trpll3, Phell4, Glull8, Glnll9, Glul20, Argl2l and Hisl24 - are all present in the C-terminal half of the S6 helix (see S.I.). While the non-polar side chain confers structural stability due to its facing the hydrophobic side, the polar side chains provide a more favourable environment for polar *K*^+^ ions and water at the intracellular side. Existence of polar amino acid residues at the end of S6 justifies easy solvent access even in the middle of the transmembrane portion of the channel. A significantly long aqueous passageway followed by a short selectivity filter is the hallmark of KcsA *K*^+^ channel [78]. This explains the very high conductance rate of *K*^+^ ions through KcsA channels.

In summary, we identify and discuss the functional significance of residues with higher values of PRI. Not infrequently, we find that the residues neighbouring an individual residue rather than the residue in question rearrange themselves to assist a specific biological activity. We also predict the likely functional significance of other residues identified by our approach, which are amenable to experimental validation.

## IV. CONCLUSION

Substantial correlations exist between the structure and function of proteins. The position and conformation of amino acid residues may be altered due to diverse factors. Conformational changes between two states of a protein arise from alterations in position and/or conformation of every residue with: (A) respect to itself, (B) its neighbouring residues, and, (C) indeed all other residues. Common visualisation software enable us to visualise structural alignment of the types A and B. However, all these conformational changes (types A, B, and C) are important as they lead to the construction and destruction of various non-covalent interactions, which influence the overall functioning of a protein. Herein, we introduce a fresh *ab initio* information-theoretic approach to study proteins and protein-ligand interactions, which accommodates all the aforementioned types of structural changes (A, B, and C).

We formulate a method to calculate “inter-state” entropy, 𝒮 _*k*_, and “intra-state” entropy, 𝒥 _*k*_. 𝒮 _*k*_ depends on the Euclidean distance between pairs of atoms and is calculated *for any given state*. 𝒥 _*k*_ depends on the *diference* in Euclidean distance between pairs of atoms and is calculated *across a pair of states*. We also propose a *de novo* measure called protein residue information (PRI), using _*𝒮 k*_ and 𝒥 _*k*_. The utility of this approach is readily visible as we employ PRI to analyze twenty eight distinct pairs of protein structures drawn from ten different classes. PRI is largely successful in pinpointing residues of structural and functional significance. Remarkably, it successfully identifies important residues, irrespective of the magnitude of conformational changes undergone by them. Besides revalidating functionally important residues cited in the literature, we furnish mechanistic and functional insights for the rest of the residues from the 3*D* structures. The identified residues are found to be functionally important in allostery, structural stability and proteinligand interactions. MD simulations and network-based techniques are powerful and established techniques. The present approach even with any attendant handicap fares better in terms of lower computation time and circumvention of cutoff distances when compared to MD simulations and network-based techniques respectively. Therefore, this formulation can serve as a fresh alternative theoretical approach to detect the contribution and importance of amino-acid residues in protein function.

## Notes

### Competing Interest Statement

The authors have declared no competing interest.

### Summary of Updates

Subsections introduced; Equations formatted; Supporting information updated.

