## Supporting Information for "An *ab initio* information-theoretic approach to proteins and protein-ligand interactions"

Using the formulation proposed herein, we can compute the intra-state entropy, $\mathcal{S}_k$, and inter-state entropy, $\mathcal{J}_k$, for every residue, $k$. The value of intra-state entropy can be computed for each state of a protein. On the other hand, the value of inter-state entropy can be calculated for any given pair of states. We then calculate the value of protein residue information, $\mathcal{M}_k$, of any individual residue, $k$, of a protein, whose structure is known. Herein, we analyse twenty eight distinct pairs of proteins drawn from ten different classes. Every pair incorporates two different states of the same protein.

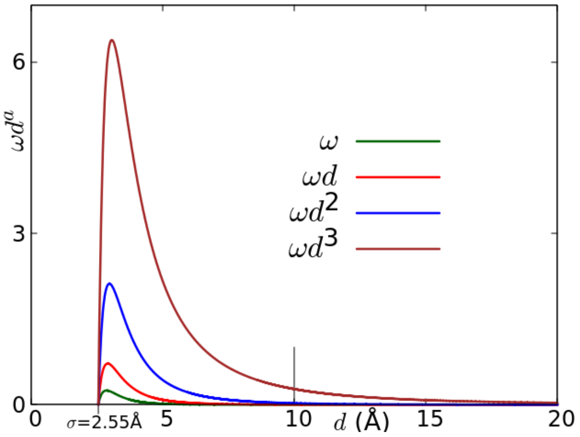

**Figure S1:** $\omega d^a$ *versus* $d$ for $a = 0, 1, 2, 3$. Here, $\omega(d) = [(\frac{\sigma}{d})^6 - (\frac{\sigma}{d})^{12}]$. Upon averaging over more than 11600 atoms of the twenty-eight pairs of distinct proteins analysed by us, we obtain $\sigma = 2.55 \pm 0.0069$ Å. We have independently verified that this functional variation is quite robust and the average $\sigma$ for any statistically significant sample of proteins in PDB does not differ much from $\sigma = 2.55$. The vast majority of non-covalent interactions are active up to $\approx 4$Å in proteins. The remaining interactions in proteins (including long-range electrostatic interactions) are far weaker beyond $\approx 4$Å and possess considerably lower relevance up to a maximum of $\approx 10$Å marked by the vertical line.

| | $\Delta A_1/\Delta A_3$ (%) | $\Delta A_2/\Delta A_3$ (%) |
|---|---|---|
| $\omega$ | 93.6 | 6.2 |
| $\omega d$ | 88.7 | 10.7 |
| $\omega d^2$ | 80.2 | 17.6 |

**Table SI:** The fraction of area in Fig. S1 for $\omega$, $\omega d$, and $\omega d^2$ at various intervals of distance, $d$. $\Delta A_1$, $\Delta A_2$, and $\Delta A_3$ denote the area for $0 \leq d \leq 5$Å, $5 < d \leq 10$Å, and $0 \leq d \leq 20$Å respectively. These areas are a signature of the number of interactions within the respective regions of distance. Thus, $\Delta A_1/\Delta A_3$ and $\Delta A_2/\Delta A_3$ signify interactions within $0 \leq d \leq 5$Å and $5 < d \leq 10$Å respectively.

We begin by determining the pairwise atomic distance between every pair of atoms in a protein to compute $\mathcal{M}_k$. The coordinates of the atoms of residues can be obtained from the relevant PDB file of the protein. Consequently, we can obtain the distribution, $\mathcal{P}(\mathcal{M}_k)$, of protein residue information, $\mathcal{M}_k$, for a given protein. We are especially interested in residues with higher values of $\mathcal{M}_k$. Towards that end, we identify and scrutinize residues which have a value of $\mathcal{M}_k$ greater than 1.5 times the standard deviation from the mean of the distribution of $\mathcal{P}(\mathcal{M}_k)$, for that pair of proteins.

As detailed in the main text, we have considered $p_i^k$ to depend on the atomic distances, $d_{ij}^{kl}$, where $k, l \in \{\mathcal{R}\}$, and $i, j \in \{\mathcal{A}\}$. Our chosen functional form is

$$p_i^k = \frac{\sum_{(j,l)} \omega_{ij}^{kl} (d_{ij}^{kl})^2}{\sum_{i=1}^{N_k} \sum_{(j,l)} \omega_{ij}^{kl} (d_{ij}^{kl})^2}, \quad (1)$$

where, $l \in \{\mathcal{R}\} \backslash \{k\}$ and $j \in \{\mathcal{A}\} \backslash \{i\}$. It should be noted that $\omega_{ij}^{kl}$ is dimensionless and is inspired by well-known Lenard-Jones (LJ) potential [1]. The LJ potential between two interacting entities separated by a distance, $r$, is commonly written as

$$V_{LJ}(r) = 4\epsilon[(\sigma/r)^{12} - (\sigma/r)^6]. \quad (2)$$

Here, $\epsilon$ and $\sigma$ are constants. $\epsilon$ signifies the strength of interaction and $\sigma$ represents the distance at which $V_{LJ}(r) = 0$. Obviously, $V_{LJ}(r) > 0$, as two atoms would repel each other when $r < \sigma$. However, for $r > \sigma$, the interaction would be attractive, implying $V_{LJ}(r) < 0$.

*
†

Herein, we consider, $\omega(d) = [(\frac{\sigma}{d})^6 - (\frac{\sigma}{d})^{12}]$. We next examine the justification of this choice by examining the general form, $p_i^k((\omega_{ij}^{kl} d_{ij}^{kl})^a)$, where $a$ is any integer.

In Fig. S1, we observe the variation of $\omega d^a$ with respect to $d$, for $a = 0, 1, 2$, and 3. For $a = 0, 1$, and 2, the values of $\omega d^a$ becomes negligible at $d > 10$Å. However, $\omega d^3$ does not vanish, even for $d > 10$Å.

It is widely known that the vast majority of non-covalent interactions are active up to $\approx 4$Å in proteins. More importantly, this relevance commonly decreases rather sharply beyond this distance. Indeed, only a small minority of these (including long-range electrostatic interactions) are relevant up to a maximum of $\approx 10$Å in proteins [2]. Thus, a distance of $d \leq 10$Å is more than sufficient for the consideration of all inter-atomic interactions within proteins. More importantly, this relevance commonly decreases rather sharply with distance. Therefore, consideration of $a \geq 3$ is not warranted.

To understand the relative significance of $\omega$, $\omega d$ and $\omega d^2$, we calculate the relevant areas in Fig. S1 using Simpson's one-third rule, as displayed in Table SI. $\Delta A_1$, $\Delta A_2$, and $\Delta A_3$ denote the area for $0 \leq d \leq 5$Å, $5 < d \leq 10$Å, and $0 \leq d \leq 20$Å respectively. These areas are a signature of the number of interactions within the respective regions of distance. Thus, $\Delta A_1/\Delta A_3$ and $\Delta A_2/\Delta A_3$ signify interactions within $0 \leq d \leq 5$Å and $5 < d \leq 10$Å respectively.

For $a = 0$ and $a = 1$, we observe that the value of $\Delta A_2/\Delta A_3$ is low. Therefore, if we choose either $\omega$ or $\omega d$, we would lose the influence of long-range electrostatic interactions towards conformational change. On the other hand, for $\omega d^2$, $\Delta A_2/\Delta A_3 = 17.6\%$, which is relatively high. In summary, choosing $a = 2$ captures the effect of both short-range and long-range covalent interactions and that too most effectively. Therefore, we have chosen $a = 2$ for the calculation of intra-state entropy, $\mathcal{S}_k$.

For the calculation of inter-state entropy, $\mathcal{J}_k^{(\mathcal{P},\mathcal{Q})}$, we consider any two states of a protein, $\mathcal{P}, \mathcal{Q}$ of a protein. Calculation of $\mathcal{J}_k^{(\mathcal{P},\mathcal{Q})}$ requires

$$q_i^k = \frac{\sum_{(j,l)} (\Delta_{ij}^{kl})^b}{\sum_{i=1}^{N_k} \sum_{(j,l)} (\Delta_{ij}^{kl})^b}, \quad (3)$$

where, $\Delta_{ij}^{kl}$ denotes the *difference* in pairwise atomic distances between the two states. The values of $\Delta_{ij}^{kl}$ help us in understanding the amount of conformational change undergone by a residue across two states.

Dimensional homogeneity of the expression of probabilities, $p_i^k = \frac{\sum_{(j,l)} \omega_{ij}^{kl} (d_{ij}^{kl})^2}{\sum_{i=1}^{N_k} \sum_{(j,l)} \omega_{ij}^{kl} (d_{ij}^{kl})^2}$, and, $q_i^k = \frac{\sum_{(j,l)} (\Delta_{ij}^{kl})^b}{\sum_{i=1}^{N_k} \sum_{(j,l)} (\Delta_{ij}^{kl})^b}$ from Eqns. 1 and 3 respectively, would strongly incline us to consider $b = 2$ for $q_i^k$. As discussed in the main text, $\mathcal{J}_k^{(\mathcal{P},\mathcal{Q})}$ is calculated from the structures of two different states of a protein, *and does not arise from direct interactions between atoms of a protein in any given state.* Therefore, physical interactions manifested through $\omega^{kl}$ and $\sigma_i^k$ in the case of $\mathcal{S}_k$, are not relevant for the determination of $\mathcal{J}_k^{(\mathcal{P},\mathcal{Q})}$.

As observed in Fig. 2 of the main text, for some pairs of atom, the value of $\Delta_{ij}^{kl}$ could be as low as 0.2Å. On the other hand, for some pairs the values of $\Delta_{ij}^{kl}$ could be as high as 4.5Å.

Therefore, if the value of $b$ is low, then the contribution of the higher values of $\Delta_{ij}^{kl}$ towards $q_j^k$ becomes negligible. On the other hand, if $b$ is high, $\mathcal{J}_k^{(\mathcal{P},\mathcal{Q})}$ would be solely dictated by higher values of $\Delta_{ij}^{kl}$. In both situations, we would lose valuable information regarding the conformational change of a protein. In our formulation, we address both subtle and significant changes in the positions of residues, and $b = 2$ is most appropriately poised to capture the effects of both lower and higher values of $\Delta_{ij}^{kl}$. Therefore, this coupled with the fact that $b = 2$ accords dimensional homogeneity across the probabilities, $p_i^k$ and $q_i^k$, makes $b = 2$ to be the optimal choice for the inter-state entropy, $\mathcal{J}_k^{(\mathcal{P},\mathcal{Q})}$.

For a given set of random events, the value of entropy tends to a maximum when the probabilities associated with each of these random events are nearly equal to each other. In other words, entropy tends to a maximum for a set of equiprobable events. To evaluate $\mathcal{S}_k$ associated with the $k^{th}$ residue, we calculate the probability, $p_i^k$, associated with each atom $i$ of $k$. $\forall i \in \{k\}$, if the values of $p_i^k$ are equal, then $\mathcal{S}_k$ would tend to a maximum. Conversely, lower values of $\mathcal{S}_k$ indicate that the associated values of $p_i^k$ possess significant inequality among themselves. $p_i^k$ is calculated by considering pairwise atomic distances as discussed above. Naturally, it includes the influence of interactions between all pairs of atoms. A tendency of equality in values $p_i^k$ (implying a larger value of $\mathcal{S}_k$), also indicates that every atom in residue $k$ is likely to experience more interactions with every atom in other residues. Let $\mathcal{S}_{k_1}$ and $\mathcal{S}_{k_2}$ denote the intra-state entropy associated respectively with residues, $k_1$ and $k_2$; such that $\mathcal{S}_{k_2}$ is significantly greater than $\mathcal{S}_{k_1}$. From the above discussion, it follows that atoms in $k_2$ are much more likely to interact with atoms of other residues, than the atoms in $k_1$.

Similarly, the highest values of $\mathcal{J}_k$ would be attained, when the associated values of $q_i^k$ are nearly equal. $q_i^k$ considers the change in pairwise atomic distances, $\Delta_{ij}^{kl}$, as discussed above. $\Delta_{ij}^{kl}$ enables us to understand the movement of atoms of a given residue between two states, both translational and rotational. Higher values of $q_i^k$ indicate a higher probability of change in position across two states. For a given $k$, if the $q_i^k$ values indicate the overall residue movement. This overall residue movement may be either large or small. However, within a given structure, a residue cannot move freely as this would result in steric clashes, and the overall stability of protein would be compromised. Therefore, it is highly likely that for higher values of $\mathcal{J}_k$, $k$ is likely to undergo subtle overall changes rather than a radical one. On the other hand, vastly varying values of $q_i^k$ would indicate that some atoms of the $k^{th}$ residue are likely experience a greater change of position. This scenario indicates an abrupt change in conformation such as a side-chain rotation, which may be important in identifying functionally important residues. As an example, we observe that Trp83 and Trp266 of MutY (Figure S4(a)) and Tyr24A of Hb (Figure S3(c)) do not go through any remarkable conformational changes. However, the values of $\mathcal{J}_k$ associated with these residues are observed to be very high. On the other hand, Tyr106 of CheY possess a lower value of $\mathcal{J}_k$ and the associated conformational change is high, as can be observed in Figure S2(a).

Herein, we are interested in identifying functionally important residues. These residues should possess higher interactions (higher $\mathcal{S}_k$) in both states and higher conformational change (lower $\mathcal{J}_k$) across two states. By definition, PRI, $\mathcal{M}_k^{(\mathcal{P},\mathcal{Q})} = \mathcal{S}_K^{(\mathcal{P})} + \mathcal{S}_k^{(\mathcal{Q})} - \mathcal{J}_k^{(\mathcal{P},\mathcal{Q})}$. If $\mathcal{J}_k^{(\mathcal{P},\mathcal{Q})} > (\mathcal{S}_K^{(\mathcal{P})} + \mathcal{S}_k^{(\mathcal{Q})})$, then $\mathcal{M}_k < 0$. This would imply a low conformational change of the $k^{th}$ residue between two states as well as fewer interactions with others. For Ala335 of aureochrome and Ala109 of Tbx5, $\mathcal{J}_k^{(\mathcal{P},\mathcal{Q})}$ is very high, indicating lower conformational change. On the other hand, their respective $\mathcal{S}_k$ values in both states are low. This implies that due to their position in the protein, they participate in rather few interactions. Therefore, these residues possessing negative PRI values, should not bear high functional importance. We consistently observe that residues with higher PRI values possess greater functional significance.

### I. CheY

Through our formulation, we identify Tyr106 as a crucial residue of CheY as it appears in all ten combinations, shown in Table 1 of the main text. Apart from Tyr106, Arg22 is found six times, while Met63 and Lys119 are found five times. Phe14 and Glu89 are found four times. We find Lys7 and Arg19 thrice while Lys26, Ser56, Lys92 and Lys126 are found twice. Glu27, Glu34, Trp58, Ile95, Phe124 and Met129 are found in one combination. Among these residues, the conformational change of Tyr106 is clearly observable in Figure Fig. S2(a). The rotational restriction of the Tyr106 side chain is crucial for activating CheY [3]. The perturbation propagation algorithm [4] also identifies Asp57$\rightarrow$ Thr87$\rightarrow$ Tyr106$\rightarrow$ ligand as the most critical allosteric pathway in CheY. Briefly, phosphorylation at Asp57 (start site) allosterically activates CheY to bind with FLiM (end site). Phosphorylation of Asp57 causes additional hydrogen bonding interactions with Thr87 preceded by the latter's movement towards the phosphorylation site. And the space vacated by Thr87 is occupied by Tyr106. It facilitates Thr87-Tyr106 coupling, which is responsible for allosteric effects in CheY [4]. Unlike Thr87, the movement of the neighbouring Glu89 appears more distinct towards Tyr106. For CheY, Asp57 is reported to be extremely important in allosteric communication. However, our formulation identifies the neighbouring residues of Asp57, i.e., Ser56 and Trp58, rather than identifying Asp57 directly. The sudtle conformational changes of Ser56 and Trp58 could be observed in Fig. S2(a) and their functional importances are discussed in the main text. We could also identify the movement of Glu89 and Lys119 in Figure Fig. S2(a). In the presence of mechanical perturbation Glu89 is observed to demonstrate relatively lower thermodynamic response. However, mechanical response is higher [5]. Lys119 moves farther away to accommodate the C-terminal end of the peptide. The side chain of Lys119 forms a salt-bridge with Asn16-Film in CheY-N16-FLiM complex [6]. Among other residues, Thr87, Ala88, and Lys109 is engaged with phosphoryl group via hydrogen bonding interaction. Mutation of Ile95 to valine enhances the clockwise rotation of the flagellar motor and is correlated with high affinity binding to FLiM. Direct interaction of Ile95 with FLiM is suggestive of signal transmission through hydrophobic interactions in other response regulators as well. Further, Glu27 is located in that area of CheY, which helps it in binding to the flagellar switch [7]. Apart from being the terminal residue, the movement of Met129 seems relevant considering the movement of the whole helical segment, upon FLiM binding. Lys26 is solvent-accessible since it is located on the surface of CheY's helix $alpha1$. The mutation of this residue does not influence the rates of phosphorylation and autodephosphorylation. However, the CheY26KE mutant develops a resistance to CheZ activity. The protein residue information values of each residue associated with the CheY proteins are displayed in Table S1(a) to Table S1(j) herein.

### II. NolR

We can easily notice a high conformational shift of the residues at the wing and DNA-binding regions of NolR from Fig. S2(b). The conformational alterations of these residues facilitate the protein's ability to bind to DNA. Alongside conformational change, these identified residues possess high PRI values. Their functional importances are discussed in detail in the main text. Table S2 illustrates the corresponding protein residue information values for these residues.

### III. Streptavidin

The conformational changes in Trp79, Trp92, Trp108 and Trp120 located at the biotin-binding site are not readily distinguishable in Figure S2(c). However, the PRI values associated with these four Trp residues have been observed to be high. These residues are very crucial for biotin binding. The tetrameric structure of streptavidin tightens upon binding with the biotin ligand [8]. Experimental observations considering key fluores-

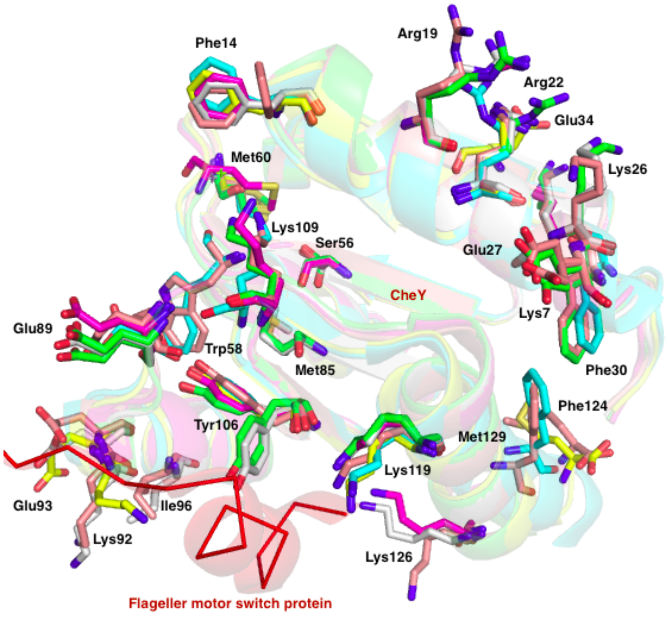

**(a)** CheY (signaling protein)

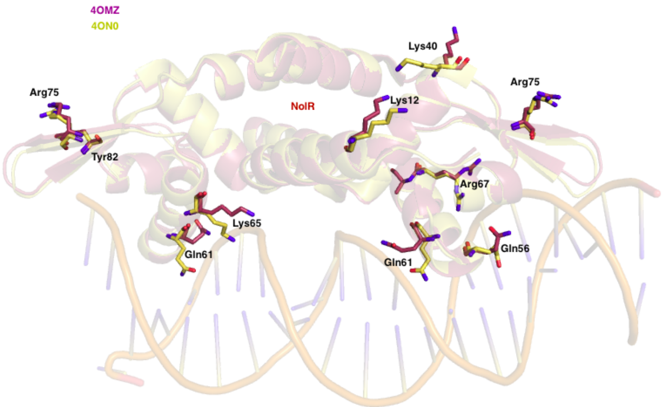

**(b)** NolR (transcription factor)

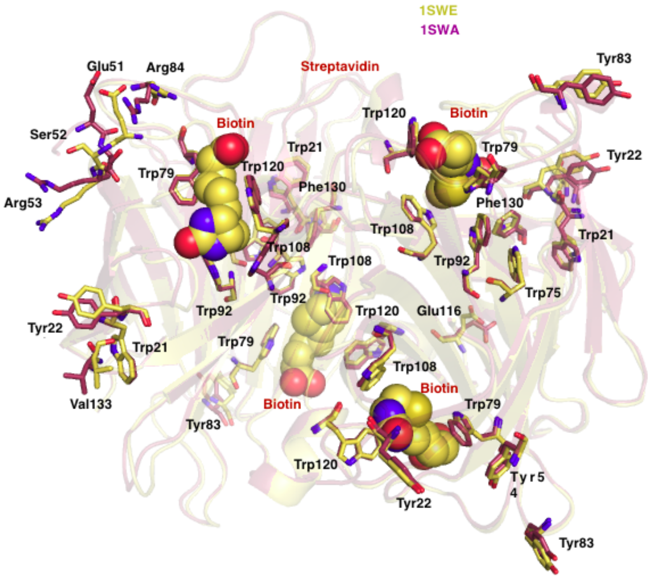

**(c)** Streptavidin (biotin-binding protein)

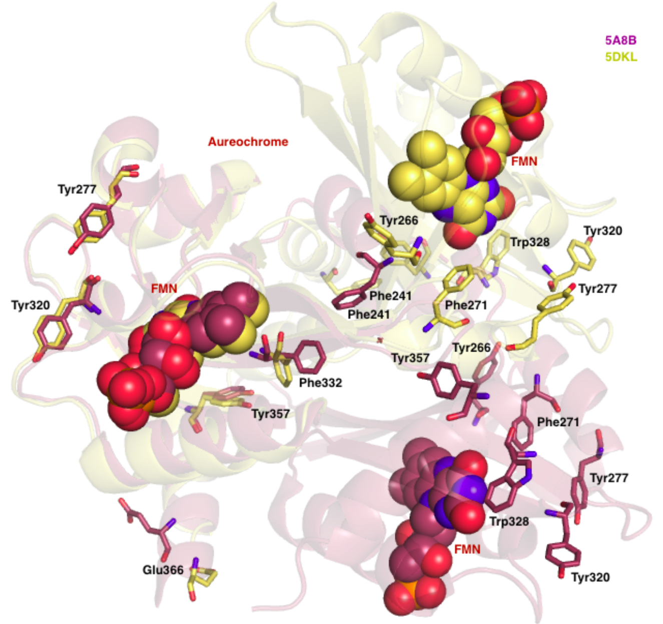

**(d)** Aureochrome (LOV protein)

**Figure S2:** Residues possessing higher values of PRI in: (a) CheY, (b) NolR, (c) Streptavidin, and (d) Aureochrome

cence parameters and mutagenesis studies suggest major structural changes in streptavidin tetramer only during the first two biotin binding events. While the third binding event induces structural changes across opposite subunits, changes are insignificant in the fourth and final binding of the biotin molecule [8]. The amino acid residues involved in biotin binding differ from one subunit of streptavidin to another. For example, almost the same set of residues is found to be important in biotin binding at subunits B and C. We do see the involvement of additional residues at subunit C during biotin binding events. Mostly, polar and aromatic residues line the biotin-binding pocket. Extensive polar/hydrogen bonding interactions enhance the stability of the complex. For instance, the synchronized movement of both Glu51 and Arg84 towards the biotin-binding pocket with the formation of a salt bridge ensures the stable burial of biotin within streptavidin. Apart from the four important Trp residues, the conformational changes of other residues with high PRI values could be observed in Figure S2(c). Most of these identified residues are polar and aromatic. Inter-residue hydrogen bonding interactions among aro-

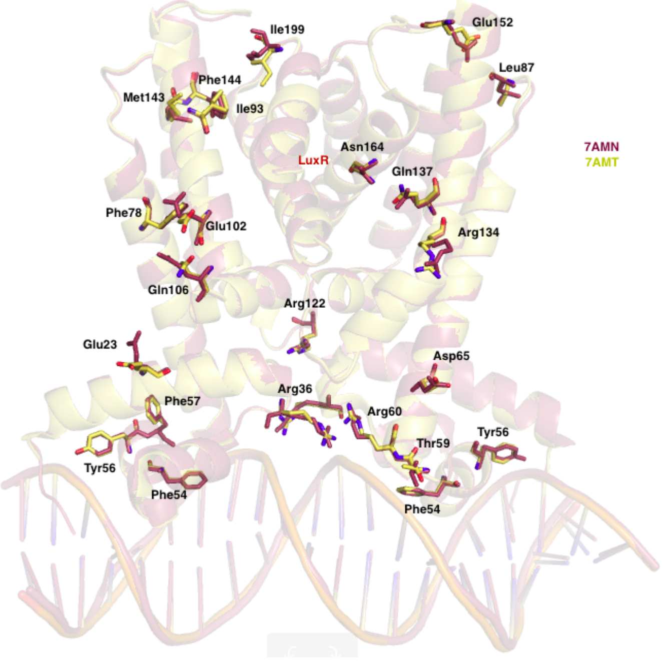

**(a)** LuxR (transcription factor)

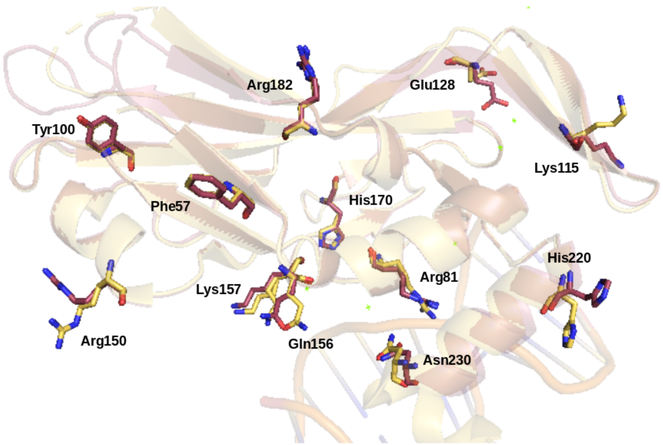

**(b)** TBX5 (transcription factor)

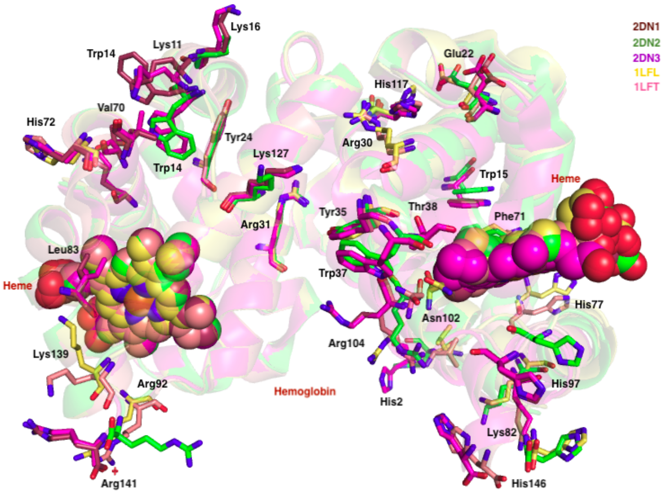

**(c)** Hemoglobin (oxygen storage/transport)

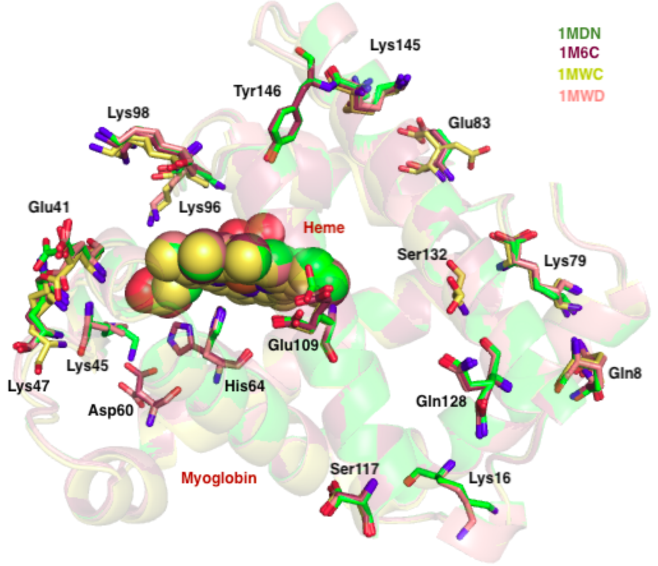

**(d)** Myoglobin (oxygen storage/transport)

**Figure S3:** Residues possessing higher values of PRI in: (a) LuxR, (b) TBX5, (c) Hemoglobin, and (d) Myoglobin

matic amino acids create the necessary 'hydrophobic box' for biotin-binding [9]. Residues like Trp21, Tyr22, and Val133 are close to the surface and engage in various interactions during the apo-holo state through structural reorientations, as discussed in the main text. These residues undergo higher conformational changes. On the other hand, Trp21 and Tyr54 residues do not experience any noticeable conformational changes. However, these residues are also crucial for birding-binding, and the main text has discussed their functional significance. Maximum participation of residues from the C subunit, followed by that from B, suggests that all subunits participate differently in the biotin-binding process – as evident in fluorescence studies [8]. Differential participation of amino acid residues from four subunits leads to cooperative structural changes and thus allostery in the biotin-binding mechanism. The associated values of protein residue information of this biotin-binding protein have been shown in Table S3.

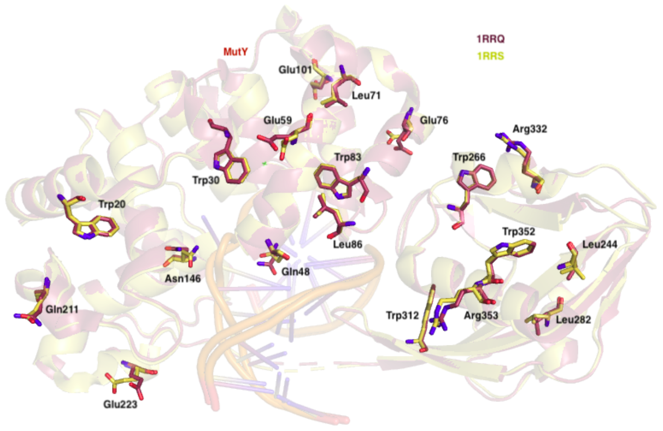

**(a)** MutY (DNA repair enzyme)

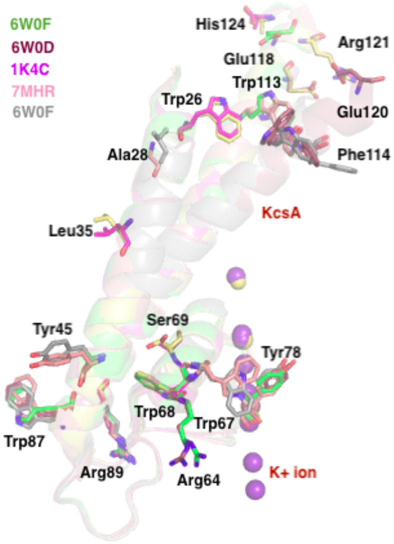

**(b)** KcsA ($K^+$ ion channel)

**Figure S4:** Residues possessing higher values of PRI in: (a) MutY and (b) KcsA

### IV. Aureochrome

Because of the rather high RMSD values in the (5A8B, 5DKL) pair, aligning these structures is difficult. With our method, we can simultaneously identify both subtle and significant changes in the orientation of the residues — as in the case of Tyr277 and Tyr320 in two distinct chains (A and B), shown in Fig. S2(d). In our method, we treat the intra-state entropy of a state as a function of pairwise atomic distances within that state. However, while calculating inter-state entropy, the difference in pairwise atomic distances between two states is taken into account. This method overlooks the non-compatibility alignment issues with this Aureochrome pair. Using calculated protein residue information, we can determine the significant residues in both A and B chains, as indicated in Table S4.

### V. LuxR

The absolute conformational changes of Glu23, Leu87, Ile93, Glu102, Glu106, Arg134 and Glu152 are clearly visible in Fig. S3(a). Most of these residues belong to the surface region of the (7AMN,7AMT) pair and facilitate various biological functions as discussed in the main text. On the other hand, Phe54, Tyr56, Phe57, Phe78 and Asn164 undergo subtle conformational changes, which are not so easily detected in Fig. S3(a). Among these residues, Phe54, Tyr56 and Phe57 are very close to the DNA binding region and likely play an important role in the repression and activation states of LuxR. Asn164, another residue identified to be significant due to its interchain hydrogen bonding interactions, as discussed in the main text. Other residues close to DNA binding regions, such as Thr59, Arg36 and Arg60, have been observed to go through higher conformational changes. The protein residue information values of all of these residues have been enlisted in Table S5.

### VI. TBX5

We can determine which residues for the 2X6U and 2X6V pair have greater conformational changes by looking at Table S6. As seen in Fig. S3(b), these residues are located on the surface and in the DNA-binding region. In the DNA bound state (2X6V), the Arg81 side chain extends to establish contact with the phosphate oxygen of DG'18. Interestingly, analyses of mutational effects from clinical studies of Halt-Oram syndrome reveal that Gly80Arg mutation completely abrogates DNA binding [10]. Unlike Arg81, Gly80 does not interact with DNA. However, our *in silico* mutagenesis studies indicate that Gly80Arg mutation results in severe steric clash with His170, located at the opposite side of Arg81. Therefore, it is clear that the presence of Gly80 is critical not only to achieve DNA binding at the major groove by Arg81, but also to maintain structural stability by accommodating His170 on the opposite side.

### VII. Hb

The $\alpha$ and $\beta$ subunits of Hb are comprised of 7 and 8 helices respectively. The helices named A to H are joined to each other by non-helical "corners". Heme, nested between E and F helices anchors a ferrous ion via a covalent linkage to the proximal His from F8. The distal site is available for the ligands to bind. Oxygen upon binding to the distal site is stabilised by hydrogen bonding interactions from the neighbouring His (E7). Ligand binding induces series of conformational changes to increase binding affinity. Initiation occurs at the heme binding pocket and extend to the E helix, CD and FG corners, central water cavity, $\alpha_1$-$\beta_2$ interface as well as other subunits. Experimental studies suggest participation of A helix, F helix and G helix residues as well in modulating Hb allostery. Cooperativity, thus triggered, shifts the allosteric equilibrium from the $T$-state to $R$-state [11]. Structure-function studies supported by computational, thermodynamic, and spectroscopic data confirm the presence of multi-state structures besides the classical $T$ and $R$ end-states. To elaborate, the relaxed state instead of being in the classical $R$-state is an ensemble of fully liganded states with discrete conformations of quaternary structure. Based on multiple structural parameters like screw-axis rotation/translation, cross-dimer interactions *etc*, these states can be assigned as $R2$, $RR2$, $R3$ and $RR3$. $R$, $R2$ and $RR2$ represent closed conformations of His(E7)-ligand channel suggesting heme ligand transport. $R3$ and $RR3$ represent open conformations facilitating the release of ligand [11]. Among all the PDBs considered herein, we prioritise two high-resolution structures 2DN1 and 2DN2, representing $R$ and $T$-states respectively. For comparison with the other ligand bound $R$-state we consider the carbon monoxide liganded Hb (2DN3) as well. Comparison among tetrameric subunits could have been the best to understand the flow of information from the unliganded to liganded state. However, due to the unavailability of suitable high-resolution structures consisting of two polypeptide chains [$\alpha$-$\beta$] in the asymmetric unit, our calculations are restricted to the flow of information in $\alpha\beta$ subunit of Hb. Residues undergoing relative conformational changes are distributed across Hbs $\alpha\beta$. The respective positions of amino acid residues in the $\alpha$ subunit are Lys11, Trp14, Lys16 in A helix, Tyr24, Arg31 in B helix, Lys60, Thr67, Val70, His72 in D helix, Leu83 in E helix, Arg92 in EF corner and Lys127, Lys139, Arg141 in G helix. The residues at the $\beta$ subunit are located at Val1, His2 in A helix; Trp15, Glu22, Arg30, Tyr35 in B helix; Trp37 in C helix; Phe71 in E helix; His77 at the EF corner; Lys82 in F helix; His97 at the FG corner; Asn102, Arg104, His117 in G helix and His146 in H helix. Therefore, besides the residues located at A, E, F and G helices, CD/FG corners, we find that amino acid residues located at other helical components (e.g. B, C, D as well as H helices), and corners (e.g. EF) also participate in $T \to R$ transition. Notably, we find that a large number of amino acids (especially from the D-helix), undergo relative conformational changes while transitioning between unliganded and CO-bound states. Such a trend is not observed during the transition from unliganded to the $O_2$-bound state. For the identified residues, we find several references in literature. For example, the 'flexible joint' and 'switch' at the $\alpha_1\beta_2$ (*SI text*) play a crucial role in $T \to R$ transition. The aforementioned 'flexible joint' consists of the FG corner, G-helix from chain A ($\alpha$ subunit) and the C-helix, CD corner from chain B ($\beta$ subunit). The 'switch' region consists of the FG corner, G-helix from chain B and C-helix, CD corner from chain A. Two 'stereochemical differences' have been underlined as the major determinants of $T \to R$ transition [12]. One prime distinction is the hydrogen bond between Asp94A and Asn102B in the $R$-state, which is absent in the $T$-state. Further, Asp99B-Asn97A/Tyr42A hydrogen bonding interactions are present in the $T$-state but absent in the $R$-state. In the same region, His97B undergoes marked differences in its conformation during the $T \to R$ transition. In fact, the change in interactions mediated by His97B in both $T$ and $R$ states is considered as a major steric barrier to the quaternary structure change. The Asp94A-Asn102B hydrogen bond connecting the flexible joint and switch region remains intact in the $R$ and $R2$ states. The C-terminal segments of both A and B are crucial components of Hb tetramer because mutations in this region can potentially disrupt the structure-function of Hb. Also, Asn130 is known to participate in the newly formed salt-bridge [13]. In Figure S3(c), we have been able to identify most of the residues with drastic conformational changes. Residues with high PRI values (Table S7) are located all throughout the Hb structure, as shown in Figure S3(c). As discussed in the main text, His146 possess a high PRI value and is crucial for $T \to R$ transition. It makes intra-, and inter-subunit interactions with $Asp94$, in the case of deoxygenated Hb. His146 is highly disordered in $R$, $R2$ and $RR3$ structures compared to the $T$ structure. His146 is the critical regulator of the Bohr effect [11].

### VIII. Mb

In Fig. S3(d), the conformational alterations of Ser117, Glu128, Lys145, and Tyr146 are visibly less than those of Gln8, His64, Lys79, Lys98, and Glu109. All of these residues, nevertheless, are crucial because they support a variety of biological processes as discussed in the main text. They are anticipated to experience greater reorientations in different states because they are mostly located in the periphery and heme-binding areas. The associated information values are displayed in Table S8. We try to explore its residues playing essential role in de-oxy $\to$ oxy/carbomonoxy transition as well as finding out the basis of varying affinity towards CO and $O_2$. In the ligand bound conformation, $Fe^{2+}$ in a planar heme surface has a low spin state. Upon the detachment of ligand, $Fe^{2+}$ converts to a high spin state and deviates from planarity by 0.3Å, thereby resulting in a dome-shaped structure. Interestingly, the space filling model of Mb shows no exit/entry pathways for $O_2/CO$. Therefore, it is certain that the conformational changes within Mb are absolutely essential for $O_2/CO$ binding [14]. However, the ligand binding/escape and therefore the opening/closing of Mb molecule is so transient that only time-resolved or high resolution structures to some extent can reveal dynamical features in Mb. Monomeric Mb consists of eight $\alpha$ helices (A to H), connected via flexible corners. The H64L/V68N mutant shows very high affinity towards oxygen — an additional H-bond from the amide group of Asn (regardless of the nature of amino acid at the 64 position) stabilises the bound Oxygen molecule. While $O_2$ affinity increases nearly three-fold, CO affinity decreases fourfold to sixfold in the V68N mutant. Even though $O_2$ binding is favoured in the V68N mutant, the Asn side chain sterically inhibits other ligands from approaching the Fe centre through the distal heme binding pocket. Comparison between wild-type/V68N de-oxy Mb and V68N Mb-CO highlights His64 as a crucial residue. High resolution crystal structures, time resolved data and UV Resonance Raman spectroscopy (UVRR) reveal distinct movements of F helix and FG corner away from/towards the heme moiety, upon dissociation of CO. UVRR along with time resolved fluorescence spectroscopy conducted on wild type and Trp mutants reveal notable structural changes between A helix and E helix. This is mostly attributable to the rearrangement of A helix due to the faster motion of E helix, which harbours distal Histidine. While the change in environment around Trp14 is transient, the same around Trp7 is substantial. Incidentally, both these Trp are located at the A helix. Calculation of correlation coefficients between "fluctuational motions" of residues as well as heme moiety in deoxy-Mb identified several residues from A, C, E, G and H helix as well as CD, EF, FG and HC corners, possessing strong dynamic interactions with heme [15]. Remarkably, all our identified residues are located at A, C, E, G and H helices as well as CD, EF and FG corners. However, Ref. [15] is only limited to deoxy-Mb. Therefore, not all the residues identified by them feature in our list.

### IX. MutY

In the case of the (1RRS, 1RRQ) pair, we observe that a number of the identified residues, mentioned in Table S9, have not undergone remarkable conformational changes. The respective importance of these residues in terms of stability of the protein and various interactions have been discussed in the main text. On the other hand, the conformational orientation of side chain of Glu26, Gln48, Glu59 and Glu223 etc can be easily visualized, as shown in Figure S4(a). The catalytic domain of MutY, having strong structural resemblance with Endonuclease III, consists of two structural modules: the 6-helix barrel and another module bearing [4Fe-4S] cluster. MutY interacts with damaged DNA right at the juncture of these two structural modules of the catalytic domain. Additionally, MutY also possesses a C-terminal domain, which is required for OG recognition. 1RRQ is a co-crystal structure containing MutY bound with damaged DNA bearing 8-OG, 1RRS is a co-crystal structure of MutY and DNA with an abasic site. Our formulation identifies Gln48, Glu53 and Glu223 as residues having high PRI, undergoing distinct conformational change while comparing between the two states. The anti-glycosidic bond of 8-OG nucleoside in MutY bound state indicates that this residue abstains from participation in base pairing interaction. However, in absence of MutY, 8-OG attains syn conformation to pair with adenine. *Syn* to *anti* conversion of 8-OG implies a 180° swivelling along the glycosidic bond and an extrusion of adenine from the helix to avoid a potential steric clash. The space thus vacated by adenine can be filled up by the Gln48 side chain. This is succeeded by $\pi$-stacking interactions of the amide bearing side chain of Gln48 with the neighbouring $3'$ base as well as hydrogen bonding interactions with the phosphate backbone [16]. Asp160 seems to impart structural stability to the adjacent helix by backbone-side chain hydrogen bonding interactions.

### X. KcsA

From Table S10 and Figure S4(b), we observe that most of the identified residues lie either in the vicinity of $K^+$ ion or in the end of $\alpha$ helices. The corresponding conformational changes are clearly visible in Fig. S4(b). The selectivity filter sequence TTVGYGD in KcsA channels is conserved in all organisms and precisely regulates the selective yet high-speed movement of $K^+$ ions across the membrane. As discussed in main text, some of the residues which flank this conserved filter region possess higher PRI values. The C-type inactivation in KcsA channel not only depends on the conductive $\rightarrow$ constricted conformational changes of the selectivity filter, but also on the residues lining the pore domain [17]. A strong allosteric coupling of residues in and/or around the filter region ensures the structural integrity of the whole region, and thus $K^+$ conductivity. Tight interaction between the tip of the helices of S5 and S6 bordering the mouth of the channel is important. Not only does it play a critical role in C-type inactivation in KcsA but also in the coupling between voltage sensor and pore domain in the Drosophila shaker channel. A comparison of structure across the open and closed state KcsA reveals translation and counter-clockwise rotation of the inner helices (S5, and S6) with respect to the centre of symmetry of the channel. Electron Paramagnetic Resonance (EPR) spectroscopy along with site-directed spin labelling methods reveal significant mobility towards the N-terminal half of the S5 helix and rather remarkable movement towards the C-terminal half of the S6 helix [18]. The S6 helix, hinged at Gly99, shows rather drastic conformational changes at these terminal residues – leading to opening/closing of the channel.

| Table S1(a): 1JBE (P), 1C4W (Q) | | | | | | | | | | | | | | |
|---|---|---|---|---|---|---|---|---|---|---|---|---|---|---|
| Residue | | | S (P) | S (Q) | J (P,Q) | PRI | | Residue | | | S (P) | S (Q) | J (P,Q) | PRI |
| ALA | 2 | A | 0.677 | 0.65 | 0.984 | 0.343 | | LEU | 66 | A | 0.734 | 0.741 | 0.992 | 0.483 |
| ASP | 3 | A | 0.744 | 0.766 | 0.99 | 0.52 | | GLU | 67 | A | 0.749 | 0.815 | 0.804 | 0.76 |
| LYS | 4 | A | 0.761 | 0.74 | 0.989 | 0.512 | | LEU | 68 | A | 0.745 | 0.758 | 1 | 0.503 |
| GLU | 5 | A | 0.807 | 0.763 | 0.99 | 0.58 | | LEU | 69 | A | 0.742 | 0.726 | 0.999 | 0.469 |
| LEU | 6 | A | 0.733 | 0.742 | 0.993 | 0.482 | | LYS | 70 | A | 0.806 | 0.795 | 0.998 | 0.603 |
| LYS | 7 | A | 0.828 | 0.846 | 0.896 | 0.778 | | THR | 71 | A | 0.712 | 0.718 | 0.999 | 0.431 |
| PHE | 8 | A | 0.826 | 0.824 | 0.998 | 0.652 | | ILE | 72 | A | 0.739 | 0.752 | 0.992 | 0.499 |
| LEU | 9 | A | 0.746 | 0.745 | 0.997 | 0.494 | | ARG | 73 | A | 0.853 | 0.853 | 0.994 | 0.712 |
| VAL | 10 | A | 0.679 | 0.686 | 0.999 | 0.366 | | ALA | 74 | A | 0.724 | 0.69 | 0.98 | 0.434 |
| VAL | 11 | A | 0.689 | 0.69 | 1 | 0.379 | | ALA | 77 | A | 0.788 | 0.757 | 0.972 | 0.573 |
| ASP | 12 | A | 0.803 | 0.783 | 0.949 | 0.637 | | MET | 78 | A | 0.849 | 0.835 | 0.936 | 0.748 |
| ASP | 13 | A | 0.736 | 0.77 | 0.979 | 0.527 | | SER | 79 | A | 0.683 | 0.705 | 0.991 | 0.397 |
| PHE | 14 | A | 0.775 | 0.773 | 0.995 | 0.553 | | ALA | 80 | A | 0.506 | 0.518 | 0.993 | 0.031 |
| SER | 15 | A | 0.736 | 0.734 | 0.994 | 0.476 | | LEU | 81 | A | 0.745 | 0.718 | 0.746 | 0.717 |
| THR | 16 | A | 0.676 | 0.706 | 0.75 | 0.632 | | PRO | 82 | A | 0.651 | 0.693 | 0.999 | 0.345 |
| MET | 17 | A | 0.713 | 0.745 | 0.997 | 0.461 | | VAL | 83 | A | 0.705 | 0.705 | 0.997 | 0.413 |
| ARG | 18 | A | 0.835 | 0.853 | 0.998 | 0.69 | | LEU | 84 | A | 0.736 | 0.735 | 0.999 | 0.472 |
| **ARG** | **19** | **A** | **0.748** | **0.818** | **0.626** | **0.94** | | MET | 85 | A | 0.763 | 0.771 | 0.737 | 0.797 |
| ILE | 20 | A | 0.8 | 0.783 | 0.996 | 0.587 | | VAL | 86 | A | 0.691 | 0.699 | 0.994 | 0.396 |
| VAL | 21 | A | 0.685 | 0.694 | 0.997 | 0.382 | | THR | 87 | A | 0.724 | 0.749 | 0.998 | 0.475 |
| **ARG** | **22** | **A** | **0.853** | **0.887** | **0.565** | **1.175** | | AALA | 88 | A | 0.651 | 0.701 | 0.98 | 0.372 |
| ASN | 23 | A | 0.81 | 0.864 | 0.891 | 0.783 | | AGLU | 89 | A | 0.694 | 0.745 | 0.967 | 0.472 |
| LEU | 24 | A | 0.774 | 0.771 | 0.993 | 0.552 | | AALA | 90 | A | 0.662 | 0.664 | 0.834 | 0.492 |
| LEU | 25 | A | 0.738 | 0.73 | 1 | 0.468 | | ALYS | 91 | A | 0.836 | 0.813 | 0.901 | 0.748 |
| LYS | 26 | A | 0.76 | 0.702 | 0.693 | 0.769 | | LYS | 92 | A | 0.779 | 0.801 | 0.979 | 0.601 |
| GLU | 27 | A | 0.812 | 0.825 | 0.885 | 0.752 | | GLU | 93 | A | 0.719 | 0.778 | 0.998 | 0.499 |
| LEU | 28 | A | 0.735 | 0.734 | 0.998 | 0.471 | | ASN | 94 | A | 0.763 | 0.77 | 0.997 | 0.536 |
| GLY | 29 | A | 0.641 | 0.633 | 0.999 | 0.275 | | ILE | 95 | A | 0.78 | 0.778 | 0.999 | 0.559 |
| PHE | 30 | A | 0.842 | 0.83 | 0.999 | 0.673 | | ILE | 96 | A | 0.805 | 0.807 | 0.992 | 0.62 |
| ASN | 31 | A | 0.746 | 0.758 | 0.998 | 0.506 | | ALA | 97 | A | 0.622 | 0.599 | 0.997 | 0.224 |
| ASN | 32 | A | 0.728 | 0.738 | 0.998 | 0.468 | | ALA | 98 | A | 0.569 | 0.581 | 0.998 | 0.152 |
| VAL | 33 | A | 0.696 | 0.712 | 0.998 | 0.41 | | ALA | 99 | A | 0.627 | 0.64 | 0.997 | 0.27 |
| GLU | 34 | A | 0.834 | 0.83 | 0.998 | 0.666 | | GLN | 100 | A | 0.815 | 0.819 | 0.988 | 0.646 |
| GLU | 35 | A | 0.75 | 0.751 | 0.998 | 0.503 | | ALA | 101 | A | 0.481 | 0.422 | 0.999 | -0.096 |
| ALA | 36 | A | 0.536 | 0.471 | 0.995 | 0.012 | | GLY | 102 | A | 0.638 | 0.636 | 0.995 | 0.279 |
| GLU | 37 | A | 0.729 | 0.783 | 0.971 | 0.541 | | ALA | 103 | A | 0.474 | 0.495 | 1 | -0.031 |
| ASP | 38 | A | 0.768 | 0.782 | 1 | 0.55 | | SER | 104 | A | 0.642 | 0.636 | 1 | 0.278 |
| GLY | 39 | A | 0.621 | 0.616 | 0.999 | 0.238 | | GLY | 105 | A | 0.628 | 0.621 | 1 | 0.249 |
| VAL | 40 | A | 0.709 | 0.724 | 0.996 | 0.437 | | **TYR** | **106** | **A** | **0.769** | **0.822** | **0.525** | **1.066** |
| ASP | 41 | A | 0.744 | 0.763 | 0.997 | 0.51 | | VAL | 107 | A | 0.655 | 0.683 | 1 | 0.338 |
| ALA | 42 | A | 0.567 | 0.572 | 1 | 0.139 | | VAL | 108 | A | 0.667 | 0.724 | 0.997 | 0.394 |
| LEU | 43 | A | 0.752 | 0.77 | 0.999 | 0.523 | | **LYS** | **109** | **A** | **0.732** | **0.796** | **0.682** | **0.846** |
| ASN | 44 | A | 0.731 | 0.737 | 0.999 | 0.469 | | PRO | 110 | A | 0.707 | 0.69 | 0.999 | 0.398 |
| LYS | 45 | A | 0.844 | 0.855 | 0.908 | 0.791 | | PHE | 111 | A | 0.81 | 0.811 | 0.993 | 0.628 |
| LEU | 46 | A | 0.737 | 0.752 | 1 | 0.489 | | THR | 112 | A | 0.731 | 0.75 | 0.998 | 0.483 |
| GLN | 47 | A | 0.746 | 0.667 | 0.969 | 0.444 | | ALA | 113 | A | 0.602 | 0.562 | 0.991 | 0.173 |
| ALA | 48 | A | 0.516 | 0.482 | 1 | -0.002 | | ALA | 114 | A | 0.672 | 0.659 | 0.998 | 0.333 |
| GLY | 49 | A | 0.643 | 0.634 | 0.999 | 0.278 | | THR | 115 | A | 0.701 | 0.707 | 0.999 | 0.409 |
| GLY | 50 | A | 0.643 | 0.631 | 0.999 | 0.275 | | LEU | 116 | A | 0.723 | 0.713 | 0.992 | 0.444 |
| TYR | 51 | A | 0.857 | 0.871 | 0.999 | 0.729 | | GLU | 117 | A | 0.795 | 0.79 | 0.83 | 0.755 |
| GLY | 52 | A | 0.63 | 0.631 | 0.999 | 0.262 | | GLU | 118 | A | 0.645 | 0.665 | 0.909 | 0.401 |
| PHE | 53 | A | 0.846 | 0.85 | 0.998 | 0.698 | | **LYS** | **119** | **A** | **0.866** | **0.854** | **0.844** | **0.876** |
| VAL | 54 | A | 0.703 | 0.698 | 0.999 | 0.402 | | LEU | 120 | A | 0.732 | 0.736 | 0.995 | 0.473 |
| ILE | 55 | A | 0.737 | 0.743 | 0.999 | 0.481 | | ASN | 121 | A | 0.747 | 0.768 | 0.991 | 0.524 |
| **SER** | **56** | **A** | **0.871** | **0.868** | **0.716** | **1.023** | | LYS | 122 | A | 0.738 | 0.692 | 0.929 | 0.501 |
| TRP | 58 | A | 0.866 | 0.873 | 0.991 | 0.748 | | ILE | 123 | A | 0.746 | 0.758 | 0.987 | 0.517 |
| ASN | 59 | A | 0.648 | 0.714 | 0.729 | 0.633 | | PHE | 124 | A | 0.836 | 0.841 | 0.985 | 0.692 |
| MET | 60 | A | 0.757 | 0.735 | 0.999 | 0.493 | | GLU | 125 | A | 0.644 | 0.647 | 0.802 | 0.489 |
| PRO | 61 | A | 0.67 | 0.683 | 0.999 | 0.354 | | LYS | 126 | A | 0.612 | 0.71 | 0.975 | 0.347 |
| ASN | 62 | A | 0.74 | 0.755 | 0.999 | 0.496 | | LEU | 127 | A | 0.706 | 0.735 | 0.797 | 0.644 |
| **MET** | **63** | **A** | **0.878** | **0.872** | **0.795** | **0.955** | | GLY | 128 | A | 0.646 | 0.656 | 0.999 | 0.303 |
| ASP | 64 | A | 0.719 | 0.602 | 0.999 | 0.322 | | MET | 129 | A | 0.864 | 0.886 | 0.995 | 0.755 |
| GLY | 65 | A | 0.628 | 0.621 | 0.999 | 0.25 | | | | | | | | |

| Table S1(b): 1JBE (P), 1F4V (Q) | | | | | | | | | | | | | | |
|---|---|---|---|---|---|---|---|---|---|---|---|---|---|---|
| Residue | | | S (P) | S (Q) | J (P,Q) | PRI | | Residue | | | S (P) | S (Q) | J (P,Q) | PRI |
| ALA | 2 | A | 0.679 | 0.627 | 0.999 | 0.307 | | GLY | 65 | A | 0.624 | 0.619 | 1 | 0.243 |
| ASP | 3 | A | 0.744 | 0.735 | 0.958 | 0.521 | | LEU | 66 | A | 0.735 | 0.76 | 0.933 | 0.562 |
| LYS | 4 | A | 0.761 | 0.789 | 0.999 | 0.551 | | GLU | 67 | A | 0.748 | 0.71 | 0.922 | 0.536 |
| GLU | 5 | A | 0.807 | 0.768 | 0.99 | 0.585 | | LEU | 68 | A | 0.744 | 0.726 | 0.922 | 0.548 |
| LEU | 6 | A | 0.733 | 0.721 | 0.961 | 0.493 | | LEU | 69 | A | 0.742 | 0.742 | 1 | 0.484 |
| LYS | 7 | A | 0.828 | 0.861 | 0.996 | 0.693 | | LYS | 70 | A | 0.806 | 0.809 | 0.997 | 0.618 |
| PHE | 8 | A | 0.826 | 0.845 | 0.996 | 0.675 | | THR | 71 | A | 0.712 | 0.77 | 0.913 | 0.569 |
| LEU | 9 | A | 0.746 | 0.741 | 1 | 0.487 | | ILE | 72 | A | 0.739 | 0.758 | 1 | 0.497 |
| VAL | 10 | A | 0.679 | 0.678 | 1 | 0.357 | | ARG | 73 | A | 0.853 | 0.849 | 1 | 0.702 |
| VAL | 11 | A | 0.693 | 0.701 | 1 | 0.394 | | ALA | 74 | A | 0.724 | 0.718 | 0.998 | 0.444 |
| ASP | 12 | A | 0.799 | 0.772 | 0.99 | 0.581 | | ALA | 77 | A | 0.788 | 0.672 | 0.986 | 0.474 |
| ASP | 13 | A | 0.735 | 0.755 | 0.999 | 0.491 | | MET | 78 | A | 0.849 | 0.849 | 0.976 | 0.722 |
| PHE | 14 | A | 0.775 | 0.864 | 0.909 | 0.73 | | SER | 79 | A | 0.683 | 0.71 | 0.999 | 0.394 |
| SER | 15 | A | 0.736 | 0.709 | 0.996 | 0.449 | | ALA | 80 | A | 0.506 | 0.51 | 1 | 0.016 |
| THR | 16 | A | 0.676 | 0.656 | 0.884 | 0.448 | | LEU | 81 | A | 0.745 | 0.728 | 0.924 | 0.549 |
| MET | 17 | A | 0.712 | 0.755 | 0.836 | 0.631 | | PRO | 82 | A | 0.651 | 0.653 | 1 | 0.304 |
| ARG | 18 | A | 0.835 | 0.83 | 0.999 | 0.666 | | VAL | 83 | A | 0.704 | 0.682 | 1 | 0.386 |
| ARG | 19 | A | 0.746 | 0.752 | 0.852 | 0.646 | | LEU | 84 | A | 0.737 | 0.729 | 1 | 0.466 |
| ILE | 20 | A | 0.799 | 0.755 | 0.996 | 0.558 | | MET | 85 | A | 0.76 | 0.707 | 0.961 | 0.506 |
| VAL | 21 | A | 0.684 | 0.705 | 0.999 | 0.39 | | VAL | 86 | A | 0.698 | 0.705 | 0.976 | 0.427 |
| **ARG** | **22** | **A** | **0.853** | **0.863** | **0.921** | **0.795** | | THR | 87 | A | 0.726 | 0.723 | 0.935 | 0.514 |
| ASN | 23 | A | 0.733 | 0.809 | 0.996 | 0.546 | | AALA | 88 | A | 0.648 | 0.698 | 0.97 | 0.376 |
| LEU | 24 | A | 0.774 | 0.773 | 0.998 | 0.549 | | AGLU | 89 | A | 0.693 | 0.802 | 0.899 | 0.596 |
| LEU | 25 | A | 0.738 | 0.731 | 0.999 | 0.47 | | AALA | 90 | A | 0.662 | 0.693 | 0.959 | 0.396 |
| LYS | 26 | A | 0.762 | 0.745 | 0.922 | 0.585 | | ALYS | 91 | A | 0.836 | 0.766 | 0.963 | 0.639 |
| GLU | 27 | A | 0.813 | 0.746 | 0.905 | 0.654 | | LYS | 92 | A | 0.779 | 0.793 | 0.997 | 0.575 |
| LEU | 28 | A | 0.736 | 0.725 | 0.999 | 0.462 | | GLU | 93 | A | 0.718 | 0.799 | 0.971 | 0.546 |
| GLY | 29 | A | 0.641 | 0.648 | 0.999 | 0.29 | | ASN | 94 | A | 0.763 | 0.747 | 0.97 | 0.54 |
| **PHE** | **30** | **A** | **0.842** | **0.847** | **0.956** | **0.733** | | ILE | 95 | A | 0.78 | 0.775 | 0.977 | 0.578 |
| ASN | 31 | A | 0.746 | 0.727 | 0.99 | 0.483 | | ILE | 96 | A | 0.805 | 0.799 | 0.999 | 0.605 |
| ASN | 32 | A | 0.728 | 0.743 | 0.999 | 0.472 | | ALA | 97 | A | 0.621 | 0.597 | 0.999 | 0.219 |
| VAL | 33 | A | 0.696 | 0.694 | 1 | 0.39 | | ALA | 98 | A | 0.568 | 0.546 | 1 | 0.114 |
| GLU | 34 | A | 0.834 | 0.809 | 0.968 | 0.675 | | ALA | 99 | A | 0.627 | 0.595 | 0.998 | 0.224 |
| GLU | 35 | A | 0.75 | 0.752 | 1 | 0.502 | | GLN | 100 | A | 0.815 | 0.775 | 0.993 | 0.597 |
| ALA | 36 | A | 0.536 | 0.511 | 1 | 0.047 | | ALA | 101 | A | 0.481 | 0.427 | 0.999 | -0.091 |
| GLU | 37 | A | 0.729 | 0.652 | 0.994 | 0.387 | | GLY | 102 | A | 0.638 | 0.642 | 1 | 0.28 |
| ASP | 38 | A | 0.768 | 0.732 | 0.942 | 0.558 | | ALA | 103 | A | 0.473 | 0.466 | 1 | -0.061 |
| GLY | 39 | A | 0.621 | 0.619 | 1 | 0.24 | | SER | 104 | A | 0.642 | 0.651 | 1 | 0.293 |
| VAL | 40 | A | 0.708 | 0.687 | 1 | 0.395 | | GLY | 105 | A | 0.628 | 0.628 | 1 | 0.256 |
| ASP | 41 | A | 0.743 | 0.641 | 0.94 | 0.444 | | **TYR** | **106** | **A** | **0.768** | **0.802** | **0.741** | **0.829** |
| ALA | 42 | A | 0.566 | 0.576 | 1 | 0.142 | | VAL | 107 | A | 0.654 | 0.705 | 0.922 | 0.437 |
| LEU | 43 | A | 0.752 | 0.789 | 0.999 | 0.542 | | VAL | 108 | A | 0.667 | 0.63 | 0.996 | 0.301 |
| ASN | 44 | A | 0.731 | 0.767 | 0.996 | 0.502 | | LYS | 109 | A | 0.766 | 0.755 | 0.971 | 0.55 |
| LYS | 45 | A | 0.844 | 0.774 | 0.977 | 0.641 | | PRO | 110 | A | 0.706 | 0.699 | 0.997 | 0.408 |
| LEU | 46 | A | 0.737 | 0.747 | 1 | 0.484 | | PHE | 111 | A | 0.81 | 0.826 | 0.991 | 0.645 |
| GLN | 47 | A | 0.746 | 0.702 | 0.996 | 0.452 | | THR | 112 | A | 0.731 | 0.717 | 0.997 | 0.451 |
| ALA | 48 | A | 0.516 | 0.558 | 1 | 0.074 | | ALA | 113 | A | 0.602 | 0.653 | 0.989 | 0.266 |
| GLY | 49 | A | 0.643 | 0.648 | 1 | 0.291 | | ALA | 114 | A | 0.672 | 0.672 | 0.994 | 0.35 |
| GLY | 50 | A | 0.644 | 0.644 | 1 | 0.288 | | THR | 115 | A | 0.701 | 0.699 | 0.996 | 0.404 |
| TYR | 51 | A | 0.857 | 0.847 | 1 | 0.704 | | LEU | 116 | A | 0.723 | 0.715 | 0.989 | 0.449 |
| GLY | 52 | A | 0.63 | 0.635 | 1 | 0.265 | | GLU | 117 | A | 0.795 | 0.705 | 0.967 | 0.533 |
| PHE | 53 | A | 0.846 | 0.839 | 1 | 0.685 | | GLU | 118 | A | 0.645 | 0.726 | 0.982 | 0.389 |
| VAL | 54 | A | 0.703 | 0.707 | 1 | 0.41 | | **LYS** | **119** | **A** | **0.866** | **0.852** | **0.87** | **0.848** |
| ILE | 55 | A | 0.738 | 0.743 | 1 | 0.481 | | LEU | 120 | A | 0.732 | 0.75 | 0.989 | 0.493 |
| SER | 56 | A | 0.76 | 0.714 | 0.993 | 0.481 | | ASN | 121 | A | 0.747 | 0.787 | 0.997 | 0.537 |
| ASP | 57 | A | 0.785 | 0.836 | 0.969 | 0.652 | | LYS | 122 | A | 0.738 | 0.7 | 0.928 | 0.51 |
| TRP | 58 | A | 0.849 | 0.863 | 0.99 | 0.722 | | ILE | 123 | A | 0.746 | 0.743 | 0.991 | 0.498 |
| ASN | 59 | A | 0.655 | 0.679 | 0.975 | 0.359 | | PHE | 124 | A | 0.835 | 0.838 | 0.97 | 0.703 |
| MET | 60 | A | 0.753 | 0.729 | 0.999 | 0.483 | | GLU | 125 | A | 0.644 | 0.679 | 0.968 | 0.355 |
| PRO | 61 | A | 0.67 | 0.677 | 0.998 | 0.349 | | LYS | 126 | A | 0.611 | 0.717 | 0.987 | 0.341 |
| ASN | 62 | A | 0.74 | 0.7 | 1 | 0.44 | | LEU | 127 | A | 0.706 | 0.627 | 0.939 | 0.394 |
| **MET** | **63** | **A** | **0.878** | **0.87** | **0.953** | **0.795** | | GLY | 128 | A | 0.645 | 0.67 | 0.988 | 0.327 |
| ASP | 64 | A | 0.719 | 0.767 | 1 | 0.486 | | MET | 129 | A | 0.843 | 0.782 | 0.92 | 0.705 |

| Table S1(c): 1JBE (P), 1FQW (Q) | | | | | | | | | | | | | | |
|---|---|---|---|---|---|---|---|---|---|---|---|---|---|---|
| Residue | | | S (P) | S (Q) | J (P,Q) | PRI | | Residue | | | S (P) | S (Q) | J (P,Q) | PRI |
| ALA | 2 | A | 0.677 | 0.659 | 0.998 | 0.338 | | GLY | 65 | A | 0.624 | 0.617 | 1 | 0.241 |
| ASP | 3 | A | 0.744 | 0.801 | 0.962 | 0.583 | | LEU | 66 | A | 0.735 | 0.773 | 0.904 | 0.604 |
| LYS | 4 | A | 0.761 | 0.804 | 0.993 | 0.572 | | GLU | 67 | A | 0.748 | 0.702 | 0.862 | 0.588 |
| GLU | 5 | A | 0.807 | 0.768 | 0.941 | 0.634 | | LEU | 68 | A | 0.744 | 0.737 | 0.999 | 0.482 |
| LEU | 6 | A | 0.733 | 0.738 | 0.998 | 0.473 | | LEU | 69 | A | 0.742 | 0.741 | 1 | 0.483 |
| LYS | 7 | A | 0.828 | 0.851 | 0.989 | 0.69 | | LYS | 70 | A | 0.806 | 0.81 | 0.998 | 0.618 |
| PHE | 8 | A | 0.826 | 0.841 | 1 | 0.667 | | THR | 71 | A | 0.712 | 0.663 | 1 | 0.375 |
| LEU | 9 | A | 0.746 | 0.754 | 1 | 0.5 | | ILE | 72 | A | 0.739 | 0.741 | 0.998 | 0.482 |
| VAL | 10 | A | 0.679 | 0.686 | 1 | 0.365 | | ARG | 73 | A | 0.853 | 0.85 | 0.999 | 0.704 |
| VAL | 11 | A | 0.693 | 0.698 | 0.998 | 0.393 | | ALA | 74 | A | 0.724 | 0.71 | 1 | 0.434 |
| ASP | 12 | A | 0.799 | 0.781 | 0.974 | 0.606 | | ALA | 77 | A | 0.788 | 0.703 | 0.989 | 0.502 |
| ASP | 13 | A | 0.735 | 0.769 | 0.996 | 0.508 | | MET | 78 | A | 0.849 | 0.85 | 0.975 | 0.724 |
| PHE | 14 | A | 0.775 | 0.841 | 0.891 | 0.725 | | SER | 79 | A | 0.683 | 0.735 | 0.996 | 0.422 |
| SER | 15 | A | 0.736 | 0.774 | 0.997 | 0.513 | | ALA | 80 | A | 0.506 | 0.541 | 0.998 | 0.049 |
| THR | 16 | A | 0.676 | 0.664 | 0.839 | 0.501 | | LEU | 81 | A | 0.745 | 0.745 | 0.899 | 0.591 |
| MET | 17 | A | 0.712 | 0.768 | 0.933 | 0.547 | | PRO | 82 | A | 0.651 | 0.671 | 1 | 0.322 |
| ARG | 18 | A | 0.835 | 0.821 | 0.999 | 0.657 | | VAL | 83 | A | 0.704 | 0.693 | 0.999 | 0.398 |
| ARG | 19 | A | 0.746 | 0.713 | 0.911 | 0.548 | | LEU | 84 | A | 0.737 | 0.756 | 1 | 0.493 |
| ILE | 20 | A | 0.799 | 0.768 | 0.996 | 0.571 | | MET | 85 | A | 0.76 | 0.713 | 0.958 | 0.515 |
| VAL | 21 | A | 0.684 | 0.7 | 0.998 | 0.386 | | VAL | 86 | A | 0.698 | 0.697 | 0.969 | 0.426 |
| **ARG** | **22** | **A** | **0.853** | **0.855** | **0.788** | **0.92** | | THR | 87 | A | 0.726 | 0.704 | 0.924 | 0.506 |
| ASN | 23 | A | 0.733 | 0.769 | 0.991 | 0.511 | | AALA | 88 | A | 0.648 | 0.693 | 0.964 | 0.377 |
| LEU | 24 | A | 0.774 | 0.785 | 0.995 | 0.564 | | AGLU | 89 | A | 0.693 | 0.768 | 0.864 | 0.597 |
| LEU | 25 | A | 0.738 | 0.741 | 0.999 | 0.48 | | AALA | 90 | A | 0.662 | 0.672 | 0.922 | 0.412 |
| LYS | 26 | A | 0.762 | 0.67 | 0.849 | 0.583 | | ALYS | 91 | A | 0.836 | 0.744 | 0.918 | 0.662 |
| GLU | 27 | A | 0.813 | 0.737 | 0.86 | 0.69 | | LYS | 92 | A | 0.779 | 0.776 | 0.641 | 0.914 |
| LEU | 28 | A | 0.736 | 0.706 | 0.999 | 0.443 | | GLU | 93 | A | 0.718 | 0.764 | 0.817 | 0.665 |
| GLY | 29 | A | 0.641 | 0.648 | 0.999 | 0.29 | | ASN | 94 | A | 0.763 | 0.741 | 0.948 | 0.556 |
| PHE | 30 | A | 0.842 | 0.849 | 0.997 | 0.694 | | ILE | 95 | A | 0.78 | 0.709 | 1 | 0.489 |
| ASN | 31 | A | 0.746 | 0.713 | 0.987 | 0.472 | | ILE | 96 | A | 0.805 | 0.831 | 0.999 | 0.637 |
| ASN | 32 | A | 0.728 | 0.728 | 0.999 | 0.457 | | ALA | 97 | A | 0.621 | 0.574 | 1 | 0.195 |
| VAL | 33 | A | 0.696 | 0.71 | 1 | 0.406 | | ALA | 98 | A | 0.568 | 0.526 | 1 | 0.094 |
| GLU | 34 | A | 0.834 | 0.835 | 0.936 | 0.733 | | ALA | 99 | A | 0.627 | 0.642 | 0.999 | 0.27 |
| GLU | 35 | A | 0.75 | 0.721 | 1 | 0.471 | | GLN | 100 | A | 0.815 | 0.676 | 0.968 | 0.523 |
| ALA | 36 | A | 0.536 | 0.493 | 1 | 0.029 | | ALA | 101 | A | 0.481 | 0.513 | 0.999 | -0.005 |
| GLU | 37 | A | 0.729 | 0.695 | 0.946 | 0.478 | | GLY | 102 | A | 0.638 | 0.64 | 0.999 | 0.279 |
| ASP | 38 | A | 0.768 | 0.761 | 0.999 | 0.53 | | ALA | 103 | A | 0.473 | 0.502 | 1 | -0.025 |
| GLY | 39 | A | 0.621 | 0.618 | 1 | 0.239 | | SER | 104 | A | 0.642 | 0.631 | 1 | 0.273 |
| VAL | 40 | A | 0.708 | 0.632 | 0.999 | 0.341 | | GLY | 105 | A | 0.628 | 0.629 | 1 | 0.257 |
| ASP | 41 | A | 0.743 | 0.738 | 1 | 0.481 | | **TYR** | **106** | **A** | **0.768** | **0.792** | **0.694** | **0.866** |
| ALA | 42 | A | 0.566 | 0.577 | 1 | 0.143 | | VAL | 107 | A | 0.654 | 0.709 | 0.894 | 0.469 |
| LEU | 43 | A | 0.752 | 0.763 | 0.999 | 0.516 | | VAL | 108 | A | 0.667 | 0.652 | 0.9 | 0.419 |
| ASN | 44 | A | 0.73 | 0.755 | 0.968 | 0.517 | | LYS | 109 | A | 0.766 | 0.77 | 0.958 | 0.578 |
| LYS | 45 | A | 0.844 | 0.858 | 0.977 | 0.725 | | PRO | 110 | A | 0.706 | 0.706 | 0.997 | 0.415 |
| LEU | 46 | A | 0.737 | 0.738 | 1 | 0.475 | | PHE | 111 | A | 0.81 | 0.814 | 0.989 | 0.635 |
| GLN | 47 | A | 0.746 | 0.682 | 0.846 | 0.582 | | THR | 112 | A | 0.731 | 0.698 | 0.993 | 0.436 |
| ALA | 48 | A | 0.516 | 0.534 | 1 | 0.05 | | ALA | 113 | A | 0.602 | 0.663 | 0.987 | 0.278 |
| GLY | 49 | A | 0.643 | 0.645 | 1 | 0.288 | | ALA | 114 | A | 0.672 | 0.65 | 0.994 | 0.328 |
| GLY | 50 | A | 0.644 | 0.642 | 0.999 | 0.287 | | THR | 115 | A | 0.701 | 0.68 | 0.993 | 0.388 |
| TYR | 51 | A | 0.857 | 0.853 | 1 | 0.71 | | LEU | 116 | A | 0.723 | 0.717 | 0.991 | 0.449 |
| GLY | 52 | A | 0.63 | 0.641 | 1 | 0.271 | | GLU | 117 | A | 0.795 | 0.774 | 0.954 | 0.615 |
| PHE | 53 | A | 0.846 | 0.843 | 1 | 0.689 | | GLU | 118 | A | 0.645 | 0.727 | 0.966 | 0.406 |
| VAL | 54 | A | 0.703 | 0.713 | 1 | 0.416 | | **LYS** | **119** | **A** | **0.866** | **0.843** | **0.867** | **0.842** |
| ILE | 55 | A | 0.738 | 0.742 | 1 | 0.48 | | LEU | 120 | A | 0.732 | 0.749 | 0.988 | 0.493 |
| SER | 56 | A | 0.76 | 0.794 | 0.927 | 0.627 | | ASN | 121 | A | 0.747 | 0.807 | 0.996 | 0.558 |
| ASP | 57 | A | 0.785 | 0.842 | 0.955 | 0.672 | | LYS | 122 | A | 0.738 | 0.66 | 0.827 | 0.571 |
| TRP | 58 | A | 0.849 | 0.868 | 0.978 | 0.739 | | ILE | 123 | A | 0.746 | 0.756 | 0.999 | 0.503 |
| ASN | 59 | A | 0.655 | 0.684 | 0.999 | 0.34 | | PHE | 124 | A | 0.836 | 0.829 | 0.999 | 0.666 |
| MET | 60 | A | 0.753 | 0.743 | 0.999 | 0.497 | | GLU | 125 | A | 0.644 | 0.692 | 0.955 | 0.381 |
| PRO | 61 | A | 0.67 | 0.687 | 0.998 | 0.359 | | LYS | 126 | A | 0.611 | 0.519 | 0.646 | 0.484 |
| ASN | 62 | A | 0.74 | 0.731 | 0.999 | 0.472 | | LEU | 127 | A | 0.706 | 0.729 | 0.994 | 0.441 |
| **MET** | **63** | **A** | **0.878** | **0.878** | **0.927** | **0.829** | | GLY | 128 | A | 0.646 | 0.659 | 1 | 0.305 |
| ASP | 64 | A | 0.719 | 0.673 | 1 | 0.392 | | MET | 129 | A | 0.864 | 0.879 | 0.993 | 0.75 |

| Table S1(d): 1JBE (P), 2B1J (Q) | | | | | | | | | | | | | | |
|---|---|---|---|---|---|---|---|---|---|---|---|---|---|---|
| Residue | | | S (P) | S (Q) | J (P,Q) | PRI | | Residue | | | S (P) | S (Q) | J (P,Q) | PRI |
| ALA | 2 | A | 0.677 | 0.669 | 0.878 | 0.468 | | GLY | 65 | A | 0.624 | 0.613 | 1 | 0.237 |
| ASP | 3 | A | 0.743 | 0.791 | 0.855 | 0.679 | | LEU | 66 | A | 0.734 | 0.735 | 1 | 0.469 |
| LYS | 4 | A | 0.761 | 0.802 | 0.985 | 0.578 | | GLU | 67 | A | 0.748 | 0.76 | 0.804 | 0.704 |
| GLU | 5 | A | 0.807 | 0.661 | 0.799 | 0.669 | | LEU | 68 | A | 0.744 | 0.742 | 1 | 0.486 |
| LEU | 6 | A | 0.732 | 0.744 | 0.995 | 0.481 | | LEU | 69 | A | 0.742 | 0.743 | 1 | 0.485 |
| **LYS** | **7** | **A** | **0.828** | **0.843** | **0.837** | **0.834** | | LYS | 70 | A | 0.805 | 0.799 | 0.999 | 0.605 |
| PHE | 8 | A | 0.826 | 0.816 | 0.998 | 0.644 | | THR | 71 | A | 0.712 | 0.678 | 1 | 0.39 |
| LEU | 9 | A | 0.746 | 0.741 | 0.999 | 0.488 | | ILE | 72 | A | 0.738 | 0.763 | 1 | 0.501 |
| VAL | 10 | A | 0.679 | 0.697 | 0.999 | 0.377 | | ARG | 73 | A | 0.853 | 0.842 | 0.999 | 0.696 |
| VAL | 11 | A | 0.693 | 0.698 | 0.999 | 0.392 | | ALA | 74 | A | 0.724 | 0.737 | 0.997 | 0.464 |
| ASP | 12 | A | 0.796 | 0.763 | 0.976 | 0.583 | | ALA | 77 | A | 0.788 | 0.769 | 0.982 | 0.575 |
| ASP | 13 | A | 0.74 | 0.75 | 0.99 | 0.5 | | MET | 78 | A | 0.849 | 0.853 | 0.96 | 0.742 |
| PHE | 14 | A | 0.739 | 0.797 | 0.831 | 0.705 | | SER | 79 | A | 0.683 | 0.69 | 0.979 | 0.394 |
| SER | 15 | A | 0.738 | 0.775 | 0.902 | 0.611 | | ALA | 80 | A | 0.505 | 0.522 | 0.998 | 0.029 |
| THR | 16 | A | 0.675 | 0.735 | 0.828 | 0.582 | | LEU | 81 | A | 0.745 | 0.69 | 0.86 | 0.575 |
| MET | 17 | A | 0.716 | 0.706 | 0.999 | 0.423 | | PRO | 82 | A | 0.651 | 0.655 | 1 | 0.306 |
| ARG | 18 | A | 0.835 | 0.827 | 0.996 | 0.666 | | VAL | 83 | A | 0.704 | 0.703 | 0.999 | 0.408 |
| ARG | 19 | A | 0.745 | 0.726 | 0.786 | 0.685 | | LEU | 84 | A | 0.737 | 0.72 | 0.999 | 0.458 |
| ILE | 20 | A | 0.798 | 0.78 | 1 | 0.578 | | MET | 85 | A | 0.76 | 0.691 | 0.924 | 0.527 |
| VAL | 21 | A | 0.684 | 0.707 | 1 | 0.391 | | VAL | 86 | A | 0.7 | 0.701 | 0.962 | 0.439 |
| **ARG** | **22** | **A** | **0.853** | **0.845** | **0.752** | **0.946** | | THR | 87 | A | 0.724 | 0.778 | 0.955 | 0.547 |
| ASN | 23 | A | 0.732 | 0.802 | 0.954 | 0.58 | | AALA | 88 | A | 0.747 | 0.823 | 0.963 | 0.607 |
| LEU | 24 | A | 0.778 | 0.766 | 1 | 0.544 | | **AGLU** | **89** | **A** | **0.771** | **0.864** | **0.727** | **0.908** |
| LEU | 25 | A | 0.739 | 0.73 | 1 | 0.469 | | AALA | 90 | A | 0.749 | 0.829 | 0.948 | 0.63 |
| **LYS** | **26** | **A** | **0.764** | **0.785** | **0.747** | **0.802** | | ALYS | 91 | A | 0.834 | 0.828 | 0.952 | 0.71 |
| **GLU** | **27** | **A** | **0.898** | **0.888** | **0.954** | **0.832** | | LYS | 92 | A | 0.778 | 0.738 | 0.932 | 0.584 |
| LEU | 28 | A | 0.735 | 0.731 | 0.999 | 0.467 | | GLU | 93 | A | 0.712 | 0.748 | 0.866 | 0.594 |
| GLY | 29 | A | 0.64 | 0.645 | 1 | 0.285 | | ASN | 94 | A | 0.757 | 0.709 | 0.992 | 0.474 |
| PHE | 30 | A | 0.842 | 0.831 | 0.999 | 0.674 | | ILE | 95 | A | 0.77 | 0.76 | 0.817 | 0.713 |
| ASN | 31 | A | 0.746 | 0.756 | 0.997 | 0.505 | | ILE | 96 | A | 0.804 | 0.749 | 0.965 | 0.588 |
| ASN | 32 | A | 0.728 | 0.725 | 0.995 | 0.458 | | ALA | 97 | A | 0.62 | 0.65 | 0.998 | 0.272 |
| VAL | 33 | A | 0.696 | 0.695 | 0.998 | 0.393 | | ALA | 98 | A | 0.566 | 0.601 | 1 | 0.167 |
| GLU | 34 | A | 0.834 | 0.761 | 0.861 | 0.734 | | ALA | 99 | A | 0.626 | 0.631 | 0.998 | 0.259 |
| GLU | 35 | A | 0.749 | 0.734 | 0.999 | 0.484 | | GLN | 100 | A | 0.814 | 0.692 | 0.935 | 0.571 |
| ALA | 36 | A | 0.537 | 0.491 | 0.997 | 0.031 | | ALA | 101 | A | 0.48 | 0.431 | 0.998 | -0.087 |
| GLU | 37 | A | 0.727 | 0.692 | 0.965 | 0.454 | | GLY | 102 | A | 0.638 | 0.644 | 0.998 | 0.284 |
| ASP | 38 | A | 0.768 | 0.752 | 0.998 | 0.522 | | ALA | 103 | A | 0.472 | 0.449 | 1 | -0.079 |
| GLY | 39 | A | 0.621 | 0.62 | 1 | 0.241 | | SER | 104 | A | 0.643 | 0.613 | 1 | 0.256 |
| VAL | 40 | A | 0.708 | 0.654 | 1 | 0.362 | | GLY | 105 | A | 0.628 | 0.632 | 1 | 0.26 |
| ASP | 41 | A | 0.743 | 0.759 | 1 | 0.502 | | **TYR** | **106** | **A** | **0.769** | **0.818** | **0.614** | **0.973** |
| ALA | 42 | A | 0.566 | 0.572 | 1 | 0.138 | | VAL | 107 | A | 0.656 | 0.74 | 0.79 | 0.606 |
| LEU | 43 | A | 0.752 | 0.744 | 1 | 0.496 | | VAL | 108 | A | 0.651 | 0.693 | 0.999 | 0.345 |
| ASN | 44 | A | 0.73 | 0.78 | 0.993 | 0.517 | | LYS | 109 | A | 0.763 | 0.776 | 0.864 | 0.675 |
| LYS | 45 | A | 0.844 | 0.852 | 0.954 | 0.742 | | PRO | 110 | A | 0.702 | 0.694 | 0.998 | 0.398 |
| LEU | 46 | A | 0.737 | 0.725 | 1 | 0.462 | | PHE | 111 | A | 0.81 | 0.82 | 0.997 | 0.633 |
| GLN | 47 | A | 0.746 | 0.679 | 0.983 | 0.442 | | THR | 112 | A | 0.729 | 0.748 | 0.998 | 0.479 |
| ALA | 48 | A | 0.516 | 0.521 | 0.987 | 0.05 | | ALA | 113 | A | 0.576 | 0.548 | 0.995 | 0.129 |
| GLY | 49 | A | 0.643 | 0.65 | 0.997 | 0.296 | | ALA | 114 | A | 0.67 | 0.713 | 0.998 | 0.385 |
| GLY | 50 | A | 0.644 | 0.646 | 0.999 | 0.291 | | THR | 115 | A | 0.7 | 0.64 | 0.998 | 0.342 |
| TYR | 51 | A | 0.857 | 0.838 | 0.997 | 0.698 | | LEU | 116 | A | 0.723 | 0.727 | 0.999 | 0.451 |
| GLY | 52 | A | 0.63 | 0.636 | 1 | 0.266 | | GLU | 117 | A | 0.794 | 0.746 | 0.904 | 0.636 |
| PHE | 53 | A | 0.846 | 0.838 | 0.999 | 0.685 | | GLU | 118 | A | 0.643 | 0.712 | 0.938 | 0.417 |
| VAL | 54 | A | 0.703 | 0.71 | 1 | 0.413 | | **LYS** | **119** | **A** | **0.865** | **0.881** | **0.909** | **0.837** |
| ILE | 55 | A | 0.738 | 0.732 | 0.999 | 0.471 | | LEU | 120 | A | 0.731 | 0.742 | 0.999 | 0.474 |
| SER | 56 | A | 0.76 | 0.782 | 0.884 | 0.658 | | ASN | 121 | A | 0.747 | 0.714 | 0.983 | 0.478 |
| ASP | 57 | A | 0.783 | 0.789 | 0.918 | 0.654 | | LYS | 122 | A | 0.737 | 0.677 | 0.835 | 0.579 |
| TRP | 58 | A | 0.847 | 0.862 | 0.986 | 0.723 | | ILE | 123 | A | 0.746 | 0.755 | 0.999 | 0.502 |
| ASN | 59 | A | 0.648 | 0.702 | 0.907 | 0.443 | | PHE | 124 | A | 0.836 | 0.828 | 0.997 | 0.667 |
| MET | 60 | A | 0.752 | 0.75 | 0.999 | 0.503 | | GLU | 125 | A | 0.643 | 0.65 | 0.829 | 0.464 |
| PRO | 61 | A | 0.669 | 0.682 | 1 | 0.351 | | LYS | 126 | A | 0.611 | 0.73 | 0.563 | 0.778 |
| ASN | 62 | A | 0.739 | 0.681 | 0.995 | 0.425 | | LEU | 127 | A | 0.706 | 0.7 | 0.917 | 0.489 |
| **MET** | **63** | **A** | **0.878** | **0.873** | **0.897** | **0.854** | | GLY | 128 | A | 0.646 | 0.66 | 0.998 | 0.308 |
| ASP | 64 | A | 0.718 | 0.601 | 0.997 | 0.322 | | MET | 129 | A | 0.864 | 0.806 | 0.967 | 0.703 |

**Table S1(e): 1JBE (P), 3CHY (Q)**

| Residue | | | S (P) | S (Q) | J (P,Q) | PRI |
|---|---|---|---|---|---|---|
| ALA | 2 | A | 0.677 | 0.525 | 0.872 | 0.33 |
| ASP | 3 | A | 0.744 | 0.772 | 0.979 | 0.537 |
| LYS | 4 | A | 0.761 | 0.78 | 0.939 | 0.602 |
| GLU | 5 | A | 0.806 | 0.807 | 0.959 | 0.654 |
| LEU | 6 | A | 0.732 | 0.723 | 0.991 | 0.464 |
| **LYS** | **7** | **A** | **0.827** | **0.86** | **0.813** | **0.874** |
| PHE | 8 | A | 0.826 | 0.822 | 0.999 | 0.649 |
| LEU | 9 | A | 0.746 | 0.75 | 0.997 | 0.499 |
| VAL | 10 | A | 0.679 | 0.689 | 0.997 | 0.371 |
| VAL | 11 | A | 0.692 | 0.699 | 0.998 | 0.393 |
| ASP | 12 | A | 0.799 | 0.801 | 0.998 | 0.602 |
| ASP | 13 | A | 0.735 | 0.738 | 0.999 | 0.474 |
| PHE | 14 | A | 0.775 | 0.783 | 0.997 | 0.561 |
| SER | 15 | A | 0.736 | 0.704 | 0.993 | 0.447 |
| THR | 16 | A | 0.675 | 0.731 | 0.581 | 0.825 |
| MET | 17 | A | 0.712 | 0.705 | 0.999 | 0.418 |
| ARG | 18 | A | 0.835 | 0.845 | 0.994 | 0.686 |
| **ARG** | **19** | **A** | **0.748** | **0.766** | **0.658** | **0.856** |
| ILE | 20 | A | 0.8 | 0.785 | 0.997 | 0.588 |
| VAL | 21 | A | 0.684 | 0.686 | 0.999 | 0.371 |
| **ARG** | **22** | **A** | **0.853** | **0.885** | **0.55** | **1.188** |
| ASN | 23 | A | 0.81 | 0.826 | 0.823 | 0.813 |
| LEU | 24 | A | 0.774 | 0.773 | 0.998 | 0.549 |
| LEU | 25 | A | 0.738 | 0.732 | 0.999 | 0.471 |
| LYS | 26 | A | 0.76 | 0.755 | 0.711 | 0.804 |
| GLU | 27 | A | 0.812 | 0.821 | 0.857 | 0.776 |
| LEU | 28 | A | 0.735 | 0.72 | 0.972 | 0.483 |
| GLY | 29 | A | 0.641 | 0.638 | 0.999 | 0.28 |
| PHE | 30 | A | 0.842 | 0.836 | 0.997 | 0.681 |
| ASN | 31 | A | 0.746 | 0.767 | 0.997 | 0.516 |
| ASN | 32 | A | 0.686 | 0.692 | 0.586 | 0.792 |
| VAL | 33 | A | 0.696 | 0.689 | 0.999 | 0.386 |
| GLU | 34 | A | 0.834 | 0.773 | 0.979 | 0.628 |
| GLU | 35 | A | 0.75 | 0.758 | 0.987 | 0.521 |
| ALA | 36 | A | 0.536 | 0.523 | 0.999 | 0.06 |
| GLU | 37 | A | 0.781 | 0.759 | 0.77 | 0.77 |
| ASP | 38 | A | 0.768 | 0.766 | 0.999 | 0.535 |
| GLY | 39 | A | 0.62 | 0.623 | 1 | 0.243 |
| VAL | 40 | A | 0.708 | 0.719 | 0.998 | 0.429 |
| ASP | 41 | A | 0.744 | 0.752 | 0.999 | 0.497 |
| ALA | 42 | A | 0.565 | 0.551 | 0.999 | 0.117 |
| LEU | 43 | A | 0.752 | 0.764 | 0.999 | 0.517 |
| ASN | 44 | A | 0.73 | 0.714 | 0.999 | 0.445 |
| LYS | 45 | A | 0.844 | 0.843 | 0.944 | 0.743 |
| LEU | 46 | A | 0.737 | 0.725 | 1 | 0.462 |
| GLN | 47 | A | 0.746 | 0.7 | 0.831 | 0.615 |
| ALA | 48 | A | 0.516 | 0.586 | 0.998 | 0.104 |
| GLY | 49 | A | 0.643 | 0.65 | 0.999 | 0.294 |
| GLY | 50 | A | 0.644 | 0.635 | 0.999 | 0.28 |
| TYR | 51 | A | 0.857 | 0.852 | 0.992 | 0.717 |
| GLY | 52 | A | 0.63 | 0.636 | 0.999 | 0.267 |
| PHE | 53 | A | 0.846 | 0.836 | 0.999 | 0.683 |
| VAL | 54 | A | 0.703 | 0.703 | 0.998 | 0.408 |
| ILE | 55 | A | 0.739 | 0.75 | 0.997 | 0.492 |
| **SER** | **56** | **A** | **0.825** | **0.857** | **0.678** | **1.004** |
| ASP | 57 | A | 0.788 | 0.783 | 0.998 | 0.573 |
| TRP | 58 | A | 0.849 | 0.843 | 0.999 | 0.693 |
| ASN | 59 | A | 0.655 | 0.629 | 1 | 0.284 |
| MET | 60 | A | 0.751 | 0.753 | 0.998 | 0.506 |
| PRO | 61 | A | 0.669 | 0.667 | 1 | 0.336 |
| ASN | 62 | A | 0.739 | 0.748 | 1 | 0.487 |
| **MET** | **63** | **A** | **0.878** | **0.88** | **0.8** | **0.958** |
| ASP | 64 | A | 0.719 | 0.638 | 0.999 | 0.358 |

| Residue | | | S (P) | S (Q) | J (P,Q) | PRI |
|---|---|---|---|---|---|---|
| GLY | 65 | A | 0.62 | 0.609 | 0.997 | 0.232 |
| LEU | 66 | A | 0.735 | 0.735 | 0.999 | 0.471 |
| GLU | 67 | A | 0.748 | 0.812 | 0.791 | 0.769 |
| LEU | 68 | A | 0.744 | 0.74 | 0.999 | 0.485 |
| LEU | 69 | A | 0.741 | 0.736 | 0.999 | 0.478 |
| LYS | 70 | A | 0.805 | 0.78 | 0.999 | 0.586 |
| THR | 71 | A | 0.712 | 0.702 | 0.999 | 0.415 |
| ILE | 72 | A | 0.738 | 0.738 | 0.995 | 0.481 |
| ARG | 73 | A | 0.853 | 0.846 | 0.998 | 0.701 |
| ALA | 74 | A | 0.724 | 0.738 | 0.982 | 0.48 |
| ALA | 77 | A | 0.788 | 0.751 | 0.966 | 0.573 |
| MET | 78 | A | 0.849 | 0.831 | 0.938 | 0.742 |
| SER | 79 | A | 0.683 | 0.703 | 0.996 | 0.39 |
| ALA | 80 | A | 0.506 | 0.534 | 0.987 | 0.053 |
| LEU | 81 | A | 0.745 | 0.691 | 0.703 | 0.733 |
| PRO | 82 | A | 0.651 | 0.659 | 0.999 | 0.311 |
| VAL | 83 | A | 0.702 | 0.708 | 0.999 | 0.411 |
| LEU | 84 | A | 0.738 | 0.727 | 0.999 | 0.466 |
| **MET** | **85** | **A** | **0.858** | **0.826** | **0.789** | **0.895** |
| VAL | 86 | A | 0.699 | 0.716 | 0.993 | 0.422 |
| THR | 87 | A | 0.724 | 0.713 | 0.993 | 0.444 |
| AALA | 88 | A | 0.647 | 0.689 | 0.999 | 0.337 |
| AGLU | 89 | A | 0.694 | 0.775 | 0.978 | 0.491 |
| AALA | 90 | A | 0.656 | 0.659 | 0.768 | 0.547 |
| ALYS | 91 | A | 0.836 | 0.823 | 0.967 | 0.692 |
| LYS | 92 | A | 0.778 | 0.799 | 0.969 | 0.608 |
| GLU | 93 | A | 0.718 | 0.783 | 0.986 | 0.515 |
| ASN | 94 | A | 0.763 | 0.763 | 0.991 | 0.535 |
| ILE | 95 | A | 0.775 | 0.754 | 0.998 | 0.531 |
| ILE | 96 | A | 0.805 | 0.81 | 0.999 | 0.616 |
| ALA | 97 | A | 0.62 | 0.612 | 0.999 | 0.233 |
| ALA | 98 | A | 0.558 | 0.557 | 0.999 | 0.116 |
| ALA | 99 | A | 0.625 | 0.638 | 0.999 | 0.264 |
| GLN | 100 | A | 0.815 | 0.809 | 0.997 | 0.627 |
| ALA | 101 | A | 0.48 | 0.462 | 0.999 | -0.057 |
| GLY | 102 | A | 0.638 | 0.638 | 1 | 0.276 |
| ALA | 103 | A | 0.467 | 0.473 | 1 | -0.06 |
| SER | 104 | A | 0.644 | 0.628 | 0.999 | 0.273 |
| GLY | 105 | A | 0.627 | 0.618 | 0.999 | 0.246 |
| **TYR** | **106** | **A** | **0.819** | **0.885** | **0.606** | **1.098** |
| VAL | 107 | A | 0.656 | 0.68 | 0.997 | 0.339 |
| VAL | 108 | A | 0.661 | 0.66 | 0.999 | 0.322 |
| LYS | 109 | A | 0.766 | 0.758 | 0.991 | 0.533 |
| PRO | 110 | A | 0.706 | 0.7 | 0.998 | 0.408 |
| PHE | 111 | A | 0.81 | 0.813 | 0.999 | 0.624 |
| THR | 112 | A | 0.731 | 0.739 | 0.997 | 0.473 |
| ALA | 113 | A | 0.602 | 0.611 | 0.999 | 0.214 |
| ALA | 114 | A | 0.672 | 0.654 | 0.999 | 0.327 |
| THR | 115 | A | 0.701 | 0.709 | 0.999 | 0.411 |
| LEU | 116 | A | 0.723 | 0.717 | 0.997 | 0.443 |
| GLU | 117 | A | 0.795 | 0.768 | 0.768 | 0.795 |
| GLU | 118 | A | 0.644 | 0.705 | 0.762 | 0.587 |
| **LYS** | **119** | **A** | **0.866** | **0.833** | **0.768** | **0.931** |
| LEU | 120 | A | 0.732 | 0.707 | 0.999 | 0.44 |
| ASN | 121 | A | 0.747 | 0.764 | 0.999 | 0.512 |
| LYS | 122 | A | 0.736 | 0.671 | 0.938 | 0.469 |
| ILE | 123 | A | 0.746 | 0.742 | 0.998 | 0.49 |
| PHE | 124 | A | 0.837 | 0.846 | 0.996 | 0.687 |
| GLU | 125 | A | 0.643 | 0.658 | 0.988 | 0.313 |
| LYS | 126 | A | 0.611 | 0.623 | 1 | 0.234 |
| LEU | 127 | A | 0.697 | 0.684 | 0.996 | 0.385 |
| GLY | 128 | A | 0.646 | 0.639 | 1 | 0.285 |
| MET | 129 | A | 0.864 | 0.813 | 0.996 | 0.681 |

| Table S1(f): 2B1J (P), 1C4W (Q) | | | | | | | | | | | | | | |
|---|---|---|---|---|---|---|---|---|---|---|---|---|---|---|
| Residue | | | S (P) | S (Q) | J (P,Q) | PRI | | Residue | | | S (P) | S (Q) | J (P,Q) | PRI |
| ALA | 2 | A | 0.669 | 0.65 | 0.894 | 0.425 | | LEU | 68 | A | 0.743 | 0.758 | 1 | 0.501 |
| ASP | 3 | A | 0.791 | 0.766 | 0.787 | 0.77 | | LEU | 69 | A | 0.743 | 0.727 | 0.999 | 0.471 |
| LYS | 4 | A | 0.803 | 0.74 | 0.926 | 0.617 | | LYS | 70 | A | 0.8 | 0.797 | 1 | 0.597 |
| GLU | 5 | A | 0.663 | 0.764 | 0.733 | 0.694 | | THR | 71 | A | 0.685 | 0.724 | 1 | 0.409 |
| LEU | 6 | A | 0.744 | 0.742 | 1 | 0.486 | | ILE | 72 | A | 0.77 | 0.762 | 1 | 0.532 |
| **LYS** | **7** | **A** | **0.799** | **0.799** | **0.759** | **0.839** | | ARG | 73 | A | 0.845 | 0.856 | 0.999 | 0.702 |
| PHE | 8 | A | 0.816 | 0.824 | 0.999 | 0.641 | | ALA | 74 | A | 0.511 | 0.49 | 0.999 | 0.002 |
| LEU | 9 | A | 0.742 | 0.746 | 0.998 | 0.49 | | ASP | 75 | A | 0.755 | 0.674 | 0.999 | 0.43 |
| VAL | 10 | A | 0.698 | 0.687 | 1 | 0.385 | | GLY | 76 | A | 0.661 | 0.652 | 0.998 | 0.315 |
| VAL | 11 | A | 0.693 | 0.691 | 0.998 | 0.386 | | ALA | 77 | A | 0.593 | 0.559 | 0.989 | 0.163 |
| ASP | 12 | A | 0.764 | 0.782 | 0.999 | 0.547 | | MET | 78 | A | 0.737 | 0.717 | 0.924 | 0.53 |
| ASP | 13 | A | 0.752 | 0.778 | 0.993 | 0.537 | | SER | 79 | A | 0.675 | 0.745 | 0.964 | 0.456 |
| **APHE** | **14** | **A** | **0.799** | **0.72** | **0.705** | **0.814** | | ALA | 80 | A | 0.518 | 0.515 | 0.976 | 0.057 |
| SER | 15 | A | 0.776 | 0.738 | 0.901 | 0.613 | | LEU | 81 | A | 0.692 | 0.72 | 0.999 | 0.413 |
| THR | 16 | A | 0.737 | 0.707 | 0.994 | 0.45 | | PRO | 82 | A | 0.655 | 0.693 | 1 | 0.348 |
| MET | 17 | A | 0.71 | 0.75 | 0.998 | 0.462 | | VAL | 83 | A | 0.705 | 0.706 | 1 | 0.411 |
| ARG | 18 | A | 0.826 | 0.851 | 0.995 | 0.682 | | LEU | 84 | A | 0.72 | 0.737 | 0.998 | 0.459 |
| **ARG** | **19** | **A** | **0.726** | **0.818** | **0.698** | **0.846** | | MET | 85 | A | 0.689 | 0.769 | 0.8 | 0.658 |
| ILE | 20 | A | 0.78 | 0.78 | 0.997 | 0.563 | | VAL | 86 | A | 0.694 | 0.7 | 0.962 | 0.432 |
| VAL | 21 | A | 0.707 | 0.693 | 0.998 | 0.402 | | THR | 87 | A | 0.779 | 0.749 | 0.929 | 0.599 |
| ARG | 22 | A | 0.801 | 0.842 | 0.995 | 0.648 | | AALA | 88 | A | 0.75 | 0.702 | 0.854 | 0.598 |
| ASN | 23 | A | 0.749 | 0.759 | 0.906 | 0.602 | | **AGLU** | **89** | **A** | **0.823** | **0.748** | **0.561** | **1.01** |
| LEU | 24 | A | 0.768 | 0.771 | 0.997 | 0.542 | | AALA | 90 | A | 0.764 | 0.648 | 0.964 | 0.448 |
| LEU | 25 | A | 0.729 | 0.73 | 1 | 0.459 | | LYS | 91 | A | 0.768 | 0.754 | 0.831 | 0.691 |
| LYS | 26 | A | 0.788 | 0.702 | 0.939 | 0.551 | | LYS | 92 | A | 0.734 | 0.795 | 0.842 | 0.687 |
| GLU | 27 | A | 0.797 | 0.826 | 0.92 | 0.703 | | GLU | 93 | A | 0.756 | 0.78 | 0.797 | 0.739 |
| LEU | 28 | A | 0.73 | 0.734 | 0.998 | 0.466 | | ASN | 94 | A | 0.72 | 0.777 | 0.99 | 0.507 |
| GLY | 29 | A | 0.646 | 0.633 | 0.999 | 0.28 | | ILE | 95 | A | 0.77 | 0.783 | 0.776 | 0.777 |
| PHE | 30 | A | 0.831 | 0.83 | 0.998 | 0.663 | | ILE | 96 | A | 0.753 | 0.808 | 0.946 | 0.615 |
| ASN | 31 | A | 0.756 | 0.758 | 0.996 | 0.518 | | ALA | 97 | A | 0.653 | 0.601 | 1 | 0.254 |
| ASN | 32 | A | 0.723 | 0.74 | 0.991 | 0.472 | | ALA | 98 | A | 0.601 | 0.58 | 1 | 0.181 |
| VAL | 33 | A | 0.7 | 0.718 | 0.998 | 0.42 | | ALA | 99 | A | 0.634 | 0.642 | 0.993 | 0.283 |
| GLU | 34 | A | 0.763 | 0.832 | 0.813 | 0.782 | | GLN | 100 | A | 0.694 | 0.82 | 0.877 | 0.637 |
| GLU | 35 | A | 0.754 | 0.76 | 0.999 | 0.515 | | ALA | 101 | A | 0.432 | 0.423 | 0.999 | -0.144 |
| ALA | 36 | A | 0.493 | 0.473 | 0.999 | -0.033 | | GLY | 102 | A | 0.644 | 0.636 | 1 | 0.28 |
| GLU | 37 | A | 0.693 | 0.784 | 0.879 | 0.598 | | ALA | 103 | A | 0.452 | 0.497 | 1 | -0.051 |
| ASP | 38 | A | 0.753 | 0.783 | 0.998 | 0.538 | | SER | 104 | A | 0.612 | 0.636 | 0.999 | 0.249 |
| GLY | 39 | A | 0.623 | 0.619 | 1 | 0.242 | | GLY | 105 | A | 0.63 | 0.619 | 0.999 | 0.25 |
| VAL | 40 | A | 0.659 | 0.728 | 1 | 0.387 | | **TYR** | **106** | **A** | **0.853** | **0.816** | **0.85** | **0.819** |
| ASP | 41 | A | 0.758 | 0.761 | 0.998 | 0.521 | | VAL | 107 | A | 0.737 | 0.68 | 0.726 | 0.691 |
| ALA | 42 | A | 0.575 | 0.574 | 0.999 | 0.15 | | VAL | 108 | A | 0.709 | 0.725 | 0.996 | 0.438 |
| LEU | 43 | A | 0.744 | 0.772 | 1 | 0.516 | | **LYS** | **109** | **A** | **0.776** | **0.796** | **0.63** | **0.942** |
| ASN | 44 | A | 0.78 | 0.737 | 0.996 | 0.521 | | PRO | 110 | A | 0.699 | 0.69 | 0.998 | 0.391 |
| LYS | 45 | A | 0.804 | 0.81 | 0.967 | 0.647 | | PHE | 111 | A | 0.821 | 0.811 | 0.982 | 0.65 |
| LEU | 46 | A | 0.725 | 0.75 | 0.999 | 0.476 | | THR | 112 | A | 0.751 | 0.751 | 0.999 | 0.503 |
| GLN | 47 | A | 0.674 | 0.721 | 0.949 | 0.446 | | ALA | 113 | A | 0.568 | 0.563 | 0.999 | 0.132 |
| ALA | 48 | A | 0.523 | 0.485 | 0.983 | 0.025 | | ALA | 114 | A | 0.715 | 0.659 | 0.994 | 0.38 |
| GLY | 49 | A | 0.651 | 0.637 | 0.999 | 0.289 | | THR | 115 | A | 0.649 | 0.712 | 0.996 | 0.365 |
| GLY | 50 | A | 0.647 | 0.633 | 1 | 0.28 | | LEU | 116 | A | 0.727 | 0.712 | 1 | 0.439 |
| TYR | 51 | A | 0.838 | 0.871 | 0.998 | 0.711 | | GLU | 117 | A | 0.748 | 0.79 | 0.996 | 0.542 |
| GLY | 52 | A | 0.636 | 0.63 | 1 | 0.266 | | GLU | 118 | A | 0.715 | 0.667 | 0.997 | 0.385 |
| PHE | 53 | A | 0.838 | 0.851 | 0.999 | 0.69 | | LYS | 119 | A | 0.788 | 0.757 | 0.861 | 0.684 |
| VAL | 54 | A | 0.711 | 0.699 | 1 | 0.41 | | LEU | 120 | A | 0.741 | 0.735 | 0.999 | 0.477 |
| ILE | 55 | A | 0.729 | 0.742 | 0.998 | 0.473 | | ASN | 121 | A | 0.714 | 0.768 | 0.953 | 0.529 |
| SER | 56 | A | 0.81 | 0.787 | 1 | 0.597 | | LYS | 122 | A | 0.682 | 0.696 | 0.76 | 0.618 |
| TRP | 58 | A | 0.883 | 0.874 | 0.985 | 0.772 | | ILE | 123 | A | 0.757 | 0.76 | 0.997 | 0.52 |
| ASN | 59 | A | 0.715 | 0.715 | 0.848 | 0.582 | | PHE | 124 | A | 0.829 | 0.841 | 0.997 | 0.673 |
| MET | 60 | A | 0.755 | 0.736 | 0.999 | 0.492 | | GLU | 125 | A | 0.652 | 0.648 | 0.973 | 0.327 |
| PRO | 61 | A | 0.683 | 0.682 | 0.999 | 0.366 | | **LYS** | **126** | **A** | **0.731** | **0.711** | **0.554** | **0.888** |
| ASN | 62 | A | 0.677 | 0.75 | 0.986 | 0.441 | | LEU | 127 | A | 0.701 | 0.735 | 0.835 | 0.601 |
| MET | 63 | A | 0.763 | 0.758 | 0.993 | 0.528 | | GLY | 128 | A | 0.66 | 0.656 | 0.992 | 0.324 |
| ASP | 64 | A | 0.603 | 0.601 | 0.996 | 0.208 | | MET | 129 | A | 0.806 | 0.886 | 0.959 | 0.733 |
| GLY | 65 | A | 0.621 | 0.624 | 1 | 0.245 | | | | | | | | |
| LEU | 66 | A | 0.735 | 0.741 | 1 | 0.476 | | | | | | | | |
| GLU | 67 | A | 0.76 | 0.818 | 0.82 | 0.758 | | | | | | | | |

| Table S1(g): 2B1J (P), 1F4V (Q) | | | | | | | | | | | | | | |
|---|---|---|---|---|---|---|---|---|---|---|---|---|---|---|
| Residue | | | S (P) | S (Q) | J (P,Q) | PRI | | Residue | | | S (P) | S (Q) | J (P,Q) | PRI |
| ALA | 2 | A | 0.67 | 0.627 | 0.905 | 0.392 | | GLU | 67 | A | 0.76 | 0.708 | 0.961 | 0.507 |
| ASP | 3 | A | 0.791 | 0.735 | 0.998 | 0.528 | | LEU | 68 | A | 0.742 | 0.725 | 0.878 | 0.589 |
| LYS | 4 | A | 0.803 | 0.79 | 0.998 | 0.595 | | LEU | 69 | A | 0.743 | 0.742 | 1 | 0.485 |
| GLU | 5 | A | 0.663 | 0.769 | 0.989 | 0.443 | | LYS | 70 | A | 0.8 | 0.81 | 1 | 0.61 |
| LEU | 6 | A | 0.744 | 0.721 | 0.969 | 0.496 | | THR | 71 | A | 0.685 | 0.779 | 0.887 | 0.577 |
| LYS | 7 | A | 0.799 | 0.809 | 0.965 | 0.643 | | ILE | 72 | A | 0.77 | 0.765 | 1 | 0.535 |
| PHE | 8 | A | 0.816 | 0.845 | 0.997 | 0.664 | | ARG | 73 | A | 0.845 | 0.851 | 1 | 0.696 |
| LEU | 9 | A | 0.742 | 0.742 | 1 | 0.484 | | ALA | 74 | A | 0.511 | 0.476 | 1 | -0.013 |
| VAL | 10 | A | 0.698 | 0.679 | 1 | 0.377 | | ASP | 75 | A | 0.755 | 0.802 | 0.998 | 0.559 |
| VAL | 11 | A | 0.699 | 0.701 | 1 | 0.4 | | GLY | 76 | A | 0.661 | 0.663 | 0.94 | 0.384 |
| ASP | 12 | A | 0.764 | 0.77 | 0.999 | 0.535 | | ALA | 77 | A | 0.593 | 0.556 | 0.996 | 0.153 |
| ASP | 13 | A | 0.751 | 0.76 | 1 | 0.511 | | MET | 78 | A | 0.737 | 0.727 | 0.965 | 0.499 |
| **APHE** | **14** | **A** | **0.798** | **0.797** | **0.835** | **0.76** | | SER | 79 | A | 0.675 | 0.692 | 0.992 | 0.375 |
| SER | 15 | A | 0.776 | 0.711 | 0.96 | 0.527 | | ALA | 80 | A | 0.518 | 0.506 | 1 | 0.024 |
| THR | 16 | A | 0.736 | 0.652 | 0.983 | 0.405 | | LEU | 81 | A | 0.692 | 0.729 | 0.999 | 0.422 |
| MET | 17 | A | 0.709 | 0.758 | 0.825 | 0.642 | | PRO | 82 | A | 0.655 | 0.653 | 1 | 0.308 |
| ARG | 18 | A | 0.826 | 0.829 | 0.999 | 0.656 | | VAL | 83 | A | 0.704 | 0.683 | 1 | 0.387 |
| ARG | 19 | A | 0.676 | 0.861 | 0.966 | 0.571 | | LEU | 84 | A | 0.72 | 0.73 | 1 | 0.45 |
| ILE | 20 | A | 0.78 | 0.753 | 0.995 | 0.538 | | MET | 85 | A | 0.689 | 0.706 | 0.996 | 0.399 |
| VAL | 21 | A | 0.707 | 0.704 | 1 | 0.411 | | VAL | 86 | A | 0.7 | 0.705 | 0.999 | 0.406 |
| **ARG** | **22** | **A** | **0.799** | **0.81** | **0.889** | **0.72** | | THR | 87 | A | 0.778 | 0.72 | 0.968 | 0.53 |
| ASN | 23 | A | 0.776 | 0.601 | 0.95 | 0.427 | | AALA | 88 | A | 0.748 | 0.699 | 0.961 | 0.486 |
| LEU | 24 | A | 0.768 | 0.773 | 0.998 | 0.543 | | **AGLU** | **89** | **A** | **0.822** | **0.803** | **0.895** | **0.73** |
| LEU | 25 | A | 0.729 | 0.732 | 0.999 | 0.462 | | AALA | 90 | A | 0.764 | 0.677 | 0.983 | 0.458 |
| LYS | 26 | A | 0.762 | 0.75 | 0.909 | 0.603 | | LYS | 91 | A | 0.767 | 0.692 | 0.834 | 0.625 |
| GLU | 27 | A | 0.797 | 0.747 | 0.92 | 0.624 | | LYS | 92 | A | 0.734 | 0.786 | 0.97 | 0.55 |
| LEU | 28 | A | 0.73 | 0.725 | 0.999 | 0.456 | | GLU | 93 | A | 0.756 | 0.81 | 0.99 | 0.576 |
| GLY | 29 | A | 0.646 | 0.648 | 1 | 0.294 | | ASN | 94 | A | 0.72 | 0.748 | 0.946 | 0.522 |
| **PHE** | **30** | **A** | **0.831** | **0.847** | **0.95** | **0.728** | | ILE | 95 | A | 0.77 | 0.777 | 0.996 | 0.551 |
| ASN | 31 | A | 0.756 | 0.727 | 0.999 | 0.484 | | ILE | 96 | A | 0.753 | 0.8 | 0.899 | 0.654 |
| ASN | 32 | A | 0.723 | 0.744 | 0.999 | 0.468 | | ALA | 97 | A | 0.653 | 0.599 | 0.999 | 0.253 |
| VAL | 33 | A | 0.699 | 0.696 | 0.999 | 0.396 | | ALA | 98 | A | 0.6 | 0.544 | 1 | 0.144 |
| GLU | 34 | A | 0.763 | 0.812 | 0.914 | 0.661 | | ALA | 99 | A | 0.633 | 0.597 | 0.999 | 0.231 |
| GLU | 35 | A | 0.754 | 0.741 | 0.999 | 0.496 | | GLN | 100 | A | 0.693 | 0.775 | 0.923 | 0.545 |
| ALA | 36 | A | 0.493 | 0.513 | 1 | 0.006 | | ALA | 101 | A | 0.432 | 0.428 | 0.999 | -0.139 |
| GLU | 37 | A | 0.693 | 0.651 | 0.983 | 0.361 | | GLY | 102 | A | 0.644 | 0.642 | 1 | 0.286 |
| ASP | 38 | A | 0.753 | 0.733 | 0.919 | 0.567 | | ALA | 103 | A | 0.451 | 0.467 | 1 | -0.082 |
| GLY | 39 | A | 0.623 | 0.622 | 1 | 0.245 | | SER | 104 | A | 0.612 | 0.649 | 1 | 0.261 |
| VAL | 40 | A | 0.658 | 0.69 | 1 | 0.348 | | GLY | 105 | A | 0.63 | 0.627 | 1 | 0.257 |
| ASP | 41 | A | 0.758 | 0.642 | 0.915 | 0.485 | | **TYR** | **106** | **A** | **0.853** | **0.837** | **0.926** | **0.764** |
| ALA | 42 | A | 0.574 | 0.578 | 1 | 0.152 | | VAL | 107 | A | 0.737 | 0.705 | 0.997 | 0.445 |
| LEU | 43 | A | 0.744 | 0.792 | 1 | 0.536 | | VAL | 108 | A | 0.709 | 0.636 | 0.997 | 0.348 |
| ASN | 44 | A | 0.78 | 0.768 | 1 | 0.548 | | LYS | 109 | A | 0.774 | 0.755 | 0.974 | 0.555 |
| LYS | 45 | A | 0.804 | 0.736 | 0.975 | 0.565 | | PRO | 110 | A | 0.699 | 0.699 | 0.998 | 0.4 |
| LEU | 46 | A | 0.725 | 0.748 | 1 | 0.473 | | PHE | 111 | A | 0.821 | 0.826 | 0.996 | 0.651 |
| GLN | 47 | A | 0.674 | 0.696 | 0.983 | 0.387 | | THR | 112 | A | 0.75 | 0.718 | 0.998 | 0.47 |
| ALA | 48 | A | 0.523 | 0.56 | 0.994 | 0.089 | | ALA | 113 | A | 0.568 | 0.653 | 0.986 | 0.235 |
| GLY | 49 | A | 0.651 | 0.649 | 0.999 | 0.301 | | ALA | 114 | A | 0.715 | 0.672 | 0.992 | 0.395 |
| GLY | 50 | A | 0.647 | 0.646 | 1 | 0.293 | | THR | 115 | A | 0.649 | 0.706 | 0.995 | 0.36 |
| TYR | 51 | A | 0.838 | 0.847 | 0.999 | 0.686 | | LEU | 116 | A | 0.727 | 0.715 | 0.989 | 0.453 |
| GLY | 52 | A | 0.636 | 0.635 | 1 | 0.271 | | GLU | 117 | A | 0.748 | 0.706 | 0.999 | 0.455 |
| PHE | 53 | A | 0.838 | 0.839 | 0.999 | 0.678 | | GLU | 118 | A | 0.715 | 0.724 | 0.997 | 0.442 |
| VAL | 54 | A | 0.711 | 0.707 | 1 | 0.418 | | LYS | 119 | A | 0.788 | 0.76 | 0.905 | 0.643 |
| ILE | 55 | A | 0.731 | 0.741 | 0.998 | 0.474 | | LEU | 120 | A | 0.741 | 0.75 | 0.992 | 0.499 |
| SER | 56 | A | 0.657 | 0.545 | 0.989 | 0.213 | | ASN | 121 | A | 0.715 | 0.787 | 0.991 | 0.511 |
| ASP | 57 | A | 0.786 | 0.833 | 1 | 0.619 | | LYS | 122 | A | 0.682 | 0.705 | 0.908 | 0.479 |
| **TRP** | **58** | **A** | **0.864** | **0.863** | **0.996** | **0.731** | | ILE | 123 | A | 0.756 | 0.745 | 0.997 | 0.504 |
| ASN | 59 | A | 0.722 | 0.681 | 0.998 | 0.405 | | **PHE** | **124** | **A** | **0.827** | **0.838** | **0.944** | **0.721** |
| MET | 60 | A | 0.751 | 0.73 | 1 | 0.481 | | GLU | 125 | A | 0.652 | 0.679 | 0.939 | 0.392 |
| PRO | 61 | A | 0.682 | 0.677 | 1 | 0.359 | | LYS | 126 | A | 0.731 | 0.717 | 0.844 | 0.604 |
| ASN | 62 | A | 0.676 | 0.695 | 0.996 | 0.375 | | LEU | 127 | A | 0.7 | 0.627 | 0.976 | 0.351 |
| MET | 63 | A | 0.764 | 0.763 | 1 | 0.527 | | GLY | 128 | A | 0.659 | 0.67 | 0.918 | 0.411 |
| ASP | 64 | A | 0.603 | 0.765 | 0.997 | 0.371 | | MET | 129 | A | 0.784 | 0.783 | 0.936 | 0.631 |
| GLY | 65 | A | 0.616 | 0.621 | 1 | 0.237 | | | | | | | | |
| LEU | 66 | A | 0.736 | 0.759 | 0.871 | 0.624 | | | | | | | | |

**Table S1(h): 2B1J (P), 1FQW (Q)**

| Residue | | | S (P) | S (Q) | J (P,Q) | PRI | Residue | | | S (P) | S (Q) | J (P,Q) | PRI |
|---|---|---|---|---|---|---|---|---|---|---|---|---|---|
| ALA | 2 | A | 0.669 | 0.659 | 0.839 | 0.489 | GLU | 67 | A | 0.76 | 0.701 | 0.895 | 0.566 |
| ASP | 3 | A | 0.791 | 0.801 | 0.944 | 0.648 | LEU | 68 | A | 0.742 | 0.736 | 0.999 | 0.479 |
| LYS | 4 | A | 0.803 | 0.804 | 0.992 | 0.615 | LEU | 69 | A | 0.743 | 0.741 | 1 | 0.484 |
| GLU | 5 | A | 0.662 | 0.768 | 0.96 | 0.47 | LYS | 70 | A | 0.8 | 0.811 | 0.999 | 0.612 |
| LEU | 6 | A | 0.744 | 0.738 | 0.991 | 0.491 | THR | 71 | A | 0.685 | 0.668 | 1 | 0.353 |
| LYS | 7 | A | 0.799 | 0.799 | 0.851 | 0.747 | ILE | 72 | A | 0.77 | 0.747 | 0.999 | 0.518 |
| PHE | 8 | A | 0.816 | 0.841 | 1 | 0.657 | ARG | 73 | A | 0.845 | 0.853 | 0.999 | 0.699 |
| LEU | 9 | A | 0.742 | 0.755 | 1 | 0.497 | ALA | 74 | A | 0.511 | 0.505 | 1 | 0.016 |
| VAL | 10 | A | 0.698 | 0.687 | 0.999 | 0.386 | ASP | 75 | A | 0.755 | 0.763 | 0.929 | 0.589 |
| VAL | 11 | A | 0.699 | 0.699 | 0.999 | 0.399 | GLY | 76 | A | 0.661 | 0.663 | 0.976 | 0.348 |
| ASP | 12 | A | 0.764 | 0.778 | 0.996 | 0.546 | ALA | 77 | A | 0.593 | 0.522 | 0.997 | 0.118 |
| ASP | 13 | A | 0.751 | 0.776 | 0.999 | 0.528 | MET | 78 | A | 0.737 | 0.744 | 0.959 | 0.522 |
| **APHE** | **14** | **A** | **0.798** | **0.799** | **0.679** | **0.918** | SER | 79 | A | 0.675 | 0.716 | 0.992 | 0.399 |
| SER | 15 | A | 0.776 | 0.778 | 0.856 | 0.698 | ALA | 80 | A | 0.517 | 0.537 | 0.998 | 0.056 |
| THR | 16 | A | 0.736 | 0.662 | 0.972 | 0.426 | LEU | 81 | A | 0.692 | 0.747 | 0.998 | 0.441 |
| MET | 17 | A | 0.709 | 0.771 | 0.893 | 0.587 | PRO | 82 | A | 0.655 | 0.671 | 1 | 0.326 |
| ARG | 18 | A | 0.826 | 0.821 | 0.996 | 0.651 | VAL | 83 | A | 0.704 | 0.694 | 0.999 | 0.399 |
| ARG | 19 | A | 0.676 | 0.745 | 0.958 | 0.463 | LEU | 84 | A | 0.72 | 0.757 | 0.999 | 0.478 |
| ILE | 20 | A | 0.78 | 0.767 | 0.994 | 0.553 | MET | 85 | A | 0.689 | 0.712 | 0.998 | 0.403 |
| VAL | 21 | A | 0.707 | 0.699 | 0.999 | 0.407 | VAL | 86 | A | 0.7 | 0.697 | 0.998 | 0.399 |
| ARG | 22 | A | 0.799 | 0.82 | 0.998 | 0.621 | THR | 87 | A | 0.778 | 0.701 | 0.941 | 0.538 |
| ASN | 23 | A | 0.776 | 0.678 | 0.868 | 0.586 | AALA | 88 | A | 0.748 | 0.693 | 0.952 | 0.489 |
| LEU | 24 | A | 0.768 | 0.784 | 0.994 | 0.558 | AGLU | 89 | A | 0.822 | 0.769 | 0.864 | 0.727 |
| LEU | 25 | A | 0.729 | 0.741 | 0.998 | 0.472 | AALA | 90 | A | 0.764 | 0.655 | 0.97 | 0.449 |
| LYS | 26 | A | 0.762 | 0.7 | 0.836 | 0.626 | LYS | 91 | A | 0.767 | 0.664 | 0.771 | 0.66 |
| GLU | 27 | A | 0.797 | 0.737 | 0.864 | 0.67 | **LYS** | **92** | **A** | **0.734** | **0.773** | **0.639** | **0.868** |
| LEU | 28 | A | 0.73 | 0.707 | 0.998 | 0.439 | GLU | 93 | A | 0.756 | 0.765 | 0.799 | 0.722 |
| GLY | 29 | A | 0.646 | 0.648 | 1 | 0.294 | ASN | 94 | A | 0.72 | 0.745 | 0.917 | 0.548 |
| PHE | 30 | A | 0.831 | 0.849 | 0.999 | 0.681 | ILE | 95 | A | 0.77 | 0.713 | 0.846 | 0.637 |
| ASN | 31 | A | 0.756 | 0.713 | 0.998 | 0.471 | ILE | 96 | A | 0.753 | 0.832 | 0.884 | 0.701 |
| ASN | 32 | A | 0.723 | 0.729 | 0.998 | 0.454 | ALA | 97 | A | 0.653 | 0.577 | 0.996 | 0.234 |
| VAL | 33 | A | 0.699 | 0.714 | 0.998 | 0.415 | ALA | 98 | A | 0.6 | 0.525 | 0.998 | 0.127 |
| **GLU** | **34** | **A** | **0.763** | **0.838** | **0.787** | **0.814** | ALA | 99 | A | 0.633 | 0.643 | 0.999 | 0.277 |
| GLU | 35 | A | 0.754 | 0.75 | 0.999 | 0.505 | GLN | 100 | A | 0.693 | 0.676 | 0.892 | 0.477 |
| ALA | 36 | A | 0.493 | 0.496 | 1 | -0.011 | ALA | 101 | A | 0.432 | 0.514 | 0.999 | -0.053 |
| GLU | 37 | A | 0.693 | 0.695 | 0.93 | 0.458 | GLY | 102 | A | 0.644 | 0.64 | 0.999 | 0.285 |
| ASP | 38 | A | 0.753 | 0.762 | 1 | 0.515 | ALA | 103 | A | 0.451 | 0.503 | 1 | -0.046 |
| GLY | 39 | A | 0.623 | 0.621 | 1 | 0.244 | SER | 104 | A | 0.612 | 0.63 | 1 | 0.242 |
| VAL | 40 | A | 0.658 | 0.637 | 1 | 0.295 | GLY | 105 | A | 0.63 | 0.627 | 1 | 0.257 |
| ASP | 41 | A | 0.758 | 0.737 | 1 | 0.495 | **TYR** | **106** | **A** | **0.853** | **0.833** | **0.883** | **0.803** |
| ALA | 42 | A | 0.574 | 0.579 | 0.999 | 0.154 | VAL | 107 | A | 0.737 | 0.709 | 0.997 | 0.449 |
| LEU | 43 | A | 0.744 | 0.766 | 0.999 | 0.511 | VAL | 108 | A | 0.709 | 0.664 | 0.886 | 0.487 |
| ASN | 44 | A | 0.78 | 0.755 | 0.926 | 0.609 | LYS | 109 | A | 0.774 | 0.77 | 0.94 | 0.604 |
| LYS | 45 | A | 0.804 | 0.806 | 0.969 | 0.641 | PRO | 110 | A | 0.699 | 0.706 | 0.998 | 0.407 |
| LEU | 46 | A | 0.725 | 0.738 | 0.999 | 0.464 | PHE | 111 | A | 0.821 | 0.814 | 0.994 | 0.641 |
| GLN | 47 | A | 0.674 | 0.681 | 0.86 | 0.495 | THR | 112 | A | 0.75 | 0.699 | 0.995 | 0.454 |
| ALA | 48 | A | 0.523 | 0.537 | 0.986 | 0.074 | ALA | 113 | A | 0.568 | 0.663 | 0.981 | 0.25 |
| GLY | 49 | A | 0.651 | 0.647 | 0.997 | 0.301 | ALA | 114 | A | 0.715 | 0.65 | 0.992 | 0.373 |
| GLY | 50 | A | 0.647 | 0.644 | 0.996 | 0.295 | THR | 115 | A | 0.649 | 0.689 | 0.986 | 0.352 |
| TYR | 51 | A | 0.838 | 0.852 | 0.998 | 0.692 | LEU | 116 | A | 0.727 | 0.717 | 0.989 | 0.455 |
| GLY | 52 | A | 0.636 | 0.641 | 1 | 0.277 | GLU | 117 | A | 0.748 | 0.774 | 0.994 | 0.528 |
| PHE | 53 | A | 0.838 | 0.843 | 0.999 | 0.682 | GLU | 118 | A | 0.715 | 0.732 | 0.909 | 0.538 |
| VAL | 54 | A | 0.711 | 0.713 | 0.999 | 0.425 | LYS | 119 | A | 0.788 | 0.728 | 0.905 | 0.611 |
| ILE | 55 | A | 0.731 | 0.741 | 0.995 | 0.477 | LEU | 120 | A | 0.741 | 0.749 | 0.992 | 0.498 |
| SER | 56 | A | 0.657 | 0.682 | 0.999 | 0.34 | ASN | 121 | A | 0.714 | 0.808 | 0.957 | 0.565 |
| ASP | 57 | A | 0.786 | 0.838 | 1 | 0.624 | LYS | 122 | A | 0.682 | 0.66 | 0.901 | 0.441 |
| TRP | 58 | A | 0.864 | 0.869 | 0.988 | 0.745 | ILE | 123 | A | 0.757 | 0.757 | 0.999 | 0.515 |
| ASN | 59 | A | 0.722 | 0.686 | 0.831 | 0.577 | PHE | 124 | A | 0.829 | 0.829 | 0.996 | 0.662 |
| MET | 60 | A | 0.751 | 0.745 | 0.998 | 0.498 | GLU | 125 | A | 0.652 | 0.692 | 0.838 | 0.506 |
| PRO | 61 | A | 0.682 | 0.687 | 0.999 | 0.37 | LYS | 126 | A | 0.731 | 0.521 | 0.88 | 0.372 |
| ASN | 62 | A | 0.676 | 0.726 | 0.998 | 0.404 | LEU | 127 | A | 0.7 | 0.729 | 0.847 | 0.582 |
| MET | 63 | A | 0.764 | 0.768 | 0.999 | 0.533 | GLY | 128 | A | 0.66 | 0.659 | 0.977 | 0.342 |
| ASP | 64 | A | 0.603 | 0.672 | 1 | 0.275 | **MET** | **129** | **A** | **0.806** | **0.879** | **0.929** | **0.756** |
| GLY | 65 | A | 0.616 | 0.619 | 1 | 0.235 | | | | | | | |
| LEU | 66 | A | 0.736 | 0.773 | 0.809 | 0.7 | | | | | | | |

**Table S1(i): 2B1J (P), 3CHY (Q)**

| Residue | | | S (P) | S (Q) | J (P,Q) | PRI |
|---|---|---|---|---|---|---|
| ALA | 2 | A | 0.669 | 0.526 | 0.876 | 0.319 |
| ASP | 3 | A | 0.791 | 0.772 | 0.846 | 0.717 |
| LYS | 4 | A | 0.803 | 0.78 | 0.929 | 0.654 |
| GLU | 5 | A | 0.66 | 0.808 | 0.811 | 0.657 |
| LEU | 6 | A | 0.744 | 0.723 | 0.995 | 0.472 |
| LYS | 7 | A | 0.816 | 0.806 | 0.858 | 0.764 |
| PHE | 8 | A | 0.816 | 0.822 | 0.998 | 0.64 |
| LEU | 9 | A | 0.742 | 0.75 | 0.997 | 0.495 |
| VAL | 10 | A | 0.698 | 0.69 | 0.999 | 0.389 |
| VAL | 11 | A | 0.699 | 0.7 | 0.999 | 0.4 |
| ASP | 12 | A | 0.764 | 0.799 | 0.984 | 0.579 |
| ASP | 13 | A | 0.751 | 0.747 | 0.986 | 0.512 |
| APHE | 14 | A | 0.798 | 0.731 | 0.769 | 0.76 |
| SER | 15 | A | 0.775 | 0.707 | 0.898 | 0.584 |
| THR | 16 | A | 0.736 | 0.731 | 0.998 | 0.469 |
| MET | 17 | A | 0.709 | 0.711 | 1 | 0.42 |
| ARG | 18 | A | 0.826 | 0.845 | 0.994 | 0.677 |
| ARG | 19 | A | 0.725 | 0.76 | 0.723 | 0.762 |
| ILE | 20 | A | 0.779 | 0.782 | 1 | 0.561 |
| VAL | 21 | A | 0.707 | 0.685 | 1 | 0.392 |
| ARG | 22 | A | 0.801 | 0.848 | 0.994 | 0.655 |
| ASN | 23 | A | 0.749 | 0.694 | 0.886 | 0.557 |
| LEU | 24 | A | 0.768 | 0.773 | 0.996 | 0.545 |
| LEU | 25 | A | 0.729 | 0.733 | 0.999 | 0.463 |
| **LYS** | **26** | **A** | **0.788** | **0.762** | **0.656** | **0.894** |
| GLU | 27 | A | 0.797 | 0.821 | 0.939 | 0.679 |
| LEU | 28 | A | 0.73 | 0.72 | 0.995 | 0.455 |
| GLY | 29 | A | 0.646 | 0.638 | 0.999 | 0.285 |
| PHE | 30 | A | 0.831 | 0.836 | 0.998 | 0.669 |
| ASN | 31 | A | 0.756 | 0.767 | 0.998 | 0.525 |
| ASN | 32 | A | 0.69 | 0.695 | 0.824 | 0.561 |
| VAL | 33 | A | 0.699 | 0.693 | 0.996 | 0.396 |
| GLU | 34 | A | 0.763 | 0.776 | 0.871 | 0.668 |
| GLU | 35 | A | 0.754 | 0.758 | 0.999 | 0.513 |
| ALA | 36 | A | 0.494 | 0.527 | 0.993 | 0.028 |
| GLU | 37 | A | 0.734 | 0.76 | 0.961 | 0.533 |
| ASP | 38 | A | 0.753 | 0.767 | 0.996 | 0.524 |
| GLY | 39 | A | 0.623 | 0.625 | 0.999 | 0.249 |
| VAL | 40 | A | 0.658 | 0.72 | 1 | 0.378 |
| ASP | 41 | A | 0.758 | 0.754 | 1 | 0.512 |
| ALA | 42 | A | 0.573 | 0.553 | 1 | 0.126 |
| LEU | 43 | A | 0.743 | 0.767 | 0.999 | 0.511 |
| ASN | 44 | A | 0.78 | 0.715 | 0.985 | 0.51 |
| LYS | 45 | A | 0.804 | 0.8 | 0.981 | 0.623 |
| LEU | 46 | A | 0.724 | 0.725 | 0.999 | 0.45 |
| GLN | 47 | A | 0.674 | 0.696 | 0.85 | 0.52 |
| ALA | 48 | A | 0.523 | 0.589 | 0.956 | 0.156 |
| GLY | 49 | A | 0.651 | 0.652 | 1 | 0.303 |
| GLY | 50 | A | 0.647 | 0.637 | 0.999 | 0.285 |
| TYR | 51 | A | 0.838 | 0.852 | 0.999 | 0.691 |
| GLY | 52 | A | 0.636 | 0.636 | 0.998 | 0.274 |
| PHE | 53 | A | 0.838 | 0.837 | 0.999 | 0.676 |
| VAL | 54 | A | 0.711 | 0.703 | 1 | 0.414 |
| ILE | 55 | A | 0.731 | 0.748 | 0.999 | 0.48 |
| SER | 56 | A | 0.742 | 0.744 | 0.84 | 0.646 |
| ASP | 57 | A | 0.789 | 0.78 | 0.913 | 0.656 |
| TRP | 58 | A | 0.865 | 0.845 | 0.981 | 0.729 |
| ASN | 59 | A | 0.722 | 0.631 | 0.876 | 0.477 |
| MET | 60 | A | 0.75 | 0.755 | 0.999 | 0.506 |
| PRO | 61 | A | 0.682 | 0.666 | 1 | 0.348 |
| ASN | 62 | A | 0.676 | 0.744 | 0.988 | 0.432 |
| MET | 63 | A | 0.764 | 0.77 | 0.995 | 0.539 |
| ASP | 64 | A | 0.603 | 0.637 | 0.998 | 0.242 |
| GLY | 65 | A | 0.613 | 0.612 | 0.998 | 0.227 |
| LEU | 66 | A | 0.736 | 0.736 | 1 | 0.472 |

| Residue | | | S (P) | S (Q) | J (P,Q) | PRI |
|---|---|---|---|---|---|---|
| GLU | 67 | A | 0.76 | 0.814 | 0.823 | 0.751 |
| LEU | 68 | A | 0.742 | 0.74 | 1 | 0.482 |
| LEU | 69 | A | 0.742 | 0.736 | 0.999 | 0.479 |
| LYS | 70 | A | 0.8 | 0.782 | 0.997 | 0.585 |
| THR | 71 | A | 0.685 | 0.708 | 0.998 | 0.395 |
| ILE | 72 | A | 0.77 | 0.746 | 1 | 0.516 |
| ARG | 73 | A | 0.845 | 0.849 | 0.999 | 0.695 |
| ALA | 74 | A | 0.511 | 0.493 | 1 | 0.004 |
| ASP | 75 | A | 0.755 | 0.778 | 1 | 0.533 |
| GLY | 76 | A | 0.661 | 0.655 | 0.991 | 0.325 |
| ALA | 77 | A | 0.592 | 0.556 | 0.984 | 0.164 |
| MET | 78 | A | 0.737 | 0.739 | 0.981 | 0.495 |
| SER | 79 | A | 0.675 | 0.693 | 0.968 | 0.4 |
| ALA | 80 | A | 0.517 | 0.53 | 1 | 0.047 |
| LEU | 81 | A | 0.692 | 0.693 | 0.998 | 0.387 |
| PRO | 82 | A | 0.655 | 0.66 | 0.999 | 0.316 |
| VAL | 83 | A | 0.702 | 0.709 | 0.999 | 0.412 |
| LEU | 84 | A | 0.722 | 0.729 | 0.998 | 0.453 |
| MET | 85 | A | 0.805 | 0.824 | 0.91 | 0.719 |
| VAL | 86 | A | 0.701 | 0.716 | 0.954 | 0.463 |
| THR | 87 | A | 0.777 | 0.717 | 0.948 | 0.546 |
| AALA | 88 | A | 0.748 | 0.69 | 0.945 | 0.493 |
| **AGLU** | **89** | **A** | **0.822** | **0.781** | **0.632** | **0.971** |
| AALA | 90 | A | 0.765 | 0.651 | 0.982 | 0.434 |
| LYS | 91 | A | 0.767 | 0.766 | 0.949 | 0.584 |
| LYS | 92 | A | 0.734 | 0.793 | 0.95 | 0.577 |
| GLU | 93 | A | 0.756 | 0.789 | 0.841 | 0.704 |
| ASN | 94 | A | 0.72 | 0.769 | 0.979 | 0.51 |
| **ILE** | **95** | **A** | **0.769** | **0.761** | **0.704** | **0.826** |
| ILE | 96 | A | 0.753 | 0.811 | 0.924 | 0.64 |
| ALA | 97 | A | 0.653 | 0.615 | 0.997 | 0.271 |
| ALA | 98 | A | 0.601 | 0.565 | 1 | 0.166 |
| ALA | 99 | A | 0.634 | 0.641 | 0.998 | 0.277 |
| GLN | 100 | A | 0.693 | 0.81 | 0.903 | 0.6 |
| ALA | 101 | A | 0.431 | 0.463 | 0.993 | -0.099 |
| GLY | 102 | A | 0.644 | 0.638 | 0.997 | 0.285 |
| ALA | 103 | A | 0.446 | 0.477 | 0.999 | -0.076 |
| SER | 104 | A | 0.613 | 0.626 | 0.999 | 0.24 |
| GLY | 105 | A | 0.629 | 0.618 | 0.999 | 0.248 |
| **TYR** | **106** | **A** | **0.83** | **0.833** | **0.549** | **1.114** |
| VAL | 107 | A | 0.736 | 0.673 | 0.741 | 0.668 |
| VAL | 108 | A | 0.711 | 0.669 | 0.999 | 0.381 |
| LYS | 109 | A | 0.774 | 0.759 | 0.827 | 0.706 |
| PRO | 110 | A | 0.699 | 0.7 | 0.998 | 0.401 |
| PHE | 111 | A | 0.821 | 0.813 | 0.994 | 0.64 |
| THR | 112 | A | 0.75 | 0.74 | 0.994 | 0.496 |
| ALA | 113 | A | 0.568 | 0.612 | 0.989 | 0.191 |
| ALA | 114 | A | 0.715 | 0.654 | 0.994 | 0.375 |
| THR | 115 | A | 0.649 | 0.714 | 0.997 | 0.366 |
| LEU | 116 | A | 0.727 | 0.717 | 0.999 | 0.445 |
| GLU | 117 | A | 0.748 | 0.768 | 0.997 | 0.519 |
| GLU | 118 | A | 0.715 | 0.709 | 0.993 | 0.431 |
| LYS | 119 | A | 0.788 | 0.732 | 0.844 | 0.676 |
| LEU | 120 | A | 0.741 | 0.706 | 0.992 | 0.455 |
| ASN | 121 | A | 0.714 | 0.764 | 0.974 | 0.504 |
| LYS | 122 | A | 0.682 | 0.674 | 0.793 | 0.563 |
| ILE | 123 | A | 0.756 | 0.745 | 0.997 | 0.504 |
| PHE | 124 | A | 0.828 | 0.846 | 0.989 | 0.685 |
| GLU | 125 | A | 0.652 | 0.659 | 0.831 | 0.48 |
| **LYS** | **126** | **A** | **0.731** | **0.625** | **0.544** | **0.812** |
| LEU | 127 | A | 0.601 | 0.684 | 0.92 | 0.365 |
| GLY | 128 | A | 0.66 | 0.639 | 0.999 | 0.3 |
| MET | 129 | A | 0.797 | 0.813 | 0.96 | 0.65 |

| Table S1(j): 3CHY (P), 1FQW (Q) | | | | | | |
|---|---|---|---|---|---|---|
| Residue | | | S (P) | S (Q) | J (P,Q) | PRI |
| ALA | 2 | A | 0.526 | 0.66 | 0.954 | 0.232 |
| ASP | 3 | A | 0.772 | 0.801 | 0.985 | 0.588 |
| LYS | 4 | A | 0.78 | 0.804 | 0.986 | 0.598 |
| GLU | 5 | A | 0.808 | 0.755 | 0.933 | 0.63 |
| LEU | 6 | A | 0.723 | 0.738 | 0.997 | 0.464 |
| LYS | 7 | A | 0.807 | 0.799 | 0.999 | 0.607 |
| PHE | 8 | A | 0.822 | 0.841 | 1 | 0.663 |
| LEU | 9 | A | 0.75 | 0.755 | 1 | 0.505 |
| VAL | 10 | A | 0.69 | 0.687 | 1 | 0.377 |
| VAL | 11 | A | 0.7 | 0.699 | 0.998 | 0.401 |
| ASP | 12 | A | 0.802 | 0.781 | 0.979 | 0.604 |
| ASP | 13 | A | 0.738 | 0.769 | 0.995 | 0.512 |
| **PHE** | **14** | **A** | **0.785** | **0.842** | **0.872** | **0.755** |
| SER | 15 | A | 0.704 | 0.775 | 0.997 | 0.482 |
| THR | 16 | A | 0.732 | 0.665 | 0.974 | 0.423 |
| MET | 17 | A | 0.707 | 0.769 | 0.947 | 0.529 |
| ARG | 18 | A | 0.845 | 0.82 | 0.998 | 0.667 |
| ARG | 19 | A | 0.761 | 0.711 | 0.822 | 0.65 |
| ILE | 20 | A | 0.784 | 0.767 | 0.995 | 0.556 |
| VAL | 21 | A | 0.685 | 0.699 | 0.998 | 0.386 |
| ARG | 22 | A | 0.848 | 0.821 | 0.996 | 0.673 |
| ASN | 23 | A | 0.693 | 0.714 | 0.975 | 0.432 |
| LEU | 24 | A | 0.771 | 0.782 | 0.993 | 0.56 |
| LEU | 25 | A | 0.732 | 0.741 | 0.999 | 0.474 |
| LYS | 26 | A | 0.761 | 0.673 | 0.745 | 0.689 |
| GLU | 27 | A | 0.785 | 0.717 | 0.788 | 0.714 |
| LEU | 28 | A | 0.72 | 0.705 | 0.998 | 0.427 |
| GLY | 29 | A | 0.638 | 0.648 | 1 | 0.286 |
| PHE | 30 | A | 0.836 | 0.849 | 0.999 | 0.686 |
| ASN | 31 | A | 0.768 | 0.713 | 0.993 | 0.488 |
| ASN | 32 | A | 0.695 | 0.69 | 0.956 | 0.429 |
| VAL | 33 | A | 0.693 | 0.714 | 0.999 | 0.408 |
| GLU | 34 | A | 0.776 | 0.839 | 0.921 | 0.694 |
| GLU | 35 | A | 0.758 | 0.75 | 1 | 0.508 |
| ALA | 36 | A | 0.526 | 0.496 | 0.999 | 0.023 |
| GLU | 37 | A | 0.76 | 0.755 | 0.917 | 0.598 |
| ASP | 38 | A | 0.767 | 0.762 | 0.998 | 0.531 |
| GLY | 39 | A | 0.625 | 0.621 | 1 | 0.246 |
| VAL | 40 | A | 0.72 | 0.637 | 0.999 | 0.358 |
| ASP | 41 | A | 0.754 | 0.738 | 1 | 0.492 |
| ALA | 42 | A | 0.553 | 0.578 | 1 | 0.131 |
| LEU | 43 | A | 0.767 | 0.766 | 0.998 | 0.535 |
| ASN | 44 | A | 0.715 | 0.755 | 0.964 | 0.506 |
| LYS | 45 | A | 0.8 | 0.806 | 0.991 | 0.615 |
| LEU | 46 | A | 0.725 | 0.738 | 1 | 0.463 |
| GLN | 47 | A | 0.696 | 0.681 | 0.791 | 0.586 |
| ALA | 48 | A | 0.589 | 0.537 | 0.998 | 0.128 |
| GLY | 49 | A | 0.651 | 0.647 | 1 | 0.298 |
| GLY | 50 | A | 0.637 | 0.645 | 1 | 0.282 |
| TYR | 51 | A | 0.852 | 0.852 | 0.999 | 0.705 |
| GLY | 52 | A | 0.636 | 0.641 | 0.999 | 0.278 |
| PHE | 53 | A | 0.837 | 0.844 | 0.999 | 0.682 |
| VAL | 54 | A | 0.703 | 0.713 | 1 | 0.416 |
| ILE | 55 | A | 0.748 | 0.741 | 1 | 0.489 |
| SER | 56 | A | 0.744 | 0.76 | 0.911 | 0.593 |
| ASP | 57 | A | 0.78 | 0.843 | 0.957 | 0.666 |
| TRP | 58 | A | 0.846 | 0.869 | 0.974 | 0.741 |
| ASN | 59 | A | 0.636 | 0.688 | 0.998 | 0.326 |
| MET | 60 | A | 0.755 | 0.744 | 0.999 | 0.5 |
| PRO | 61 | A | 0.667 | 0.687 | 0.999 | 0.355 |
| ASN | 62 | A | 0.744 | 0.726 | 0.995 | 0.475 |
| MET | 63 | A | 0.77 | 0.768 | 0.999 | 0.539 |
| ASP | 64 | A | 0.638 | 0.673 | 0.998 | 0.313 |
| GLY | 65 | A | 0.612 | 0.617 | 1 | 0.229 |
| LEU | 66 | A | 0.736 | 0.774 | 0.883 | 0.627 |
| GLU | 67 | A | 0.815 | 0.701 | 0.902 | 0.614 |
| LEU | 68 | A | 0.74 | 0.736 | 0.999 | 0.477 |
| LEU | 69 | A | 0.736 | 0.741 | 0.999 | 0.478 |
| LYS | 70 | A | 0.782 | 0.811 | 0.995 | 0.598 |
| THR | 71 | A | 0.708 | 0.668 | 0.999 | 0.377 |
| ILE | 72 | A | 0.746 | 0.747 | 0.999 | 0.494 |
| ARG | 73 | A | 0.849 | 0.853 | 0.999 | 0.703 |
| ALA | 74 | A | 0.493 | 0.505 | 0.999 | -0.001 |
| ASP | 75 | A | 0.778 | 0.763 | 0.95 | 0.591 |
| GLY | 76 | A | 0.655 | 0.663 | 1 | 0.318 |
| ALA | 77 | A | 0.556 | 0.522 | 0.998 | 0.08 |
| MET | 78 | A | 0.74 | 0.744 | 0.987 | 0.497 |
| SER | 79 | A | 0.693 | 0.716 | 0.999 | 0.41 |
| ALA | 80 | A | 0.53 | 0.537 | 0.998 | 0.069 |
| LEU | 81 | A | 0.693 | 0.747 | 0.999 | 0.441 |
| PRO | 82 | A | 0.66 | 0.656 | 1 | 0.316 |
| VAL | 83 | A | 0.709 | 0.692 | 0.999 | 0.402 |
| LEU | 84 | A | 0.729 | 0.758 | 0.998 | 0.489 |
| MET | 85 | A | 0.824 | 0.826 | 0.958 | 0.692 |
| VAL | 86 | A | 0.715 | 0.697 | 0.969 | 0.443 |
| THR | 87 | A | 0.718 | 0.699 | 0.91 | 0.507 |
| ALA | 88 | A | 0.531 | 0.525 | 0.942 | 0.114 |
| GLU | 89 | A | 0.737 | 0.648 | 0.863 | 0.522 |
| ALA | 90 | A | 0.481 | 0.508 | 0.957 | 0.032 |
| LYS | 91 | A | 0.833 | 0.668 | 0.869 | 0.632 |
| **LYS** | **92** | **A** | **0.793** | **0.779** | **0.648** | **0.924** |
| **GLU** | **93** | **A** | **0.79** | **0.766** | **0.793** | **0.763** |
| ASN | 94 | A | 0.773 | 0.726 | 0.947 | 0.552 |
| ILE | 95 | A | 0.769 | 0.723 | 0.999 | 0.493 |
| ILE | 96 | A | 0.811 | 0.832 | 0.999 | 0.644 |
| ALA | 97 | A | 0.615 | 0.578 | 1 | 0.193 |
| ALA | 98 | A | 0.566 | 0.525 | 0.999 | 0.092 |
| ALA | 99 | A | 0.642 | 0.644 | 1 | 0.286 |
| GLN | 100 | A | 0.81 | 0.677 | 0.949 | 0.538 |
| ALA | 101 | A | 0.463 | 0.514 | 0.999 | -0.022 |
| GLY | 102 | A | 0.638 | 0.64 | 0.999 | 0.279 |
| ALA | 103 | A | 0.477 | 0.499 | 0.999 | -0.023 |
| SER | 104 | A | 0.627 | 0.631 | 1 | 0.258 |
| GLY | 105 | A | 0.618 | 0.627 | 1 | 0.245 |
| **TYR** | **106** | **A** | **0.834** | **0.798** | **0.617** | **1.015** |
| VAL | 107 | A | 0.671 | 0.707 | 0.875 | 0.503 |
| VAL | 108 | A | 0.681 | 0.661 | 0.887 | 0.455 |
| LYS | 109 | A | 0.76 | 0.77 | 0.953 | 0.577 |
| PRO | 110 | A | 0.703 | 0.708 | 0.999 | 0.412 |
| PHE | 111 | A | 0.813 | 0.814 | 0.99 | 0.637 |
| THR | 112 | A | 0.74 | 0.7 | 0.995 | 0.445 |
| ALA | 113 | A | 0.615 | 0.665 | 0.986 | 0.294 |
| ALA | 114 | A | 0.654 | 0.65 | 0.993 | 0.311 |
| THR | 115 | A | 0.714 | 0.689 | 0.991 | 0.412 |
| LEU | 116 | A | 0.717 | 0.717 | 0.994 | 0.44 |
| GLU | 117 | A | 0.768 | 0.774 | 0.998 | 0.544 |
| GLU | 118 | A | 0.709 | 0.733 | 0.908 | 0.534 |
| LYS | 119 | A | 0.733 | 0.728 | 0.787 | 0.674 |
| LEU | 120 | A | 0.706 | 0.749 | 0.977 | 0.478 |
| ASN | 121 | A | 0.764 | 0.808 | 0.992 | 0.58 |
| LYS | 122 | A | 0.674 | 0.661 | 0.849 | 0.486 |
| ILE | 123 | A | 0.746 | 0.757 | 0.995 | 0.508 |
| PHE | 124 | A | 0.846 | 0.829 | 0.999 | 0.676 |
| GLU | 125 | A | 0.659 | 0.693 | 0.931 | 0.421 |
| LYS | 126 | A | 0.625 | 0.52 | 0.649 | 0.496 |
| LEU | 127 | A | 0.684 | 0.692 | 0.997 | 0.379 |
| GLY | 128 | A | 0.639 | 0.659 | 0.999 | 0.299 |
| MET | 129 | A | 0.813 | 0.875 | 0.989 | 0.699 |

**Table S2: 4OMZ (P), 4ON0 (Q)**

| Residue | | | S (P) | S (Q) | J | PRI |
|---|---|---|---|---|---|---|
| GLN | 6 | A | 0.709 | 0.778 | 0.858 | 0.629 |
| PRO | 7 | A | 0.68 | 0.77 | 0.999 | 0.451 |
| LEU | 8 | A | 0.737 | 0.753 | 0.999 | 0.491 |
| SER | 9 | A | 0.69 | 0.724 | 0.997 | 0.417 |
| PRO | 10 | A | 0.754 | 0.741 | 1 | 0.495 |
| GLU | 11 | A | 0.655 | 0.722 | 0.769 | 0.608 |
| **LYS** | **12** | **A** | **0.713** | **0.829** | **0.746** | **0.796** |
| HIS | 13 | A | 0.793 | 0.771 | 0.958 | 0.606 |
| GLU | 14 | A | 0.768 | 0.708 | 0.737 | 0.739 |
| GLU | 15 | A | 0.705 | 0.724 | 0.773 | 0.656 |
| ALA | 16 | A | 0.548 | 0.544 | 0.999 | 0.093 |
| GLU | 17 | A | 0.761 | 0.754 | 0.996 | 0.519 |
| ILE | 18 | A | 0.798 | 0.8 | 1 | 0.598 |
| ALA | 19 | A | 0.583 | 0.582 | 1 | 0.165 |
| ALA | 20 | A | 0.564 | 0.571 | 1 | 0.135 |
| GLY | 21 | A | 0.638 | 0.64 | 1 | 0.278 |
| PHE | 22 | A | 0.84 | 0.823 | 1 | 0.663 |
| LEU | 23 | A | 0.7 | 0.713 | 1 | 0.413 |
| SER | 24 | A | 0.735 | 0.674 | 0.999 | 0.41 |
| ALA | 25 | A | 0.542 | 0.553 | 0.999 | 0.096 |
| MET | 26 | A | 0.74 | 0.748 | 1 | 0.488 |
| ALA | 27 | A | 0.495 | 0.51 | 1 | 0.005 |
| ASN | 28 | A | 0.794 | 0.798 | 0.997 | 0.595 |
| PRO | 29 | A | 0.733 | 0.758 | 1 | 0.491 |
| LYS | 30 | A | 0.75 | 0.782 | 0.999 | 0.533 |
| ARG | 31 | A | 0.847 | 0.849 | 0.997 | 0.699 |
| LEU | 32 | A | 0.679 | 0.686 | 1 | 0.365 |
| LEU | 33 | A | 0.773 | 0.732 | 0.999 | 0.506 |
| ILE | 34 | A | 0.775 | 0.771 | 1 | 0.546 |
| LEU | 35 | A | 0.724 | 0.743 | 1 | 0.467 |
| ASP | 36 | A | 0.761 | 0.793 | 0.994 | 0.56 |
| SER | 37 | A | 0.7 | 0.7 | 1 | 0.4 |
| LEU | 38 | A | 0.751 | 0.745 | 1 | 0.496 |
| VAL | 39 | A | 0.714 | 0.711 | 0.998 | 0.427 |
| LYS | 40 | A | 0.794 | 0.783 | 0.965 | 0.612 |
| GLU | 41 | A | 0.751 | 0.71 | 0.959 | 0.502 |
| GLU | 42 | A | 0.69 | 0.798 | 0.998 | 0.49 |
| MET | 43 | A | 0.752 | 0.746 | 0.784 | 0.714 |
| ALA | 44 | A | 0.595 | 0.599 | 1 | 0.194 |
| VAL | 45 | A | 0.711 | 0.67 | 0.999 | 0.382 |
| GLY | 46 | A | 0.627 | 0.637 | 0.999 | 0.265 |
| ALA | 47 | A | 0.615 | 0.617 | 1 | 0.232 |
| LEU | 48 | A | 0.723 | 0.684 | 0.934 | 0.473 |
| ALA | 49 | A | 0.55 | 0.57 | 0.994 | 0.126 |
| ASN | 50 | A | 0.823 | 0.686 | 0.973 | 0.536 |
| LYS | 51 | A | 0.743 | 0.818 | 0.936 | 0.625 |
| VAL | 52 | A | 0.661 | 0.66 | 0.998 | 0.323 |
| GLY | 53 | A | 0.628 | 0.646 | 0.999 | 0.275 |
| LEU | 54 | A | 0.727 | 0.718 | 0.861 | 0.584 |
| SER | 55 | A | 0.65 | 0.693 | 0.996 | 0.347 |
| GLN | 56 | A | 0.705 | 0.797 | 0.857 | 0.645 |
| SER | 57 | A | 0.706 | 0.747 | 0.998 | 0.455 |
| ALA | 58 | A | 0.679 | 0.646 | 0.997 | 0.328 |
| LEU | 59 | A | 0.726 | 0.754 | 0.999 | 0.481 |
| SER | 60 | A | 0.717 | 0.608 | 0.927 | 0.398 |
| **GLN** | **61** | **A** | **0.858** | **0.651** | **0.752** | **0.757** |
| HIS | 62 | A | 0.86 | 0.813 | 0.954 | 0.719 |
| LEU | 63 | A | 0.72 | 0.726 | 0.997 | 0.449 |
| SER | 64 | A | 0.716 | 0.69 | 0.977 | 0.429 |
| **LYS** | **65** | **A** | **0.771** | **0.735** | **0.703** | **0.803** |
| LEU | 66 | A | 0.735 | 0.742 | 0.999 | 0.478 |
| ARG | 67 | A | 0.784 | 0.752 | 0.85 | 0.686 |
| ALA | 68 | A | 0.539 | 0.543 | 0.999 | 0.083 |
| GLN | 69 | A | 0.804 | 0.784 | 0.999 | 0.589 |
| ASN | 70 | A | 0.725 | 0.793 | 0.992 | 0.526 |
| LEU | 71 | A | 0.738 | 0.699 | 0.894 | 0.543 |
| VAL | 72 | A | 0.671 | 0.666 | 1 | 0.337 |
| SER | 73 | A | 0.648 | 0.637 | 0.912 | 0.373 |
| THR | 74 | A | 0.679 | 0.679 | 1 | 0.358 |
| **ARG** | **75** | **A** | **0.828** | **0.86** | **0.75** | **0.938** |
| ARG | 76 | A | 0.775 | 0.671 | 0.723 | 0.723 |
| ASP | 77 | A | 0.705 | 0.731 | 0.976 | 0.46 |
| ALA | 78 | A | 0.739 | 0.7 | 0.972 | 0.467 |
| GLN | 79 | A | 0.818 | 0.67 | 0.966 | 0.522 |
| THR | 80 | A | 0.732 | 0.74 | 0.985 | 0.487 |
| ILE | 81 | A | 0.763 | 0.685 | 0.968 | 0.48 |
| **TYR** | **82** | **A** | **0.883** | **0.873** | **1** | **0.756** |
| TYR | 83 | A | 0.843 | 0.832 | 0.986 | 0.689 |
| SER | 84 | A | 0.604 | 0.742 | 0.931 | 0.415 |
| SER | 85 | A | 0.59 | 0.61 | 0.997 | 0.203 |
| SER | 86 | A | 0.634 | 0.57 | 0.91 | 0.294 |
| SER | 87 | A | 0.617 | 0.562 | 0.989 | 0.19 |
| ASP | 88 | A | 0.782 | 0.712 | 0.92 | 0.574 |
| SER | 89 | A | 0.686 | 0.714 | 0.947 | 0.453 |
| VAL | 90 | A | 0.698 | 0.68 | 0.998 | 0.38 |
| MET | 91 | A | 0.769 | 0.723 | 0.805 | 0.687 |
| LYS | 92 | A | 0.827 | 0.793 | 0.947 | 0.673 |
| ILE | 93 | A | 0.734 | 0.714 | 1 | 0.448 |
| LEU | 94 | A | 0.674 | 0.713 | 0.999 | 0.388 |
| GLY | 95 | A | 0.642 | 0.64 | 0.999 | 0.283 |
| ALA | 96 | A | 0.596 | 0.583 | 0.999 | 0.18 |
| LEU | 97 | A | 0.73 | 0.697 | 1 | 0.427 |
| SER | 98 | A | 0.639 | 0.674 | 0.979 | 0.334 |
| GLU | 99 | A | 0.74 | 0.772 | 0.996 | 0.516 |
| ILE | 100 | A | 0.743 | 0.721 | 0.993 | 0.471 |
| TYR | 101 | A | 0.853 | 0.842 | 0.999 | 0.696 |
| GLY | 102 | A | 0.722 | 0.787 | 0.851 | 0.658 |
| GLN | 6 | B | 0.809 | 0.702 | 0.766 | 0.745 |
| PRO | 7 | B | 0.683 | 0.692 | 0.985 | 0.39 |
| LEU | 8 | B | 0.755 | 0.706 | 0.998 | 0.463 |
| SER | 9 | B | 0.71 | 0.695 | 0.998 | 0.407 |
| PRO | 10 | B | 0.79 | 0.738 | 0.992 | 0.536 |
| GLU | 11 | B | 0.694 | 0.761 | 0.794 | 0.661 |
| **LYS** | **12** | **B** | **0.792** | **0.787** | **0.731** | **0.848** |
| HIS | 13 | B | 0.813 | 0.796 | 0.947 | 0.662 |
| GLU | 14 | B | 0.776 | 0.704 | 0.798 | 0.682 |
| GLU | 15 | B | 0.83 | 0.743 | 0.951 | 0.622 |
| ALA | 16 | B | 0.533 | 0.546 | 1 | 0.079 |
| GLU | 17 | B | 0.783 | 0.632 | 0.848 | 0.567 |
| ILE | 18 | B | 0.792 | 0.792 | 0.998 | 0.586 |
| ALA | 19 | B | 0.582 | 0.574 | 1 | 0.156 |
| ALA | 20 | B | 0.55 | 0.573 | 1 | 0.123 |
| GLY | 21 | B | 0.638 | 0.64 | 1 | 0.278 |
| PHE | 22 | B | 0.848 | 0.837 | 0.999 | 0.686 |
| LEU | 23 | B | 0.708 | 0.684 | 0.747 | 0.645 |
| SER | 24 | B | 0.725 | 0.692 | 0.999 | 0.418 |
| ALA | 25 | B | 0.535 | 0.548 | 1 | 0.083 |
| MET | 26 | B | 0.739 | 0.751 | 1 | 0.49 |
| ALA | 27 | B | 0.49 | 0.496 | 1 | -0.014 |
| ASN | 28 | B | 0.793 | 0.802 | 1 | 0.595 |
| PRO | 29 | B | 0.735 | 0.752 | 1 | 0.487 |
| LYS | 30 | B | 0.748 | 0.757 | 0.952 | 0.553 |
| ARG | 31 | B | 0.838 | 0.848 | 0.999 | 0.687 |
| LEU | 32 | B | 0.75 | 0.679 | 0.867 | 0.562 |
| LEU | 33 | B | 0.78 | 0.725 | 0.999 | 0.506 |
| ILE | 34 | B | 0.778 | 0.763 | 0.998 | 0.543 |
| LEU | 35 | B | 0.731 | 0.746 | 0.999 | 0.478 |
| ASP | 36 | B | 0.713 | 0.836 | 0.961 | 0.588 |
| SER | 37 | B | 0.693 | 0.697 | 0.997 | 0.393 |
| LEU | 38 | B | 0.747 | 0.744 | 0.997 | 0.494 |
| **VAL** | **39** | **B** | **0.704** | **0.714** | **0.883** | **0.535** |
| LYS | 40 | B | 0.805 | 0.731 | 0.746 | 0.79 |
| GLU | 41 | B | 0.773 | 0.723 | 0.75 | 0.746 |
| GLU | 42 | B | 0.64 | 0.769 | 0.992 | 0.417 |
| MET | 43 | B | 0.731 | 0.671 | 0.757 | 0.645 |
| ALA | 44 | B | 0.611 | 0.6 | 0.999 | 0.212 |
| VAL | 45 | B | 0.693 | 0.659 | 0.993 | 0.359 |
| GLY | 46 | B | 0.637 | 0.631 | 1 | 0.268 |
| ALA | 47 | B | 0.613 | 0.617 | 1 | 0.23 |
| LEU | 48 | B | 0.718 | 0.661 | 0.91 | 0.469 |
| ALA | 49 | B | 0.568 | 0.59 | 0.992 | 0.166 |
| ASN | 50 | B | 0.703 | 0.798 | 0.928 | 0.573 |
| LYS | 51 | B | 0.741 | 0.761 | 0.988 | 0.514 |
| VAL | 52 | B | 0.667 | 0.665 | 0.933 | 0.399 |
| GLY | 53 | B | 0.645 | 0.641 | 0.987 | 0.299 |
| LEU | 54 | B | 0.729 | 0.749 | 0.828 | 0.65 |
| SER | 55 | B | 0.662 | 0.699 | 0.984 | 0.377 |
| **GLN** | **56** | **B** | **0.768** | **0.796** | **0.732** | **0.832** |
| SER | 57 | B | 0.752 | 0.738 | 0.99 | 0.5 |
| ALA | 58 | B | 0.613 | 0.638 | 0.987 | 0.264 |
| LEU | 59 | B | 0.72 | 0.751 | 0.999 | 0.472 |
| SER | 60 | B | 0.702 | 0.595 | 0.964 | 0.333 |
| **GLN** | **61** | **B** | **0.812** | **0.651** | **0.667** | **0.796** |
| HIS | 62 | B | 0.858 | 0.799 | 0.998 | 0.659 |
| LEU | 63 | B | 0.719 | 0.726 | 0.999 | 0.446 |
| SER | 64 | B | 0.711 | 0.715 | 0.97 | 0.456 |
| LYS | 65 | B | 0.818 | 0.757 | 0.945 | 0.63 |
| LEU | 66 | B | 0.748 | 0.745 | 1 | 0.493 |
| **ARG** | **67** | **B** | **0.753** | **0.807** | **0.758** | **0.802** |
| ALA | 68 | B | 0.536 | 0.548 | 0.999 | 0.085 |
| GLN | 69 | B | 0.807 | 0.796 | 0.999 | 0.604 |
| ASN | 70 | B | 0.724 | 0.796 | 0.997 | 0.523 |
| LEU | 71 | B | 0.751 | 0.714 | 1 | 0.465 |
| VAL | 72 | B | 0.67 | 0.658 | 0.998 | 0.33 |
| SER | 73 | B | 0.65 | 0.685 | 0.997 | 0.338 |
| THR | 74 | B | 0.707 | 0.68 | 0.996 | 0.391 |
| **ARG** | **75** | **B** | **0.815** | **0.844** | **0.882** | **0.777** |
| ARG | 76 | B | 0.822 | 0.795 | 0.941 | 0.676 |
| ASP | 77 | B | 0.705 | 0.703 | 0.987 | 0.421 |
| ALA | 78 | B | 0.751 | 0.749 | 0.993 | 0.507 |
| GLN | 79 | B | 0.682 | 0.763 | 0.981 | 0.464 |
| THR | 80 | B | 0.749 | 0.729 | 0.988 | 0.49 |
| ILE | 81 | B | 0.778 | 0.653 | 0.995 | 0.436 |
| TYR | 82 | B | 0.865 | 0.871 | 0.998 | 0.738 |
| TYR | 83 | B | 0.851 | 0.816 | 0.987 | 0.68 |
| SER | 84 | B | 0.616 | 0.657 | 0.997 | 0.276 |
| SER | 85 | B | 0.596 | 0.609 | 0.995 | 0.21 |
| SER | 86 | B | 0.638 | 0.664 | 0.998 | 0.304 |
| SER | 87 | B | 0.617 | 0.612 | 0.999 | 0.23 |
| ASP | 88 | B | 0.839 | 0.758 | 0.918 | 0.679 |
| SER | 89 | B | 0.68 | 0.643 | 1 | 0.323 |
| VAL | 90 | B | 0.704 | 0.693 | 1 | 0.397 |
| MET | 91 | B | 0.763 | 0.723 | 0.802 | 0.684 |
| LYS | 92 | B | 0.783 | 0.799 | 0.903 | 0.679 |
| ILE | 93 | B | 0.745 | 0.703 | 1 | 0.448 |
| LEU | 94 | B | 0.711 | 0.725 | 1 | 0.436 |
| GLY | 95 | B | 0.641 | 0.643 | 0.999 | 0.285 |
| ALA | 96 | B | 0.593 | 0.581 | 1 | 0.174 |
| LEU | 97 | B | 0.693 | 0.713 | 0.999 | 0.407 |
| SER | 98 | B | 0.739 | 0.642 | 0.915 | 0.466 |
| GLU | 99 | B | 0.656 | 0.763 | 0.925 | 0.494 |
| ILE | 100 | B | 0.749 | 0.722 | 0.992 | 0.479 |
| TYR | 101 | B | 0.84 | 0.839 | 0.999 | 0.68 |
| GLY | 102 | B | 0.74 | 0.757 | 0.994 | 0.503 |

### Table S3: 1SWE (P), 1SWA (Q)

| Residue | | | S (P) | S (Q) | J | PRI |
|---|---|---|---|---|---|---|
| GLY | 16 | A | 0.758 | 0.809 | 0.997 | 0.57 |
| ILE | 17 | A | 0.719 | 0.71 | 0.999 | 0.43 |
| THR | 18 | A | 0.747 | 0.741 | 1 | 0.488 |
| GLY | 19 | A | 0.632 | 0.622 | 1 | 0.254 |
| THR | 20 | A | 0.727 | 0.716 | 1 | 0.443 |
| TRP | 21 | A | 0.866 | 0.879 | 1 | 0.745 |
| **TYR** | **22** | **A** | **0.883** | **0.867** | **1** | **0.75** |
| ASN | 23 | A | 0.755 | 0.746 | 1 | 0.501 |
| GLN | 24 | A | 0.743 | 0.748 | 0.997 | 0.494 |
| LEU | 25 | A | 0.731 | 0.703 | 0.967 | 0.467 |
| GLY | 26 | A | 0.644 | 0.655 | 0.999 | 0.3 |
| SER | 27 | A | 0.643 | 0.664 | 1 | 0.307 |
| THR | 28 | A | 0.726 | 0.729 | 0.999 | 0.456 |
| PHE | 29 | A | 0.836 | 0.829 | 0.999 | 0.666 |
| ILE | 30 | A | 0.759 | 0.774 | 0.998 | 0.535 |
| VAL | 31 | A | 0.695 | 0.687 | 1 | 0.382 |
| THR | 32 | A | 0.735 | 0.761 | 0.964 | 0.532 |
| ALA | 33 | A | 0.382 | 0.391 | 0.999 | -0.226 |
| GLY | 34 | A | 0.636 | 0.633 | 1 | 0.269 |
| ALA | 35 | A | 0.648 | 0.56 | 0.999 | 0.209 |
| ASP | 36 | A | 0.767 | 0.744 | 0.992 | 0.519 |
| GLY | 37 | A | 0.626 | 0.624 | 1 | 0.25 |
| ALA | 38 | A | 0.609 | 0.599 | 1 | 0.208 |
| LEU | 39 | A | 0.722 | 0.701 | 1 | 0.423 |
| THR | 40 | A | 0.734 | 0.763 | 1 | 0.497 |
| GLY | 41 | A | 0.625 | 0.63 | 1 | 0.255 |
| THR | 42 | A | 0.702 | 0.737 | 1 | 0.439 |
| TYR | 43 | A | 0.86 | 0.847 | 1 | 0.707 |
| GLU | 44 | A | 0.781 | 0.687 | 0.994 | 0.474 |
| SER | 45 | A | 0.628 | 0.56 | 0.999 | 0.189 |
| ALA | 46 | A | 0.548 | 0.59 | 1 | 0.138 |
| VAL | 47 | A | 0.73 | 0.697 | 1 | 0.427 |
| GLY | 48 | A | 0.632 | 0.634 | 1 | 0.266 |
| ASN | 49 | A | 0.736 | 0.731 | 0.995 | 0.472 |
| ALA | 50 | A | 0.516 | 0.517 | 1 | 0.033 |
| GLU | 51 | A | 0.704 | 0.811 | 0.999 | 0.516 |
| SER | 52 | A | 0.722 | 0.714 | 0.999 | 0.437 |
| ARG | 53 | A | 0.849 | 0.765 | 0.942 | 0.672 |
| **TYR** | **54** | **A** | **0.874** | **0.873** | **1** | **0.747** |
| VAL | 55 | A | 0.635 | 0.665 | 1 | 0.3 |
| LEU | 56 | A | 0.739 | 0.731 | 1 | 0.47 |
| THR | 57 | A | 0.755 | 0.677 | 0.999 | 0.433 |
| GLY | 58 | A | 0.617 | 0.615 | 1 | 0.232 |
| ARG | 59 | A | 0.852 | 0.862 | 0.999 | 0.715 |
| TYR | 60 | A | 0.851 | 0.835 | 1 | 0.686 |
| ASP | 61 | A | 0.698 | 0.698 | 1 | 0.396 |
| SER | 62 | A | 0.526 | 0.544 | 1 | 0.07 |
| ALA | 63 | A | 0.596 | 0.601 | 0.999 | 0.198 |
| PRO | 64 | A | 0.648 | 0.678 | 0.999 | 0.327 |
| ALA | 65 | A | 0.553 | 0.53 | 1 | 0.083 |
| THR | 66 | A | 0.719 | 0.682 | 0.997 | 0.404 |
| ASP | 67 | A | 0.749 | 0.744 | 1 | 0.493 |
| GLY | 68 | A | 0.662 | 0.67 | 1 | 0.332 |
| SER | 69 | A | 0.676 | 0.695 | 0.995 | 0.376 |
| GLY | 70 | A | 0.625 | 0.629 | 1 | 0.254 |
| THR | 71 | A | 0.698 | 0.712 | 1 | 0.41 |
| ALA | 72 | A | 0.481 | 0.467 | 1 | -0.052 |
| LEU | 73 | A | 0.686 | 0.686 | 0.995 | 0.377 |
| GLY | 74 | A | 0.616 | 0.61 | 0.999 | 0.227 |
| TRP | 75 | A | 0.859 | 0.885 | 1 | 0.744 |
| THR | 76 | A | 0.712 | 0.709 | 1 | 0.421 |
| VAL | 77 | A | 0.701 | 0.677 | 1 | 0.378 |
| ALA | 78 | A | 0.423 | 0.421 | 1 | -0.156 |
| **TRP** | **79** | **A** | **0.89** | **0.879** | **1** | **0.769** |
| LYS | 80 | A | 0.744 | 0.815 | 0.958 | 0.601 |

| Residue | | | S (P) | S (Q) | J | PRI |
|---|---|---|---|---|---|---|
| ASN | 81 | A | 0.772 | 0.774 | 1 | 0.546 |
| ASN | 82 | A | 0.646 | 0.676 | 1 | 0.322 |
| TYR | 83 | A | 0.806 | 0.831 | 0.993 | 0.644 |
| **ARG** | **84** | **A** | **0.846** | **0.849** | **0.905** | **0.79** |
| ASN | 85 | A | 0.77 | 0.74 | 1 | 0.51 |
| ALA | 86 | A | 0.479 | 0.439 | 1 | -0.082 |
| HIS | 87 | A | 0.81 | 0.778 | 1 | 0.588 |
| SER | 88 | A | 0.668 | 0.699 | 1 | 0.367 |
| ALA | 89 | A | 0.51 | 0.468 | 1 | -0.022 |
| THR | 90 | A | 0.716 | 0.73 | 1 | 0.446 |
| THR | 91 | A | 0.699 | 0.721 | 1 | 0.42 |
| TRP | 92 | A | 0.869 | 0.876 | 1 | 0.745 |
| SER | 93 | A | 0.629 | 0.61 | 0.999 | 0.24 |
| GLY | 94 | A | 0.623 | 0.628 | 1 | 0.251 |
| GLN | 95 | A | 0.742 | 0.744 | 1 | 0.486 |
| TYR | 96 | A | 0.858 | 0.864 | 1 | 0.722 |
| VAL | 97 | A | 0.706 | 0.705 | 0.996 | 0.415 |
| GLY | 98 | A | 0.63 | 0.616 | 0.999 | 0.247 |
| GLY | 99 | A | 0.632 | 0.64 | 1 | 0.272 |
| ALA | 100 | A | 0.735 | 0.696 | 0.988 | 0.443 |
| GLU | 101 | A | 0.67 | 0.757 | 0.949 | 0.478 |
| ALA | 102 | A | 0.534 | 0.503 | 1 | 0.037 |
| ARG | 103 | A | 0.833 | 0.775 | 0.966 | 0.642 |
| ILE | 104 | A | 0.75 | 0.738 | 1 | 0.488 |
| ASN | 105 | A | 0.667 | 0.748 | 0.999 | 0.416 |
| THR | 106 | A | 0.723 | 0.671 | 1 | 0.394 |
| GLN | 107 | A | 0.71 | 0.783 | 0.997 | 0.496 |
| **TRP** | **108** | **A** | **0.889** | **0.875** | **1** | **0.764** |
| LEU | 109 | A | 0.739 | 0.741 | 1 | 0.48 |
| LEU | 110 | A | 0.754 | 0.766 | 1 | 0.52 |
| THR | 111 | A | 0.663 | 0.705 | 1 | 0.368 |
| SER | 112 | A | 0.568 | 0.577 | 1 | 0.145 |
| GLY | 113 | A | 0.609 | 0.597 | 1 | 0.206 |
| THR | 114 | A | 0.715 | 0.758 | 0.999 | 0.474 |
| THR | 115 | A | 0.687 | 0.687 | 1 | 0.374 |
| **GLU** | **116** | **A** | **0.756** | **0.798** | **0.741** | **0.813** |
| ALA | 117 | A | 0.617 | 0.571 | 0.999 | 0.189 |
| ASN | 118 | A | 0.76 | 0.777 | 0.998 | 0.539 |
| ALA | 119 | A | 0.494 | 0.56 | 1 | 0.054 |
| **TRP** | **120** | **A** | **0.883** | **0.885** | **0.997** | **0.771** |
| LYS | 121 | A | 0.799 | 0.773 | 0.985 | 0.587 |
| SER | 122 | A | 0.659 | 0.646 | 0.999 | 0.306 |
| THR | 123 | A | 0.71 | 0.691 | 1 | 0.401 |
| LEU | 124 | A | 0.701 | 0.715 | 1 | 0.416 |
| VAL | 125 | A | 0.672 | 0.696 | 1 | 0.368 |
| GLY | 126 | A | 0.617 | 0.618 | 1 | 0.235 |
| HIS | 127 | A | 0.838 | 0.83 | 1 | 0.668 |
| ASP | 128 | A | 0.743 | 0.773 | 1 | 0.516 |
| THR | 129 | A | 0.731 | 0.702 | 1 | 0.433 |
| PHE | 130 | A | 0.837 | 0.836 | 1 | 0.673 |
| THR | 131 | A | 0.647 | 0.672 | 0.928 | 0.391 |
| LYS | 132 | A | 0.823 | 0.862 | 0.973 | 0.712 |
| GLY | 16 | B | 0.802 | 0.787 | 0.988 | 0.601 |
| ILE | 17 | B | 0.724 | 0.715 | 0.998 | 0.441 |
| THR | 18 | B | 0.744 | 0.729 | 0.998 | 0.475 |
| GLY | 19 | B | 0.623 | 0.632 | 0.998 | 0.257 |
| THR | 20 | B | 0.719 | 0.726 | 0.998 | 0.447 |
| **TRP** | **21** | **B** | **0.877** | **0.893** | **0.999** | **0.771** |
| **TYR** | **22** | **B** | **0.887** | **0.882** | **0.968** | **0.801** |
| ASN | 23 | B | 0.763 | 0.765 | 0.993 | 0.535 |
| GLN | 24 | B | 0.716 | 0.737 | 0.964 | 0.489 |
| LEU | 25 | B | 0.638 | 0.697 | 0.994 | 0.341 |
| GLY | 26 | B | 0.652 | 0.66 | 1 | 0.312 |
| SER | 27 | B | 0.618 | 0.654 | 0.996 | 0.276 |
| THR | 28 | B | 0.755 | 0.732 | 1 | 0.487 |

| Residue | | | S (P) | S (Q) | J | PRI |
|---|---|---|---|---|---|---|
| PHE | 29 | B | 0.829 | 0.832 | 0.997 | 0.664 |
| ILE | 30 | B | 0.748 | 0.775 | 0.995 | 0.528 |
| VAL | 31 | B | 0.683 | 0.685 | 0.999 | 0.369 |
| THR | 32 | B | 0.733 | 0.759 | 0.999 | 0.493 |
| ALA | 33 | B | 0.377 | 0.386 | 1 | -0.237 |
| GLY | 34 | B | 0.631 | 0.633 | 0.999 | 0.265 |
| ALA | 35 | B | 0.616 | 0.618 | 0.998 | 0.236 |
| ASP | 36 | B | 0.698 | 0.708 | 0.927 | 0.479 |
| GLY | 37 | B | 0.622 | 0.621 | 1 | 0.243 |
| ALA | 38 | B | 0.544 | 0.615 | 1 | 0.159 |
| LEU | 39 | B | 0.667 | 0.714 | 1 | 0.381 |
| THR | 40 | B | 0.757 | 0.716 | 0.999 | 0.474 |
| GLY | 41 | B | 0.631 | 0.625 | 1 | 0.256 |
| THR | 42 | B | 0.762 | 0.727 | 0.999 | 0.49 |
| TYR | 43 | B | 0.859 | 0.835 | 0.997 | 0.697 |
| GLU | 44 | B | 0.796 | 0.796 | 0.984 | 0.608 |
| SER | 45 | B | 0.755 | 0.797 | 0.993 | 0.559 |
| ASN | 49 | B | 0.864 | 0.768 | 0.987 | 0.645 |
| ALA | 50 | B | 0.675 | 0.511 | 0.954 | 0.232 |
| GLU | 51 | B | 0.734 | 0.763 | 0.917 | 0.58 |
| SER | 52 | B | 0.627 | 0.723 | 0.887 | 0.463 |
| ARG | 53 | B | 0.754 | 0.86 | 0.941 | 0.673 |
| TYR | 54 | B | 0.863 | 0.874 | 1 | 0.737 |
| VAL | 55 | B | 0.624 | 0.629 | 1 | 0.253 |
| LEU | 56 | B | 0.741 | 0.749 | 0.997 | 0.493 |
| THR | 57 | B | 0.72 | 0.711 | 1 | 0.431 |
| GLY | 58 | B | 0.618 | 0.613 | 1 | 0.231 |
| ARG | 59 | B | 0.853 | 0.853 | 1 | 0.706 |
| TYR | 60 | B | 0.855 | 0.845 | 1 | 0.7 |
| ASP | 61 | B | 0.705 | 0.703 | 1 | 0.408 |
| SER | 62 | B | 0.497 | 0.616 | 0.999 | 0.114 |
| ALA | 63 | B | 0.555 | 0.566 | 1 | 0.121 |
| PRO | 64 | B | 0.681 | 0.641 | 1 | 0.322 |
| ALA | 65 | B | 0.59 | 0.625 | 0.999 | 0.216 |
| THR | 66 | B | 0.697 | 0.694 | 0.985 | 0.406 |
| ASP | 67 | B | 0.767 | 0.742 | 0.984 | 0.525 |
| GLY | 68 | B | 0.66 | 0.654 | 1 | 0.314 |
| SER | 69 | B | 0.75 | 0.663 | 1 | 0.413 |
| GLY | 70 | B | 0.631 | 0.633 | 0.999 | 0.265 |
| THR | 71 | B | 0.709 | 0.702 | 1 | 0.411 |
| ALA | 72 | B | 0.476 | 0.473 | 1 | -0.051 |
| LEU | 73 | B | 0.694 | 0.702 | 1 | 0.396 |
| GLY | 74 | B | 0.611 | 0.613 | 1 | 0.224 |
| **TRP** | **75** | **B** | **0.873** | **0.88** | **1** | **0.753** |
| THR | 76 | B | 0.717 | 0.705 | 1 | 0.422 |
| VAL | 77 | B | 0.685 | 0.709 | 1 | 0.394 |
| ALA | 78 | B | 0.444 | 0.41 | 1 | -0.146 |
| **TRP** | **79** | **B** | **0.888** | **0.89** | **1** | **0.778** |
| LYS | 80 | B | 0.801 | 0.76 | 0.98 | 0.581 |
| ASN | 81 | B | 0.749 | 0.767 | 0.999 | 0.517 |
| ASN | 82 | B | 0.72 | 0.719 | 0.961 | 0.478 |
| **TYR** | **83** | **B** | **0.846** | **0.832** | **0.919** | **0.759** |
| ARG | 84 | B | 0.809 | 0.835 | 0.921 | 0.723 |
| ASN | 85 | B | 0.725 | 0.767 | 0.999 | 0.493 |
| ALA | 86 | B | 0.628 | 0.445 | 1 | 0.073 |
| HIS | 87 | B | 0.789 | 0.779 | 0.999 | 0.569 |
| SER | 88 | B | 0.7 | 0.668 | 0.997 | 0.371 |
| ALA | 89 | B | 0.48 | 0.495 | 1 | -0.025 |
| THR | 90 | B | 0.738 | 0.723 | 1 | 0.461 |
| THR | 91 | B | 0.721 | 0.706 | 1 | 0.427 |
| **TRP** | **92** | **B** | **0.883** | **0.892** | **1** | **0.775** |
| SER | 93 | B | 0.575 | 0.617 | 1 | 0.192 |
| GLY | 94 | B | 0.629 | 0.619 | 1 | 0.248 |
| GLN | 95 | B | 0.753 | 0.773 | 0.999 | 0.527 |

**Table S3: 1SWE (P), 1SWA (Q)**, continued..

| Residue | | | S (P) | S (Q) | J (P,Q) | PRI |
|---|---|---|---|---|---|---|
| TYR | 96 | B | 0.856 | 0.86 | 1 | 0.716 |
| VAL | 97 | B | 0.7 | 0.705 | 0.999 | 0.406 |
| GLY | 98 | B | 0.614 | 0.609 | 0.999 | 0.224 |
| GLY | 99 | B | 0.645 | 0.635 | 0.999 | 0.281 |
| ALA | 100 | B | 0.743 | 0.73 | 0.979 | 0.494 |
| GLU | 101 | B | 0.74 | 0.764 | 0.929 | 0.575 |
| ALA | 102 | B | 0.51 | 0.476 | 1 | -0.014 |
| ARG | 103 | B | 0.824 | 0.825 | 0.998 | 0.651 |
| ILE | 104 | B | 0.744 | 0.742 | 1 | 0.486 |
| ASN | 105 | B | 0.75 | 0.741 | 1 | 0.491 |
| THR | 106 | B | 0.722 | 0.715 | 1 | 0.437 |
| GLN | 107 | B | 0.742 | 0.764 | 0.997 | 0.509 |
| **TRP** | **108** | **B** | **0.885** | **0.881** | **1** | **0.766** |
| LEU | 109 | B | 0.738 | 0.729 | 1 | 0.467 |
| LEU | 110 | B | 0.738 | 0.744 | 1 | 0.482 |
| THR | 111 | B | 0.731 | 0.697 | 1 | 0.428 |
| SER | 112 | B | 0.598 | 0.584 | 1 | 0.182 |
| GLY | 113 | B | 0.596 | 0.611 | 1 | 0.207 |
| THR | 114 | B | 0.766 | 0.707 | 0.999 | 0.474 |
| THR | 115 | B | 0.612 | 0.578 | 1 | 0.19 |
| GLU | 116 | B | 0.733 | 0.755 | 0.994 | 0.494 |
| ALA | 117 | B | 0.588 | 0.637 | 0.999 | 0.226 |
| ASN | 118 | B | 0.735 | 0.747 | 0.999 | 0.483 |
| ALA | 119 | B | 0.504 | 0.583 | 1 | 0.087 |
| **TRP** | **120** | **B** | **0.882** | **0.87** | **1** | **0.752** |
| LYS | 121 | B | 0.755 | 0.809 | 0.997 | 0.567 |
| SER | 122 | B | 0.625 | 0.616 | 1 | 0.241 |
| THR | 123 | B | 0.704 | 0.687 | 1 | 0.391 |
| LEU | 124 | B | 0.703 | 0.702 | 1 | 0.405 |
| VAL | 125 | B | 0.673 | 0.688 | 1 | 0.361 |
| GLY | 126 | B | 0.616 | 0.617 | 1 | 0.233 |
| HIS | 127 | B | 0.815 | 0.836 | 0.999 | 0.652 |
| ASP | 128 | B | 0.742 | 0.773 | 0.918 | 0.597 |
| THR | 129 | B | 0.733 | 0.714 | 1 | 0.447 |
| **PHE** | **130** | **B** | **0.839** | **0.84** | **0.925** | **0.754** |
| THR | 131 | B | 0.718 | 0.755 | 0.998 | 0.475 |
| LYS | 132 | B | 0.744 | 0.776 | 0.963 | 0.557 |
| **VAL** | **133** | **B** | **0.859** | **0.814** | **0.844** | **0.829** |
| GLY | 16 | C | 0.814 | 0.791 | 1 | 0.605 |
| ILE | 17 | C | 0.709 | 0.708 | 0.999 | 0.418 |
| THR | 18 | C | 0.737 | 0.743 | 0.998 | 0.482 |
| GLY | 19 | C | 0.631 | 0.634 | 1 | 0.265 |
| THR | 20 | C | 0.725 | 0.719 | 0.999 | 0.445 |
| **TRP** | **21** | **C** | **0.883** | **0.874** | **1** | **0.757** |
| **TYR** | **22** | **C** | **0.88** | **0.87** | **0.995** | **0.755** |
| ASN | 23 | C | 0.79 | 0.76 | 0.992 | 0.558 |
| GLN | 24 | C | 0.736 | 0.785 | 0.986 | 0.535 |
| LEU | 25 | C | 0.712 | 0.672 | 0.996 | 0.388 |
| GLY | 26 | C | 0.662 | 0.653 | 0.99 | 0.325 |
| SER | 27 | C | 0.609 | 0.659 | 0.998 | 0.27 |
| THR | 28 | C | 0.73 | 0.73 | 0.995 | 0.465 |
| PHE | 29 | C | 0.832 | 0.84 | 0.973 | 0.699 |
| ILE | 30 | C | 0.765 | 0.765 | 0.995 | 0.535 |
| VAL | 31 | C | 0.64 | 0.718 | 0.998 | 0.36 |
| THR | 32 | C | 0.741 | 0.752 | 0.995 | 0.498 |
| ALA | 33 | C | 0.383 | 0.417 | 0.999 | -0.199 |
| GLY | 34 | C | 0.628 | 0.624 | 1 | 0.252 |
| ALA | 35 | C | 0.619 | 0.588 | 1 | 0.207 |
| ASP | 36 | C | 0.768 | 0.763 | 1 | 0.531 |
| GLY | 37 | C | 0.622 | 0.628 | 1 | 0.25 |
| ALA | 38 | C | 0.558 | 0.583 | 1 | 0.141 |
| LEU | 39 | C | 0.733 | 0.744 | 1 | 0.477 |
| THR | 40 | C | 0.734 | 0.723 | 0.993 | 0.464 |
| GLY | 41 | C | 0.628 | 0.637 | 1 | 0.265 |
| THR | 42 | C | 0.745 | 0.706 | 0.991 | 0.46 |
| TYR | 43 | C | 0.866 | 0.835 | 0.999 | 0.702 |
| GLU | 44 | C | 0.737 | 0.798 | 0.951 | 0.584 |
| SER | 45 | C | 0.755 | 0.804 | 0.839 | 0.72 |
| ASN | 49 | C | 0.861 | 0.791 | 0.986 | 0.666 |
| ALA | 50 | C | 0.583 | 0.495 | 0.952 | 0.126 |
| **GLU** | **51** | **C** | **0.833** | **0.797** | **0.875** | **0.755** |
| **SER** | **52** | **C** | **0.776** | **0.752** | **0.767** | **0.761** |
| **ARG** | **53** | **C** | **0.859** | **0.835** | **0.878** | **0.816** |
| TYR | 54 | C | 0.868 | 0.868 | 0.998 | 0.738 |
| VAL | 55 | C | 0.635 | 0.665 | 1 | 0.3 |
| LEU | 56 | C | 0.728 | 0.737 | 0.998 | 0.467 |
| THR | 57 | C | 0.727 | 0.721 | 0.999 | 0.449 |
| GLY | 58 | C | 0.615 | 0.609 | 1 | 0.224 |
| ARG | 59 | C | 0.852 | 0.837 | 0.999 | 0.69 |
| TYR | 60 | C | 0.861 | 0.867 | 0.999 | 0.729 |
| ASP | 61 | C | 0.699 | 0.687 | 0.999 | 0.387 |
| SER | 62 | C | 0.57 | 0.575 | 0.999 | 0.146 |
| ALA | 63 | C | 0.571 | 0.543 | 1 | 0.114 |
| PRO | 64 | C | 0.655 | 0.653 | 0.999 | 0.309 |
| ALA | 65 | C | 0.529 | 0.545 | 0.999 | 0.075 |
| THR | 66 | C | 0.609 | 0.734 | 0.97 | 0.373 |
| ASP | 67 | C | 0.736 | 0.766 | 0.997 | 0.505 |
| GLY | 68 | C | 0.656 | 0.659 | 1 | 0.315 |
| SER | 69 | C | 0.685 | 0.68 | 1 | 0.365 |
| GLY | 70 | C | 0.626 | 0.617 | 1 | 0.243 |
| THR | 71 | C | 0.704 | 0.704 | 1 | 0.408 |
| ALA | 72 | C | 0.483 | 0.489 | 1 | -0.028 |
| LEU | 73 | C | 0.735 | 0.662 | 1 | 0.397 |
| GLY | 74 | C | 0.613 | 0.608 | 0.999 | 0.222 |
| TRP | 75 | C | 0.848 | 0.891 | 0.999 | 0.74 |
| THR | 76 | C | 0.715 | 0.73 | 1 | 0.445 |
| VAL | 77 | C | 0.696 | 0.69 | 1 | 0.386 |
| ALA | 78 | C | 0.406 | 0.436 | 1 | -0.158 |
| **TRP** | **79** | **C** | **0.888** | **0.892** | **0.999** | **0.781** |
| LYS | 80 | C | 0.795 | 0.818 | 0.994 | 0.619 |
| ASN | 81 | C | 0.732 | 0.76 | 1 | 0.492 |
| ASN | 82 | C | 0.705 | 0.643 | 0.887 | 0.461 |
| TYR | 83 | C | 0.852 | 0.836 | 0.991 | 0.697 |
| **ARG** | **84** | **C** | **0.835** | **0.831** | **0.911** | **0.755** |
| ASN | 85 | C | 0.732 | 0.743 | 0.999 | 0.476 |
| ALA | 86 | C | 0.462 | 0.439 | 0.999 | -0.098 |
| HIS | 87 | C | 0.751 | 0.756 | 1 | 0.507 |
| SER | 88 | C | 0.636 | 0.673 | 0.999 | 0.31 |
| ALA | 89 | C | 0.52 | 0.493 | 1 | 0.013 |
| THR | 90 | C | 0.715 | 0.744 | 0.998 | 0.461 |
| THR | 91 | C | 0.706 | 0.72 | 1 | 0.426 |
| **TRP** | **92** | **C** | **0.88** | **0.884** | **0.999** | **0.765** |
| SER | 93 | C | 0.611 | 0.566 | 1 | 0.177 |
| GLY | 94 | C | 0.621 | 0.627 | 1 | 0.248 |
| GLN | 95 | C | 0.765 | 0.758 | 1 | 0.523 |
| TYR | 96 | C | 0.864 | 0.854 | 1 | 0.718 |
| VAL | 97 | C | 0.712 | 0.658 | 0.999 | 0.371 |
| GLY | 98 | C | 0.625 | 0.633 | 0.986 | 0.272 |
| GLY | 99 | C | 0.63 | 0.654 | 0.995 | 0.289 |
| ALA | 100 | C | 0.743 | 0.691 | 0.988 | 0.446 |
| GLU | 101 | C | 0.769 | 0.709 | 0.852 | 0.626 |
| ALA | 102 | C | 0.545 | 0.533 | 1 | 0.078 |
| ARG | 103 | C | 0.788 | 0.783 | 0.94 | 0.631 |
| ILE | 104 | C | 0.752 | 0.744 | 1 | 0.496 |
| ASN | 105 | C | 0.753 | 0.738 | 0.999 | 0.492 |
| THR | 106 | C | 0.725 | 0.71 | 1 | 0.435 |
| GLN | 107 | C | 0.751 | 0.744 | 0.997 | 0.498 |
| **TRP** | **108** | **C** | **0.876** | **0.877** | **1** | **0.753** |
| LEU | 109 | C | 0.747 | 0.755 | 1 | 0.502 |
| LEU | 110 | C | 0.766 | 0.765 | 1 | 0.531 |
| THR | 111 | C | 0.692 | 0.679 | 1 | 0.371 |
| SER | 112 | C | 0.643 | 0.589 | 0.997 | 0.235 |
| GLY | 113 | C | 0.603 | 0.606 | 0.999 | 0.21 |
| THR | 114 | C | 0.727 | 0.756 | 0.999 | 0.484 |
| THR | 115 | C | 0.628 | 0.629 | 1 | 0.257 |
| GLU | 116 | C | 0.652 | 0.727 | 0.841 | 0.538 |
| ALA | 117 | C | 0.578 | 0.59 | 0.999 | 0.169 |
| ASN | 118 | C | 0.743 | 0.773 | 1 | 0.516 |
| ALA | 119 | C | 0.567 | 0.477 | 1 | 0.044 |
| **TRP** | **120** | **C** | **0.889** | **0.883** | **0.999** | **0.773** |
| LYS | 121 | C | 0.783 | 0.785 | 0.994 | 0.574 |
| SER | 122 | C | 0.604 | 0.573 | 1 | 0.177 |

**Table S3: 1SWE (P), 1SWA (Q)**, continued..

| Residue | | | S (P) | S (Q) | J (P,Q) | PRI |
|---|---|---|---|---|---|---|
| THR | 123 | C | 0.721 | 0.694 | 1 | 0.415 |
| LEU | 124 | C | 0.7 | 0.71 | 0.999 | 0.411 |
| VAL | 125 | C | 0.677 | 0.711 | 1 | 0.388 |
| GLY | 126 | C | 0.62 | 0.613 | 1 | 0.233 |
| HIS | 127 | C | 0.841 | 0.832 | 0.997 | 0.676 |
| ASP | 128 | C | 0.738 | 0.781 | 0.937 | 0.582 |
| THR | 129 | C | 0.718 | 0.711 | 1 | 0.429 |
| PHE | 130 | C | 0.835 | 0.843 | 0.999 | 0.679 |
| THR | 131 | C | 0.718 | 0.758 | 0.965 | 0.511 |
| LYS | 132 | C | 0.761 | 0.744 | 0.999 | 0.506 |
| **VAL** | **133** | **C** | **0.861** | **0.858** | **0.915** | **0.804** |
| GLY | 16 | D | 0.798 | 0.786 | 1 | 0.584 |
| ILE | 17 | D | 0.727 | 0.715 | 0.999 | 0.443 |
| THR | 18 | D | 0.73 | 0.753 | 1 | 0.483 |
| GLY | 19 | D | 0.63 | 0.629 | 0.999 | 0.26 |
| THR | 20 | D | 0.733 | 0.749 | 0.998 | 0.484 |
| **TRP** | **21** | **D** | **0.886** | **0.882** | **0.999** | **0.769** |
| **TYR** | **22** | **D** | **0.888** | **0.866** | **1** | **0.754** |
| ASN | 23 | D | 0.712 | 0.765 | 0.98 | 0.497 |
| GLN | 24 | D | 0.666 | 0.774 | 0.973 | 0.467 |
| LEU | 25 | D | 0.655 | 0.689 | 0.991 | 0.353 |
| GLY | 26 | D | 0.653 | 0.64 | 0.998 | 0.295 |
| SER | 27 | D | 0.571 | 0.591 | 0.978 | 0.184 |
| THR | 28 | D | 0.738 | 0.74 | 0.997 | 0.481 |
| PHE | 29 | D | 0.836 | 0.839 | 0.987 | 0.688 |
| ILE | 30 | D | 0.782 | 0.784 | 0.992 | 0.574 |
| VAL | 31 | D | 0.652 | 0.709 | 0.995 | 0.366 |
| THR | 32 | D | 0.748 | 0.769 | 0.999 | 0.518 |
| ALA | 33 | D | 0.405 | 0.383 | 1 | -0.212 |
| GLY | 34 | D | 0.625 | 0.63 | 0.999 | 0.256 |
| ALA | 35 | D | 0.58 | 0.573 | 0.997 | 0.156 |
| ASP | 36 | D | 0.749 | 0.744 | 0.862 | 0.631 |
| GLY | 37 | D | 0.617 | 0.623 | 1 | 0.24 |
| ALA | 38 | D | 0.624 | 0.595 | 1 | 0.219 |
| LEU | 39 | D | 0.672 | 0.735 | 0.998 | 0.409 |
| THR | 40 | D | 0.702 | 0.716 | 0.999 | 0.419 |
| GLY | 41 | D | 0.627 | 0.619 | 1 | 0.246 |
| THR | 42 | D | 0.68 | 0.715 | 0.984 | 0.411 |
| TYR | 43 | D | 0.86 | 0.843 | 0.995 | 0.708 |
| GLU | 44 | D | 0.797 | 0.773 | 0.943 | 0.627 |
| SER | 45 | D | 0.84 | 0.793 | 0.978 | 0.655 |
| ASN | 49 | D | 0.819 | 0.769 | 0.987 | 0.601 |
| ALA | 50 | D | 0.587 | 0.451 | 0.955 | 0.083 |
| GLU | 51 | D | 0.738 | 0.757 | 0.89 | 0.605 |
| SER | 52 | D | 0.641 | 0.693 | 0.797 | 0.537 |
| ARG | 53 | D | 0.841 | 0.722 | 0.979 | 0.584 |
| TYR | 54 | D | 0.861 | 0.866 | 0.998 | 0.729 |
| VAL | 55 | D | 0.674 | 0.596 | 0.999 | 0.271 |
| LEU | 56 | D | 0.73 | 0.733 | 1 | 0.463 |
| THR | 57 | D | 0.719 | 0.73 | 0.998 | 0.451 |
| GLY | 58 | D | 0.617 | 0.615 | 1 | 0.232 |
| ARG | 59 | D | 0.843 | 0.86 | 1 | 0.703 |
| TYR | 60 | D | 0.849 | 0.854 | 0.999 | 0.704 |
| ASP | 61 | D | 0.685 | 0.678 | 0.998 | 0.365 |
| SER | 62 | D | 0.575 | 0.585 | 0.999 | 0.161 |
| ALA | 63 | D | 0.6 | 0.602 | 1 | 0.202 |
| PRO | 64 | D | 0.694 | 0.682 | 1 | 0.376 |
| ALA | 65 | D | 0.578 | 0.526 | 0.999 | 0.105 |
| THR | 66 | D | 0.677 | 0.699 | 0.996 | 0.38 |
| ASP | 67 | D | 0.753 | 0.766 | 0.998 | 0.521 |
| GLY | 68 | D | 0.66 | 0.66 | 1 | 0.32 |
| SER | 69 | D | 0.646 | 0.757 | 0.997 | 0.406 |
| GLY | 70 | D | 0.627 | 0.625 | 1 | 0.252 |
| THR | 71 | D | 0.708 | 0.715 | 0.999 | 0.424 |
| ALA | 72 | D | 0.479 | 0.486 | 0.999 | -0.034 |
| LEU | 73 | D | 0.724 | 0.69 | 1 | 0.414 |
| GLY | 74 | D | 0.602 | 0.61 | 0.999 | 0.213 |
| **TRP** | **75** | **D** | **0.861** | **0.891** | **0.999** | **0.753** |
| THR | 76 | D | 0.703 | 0.731 | 0.999 | 0.435 |
| VAL | 77 | D | 0.703 | 0.703 | 0.999 | 0.407 |
| ALA | 78 | D | 0.445 | 0.425 | 0.999 | -0.129 |
| **TRP** | **79** | **D** | **0.888** | **0.882** | **0.999** | **0.771** |
| LYS | 80 | D | 0.797 | 0.786 | 0.981 | 0.602 |
| ASN | 81 | D | 0.74 | 0.75 | 0.994 | 0.496 |
| ASN | 82 | D | 0.66 | 0.683 | 0.982 | 0.361 |
| **TYR** | **83** | **D** | **0.83** | **0.852** | **0.915** | **0.767** |
| ARG | 84 | D | 0.65 | 0.815 | 0.844 | 0.621 |
| ASN | 85 | D | 0.754 | 0.724 | 0.999 | 0.479 |
| ALA | 86 | D | 0.445 | 0.45 | 0.997 | -0.102 |
| HIS | 87 | D | 0.767 | 0.797 | 0.999 | 0.565 |
| SER | 88 | D | 0.676 | 0.662 | 1 | 0.338 |
| ALA | 89 | D | 0.518 | 0.511 | 1 | 0.029 |
| THR | 90 | D | 0.745 | 0.74 | 1 | 0.485 |
| THR | 91 | D | 0.709 | 0.726 | 1 | 0.435 |
| **TRP** | **92** | **D** | **0.885** | **0.877** | **0.999** | **0.763** |
| SER | 93 | D | 0.543 | 0.59 | 0.999 | 0.134 |
| GLY | 94 | D | 0.622 | 0.62 | 1 | 0.242 |
| GLN | 95 | D | 0.757 | 0.732 | 0.996 | 0.493 |
| TYR | 96 | D | 0.861 | 0.874 | 1 | 0.735 |
| VAL | 97 | D | 0.698 | 0.716 | 0.998 | 0.416 |
| GLY | 98 | D | 0.62 | 0.618 | 0.999 | 0.239 |
| GLY | 99 | D | 0.643 | 0.642 | 0.994 | 0.291 |
| ALA | 100 | D | 0.638 | 0.661 | 0.897 | 0.402 |
| GLU | 101 | D | 0.695 | 0.738 | 0.842 | 0.591 |
| ALA | 102 | D | 0.463 | 0.503 | 0.998 | -0.032 |
| ARG | 103 | D | 0.848 | 0.802 | 0.962 | 0.688 |
| ILE | 104 | D | 0.744 | 0.747 | 0.999 | 0.492 |
| ASN | 105 | D | 0.72 | 0.729 | 0.999 | 0.45 |
| THR | 106 | D | 0.702 | 0.719 | 1 | 0.421 |
| GLN | 107 | D | 0.755 | 0.765 | 0.991 | 0.529 |
| **TRP** | **108** | **D** | **0.882** | **0.887** | **1** | **0.769** |
| LEU | 109 | D | 0.751 | 0.745 | 1 | 0.496 |
| LEU | 110 | D | 0.765 | 0.741 | 1 | 0.506 |
| THR | 111 | D | 0.713 | 0.702 | 0.999 | 0.416 |
| SER | 112 | D | 0.629 | 0.619 | 0.999 | 0.249 |
| GLY | 113 | D | 0.596 | 0.594 | 1 | 0.19 |
| THR | 114 | D | 0.738 | 0.738 | 1 | 0.476 |
| THR | 115 | D | 0.586 | 0.709 | 1 | 0.295 |
| GLU | 116 | D | 0.767 | 0.754 | 0.967 | 0.554 |
| ALA | 117 | D | 0.565 | 0.549 | 1 | 0.114 |
| ASN | 118 | D | 0.724 | 0.767 | 0.999 | 0.492 |
| ALA | 119 | D | 0.547 | 0.672 | 1 | 0.219 |
| **TRP** | **120** | **D** | **0.901** | **0.885** | **1** | **0.786** |
| LYS | 121 | D | 0.762 | 0.81 | 0.98 | 0.592 |
| SER | 122 | D | 0.614 | 0.599 | 1 | 0.213 |
| THR | 123 | D | 0.706 | 0.714 | 1 | 0.42 |
| LEU | 124 | D | 0.713 | 0.699 | 1 | 0.412 |
| VAL | 125 | D | 0.685 | 0.672 | 1 | 0.357 |
| GLY | 126 | D | 0.627 | 0.612 | 1 | 0.239 |
| HIS | 127 | D | 0.82 | 0.833 | 1 | 0.653 |
| ASP | 128 | D | 0.697 | 0.79 | 0.941 | 0.546 |
| THR | 129 | D | 0.722 | 0.71 | 0.998 | 0.434 |
| **PHE** | **130** | **D** | **0.841** | **0.819** | **0.901** | **0.759** |
| THR | 131 | D | 0.733 | 0.711 | 0.999 | 0.445 |
| LYS | 132 | D | 0.748 | 0.74 | 0.99 | 0.498 |
| VAL | 133 | D | 0.783 | 0.835 | 0.955 | 0.663 |

**Table S4: 5A8B (P), 5DKL (Q)**

| Residue | | | S (P) | S (Q) | J (P,Q) | PRI | Residue | | | S (P) | S (Q) | J (P,Q) | PRI |
|---|---|---|---|---|---|---|---|---|---|---|---|---|---|
| PHE | 239 | A | 0.825 | 0.848 | 0.995 | 0.678 | GLY | 310 | A | 0.635 | 0.636 | 0.999 | 0.272 |
| SER | 240 | A | 0.624 | 0.644 | 0.997 | 0.271 | ASN | 311 | A | 0.761 | 0.76 | 0.994 | 0.527 |
| **PHE** | **241** | **A** | **0.869** | **0.831** | **0.986** | **0.714** | ASP | 312 | A | 0.758 | 0.759 | 0.995 | 0.522 |
| ILE | 242 | A | 0.762 | 0.738 | 0.987 | 0.513 | MET | 313 | A | 0.788 | 0.699 | 0.987 | 0.5 |
| LYS | 243 | A | 0.632 | 0.621 | 0.997 | 0.256 | SER | 314 | A | 0.629 | 0.64 | 0.998 | 0.271 |
| ALA | 244 | A | 0.567 | 0.573 | 0.998 | 0.142 | VAL | 315 | A | 0.72 | 0.737 | 0.996 | 0.461 |
| LEU | 245 | A | 0.727 | 0.735 | 0.994 | 0.468 | CYS | 316 | A | 0.577 | 0.528 | 0.999 | 0.106 |
| GLN | 246 | A | 0.817 | 0.793 | 0.982 | 0.628 | LEU | 317 | A | 0.723 | 0.687 | 0.998 | 0.412 |
| THR | 247 | A | 0.761 | 0.766 | 0.998 | 0.529 | LEU | 318 | A | 0.76 | 0.755 | 0.998 | 0.517 |
| ALA | 248 | A | 0.551 | 0.515 | 0.994 | 0.072 | ASN | 319 | A | 0.737 | 0.746 | 0.998 | 0.485 |
| GLN | 249 | A | 0.759 | 0.699 | 0.971 | 0.487 | **TYR** | **320** | **A** | **0.867** | **0.868** | **0.992** | **0.743** |
| GLN | 250 | A | 0.745 | 0.728 | 0.995 | 0.478 | ARG | 321 | A | 0.837 | 0.828 | 0.996 | 0.669 |
| ASN | 251 | A | 0.754 | 0.772 | 0.995 | 0.531 | VAL | 322 | A | 0.695 | 0.707 | 0.997 | 0.405 |
| PHE | 252 | A | 0.829 | 0.843 | 0.991 | 0.681 | ASP | 323 | A | 0.744 | 0.746 | 0.999 | 0.491 |
| VAL | 253 | A | 0.674 | 0.698 | 0.999 | 0.373 | GLY | 324 | A | 0.651 | 0.649 | 0.999 | 0.301 |
| VAL | 254 | A | 0.728 | 0.721 | 0.998 | 0.451 | THR | 325 | A | 0.774 | 0.744 | 0.999 | 0.519 |
| THR | 255 | A | 0.703 | 0.699 | 0.997 | 0.405 | THR | 326 | A | 0.652 | 0.688 | 0.999 | 0.341 |
| ASP | 256 | A | 0.735 | 0.757 | 0.997 | 0.495 | PHE | 327 | A | 0.837 | 0.83 | 0.994 | 0.673 |
| PRO | 257 | A | 0.691 | 0.654 | 0.999 | 0.346 | **TRP** | **328** | **A** | **0.87** | **0.863** | **0.992** | **0.741** |
| SER | 258 | A | 0.738 | 0.675 | 1 | 0.413 | ASN | 329 | A | 0.742 | 0.727 | 0.991 | 0.478 |
| LEU | 259 | A | 0.737 | 0.77 | 0.999 | 0.508 | GLN | 330 | A | 0.81 | 0.789 | 0.988 | 0.611 |
| PRO | 260 | A | 0.746 | 0.733 | 0.999 | 0.48 | PHE | 331 | A | 0.828 | 0.854 | 0.998 | 0.684 |
| ASP | 261 | A | 0.776 | 0.797 | 0.999 | 0.574 | **PHE** | **332** | **A** | **0.865** | **0.845** | **0.996** | **0.714** |
| ASN | 262 | A | 0.8 | 0.761 | 0.999 | 0.562 | ILE | 333 | A | 0.753 | 0.744 | 0.998 | 0.499 |
| PRO | 263 | A | 0.68 | 0.67 | 0.998 | 0.352 | ALA | 334 | A | 0.528 | 0.521 | 0.997 | 0.052 |
| ILE | 264 | A | 0.767 | 0.764 | 0.995 | 0.536 | ALA | 335 | A | 0.499 | 0.405 | 0.998 | -0.094 |
| VAL | 265 | A | 0.709 | 0.692 | 0.999 | 0.402 | LEU | 336 | A | 0.735 | 0.736 | 0.997 | 0.474 |
| TYR | 266 | A | 0.879 | 0.816 | 0.995 | 0.7 | ARG | 337 | A | 0.781 | 0.796 | 0.998 | 0.579 |
| ALA | 267 | A | 0.471 | 0.452 | 0.997 | -0.074 | ASP | 338 | A | 0.8 | 0.771 | 0.999 | 0.572 |
| SER | 268 | A | 0.676 | 0.695 | 0.998 | 0.373 | ALA | 339 | A | 0.473 | 0.614 | 1 | 0.087 |
| GLN | 269 | A | 0.789 | 0.738 | 0.996 | 0.531 | GLY | 340 | A | 0.654 | 0.641 | 0.999 | 0.296 |
| GLY | 270 | A | 0.632 | 0.63 | 0.991 | 0.271 | GLY | 341 | A | 0.648 | 0.618 | 1 | 0.266 |
| PHE | 271 | A | 0.853 | 0.836 | 0.996 | 0.693 | ASN | 342 | A | 0.802 | 0.749 | 0.998 | 0.553 |
| LEU | 272 | A | 0.777 | 0.715 | 0.994 | 0.498 | VAL | 343 | A | 0.677 | 0.672 | 0.999 | 0.35 |
| ASN | 273 | A | 0.756 | 0.814 | 0.993 | 0.577 | THR | 344 | A | 0.636 | 0.746 | 1 | 0.382 |
| LEU | 274 | A | 0.725 | 0.71 | 0.996 | 0.439 | ASN | 345 | A | 0.802 | 0.839 | 0.998 | 0.643 |
| THR | 275 | A | 0.711 | 0.729 | 0.998 | 0.442 | PHE | 346 | A | 0.84 | 0.855 | 0.995 | 0.7 |
| GLY | 276 | A | 0.633 | 0.629 | 0.997 | 0.265 | VAL | 347 | A | 0.712 | 0.703 | 0.997 | 0.418 |
| **TYR** | **277** | **A** | **0.86** | **0.864** | **0.988** | **0.736** | GLY | 348 | A | 0.62 | 0.617 | 0.997 | 0.24 |
| SER | 278 | A | 0.746 | 0.684 | 0.999 | 0.431 | VAL | 349 | A | 0.68 | 0.709 | 0.998 | 0.391 |
| LEU | 279 | A | 0.676 | 0.649 | 0.997 | 0.328 | GLN | 350 | A | 0.792 | 0.801 | 0.999 | 0.594 |
| ASP | 280 | A | 0.785 | 0.646 | 1 | 0.431 | CYS | 351 | A | 0.61 | 0.583 | 0.998 | 0.195 |
| GLN | 281 | A | 0.798 | 0.737 | 0.996 | 0.539 | LYS | 352 | A | 0.724 | 0.753 | 0.996 | 0.481 |
| ILE | 282 | A | 0.753 | 0.755 | 0.998 | 0.51 | VAL | 353 | A | 0.678 | 0.668 | 0.999 | 0.347 |
| LEU | 283 | A | 0.735 | 0.687 | 0.995 | 0.427 | SER | 354 | A | 0.736 | 0.702 | 0.998 | 0.44 |
| GLY | 284 | A | 0.642 | 0.646 | 1 | 0.288 | ASP | 355 | A | 0.591 | 0.599 | 0.997 | 0.193 |
| ARG | 285 | A | 0.766 | 0.866 | 0.999 | 0.633 | GLN | 356 | A | 0.726 | 0.739 | 0.998 | 0.467 |
| ASN | 286 | A | 0.774 | 0.775 | 0.997 | 0.552 | **TYR** | **357** | **A** | **0.871** | **0.865** | **0.999** | **0.737** |
| CYS | 287 | A | 0.515 | 0.615 | 0.998 | 0.132 | ALA | 358 | A | 0.578 | 0.53 | 0.997 | 0.111 |
| ARG | 288 | A | 0.765 | 0.825 | 0.992 | 0.598 | ALA | 359 | A | 0.675 | 0.653 | 0.998 | 0.33 |
| PHE | 289 | A | 0.812 | 0.846 | 0.997 | 0.661 | THR | 360 | A | 0.76 | 0.755 | 0.999 | 0.516 |
| LEU | 290 | A | 0.751 | 0.759 | 0.997 | 0.513 | VAL | 361 | A | 0.715 | 0.707 | 0.999 | 0.423 |
| GLN | 291 | A | 0.803 | 0.776 | 0.993 | 0.586 | THR | 362 | A | 0.759 | 0.715 | 0.997 | 0.477 |
| GLY | 292 | A | 0.63 | 0.628 | 0.997 | 0.261 | LYS | 363 | A | 0.797 | 0.636 | 0.999 | 0.434 |
| PRO | 293 | A | 0.75 | 0.706 | 0.995 | 0.461 | GLN | 364 | A | 0.718 | 0.776 | 0.999 | 0.495 |
| GLU | 294 | A | 0.65 | 0.7 | 0.999 | 0.351 | GLN | 365 | A | 0.783 | 0.763 | 0.997 | 0.549 |
| THR | 295 | A | 0.71 | 0.707 | 0.995 | 0.422 | **GLU** | **366** | **A** | **0.844** | **0.882** | **0.998** | **0.728** |
| ASP | 296 | A | 0.726 | 0.759 | 0.996 | 0.489 | SER | 240 | B | 0.657 | 0.716 | 0.998 | 0.375 |
| PRO | 297 | A | 0.787 | 0.721 | 0.999 | 0.509 | PHE | 241 | B | 0.826 | 0.789 | 0.978 | 0.637 |
| LYS | 298 | A | 0.604 | 0.752 | 0.998 | 0.358 | ILE | 242 | B | 0.72 | 0.7 | 0.991 | 0.429 |
| ALA | 299 | A | 0.61 | 0.559 | 0.999 | 0.17 | LYS | 243 | B | 0.641 | 0.621 | 0.999 | 0.263 |
| VAL | 300 | A | 0.741 | 0.701 | 0.997 | 0.445 | ALA | 244 | B | 0.571 | 0.595 | 1 | 0.166 |
| GLU | 301 | A | 0.726 | 0.753 | 0.995 | 0.484 | LEU | 245 | B | 0.638 | 0.766 | 0.998 | 0.406 |
| ARG | 302 | A | 0.857 | 0.753 | 0.994 | 0.616 | GLN | 246 | B | 0.797 | 0.665 | 0.996 | 0.466 |
| ILE | 303 | A | 0.722 | 0.735 | 0.994 | 0.463 | THR | 247 | B | 0.71 | 0.766 | 0.998 | 0.478 |
| ARG | 304 | A | 0.846 | 0.827 | 0.999 | 0.674 | ALA | 248 | B | 0.463 | 0.489 | 0.999 | -0.047 |
| LYS | 305 | A | 0.782 | 0.763 | 0.995 | 0.55 | GLN | 249 | B | 0.756 | 0.683 | 0.99 | 0.449 |
| ALA | 306 | A | 0.599 | 0.557 | 0.999 | 0.157 | GLN | 250 | B | 0.785 | 0.785 | 0.986 | 0.584 |
| ILE | 307 | A | 0.735 | 0.755 | 0.995 | 0.495 | ASN | 251 | B | 0.765 | 0.757 | 0.998 | 0.524 |
| GLU | 308 | A | 0.67 | 0.795 | 0.999 | 0.466 | PHE | 252 | B | 0.832 | 0.85 | 0.994 | 0.688 |
| GLN | 309 | A | 0.782 | 0.789 | 0.999 | 0.572 | VAL | 253 | B | 0.729 | 0.678 | 0.999 | 0.408 |

**Table S4: 5A8B (P), 5DKL (Q),** continued..

| Residue | | | S (P) | S (Q) | J (P,Q) | PRI |
|---|---|---|---|---|---|---|
| VAL | 254 | B | 0.727 | 0.718 | 0.999 | 0.446 |
| THR | 255 | B | 0.674 | 0.685 | 0.998 | 0.361 |
| ASP | 256 | B | 0.777 | 0.747 | 0.998 | 0.526 |
| PRO | 257 | B | 0.636 | 0.649 | 0.999 | 0.286 |
| SER | 258 | B | 0.661 | 0.639 | 1 | 0.3 |
| LEU | 259 | B | 0.767 | 0.752 | 1 | 0.519 |
| PRO | 260 | B | 0.732 | 0.739 | 0.999 | 0.472 |
| ASP | 261 | B | 0.807 | 0.749 | 0.999 | 0.557 |
| ASN | 262 | B | 0.755 | 0.759 | 0.999 | 0.515 |
| PRO | 263 | B | 0.656 | 0.667 | 0.999 | 0.324 |
| ILE | 264 | B | 0.774 | 0.746 | 0.996 | 0.524 |
| VAL | 265 | B | 0.716 | 0.678 | 0.999 | 0.395 |
| **TYR** | **266** | **B** | **0.863** | **0.868** | **0.997** | **0.734** |
| ALA | 267 | B | 0.533 | 0.456 | 0.998 | -0.009 |
| SER | 268 | B | 0.687 | 0.707 | 0.999 | 0.395 |
| GLN | 269 | B | 0.711 | 0.721 | 0.996 | 0.436 |
| GLY | 270 | B | 0.635 | 0.635 | 0.993 | 0.277 |
| **PHE** | **271** | **B** | **0.855** | **0.848** | **0.997** | **0.706** |
| LEU | 272 | B | 0.743 | 0.768 | 0.993 | 0.518 |
| ASN | 273 | B | 0.782 | 0.821 | 0.991 | 0.612 |
| LEU | 274 | B | 0.721 | 0.732 | 0.994 | 0.459 |
| THR | 275 | B | 0.737 | 0.699 | 0.999 | 0.437 |
| GLY | 276 | B | 0.624 | 0.643 | 0.998 | 0.269 |
| **TYR** | **277** | **B** | **0.863** | **0.875** | **0.985** | **0.753** |
| SER | 278 | B | 0.729 | 0.691 | 0.998 | 0.422 |
| LEU | 279 | B | 0.749 | 0.729 | 0.997 | 0.481 |
| ASP | 280 | B | 0.619 | 0.677 | 0.998 | 0.298 |
| GLN | 281 | B | 0.76 | 0.783 | 0.997 | 0.546 |
| ILE | 282 | B | 0.763 | 0.764 | 0.997 | 0.53 |
| LEU | 283 | B | 0.748 | 0.75 | 0.997 | 0.501 |
| GLY | 284 | B | 0.641 | 0.641 | 0.999 | 0.283 |
| ARG | 285 | B | 0.807 | 0.685 | 0.998 | 0.494 |
| ASN | 286 | B | 0.778 | 0.77 | 0.998 | 0.55 |
| CYS | 287 | B | 0.49 | 0.604 | 0.998 | 0.096 |
| ARG | 288 | B | 0.845 | 0.812 | 0.991 | 0.666 |
| PHE | 289 | B | 0.824 | 0.837 | 0.998 | 0.663 |
| LEU | 290 | B | 0.765 | 0.729 | 0.999 | 0.495 |
| GLN | 291 | B | 0.781 | 0.776 | 0.992 | 0.565 |
| GLY | 292 | B | 0.632 | 0.629 | 0.997 | 0.264 |
| PRO | 293 | B | 0.704 | 0.715 | 0.995 | 0.424 |
| GLU | 294 | B | 0.688 | 0.654 | 0.998 | 0.344 |
| THR | 295 | B | 0.711 | 0.707 | 0.995 | 0.423 |
| ASP | 296 | B | 0.699 | 0.67 | 0.996 | 0.373 |
| PRO | 297 | B | 0.772 | 0.739 | 0.999 | 0.512 |
| LYS | 298 | B | 0.648 | 0.602 | 0.997 | 0.253 |
| ALA | 299 | B | 0.596 | 0.562 | 0.999 | 0.159 |
| VAL | 300 | B | 0.73 | 0.718 | 0.997 | 0.451 |
| GLU | 301 | B | 0.72 | 0.771 | 0.993 | 0.498 |
| ARG | 302 | B | 0.84 | 0.787 | 0.998 | 0.629 |
| ILE | 303 | B | 0.754 | 0.739 | 0.993 | 0.5 |
| ARG | 304 | B | 0.859 | 0.831 | 0.999 | 0.691 |
| LYS | 305 | B | 0.726 | 0.768 | 0.996 | 0.498 |
| ALA | 306 | B | 0.599 | 0.554 | 0.999 | 0.154 |
| ILE | 307 | B | 0.698 | 0.748 | 0.995 | 0.451 |
| GLU | 308 | B | 0.765 | 0.806 | 0.999 | 0.572 |
| GLN | 309 | B | 0.821 | 0.732 | 0.998 | 0.555 |
| GLY | 310 | B | 0.635 | 0.627 | 0.999 | 0.263 |
| ASN | 311 | B | 0.768 | 0.729 | 0.992 | 0.505 |
| ASP | 312 | B | 0.689 | 0.774 | 0.995 | 0.468 |
| MET | 313 | B | 0.755 | 0.715 | 0.986 | 0.484 |
| SER | 314 | B | 0.626 | 0.64 | 0.998 | 0.268 |
| VAL | 315 | B | 0.687 | 0.735 | 0.995 | 0.427 |
| CYS | 316 | B | 0.565 | 0.511 | 0.999 | 0.077 |
| LEU | 317 | B | 0.73 | 0.691 | 0.997 | 0.424 |
| LEU | 318 | B | 0.745 | 0.763 | 0.999 | 0.509 |
| ASN | 319 | B | 0.746 | 0.74 | 0.998 | 0.488 |
| **TYR** | **320** | **B** | **0.874** | **0.872** | **0.997** | **0.749** |
| ARG | 321 | B | 0.846 | 0.818 | 0.997 | 0.667 |
| VAL | 322 | B | 0.71 | 0.701 | 0.996 | 0.415 |
| ASP | 323 | B | 0.741 | 0.702 | 0.994 | 0.449 |
| GLY | 324 | B | 0.659 | 0.652 | 0.999 | 0.312 |

| Residue | | | S (P) | S (Q) | J (P,Q) | PRI |
|---|---|---|---|---|---|---|
| THR | 325 | B | 0.796 | 0.732 | 0.997 | 0.531 |
| THR | 326 | B | 0.714 | 0.694 | 0.999 | 0.409 |
| PHE | 327 | B | 0.834 | 0.829 | 0.99 | 0.673 |
| **TRP** | **328** | **B** | **0.879** | **0.879** | **0.993** | **0.765** |
| ASN | 329 | B | 0.76 | 0.72 | 0.994 | 0.486 |
| GLN | 330 | B | 0.79 | 0.797 | 0.988 | 0.599 |
| PHE | 331 | B | 0.83 | 0.852 | 0.998 | 0.684 |
| PHE | 332 | B | 0.853 | 0.846 | 0.996 | 0.703 |
| ILE | 333 | B | 0.733 | 0.752 | 0.997 | 0.488 |
| ALA | 334 | B | 0.457 | 0.503 | 0.996 | -0.036 |
| ALA | 335 | B | 0.485 | 0.392 | 0.998 | -0.121 |
| LEU | 336 | B | 0.741 | 0.729 | 0.998 | 0.472 |
| ARG | 337 | B | 0.791 | 0.814 | 0.997 | 0.608 |
| ASP | 338 | B | 0.779 | 0.763 | 0.999 | 0.543 |
| ALA | 339 | B | 0.616 | 0.613 | 0.999 | 0.23 |
| GLY | 340 | B | 0.663 | 0.642 | 0.999 | 0.306 |
| GLY | 341 | B | 0.651 | 0.612 | 1 | 0.263 |
| ASN | 342 | B | 0.764 | 0.73 | 0.998 | 0.496 |
| VAL | 343 | B | 0.679 | 0.684 | 0.999 | 0.364 |
| THR | 344 | B | 0.73 | 0.731 | 0.999 | 0.462 |
| ASN | 345 | B | 0.781 | 0.83 | 0.999 | 0.612 |
| PHE | 346 | B | 0.842 | 0.854 | 0.996 | 0.7 |
| VAL | 347 | B | 0.669 | 0.704 | 0.997 | 0.376 |
| GLY | 348 | B | 0.617 | 0.615 | 0.998 | 0.234 |
| VAL | 349 | B | 0.689 | 0.706 | 0.999 | 0.396 |
| GLN | 350 | B | 0.791 | 0.792 | 0.998 | 0.585 |
| CYS | 351 | B | 0.686 | 0.573 | 0.999 | 0.26 |
| LYS | 352 | B | 0.736 | 0.778 | 0.992 | 0.522 |
| VAL | 353 | B | 0.664 | 0.666 | 0.999 | 0.331 |
| SER | 354 | B | 0.743 | 0.735 | 0.998 | 0.48 |
| ASP | 355 | B | 0.59 | 0.583 | 0.997 | 0.176 |
| GLN | 356 | B | 0.644 | 0.771 | 0.997 | 0.418 |
| **TYR** | **357** | **B** | **0.861** | **0.866** | **0.999** | **0.728** |
| ALA | 358 | B | 0.6 | 0.553 | 0.997 | 0.156 |
| ALA | 359 | B | 0.705 | 0.649 | 0.998 | 0.356 |
| THR | 360 | B | 0.704 | 0.711 | 0.999 | 0.416 |
| VAL | 361 | B | 0.729 | 0.691 | 0.998 | 0.422 |
| THR | 362 | B | 0.572 | 0.711 | 0.997 | 0.286 |
| LYS | 363 | B | 0.769 | 0.73 | 0.999 | 0.5 |
| GLN | 364 | B | 0.68 | 0.778 | 1 | 0.458 |
| GLN | 365 | B | 0.796 | 0.753 | 0.999 | 0.55 |
| GLU | 366 | B | 0.768 | 0.788 | 0.997 | 0.559 |
| GLU | 367 | B | 0.786 | 0.778 | 1 | 0.564 |
| GLU | 368 | B | 0.745 | 0.74 | 0.999 | 0.486 |
| GLU | 369 | B | 0.78 | 0.672 | 0.997 | 0.455 |
| GLU | 370 | B | 0.793 | 0.786 | 0.997 | 0.582 |

| Table S5: 7AMN (P), 7AMT (Q) | | | | | | | | | | | | | |
|---|---|---|---|---|---|---|---|---|---|---|---|---|---|
| **Residue** | | | **S (P)** | **S (Q)** | **J (P,Q)** | **PRI** | **Residue** | | | **S (P)** | **S (Q)** | **J (P,Q)** | **PRI** |
| THR | 10 | A | 0.729 | 0.708 | 0.952 | 0.485 | ASP | 81 | A | 0.759 | 0.787 | 0.843 | 0.703 |
| ARG | 11 | A | 0.69 | 0.705 | 0.992 | 0.403 | ASN | 82 | A | 0.744 | 0.738 | 0.998 | 0.484 |
| LEU | 12 | A | 0.727 | 0.721 | 0.99 | 0.458 | ILE | 83 | A | 0.75 | 0.717 | 0.999 | 0.468 |
| SER | 13 | A | 0.726 | 0.715 | 0.885 | 0.556 | ASP | 84 | A | 0.691 | 0.754 | 0.995 | 0.45 |
| PRO | 14 | A | 0.734 | 0.739 | 0.999 | 0.474 | LEU | 85 | A | 0.706 | 0.709 | 0.986 | 0.429 |
| ILE | 15 | A | 0.632 | 0.663 | 0.999 | 0.296 | ASP | 86 | A | 0.622 | 0.632 | 0.706 | 0.548 |
| LYS | 16 | A | 0.749 | 0.798 | 0.799 | 0.748 | **LEU** | **87** | **A** | **0.736** | **0.779** | **0.627** | **0.888** |
| ARG | 17 | A | 0.818 | 0.828 | 0.999 | 0.647 | HIS | 88 | A | 0.85 | 0.76 | 0.876 | 0.734 |
| LYS | 18 | A | 0.686 | 0.689 | 1 | 0.375 | ALA | 89 | A | 0.595 | 0.64 | 0.991 | 0.244 |
| GLN | 19 | A | 0.839 | 0.827 | 0.989 | 0.677 | ARG | 90 | A | 0.776 | 0.776 | 0.829 | 0.723 |
| GLN | 20 | A | 0.782 | 0.776 | 0.994 | 0.564 | GLU | 91 | A | 0.794 | 0.744 | 0.964 | 0.574 |
| LEU | 21 | A | 0.707 | 0.735 | 0.997 | 0.445 | ASN | 92 | A | 0.806 | 0.795 | 0.998 | 0.603 |
| MET | 22 | A | 0.713 | 0.729 | 1 | 0.442 | ILE | 93 | A | 0.765 | 0.76 | 0.996 | 0.529 |
| GLU | 23 | A | 0.631 | 0.633 | 0.999 | 0.265 | ALA | 94 | A | 0.635 | 0.647 | 0.999 | 0.283 |
| ILE | 24 | A | 0.768 | 0.767 | 1 | 0.535 | ASN | 95 | A | 0.77 | 0.763 | 0.999 | 0.534 |
| ALA | 25 | A | 0.566 | 0.564 | 1 | 0.13 | ILE | 96 | A | 0.754 | 0.754 | 0.998 | 0.51 |
| LEU | 26 | A | 0.751 | 0.759 | 0.998 | 0.512 | THR | 97 | A | 0.718 | 0.72 | 0.999 | 0.439 |
| GLU | 27 | A | 0.723 | 0.736 | 0.99 | 0.469 | ASN | 98 | A | 0.731 | 0.796 | 0.993 | 0.534 |
| VAL | 28 | A | 0.708 | 0.71 | 1 | 0.418 | ALA | 99 | A | 0.594 | 0.582 | 0.999 | 0.177 |
| PHE | 29 | A | 0.837 | 0.835 | 0.986 | 0.686 | MET | 100 | A | 0.691 | 0.743 | 0.997 | 0.437 |
| ALA | 30 | A | 0.509 | 0.502 | 1 | 0.011 | ILE | 101 | A | 0.779 | 0.78 | 0.965 | 0.594 |
| ARG | 31 | A | 0.645 | 0.731 | 0.874 | 0.502 | GLU | 102 | A | 0.692 | 0.789 | 0.708 | 0.773 |
| ARG | 32 | A | 0.742 | 0.782 | 0.999 | 0.525 | LEU | 103 | A | 0.74 | 0.737 | 1 | 0.477 |
| GLY | 33 | A | 0.62 | 0.617 | 0.999 | 0.238 | VAL | 104 | A | 0.711 | 0.703 | 0.997 | 0.417 |
| ILE | 34 | A | 0.718 | 0.715 | 0.999 | 0.434 | SER | 105 | A | 0.666 | 0.65 | 1 | 0.316 |
| GLY | 35 | A | 0.622 | 0.626 | 1 | 0.248 | GLN | 106 | A | 0.756 | 0.722 | 0.894 | 0.584 |
| **ARG** | **36** | **A** | **0.821** | **0.84** | **0.853** | **0.808** | ASP | 107 | A | 0.615 | 0.651 | 0.99 | 0.276 |
| GLY | 37 | A | 0.609 | 0.612 | 0.998 | 0.223 | CYS | 108 | A | 0.623 | 0.654 | 1 | 0.277 |
| GLY | 38 | A | 0.626 | 0.626 | 0.998 | 0.254 | HIS | 109 | A | 0.82 | 0.821 | 0.999 | 0.642 |
| HIS | 39 | A | 0.817 | 0.821 | 0.999 | 0.639 | TRP | 110 | A | 0.871 | 0.88 | 1 | 0.751 |
| ALA | 40 | A | 0.652 | 0.648 | 1 | 0.3 | LEU | 111 | A | 0.733 | 0.729 | 1 | 0.462 |
| ASP | 41 | A | 0.738 | 0.745 | 0.997 | 0.486 | LYS | 112 | A | 0.839 | 0.81 | 0.993 | 0.656 |
| ILE | 42 | A | 0.742 | 0.752 | 0.999 | 0.495 | VAL | 113 | A | 0.677 | 0.692 | 0.999 | 0.37 |
| ALA | 43 | A | 0.573 | 0.58 | 0.999 | 0.154 | TRP | 114 | A | 0.863 | 0.867 | 1 | 0.73 |
| GLU | 44 | A | 0.648 | 0.694 | 0.855 | 0.487 | PHE | 115 | A | 0.835 | 0.841 | 1 | 0.676 |
| ILE | 45 | A | 0.74 | 0.742 | 0.996 | 0.486 | GLU | 116 | A | 0.792 | 0.785 | 1 | 0.577 |
| ALA | 46 | A | 0.433 | 0.492 | 0.999 | -0.074 | TRP | 117 | A | 0.863 | 0.872 | 1 | 0.735 |
| GLN | 47 | A | 0.644 | 0.684 | 0.95 | 0.378 | SER | 118 | A | 0.675 | 0.654 | 1 | 0.329 |
| VAL | 48 | A | 0.699 | 0.717 | 0.999 | 0.417 | ALA | 119 | A | 0.407 | 0.43 | 1 | -0.163 |
| SER | 49 | A | 0.735 | 0.731 | 1 | 0.466 | SER | 120 | A | 0.557 | 0.551 | 1 | 0.108 |
| VAL | 50 | A | 0.7 | 0.706 | 0.999 | 0.407 | THR | 121 | A | 0.671 | 0.627 | 0.998 | 0.3 |
| ALA | 51 | A | 0.629 | 0.635 | 1 | 0.264 | ARG | 122 | A | 0.813 | 0.827 | 0.999 | 0.641 |
| THR | 52 | A | 0.723 | 0.714 | 1 | 0.437 | ASP | 123 | A | 0.558 | 0.563 | 0.99 | 0.131 |
| VAL | 53 | A | 0.665 | 0.681 | 0.999 | 0.347 | GLU | 124 | A | 0.73 | 0.739 | 0.993 | 0.476 |
| **PHE** | **54** | **A** | **0.88** | **0.869** | **0.864** | **0.885** | VAL | 125 | A | 0.705 | 0.714 | 1 | 0.419 |
| ASN | 55 | A | 0.704 | 0.754 | 0.961 | 0.497 | TRP | 126 | A | 0.874 | 0.872 | 1 | 0.746 |
| **TYR** | **56** | **A** | **0.863** | **0.851** | **0.847** | **0.867** | PRO | 127 | A | 0.692 | 0.689 | 1 | 0.381 |
| PHE | 57 | A | 0.825 | 0.833 | 1 | 0.658 | LEU | 128 | A | 0.721 | 0.742 | 0.999 | 0.464 |
| PRO | 58 | A | 0.681 | 0.684 | 0.998 | 0.367 | PHE | 129 | A | 0.822 | 0.822 | 1 | 0.644 |
| **THR** | **59** | **A** | **0.789** | **0.787** | **0.61** | **0.966** | VAL | 130 | A | 0.699 | 0.705 | 0.995 | 0.409 |
| **ARG** | **60** | **A** | **0.87** | **0.852** | **0.749** | **0.973** | SER | 131 | A | 0.672 | 0.694 | 0.999 | 0.367 |
| GLU | 61 | A | 0.607 | 0.719 | 0.679 | 0.647 | THR | 132 | A | 0.72 | 0.715 | 0.997 | 0.438 |
| ASP | 62 | A | 0.705 | 0.719 | 1 | 0.424 | ASN | 133 | A | 0.743 | 0.741 | 1 | 0.484 |
| LEU | 63 | A | 0.724 | 0.718 | 1 | 0.442 | **ARG** | **134** | **A** | **0.84** | **0.764** | **0.798** | **0.806** |
| VAL | 64 | A | 0.73 | 0.726 | 1 | 0.456 | THR | 135 | A | 0.618 | 0.65 | 0.998 | 0.27 |
| **ASP** | **65** | **A** | **0.824** | **0.811** | **0.676** | **0.959** | ASN | 136 | A | 0.725 | 0.735 | 1 | 0.46 |
| GLU | 66 | A | 0.844 | 0.839 | 1 | 0.683 | **GLN** | **137** | **A** | **0.801** | **0.753** | **0.724** | **0.83** |
| VAL | 67 | A | 0.704 | 0.718 | 1 | 0.422 | LEU | 138 | A | 0.669 | 0.674 | 0.999 | 0.344 |
| LEU | 68 | A | 0.719 | 0.708 | 1 | 0.427 | LEU | 139 | A | 0.638 | 0.685 | 1 | 0.323 |
| ASN | 69 | A | 0.67 | 0.724 | 1 | 0.394 | VAL | 140 | A | 0.69 | 0.692 | 1 | 0.382 |
| HIS | 70 | A | 0.814 | 0.838 | 1 | 0.652 | GLN | 141 | A | 0.802 | 0.8 | 0.998 | 0.604 |
| VAL | 71 | A | 0.731 | 0.725 | 1 | 0.456 | ASN | 142 | A | 0.773 | 0.809 | 0.967 | 0.615 |
| VAL | 72 | A | 0.719 | 0.72 | 0.999 | 0.44 | MET | 143 | A | 0.725 | 0.741 | 0.9 | 0.566 |
| ARG | 73 | A | 0.852 | 0.839 | 0.984 | 0.707 | PHE | 144 | A | 0.85 | 0.837 | 0.985 | 0.702 |
| GLN | 74 | A | 0.806 | 0.793 | 0.999 | 0.6 | ILE | 145 | A | 0.785 | 0.785 | 0.998 | 0.572 |
| PHE | 75 | A | 0.809 | 0.818 | 1 | 0.627 | LYS | 146 | A | 0.799 | 0.788 | 0.995 | 0.592 |
| SER | 76 | A | 0.726 | 0.701 | 0.999 | 0.428 | ALA | 147 | A | 0.601 | 0.637 | 0.998 | 0.24 |
| ASN | 77 | A | 0.782 | 0.792 | 0.989 | 0.585 | ILE | 148 | A | 0.779 | 0.743 | 0.992 | 0.53 |
| PHE | 78 | A | 0.819 | 0.789 | 0.997 | 0.611 | GLU | 149 | A | 0.815 | 0.8 | 0.993 | 0.622 |
| LEU | 79 | A | 0.718 | 0.679 | 0.999 | 0.398 | ARG | 150 | A | 0.758 | 0.787 | 0.897 | 0.648 |
| SER | 80 | A | 0.692 | 0.706 | 0.999 | 0.399 | GLY | 151 | A | 0.636 | 0.637 | 1 | 0.273 |

**Table S5: 7AMN (P), 7AMT (Q),** continued..

| Residue | | | S (P) | S (Q) | J (P,Q) | PRI |
|---|---|---|---|---|---|---|
| **GLU** | **152** | **A** | **0.798** | **0.809** | **0.711** | **0.896** |
| VAL | 153 | A | 0.688 | 0.645 | 0.996 | 0.337 |
| CYS | 154 | A | 0.683 | 0.651 | 0.967 | 0.367 |
| ASP | 155 | A | 0.638 | 0.702 | 0.94 | 0.4 |
| GLN | 156 | A | 0.681 | 0.621 | 0.926 | 0.376 |
| HIS | 157 | A | 0.794 | 0.791 | 0.88 | 0.705 |
| ASP | 158 | A | 0.713 | 0.658 | 0.949 | 0.422 |
| SER | 159 | A | 0.712 | 0.71 | 1 | 0.422 |
| GLU | 160 | A | 0.794 | 0.757 | 0.903 | 0.648 |
| HIS | 161 | A | 0.778 | 0.766 | 0.999 | 0.545 |
| LEU | 162 | A | 0.759 | 0.772 | 0.997 | 0.534 |
| ALA | 163 | A | 0.587 | 0.588 | 0.999 | 0.176 |
| **ASN** | **164** | **A** | **0.808** | **0.805** | **0.707** | **0.906** |
| LEU | 165 | A | 0.753 | 0.736 | 0.991 | 0.498 |
| PHE | 166 | A | 0.819 | 0.821 | 1 | 0.64 |
| HIS | 167 | A | 0.783 | 0.759 | 1 | 0.542 |
| GLY | 168 | A | 0.622 | 0.623 | 1 | 0.245 |
| ILE | 169 | A | 0.757 | 0.765 | 0.999 | 0.523 |
| CYS | 170 | A | 0.669 | 0.664 | 1 | 0.333 |
| TYR | 171 | A | 0.833 | 0.864 | 0.998 | 0.699 |
| SER | 172 | A | 0.679 | 0.676 | 0.999 | 0.356 |
| LEU | 173 | A | 0.726 | 0.714 | 0.973 | 0.467 |
| PHE | 174 | A | 0.857 | 0.852 | 1 | 0.709 |
| VAL | 175 | A | 0.701 | 0.702 | 0.998 | 0.405 |
| GLN | 176 | A | 0.718 | 0.782 | 0.857 | 0.643 |
| ALA | 177 | A | 0.491 | 0.468 | 0.999 | -0.04 |
| ASN | 178 | A | 0.703 | 0.736 | 0.999 | 0.44 |
| ARG | 179 | A | 0.822 | 0.749 | 0.776 | 0.795 |
| ALA | 180 | A | 0.629 | 0.659 | 0.976 | 0.312 |
| ALA | 184 | A | 0.818 | 0.77 | 0.992 | 0.596 |
| GLU | 185 | A | 0.67 | 0.667 | 0.713 | 0.624 |
| LEU | 186 | A | 0.72 | 0.715 | 0.958 | 0.477 |
| LYS | 187 | A | 0.655 | 0.655 | 0.879 | 0.431 |
| HIS | 188 | A | 0.777 | 0.757 | 0.863 | 0.671 |
| LEU | 189 | A | 0.746 | 0.734 | 0.765 | 0.715 |
| VAL | 190 | A | 0.735 | 0.724 | 0.998 | 0.461 |
| ASN | 191 | A | 0.633 | 0.756 | 0.714 | 0.675 |
| SER | 192 | A | 0.65 | 0.641 | 0.774 | 0.517 |
| TYR | 193 | A | 0.833 | 0.858 | 0.999 | 0.692 |
| LEU | 194 | A | 0.708 | 0.661 | 0.596 | 0.773 |
| ASP | 195 | A | 0.678 | 0.724 | 0.98 | 0.422 |
| MET | 196 | A | 0.711 | 0.718 | 0.909 | 0.52 |
| LEU | 197 | A | 0.725 | 0.744 | 0.999 | 0.47 |
| CYS | 198 | A | 0.683 | 0.787 | 0.987 | 0.483 |
| LEU | 12 | B | 0.749 | 0.74 | 0.991 | 0.498 |
| SER | 13 | B | 0.719 | 0.701 | 0.97 | 0.45 |
| PRO | 14 | B | 0.743 | 0.743 | 0.997 | 0.489 |
| ILE | 15 | B | 0.744 | 0.82 | 1 | 0.564 |
| LYS | 16 | B | 0.828 | 0.755 | 0.948 | 0.635 |
| ARG | 17 | B | 0.838 | 0.839 | 0.999 | 0.678 |
| LYS | 18 | B | 0.745 | 0.65 | 0.997 | 0.398 |
| GLN | 19 | B | 0.823 | 0.839 | 0.995 | 0.667 |
| GLN | 20 | B | 0.819 | 0.799 | 0.989 | 0.629 |
| LEU | 21 | B | 0.701 | 0.737 | 1 | 0.438 |
| MET | 22 | B | 0.744 | 0.733 | 0.998 | 0.479 |
| **GLU** | **23** | **B** | **0.677** | **0.754** | **0.616** | **0.815** |
| ILE | 24 | B | 0.762 | 0.763 | 0.994 | 0.531 |
| ALA | 25 | B | 0.567 | 0.562 | 1 | 0.129 |
| LEU | 26 | B | 0.759 | 0.742 | 0.999 | 0.502 |
| GLU | 27 | B | 0.806 | 0.808 | 0.861 | 0.753 |
| VAL | 28 | B | 0.677 | 0.701 | 1 | 0.378 |
| PHE | 29 | B | 0.842 | 0.844 | 0.994 | 0.692 |
| ALA | 30 | B | 0.491 | 0.51 | 1 | 0.001 |
| ARG | 31 | B | 0.793 | 0.781 | 0.99 | 0.584 |
| ARG | 32 | B | 0.793 | 0.8 | 0.996 | 0.597 |
| GLY | 33 | B | 0.618 | 0.619 | 1 | 0.237 |
| ILE | 34 | B | 0.732 | 0.718 | 0.999 | 0.451 |
| GLY | 35 | B | 0.624 | 0.626 | 0.999 | 0.251 |
| **ARG** | **36** | **B** | **0.823** | **0.814** | **0.772** | **0.865** |
| GLY | 37 | B | 0.614 | 0.613 | 1 | 0.227 |
| GLY | 38 | B | 0.624 | 0.625 | 1 | 0.249 |

| Residue | | | S (P) | S (Q) | J (P,Q) | PRI |
|---|---|---|---|---|---|---|
| HIS | 39 | B | 0.819 | 0.827 | 1 | 0.646 |
| ALA | 40 | B | 0.65 | 0.656 | 1 | 0.306 |
| ASP | 41 | B | 0.749 | 0.745 | 0.996 | 0.498 |
| ILE | 42 | B | 0.74 | 0.749 | 0.999 | 0.49 |
| ALA | 43 | B | 0.562 | 0.565 | 1 | 0.127 |
| GLU | 44 | B | 0.644 | 0.66 | 0.977 | 0.327 |
| ILE | 45 | B | 0.754 | 0.763 | 0.998 | 0.519 |
| ALA | 46 | B | 0.493 | 0.511 | 0.998 | 0.006 |
| GLN | 47 | B | 0.66 | 0.661 | 0.965 | 0.356 |
| VAL | 48 | B | 0.682 | 0.662 | 0.999 | 0.345 |
| SER | 49 | B | 0.746 | 0.73 | 1 | 0.476 |
| VAL | 50 | B | 0.678 | 0.706 | 1 | 0.384 |
| ALA | 51 | B | 0.608 | 0.622 | 1 | 0.23 |
| THR | 52 | B | 0.695 | 0.716 | 0.997 | 0.414 |
| VAL | 53 | B | 0.684 | 0.686 | 1 | 0.37 |
| **PHE** | **54** | **B** | **0.862** | **0.864** | **0.852** | **0.874** |
| ASN | 55 | B | 0.711 | 0.733 | 0.994 | 0.45 |
| **TYR** | **56** | **B** | **0.854** | **0.853** | **0.87** | **0.837** |
| **PHE** | **57** | **B** | **0.806** | **0.828** | **0.808** | **0.826** |
| PRO | 58 | B | 0.75 | 0.743 | 1 | 0.493 |
| THR | 59 | B | 0.745 | 0.791 | 1 | 0.536 |
| ARG | 60 | B | 0.828 | 0.841 | 0.999 | 0.67 |
| GLU | 61 | B | 0.641 | 0.642 | 0.979 | 0.304 |
| ASP | 62 | B | 0.671 | 0.67 | 0.978 | 0.363 |
| LEU | 63 | B | 0.726 | 0.722 | 0.999 | 0.449 |
| VAL | 64 | B | 0.709 | 0.732 | 0.999 | 0.442 |
| ASP | 65 | B | 0.673 | 0.708 | 0.824 | 0.557 |
| GLU | 66 | B | 0.783 | 0.713 | 0.826 | 0.67 |
| VAL | 67 | B | 0.713 | 0.724 | 0.999 | 0.438 |
| LEU | 68 | B | 0.75 | 0.717 | 0.999 | 0.468 |
| ASN | 69 | B | 0.721 | 0.737 | 0.997 | 0.461 |
| HIS | 70 | B | 0.843 | 0.837 | 0.999 | 0.681 |
| VAL | 71 | B | 0.729 | 0.727 | 1 | 0.456 |
| VAL | 72 | B | 0.662 | 0.706 | 1 | 0.368 |
| ARG | 73 | B | 0.765 | 0.733 | 0.785 | 0.713 |
| GLN | 74 | B | 0.781 | 0.76 | 0.947 | 0.594 |
| PHE | 75 | B | 0.791 | 0.807 | 0.995 | 0.603 |
| SER | 76 | B | 0.73 | 0.68 | 0.998 | 0.412 |
| ASN | 77 | B | 0.794 | 0.791 | 1 | 0.585 |
| **PHE** | **78** | **B** | **0.815** | **0.815** | **0.811** | **0.819** |
| LEU | 79 | B | 0.669 | 0.727 | 0.859 | 0.537 |
| SER | 80 | B | 0.721 | 0.65 | 0.924 | 0.447 |
| ASP | 81 | B | 0.778 | 0.722 | 0.999 | 0.501 |
| ASN | 82 | B | 0.732 | 0.716 | 0.988 | 0.46 |
| ILE | 83 | B | 0.729 | 0.746 | 0.999 | 0.476 |
| ASP | 84 | B | 0.625 | 0.609 | 0.977 | 0.257 |
| LEU | 85 | B | 0.67 | 0.576 | 0.758 | 0.488 |
| ASP | 86 | B | 0.655 | 0.726 | 0.791 | 0.59 |
| LEU | 87 | B | 0.764 | 0.684 | 0.999 | 0.449 |
| HIS | 88 | B | 0.821 | 0.769 | 0.881 | 0.709 |
| ALA | 89 | B | 0.681 | 0.658 | 0.999 | 0.34 |
| ARG | 90 | B | 0.683 | 0.729 | 0.829 | 0.583 |
| GLU | 91 | B | 0.682 | 0.708 | 0.791 | 0.599 |
| ASN | 92 | B | 0.732 | 0.782 | 0.874 | 0.64 |
| **ILE** | **93** | **B** | **0.765** | **0.772** | **0.602** | **0.935** |
| ALA | 94 | B | 0.605 | 0.612 | 1 | 0.217 |
| ASN | 95 | B | 0.605 | 0.584 | 0.991 | 0.198 |
| ILE | 96 | B | 0.773 | 0.759 | 1 | 0.532 |
| THR | 97 | B | 0.74 | 0.762 | 0.809 | 0.693 |
| ASN | 98 | B | 0.692 | 0.791 | 0.988 | 0.495 |
| ALA | 99 | B | 0.568 | 0.61 | 1 | 0.178 |
| MET | 100 | B | 0.692 | 0.704 | 0.968 | 0.428 |
| ILE | 101 | B | 0.8 | 0.757 | 1 | 0.557 |
| **GLU** | **102** | **B** | **0.692** | **0.812** | **0.689** | **0.815** |
| LEU | 103 | B | 0.767 | 0.77 | 0.995 | 0.542 |
| VAL | 104 | B | 0.703 | 0.682 | 0.999 | 0.386 |
| SER | 105 | B | 0.649 | 0.655 | 0.999 | 0.305 |
| **GLN** | **106** | **B** | **0.761** | **0.78** | **0.73** | **0.811** |
| ASP | 107 | B | 0.663 | 0.68 | 0.956 | 0.387 |
| CYS | 108 | B | 0.659 | 0.658 | 1 | 0.317 |
| HIS | 109 | B | 0.827 | 0.835 | 1 | 0.662 |

**Table S5: 7AMN (P), 7AMT (Q),** continued..

| Residue | | | S (P) | S (Q) | J (P,Q) | PRI |
|---|---|---|---|---|---|---|
| TRP | 110 | B | 0.888 | 0.888 | 1 | 0.776 |
| LEU | 111 | B | 0.721 | 0.729 | 1 | 0.45 |
| LYS | 112 | B | 0.813 | 0.81 | 0.952 | 0.671 |
| VAL | 113 | B | 0.693 | 0.689 | 1 | 0.382 |
| TRP | 114 | B | 0.871 | 0.869 | 1 | 0.74 |
| PHE | 115 | B | 0.839 | 0.833 | 1 | 0.672 |
| GLU | 116 | B | 0.789 | 0.792 | 1 | 0.581 |
| TRP | 117 | B | 0.872 | 0.862 | 0.999 | 0.735 |
| SER | 118 | B | 0.63 | 0.605 | 1 | 0.235 |
| ALA | 119 | B | 0.478 | 0.479 | 1 | -0.043 |
| SER | 120 | B | 0.56 | 0.538 | 1 | 0.098 |
| THR | 121 | B | 0.667 | 0.681 | 0.999 | 0.349 |
| **ARG** | **122** | **B** | **0.806** | **0.826** | **0.794** | **0.838** |
| ASP | 123 | B | 0.624 | 0.555 | 0.959 | 0.22 |
| GLU | 124 | B | 0.735 | 0.759 | 0.997 | 0.497 |
| VAL | 125 | B | 0.707 | 0.709 | 1 | 0.416 |
| TRP | 126 | B | 0.871 | 0.869 | 1 | 0.74 |
| PRO | 127 | B | 0.762 | 0.749 | 1 | 0.511 |
| LEU | 128 | B | 0.731 | 0.733 | 1 | 0.464 |
| PHE | 129 | B | 0.834 | 0.836 | 1 | 0.67 |
| VAL | 130 | B | 0.706 | 0.725 | 0.999 | 0.432 |
| SER | 131 | B | 0.754 | 0.752 | 0.999 | 0.507 |
| THR | 132 | B | 0.702 | 0.709 | 0.75 | 0.661 |
| ASN | 133 | B | 0.777 | 0.753 | 0.999 | 0.531 |
| ARG | 134 | B | 0.816 | 0.829 | 0.862 | 0.783 |
| THR | 135 | B | 0.687 | 0.686 | 0.977 | 0.396 |
| ASN | 136 | B | 0.799 | 0.754 | 0.984 | 0.569 |
| GLN | 137 | B | 0.76 | 0.759 | 0.848 | 0.671 |
| LEU | 138 | B | 0.767 | 0.744 | 0.78 | 0.731 |
| LEU | 139 | B | 0.66 | 0.702 | 0.829 | 0.533 |
| VAL | 140 | B | 0.681 | 0.66 | 1 | 0.341 |
| GLN | 141 | B | 0.828 | 0.76 | 0.987 | 0.601 |
| ASN | 142 | B | 0.791 | 0.805 | 0.986 | 0.61 |
| **MET** | **143** | **B** | **0.768** | **0.711** | **0.618** | **0.861** |
| **PHE** | **144** | **B** | **0.844** | **0.831** | **0.766** | **0.909** |
| ILE | 145 | B | 0.781 | 0.764 | 0.999 | 0.546 |
| LYS | 146 | B | 0.712 | 0.731 | 0.755 | 0.688 |
| ALA | 147 | B | 0.608 | 0.629 | 0.99 | 0.247 |
| ILE | 148 | B | 0.781 | 0.671 | 0.738 | 0.714 |
| GLU | 149 | B | 0.688 | 0.722 | 0.931 | 0.479 |
| ARG | 150 | B | 0.833 | 0.729 | 0.856 | 0.706 |
| GLY | 151 | B | 0.63 | 0.644 | 0.978 | 0.296 |
| GLU | 152 | B | 0.739 | 0.799 | 0.875 | 0.663 |
| VAL | 153 | B | 0.735 | 0.747 | 0.725 | 0.757 |
| CYS | 154 | B | 0.789 | 0.722 | 0.879 | 0.632 |
| HIS | 157 | B | 0.843 | 0.846 | 0.986 | 0.703 |
| ASP | 158 | B | 0.729 | 0.773 | 0.865 | 0.637 |
| SER | 159 | B | 0.693 | 0.703 | 0.994 | 0.402 |
| GLU | 160 | B | 0.791 | 0.743 | 0.775 | 0.759 |
| HIS | 161 | B | 0.766 | 0.786 | 0.992 | 0.56 |
| LEU | 162 | B | 0.777 | 0.785 | 0.998 | 0.564 |
| ALA | 163 | B | 0.592 | 0.592 | 0.999 | 0.185 |
| ASN | 164 | B | 0.769 | 0.824 | 0.997 | 0.596 |
| LEU | 165 | B | 0.651 | 0.666 | 0.958 | 0.359 |
| PHE | 166 | B | 0.841 | 0.842 | 1 | 0.683 |
| HIS | 167 | B | 0.784 | 0.765 | 0.999 | 0.55 |
| GLY | 168 | B | 0.62 | 0.62 | 1 | 0.24 |
| ILE | 169 | B | 0.775 | 0.736 | 0.999 | 0.512 |
| CYS | 170 | B | 0.67 | 0.68 | 1 | 0.35 |
| TYR | 171 | B | 0.843 | 0.862 | 0.999 | 0.706 |
| SER | 172 | B | 0.695 | 0.686 | 1 | 0.381 |
| LEU | 173 | B | 0.708 | 0.699 | 0.838 | 0.569 |
| PHE | 174 | B | 0.85 | 0.858 | 1 | 0.708 |
| VAL | 175 | B | 0.712 | 0.717 | 0.767 | 0.662 |
| GLN | 176 | B | 0.8 | 0.785 | 0.999 | 0.586 |
| ALA | 177 | B | 0.521 | 0.517 | 0.999 | 0.039 |
| ASN | 178 | B | 0.725 | 0.696 | 0.999 | 0.422 |
| ARG | 179 | B | 0.806 | 0.715 | 0.984 | 0.537 |
| ALA | 180 | B | 0.782 | 0.683 | 0.994 | 0.471 |
| ALA | 184 | B | 0.813 | 0.823 | 0.999 | 0.637 |
| GLU | 185 | B | 0.764 | 0.64 | 0.77 | 0.634 |
| LEU | 186 | B | 0.724 | 0.704 | 0.827 | 0.601 |
| LYS | 187 | B | 0.669 | 0.597 | 0.885 | 0.381 |
| HIS | 188 | B | 0.715 | 0.705 | 0.759 | 0.661 |
| LEU | 189 | B | 0.695 | 0.72 | 0.712 | 0.703 |
| VAL | 190 | B | 0.747 | 0.763 | 0.874 | 0.636 |
| ASN | 191 | B | 0.751 | 0.692 | 0.951 | 0.492 |
| SER | 192 | B | 0.684 | 0.692 | 0.991 | 0.385 |
| TYR | 193 | B | 0.823 | 0.813 | 0.997 | 0.639 |
| LEU | 194 | B | 0.748 | 0.737 | 0.696 | 0.789 |
| ASP | 195 | B | 0.776 | 0.794 | 0.955 | 0.615 |
| MET | 196 | B | 0.774 | 0.766 | 0.985 | 0.555 |
| LEU | 197 | B | 0.74 | 0.724 | 0.769 | 0.695 |
| CYS | 198 | B | 0.568 | 0.619 | 0.922 | 0.265 |
| **ILE** | **199** | **B** | **0.832** | **0.867** | **0.7** | **0.999** |

**Table S6: 2X6U (P), 2X6V (Q)**

| Residue | | | S (P) | S (Q) | J | PRI |
|---|---|---|---|---|---|---|
| GLY | 53 | A | 0.435 | 0.828 | 0.841 | 0.422 |
| ILE | 54 | A | 0.62 | 0.633 | 0.989 | 0.264 |
| LYS | 55 | A | 0.79 | 0.751 | 0.856 | 0.685 |
| VAL | 56 | A | 0.7 | 0.711 | 0.997 | 0.414 |
| **PHE** | **57** | **A** | **0.823** | **0.838** | **0.765** | **0.896** |
| LEU | 58 | A | 0.736 | 0.733 | 0.999 | 0.47 |
| HIS | 59 | A | 0.856 | 0.855 | 0.994 | 0.717 |
| GLU | 60 | A | 0.745 | 0.722 | 0.972 | 0.495 |
| ARG | 61 | A | 0.859 | 0.827 | 0.982 | 0.704 |
| AGLU | 62 | A | 0.744 | 0.628 | 0.651 | 0.721 |
| LEU | 63 | A | 0.737 | 0.722 | 0.992 | 0.467 |
| TRP | 64 | A | 0.889 | 0.887 | 0.999 | 0.777 |
| LEU | 65 | A | 0.773 | 0.772 | 0.999 | 0.546 |
| LYS | 66 | A | 0.684 | 0.681 | 0.95 | 0.415 |
| PHE | 67 | A | 0.805 | 0.815 | 0.996 | 0.624 |
| HIS | 68 | A | 0.81 | 0.811 | 0.998 | 0.623 |
| GLU | 69 | A | 0.775 | 0.758 | 0.988 | 0.545 |
| VAL | 70 | A | 0.682 | 0.692 | 0.999 | 0.375 |
| GLY | 71 | A | 0.637 | 0.636 | 0.998 | 0.275 |
| THR | 72 | A | 0.723 | 0.728 | 1 | 0.451 |
| GLU | 73 | A | 0.762 | 0.769 | 0.997 | 0.534 |
| MET | 74 | A | 0.775 | 0.772 | 1 | 0.547 |
| ILE | 75 | A | 0.765 | 0.771 | 0.995 | 0.541 |
| ILE | 76 | A | 0.757 | 0.748 | 0.998 | 0.507 |
| THR | 77 | A | 0.746 | 0.771 | 0.998 | 0.519 |
| LYS | 78 | A | 0.731 | 0.746 | 0.998 | 0.479 |
| ALA | 79 | A | 0.548 | 0.576 | 0.997 | 0.127 |
| GLY | 80 | A | 0.616 | 0.611 | 0.99 | 0.237 |
| **ARG** | **81** | **A** | **0.889** | **0.883** | **0.973** | **0.799** |
| ARG | 82 | A | 0.857 | 0.846 | 0.915 | 0.788 |
| MET | 83 | A | 0.697 | 0.691 | 0.998 | 0.39 |
| PHE | 84 | A | 0.858 | 0.854 | 0.97 | 0.742 |
| PRO | 85 | A | 0.629 | 0.631 | 1 | 0.26 |
| SER | 86 | A | 0.634 | 0.594 | 0.993 | 0.235 |
| TYR | 87 | A | 0.851 | 0.83 | 0.996 | 0.685 |
| LYS | 88 | A | 0.76 | 0.804 | 0.983 | 0.581 |
| VAL | 89 | A | 0.648 | 0.645 | 0.999 | 0.294 |
| LYS | 90 | A | 0.75 | 0.733 | 0.997 | 0.486 |
| AVAL | 91 | A | 0.755 | 0.752 | 0.999 | 0.508 |
| THR | 92 | A | 0.745 | 0.73 | 0.999 | 0.476 |
| GLY | 93 | A | 0.645 | 0.645 | 1 | 0.29 |
| LEU | 94 | A | 0.633 | 0.654 | 0.997 | 0.29 |
| ASN | 95 | A | 0.693 | 0.709 | 0.998 | 0.404 |
| PRO | 96 | A | 0.719 | 0.714 | 0.992 | 0.441 |
| LYS | 97 | A | 0.633 | 0.67 | 0.824 | 0.479 |
| THR | 98 | A | 0.704 | 0.723 | 0.998 | 0.429 |
| LYS | 99 | A | 0.75 | 0.734 | 0.961 | 0.523 |
| **TYR** | **100** | **A** | **0.854** | **0.855** | **0.871** | **0.838** |
| ILE | 101 | A | 0.668 | 0.709 | 0.999 | 0.378 |
| LEU | 102 | A | 0.717 | 0.727 | 0.999 | 0.445 |
| ALEU | 103 | A | 0.663 | 0.664 | 0.999 | 0.328 |
| MET | 104 | A | 0.751 | 0.722 | 0.988 | 0.485 |
| ASP | 105 | A | 0.768 | 0.795 | 0.989 | 0.574 |
| ILE | 106 | A | 0.744 | 0.742 | 0.998 | 0.488 |
| VAL | 107 | A | 0.712 | 0.726 | 0.999 | 0.439 |
| PRO | 108 | A | 0.651 | 0.653 | 0.999 | 0.305 |
| ALA | 109 | A | 0.434 | 0.421 | 0.997 | -0.142 |
| ASP | 110 | A | 0.749 | 0.732 | 0.999 | 0.482 |
| AASP | 111 | A | 0.74 | 0.675 | 0.862 | 0.553 |
| HIS | 112 | A | 0.815 | 0.819 | 0.978 | 0.656 |
| ARG | 113 | A | 0.857 | 0.867 | 0.958 | 0.766 |
| TYR | 114 | A | 0.821 | 0.836 | 0.991 | 0.666 |
| **LYS** | **115** | **A** | **0.802** | **0.717** | **0.725** | **0.794** |
| PHE | 116 | A | 0.849 | 0.846 | 0.997 | 0.698 |
| ALA | 117 | A | 0.65 | 0.619 | 0.998 | 0.271 |
| ASP | 118 | A | 0.664 | 0.732 | 0.97 | 0.426 |
| ASN | 119 | A | 0.643 | 0.699 | 0.666 | 0.676 |
| LYS | 120 | A | 0.756 | 0.793 | 0.848 | 0.701 |
| TRP | 121 | A | 0.884 | 0.886 | 0.997 | 0.773 |
| SER | 122 | A | 0.62 | 0.69 | 0.944 | 0.366 |
| VAL | 123 | A | 0.675 | 0.702 | 0.997 | 0.38 |
| THR | 124 | A | 0.675 | 0.667 | 0.736 | 0.606 |
| GLY | 125 | A | 0.625 | 0.626 | 1 | 0.251 |
| LYS | 126 | A | 0.724 | 0.801 | 0.802 | 0.723 |
| ALA | 127 | A | 0.513 | 0.493 | 1 | 0.006 |
| **GLU** | **128** | **A** | **0.775** | **0.771** | **0.717** | **0.829** |
| PRO | 129 | A | 0.714 | 0.716 | 0.979 | 0.451 |
| ALA | 130 | A | 0.766 | 0.732 | 0.939 | 0.559 |
| ARG | 134 | A | 0.826 | 0.829 | 0.996 | 0.659 |
| LEU | 135 | A | 0.601 | 0.554 | 0.998 | 0.157 |
| TYR | 136 | A | 0.858 | 0.843 | 0.995 | 0.706 |
| VAL | 137 | A | 0.676 | 0.687 | 0.999 | 0.364 |
| HIS | 138 | A | 0.839 | 0.838 | 0.997 | 0.68 |
| PRO | 139 | A | 0.649 | 0.646 | 1 | 0.295 |
| ASP | 140 | A | 0.751 | 0.753 | 0.999 | 0.505 |
| SER | 141 | A | 0.632 | 0.645 | 0.993 | 0.284 |
| PRO | 142 | A | 0.668 | 0.684 | 0.997 | 0.355 |
| ALA | 143 | A | 0.569 | 0.568 | 0.999 | 0.138 |
| THR | 144 | A | 0.672 | 0.668 | 0.999 | 0.341 |
| GLY | 145 | A | 0.621 | 0.62 | 1 | 0.241 |
| ALA | 146 | A | 0.632 | 0.621 | 0.999 | 0.254 |
| HIS | 147 | A | 0.759 | 0.756 | 0.999 | 0.516 |
| TRP | 148 | A | 0.874 | 0.873 | 0.999 | 0.748 |
| MET | 149 | A | 0.768 | 0.763 | 0.999 | 0.532 |
| **ARG** | **150** | **A** | **0.687** | **0.815** | **0.71** | **0.792** |
| GLN | 151 | A | 0.806 | 0.78 | 0.996 | 0.59 |
| LEU | 152 | A | 0.666 | 0.658 | 0.998 | 0.326 |
| VAL | 153 | A | 0.684 | 0.681 | 1 | 0.365 |
| SER | 154 | A | 0.658 | 0.651 | 0.999 | 0.31 |
| PHE | 155 | A | 0.848 | 0.845 | 0.999 | 0.694 |
| **GLN** | **156** | **A** | **0.822** | **0.777** | **0.659** | **0.94** |
| **ALYS** | **157** | **A** | **0.721** | **0.785** | **0.708** | **0.798** |
| LEU | 158 | A | 0.744 | 0.759 | 0.997 | 0.506 |
| LYS | 159 | A | 0.789 | 0.778 | 0.982 | 0.585 |
| LEU | 160 | A | 0.753 | 0.739 | 0.997 | 0.495 |
| THR | 161 | A | 0.692 | 0.704 | 0.999 | 0.397 |
| ASN | 162 | A | 0.657 | 0.693 | 0.741 | 0.609 |
| ASN | 163 | A | 0.747 | 0.756 | 0.999 | 0.504 |
| HIS | 164 | A | 0.71 | 0.728 | 0.999 | 0.439 |
| LEU | 165 | A | 0.705 | 0.659 | 0.999 | 0.365 |
| ASP | 166 | A | 0.774 | 0.766 | 0.999 | 0.541 |
| PRO | 167 | A | 0.69 | 0.7 | 1 | 0.39 |
| PHE | 168 | A | 0.829 | 0.801 | 0.999 | 0.631 |
| GLY | 169 | A | 0.647 | 0.644 | 1 | 0.291 |
| **HIS** | **170** | **A** | **0.816** | **0.812** | **0.786** | **0.842** |
| ILE | 171 | A | 0.617 | 0.611 | 1 | 0.228 |
| ILE | 172 | A | 0.752 | 0.763 | 0.998 | 0.517 |
| LEU | 173 | A | 0.719 | 0.717 | 0.999 | 0.437 |
| ASN | 174 | A | 0.681 | 0.713 | 0.997 | 0.397 |
| SER | 175 | A | 0.689 | 0.688 | 0.999 | 0.378 |
| MET | 176 | A | 0.751 | 0.743 | 0.992 | 0.502 |
| HIS | 177 | A | 0.799 | 0.803 | 0.837 | 0.765 |
| LYS | 178 | A | 0.701 | 0.672 | 0.998 | 0.375 |
| TYR | 179 | A | 0.861 | 0.855 | 0.999 | 0.717 |
| GLN | 180 | A | 0.81 | 0.81 | 1 | 0.62 |
| PRO | 181 | A | 0.678 | 0.674 | 0.999 | 0.353 |
| **ARG** | **182** | **A** | **0.876** | **0.877** | **0.952** | **0.801** |
| LEU | 183 | A | 0.758 | 0.734 | 0.998 | 0.494 |
| HIS | 184 | A | 0.814 | 0.777 | 0.983 | 0.608 |
| ILE | 185 | A | 0.72 | 0.717 | 0.999 | 0.438 |
| VAL | 186 | A | 0.71 | 0.717 | 0.997 | **0.43** |
| LYS | 187 | A | 0.775 | 0.788 | 0.94 | 0.623 |
| ALA | 188 | A | 0.615 | 0.612 | 0.985 | 0.242 |
| ASP | 189 | A | 0.798 | 0.75 | 0.908 | 0.64 |
| THR | 199 | A | 0.705 | 0.805 | 0.942 | 0.568 |
| ALA | 200 | A | 0.591 | 0.605 | 0.978 | 0.218 |
| PHE | 201 | A | 0.744 | 0.783 | 0.96 | 0.567 |
| CYS | 202 | A | 0.686 | 0.532 | 0.435 | 0.783 |
| THR | 203 | A | 0.726 | 0.68 | 0.996 | 0.41 |
| HIS | 204 | A | 0.842 | 0.836 | 0.957 | 0.721 |
| VAL | 205 | A | 0.704 | 0.696 | 0.998 | 0.402 |
| PHE | 206 | A | 0.845 | 0.846 | 0.988 | 0.703 |
| PRO | 207 | A | 0.72 | 0.717 | 0.989 | 0.448 |
| GLU | 208 | A | 0.767 | 0.758 | 0.999 | 0.526 |
| THR | 209 | A | 0.643 | 0.632 | 0.998 | 0.277 |
| ALA | 210 | A | 0.556 | 0.549 | 0.999 | 0.106 |
| PHE | 211 | A | 0.831 | 0.83 | 1 | 0.661 |
| ILE | 212 | A | 0.753 | 0.742 | 0.997 | 0.498 |
| ALA | 213 | A | 0.453 | 0.46 | 0.998 | -0.085 |
| VAL | 214 | A | 0.701 | 0.714 | 1 | 0.415 |
| THR | 215 | A | 0.684 | 0.671 | 0.997 | 0.358 |
| SER | 216 | A | 0.648 | 0.656 | 0.998 | 0.306 |
| TYR | 217 | A | 0.846 | 0.841 | 1 | 0.687 |
| GLN | 218 | A | 0.747 | 0.707 | 0.999 | 0.455 |
| ASN | 219 | A | 0.735 | 0.761 | 0.984 | 0.512 |
| **HIS** | **220** | **A** | **0.791** | **0.847** | **0.827** | **0.811** |
| LYS | 221 | A | 0.645 | 0.691 | 0.998 | 0.338 |
| ILE | 222 | A | 0.759 | 0.758 | 0.981 | 0.536 |
| THR | 223 | A | 0.708 | 0.711 | 0.995 | 0.424 |
| GLN | 224 | A | 0.765 | 0.748 | 0.833 | 0.68 |
| LEU | 225 | A | 0.734 | 0.742 | 0.841 | 0.635 |
| LYS | 226 | A | 0.825 | 0.837 | 0.996 | 0.666 |
| ILE | 227 | A | 0.795 | 0.796 | 0.998 | 0.593 |
| GLU | 228 | A | 0.8 | 0.784 | 0.992 | 0.592 |
| ASN | 229 | A | 0.742 | 0.742 | 0.995 | 0.489 |
| **ASN** | **230** | **A** | **0.88** | **0.862** | **0.91** | **0.832** |

| Table S7(a): 1LFL (P), 1LFT (Q) | | | | | | | | | | | | | |
|---|---|---|---|---|---|---|---|---|---|---|---|---|---|
| **Residue** | | | **S (P)** | **S (Q)** | **J (P,Q)** | **PRI** | **Residue** | | | **S (P)** | **S (Q)** | **J (P,Q)** | **PRI** |
| VAL | 1 | A | 0.691 | 0.75 | 0.859 | 0.582 | ASP | 74 | A | 0.733 | 0.697 | 0.969 | 0.461 |
| LEU | 2 | A | 0.644 | 0.685 | 0.992 | 0.337 | ASP | 75 | A | 0.708 | 0.776 | 0.958 | 0.526 |
| SER | 3 | A | 0.728 | 0.736 | 1 | 0.464 | MET | 76 | A | 0.726 | 0.696 | 0.999 | 0.423 |
| PRO | 4 | A | 0.748 | 0.76 | 0.999 | 0.509 | PRO | 77 | A | 0.69 | 0.744 | 0.998 | 0.436 |
| ALA | 5 | A | 0.658 | 0.648 | 1 | 0.306 | ASN | 78 | A | 0.692 | 0.744 | 0.897 | 0.539 |
| ASP | 6 | A | 0.765 | 0.791 | 0.942 | 0.614 | ALA | 79 | A | 0.458 | 0.508 | 0.997 | -0.031 |
| LYS | 7 | A | 0.85 | 0.73 | 0.872 | 0.708 | LEU | 80 | A | 0.735 | 0.713 | 1 | 0.448 |
| THR | 8 | A | 0.727 | 0.728 | 1 | 0.455 | SER | 81 | A | 0.736 | 0.755 | 0.999 | 0.492 |
| ASN | 9 | A | 0.706 | 0.767 | 0.951 | 0.522 | ALA | 82 | A | 0.679 | 0.686 | 1 | 0.365 |
| VAL | 10 | A | 0.707 | 0.689 | 0.999 | 0.397 | LEU | 83 | A | 0.677 | 0.696 | 0.999 | 0.374 |
| LYS | 11 | A | 0.809 | 0.773 | 0.925 | 0.657 | SER | 84 | A | 0.601 | 0.733 | 0.891 | 0.443 |
| ALA | 12 | A | 0.64 | 0.634 | 0.996 | 0.278 | ASP | 85 | A | 0.824 | 0.735 | 0.978 | 0.581 |
| ALA | 13 | A | 0.546 | 0.544 | 1 | 0.09 | LEU | 86 | A | 0.759 | 0.729 | 0.877 | 0.611 |
| TRP | 14 | A | 0.858 | 0.843 | 1 | 0.701 | HIS | 87 | A | 0.817 | 0.729 | 0.989 | 0.557 |
| GLY | 15 | A | 0.659 | 0.654 | 0.999 | 0.314 | ALA | 88 | A | 0.54 | 0.521 | 0.989 | 0.072 |
| LYS | 16 | A | 0.816 | 0.793 | 0.912 | 0.697 | HIS | 89 | A | 0.811 | 0.809 | 0.986 | 0.634 |
| VAL | 17 | A | 0.703 | 0.674 | 0.999 | 0.378 | LYS | 90 | A | 0.738 | 0.655 | 0.868 | 0.525 |
| GLY | 18 | A | 0.651 | 0.647 | 0.998 | 0.3 | LEU | 91 | A | 0.696 | 0.582 | 0.996 | 0.282 |
| ALA | 19 | A | 0.513 | 0.542 | 1 | 0.055 | **ARG** | **92** | **A** | **0.865** | **0.831** | **0.965** | **0.731** |
| HIS | 20 | A | 0.787 | 0.77 | 0.999 | 0.558 | VAL | 93 | A | 0.718 | 0.684 | 0.996 | 0.406 |
| ALA | 21 | A | 0.599 | 0.66 | 1 | 0.259 | ASP | 94 | A | 0.77 | 0.695 | 0.92 | 0.545 |
| GLY | 22 | A | 0.634 | 0.633 | 1 | 0.267 | PRO | 95 | A | 0.682 | 0.684 | 0.998 | 0.368 |
| GLU | 23 | A | 0.682 | 0.71 | 0.926 | 0.466 | VAL | 96 | A | 0.69 | 0.597 | 0.995 | 0.292 |
| **TYR** | **24** | **A** | **0.862** | **0.869** | **1** | **0.731** | ASN | 97 | A | 0.773 | 0.742 | 0.861 | 0.654 |
| GLY | 25 | A | 0.627 | 0.632 | 1 | 0.259 | PHE | 98 | A | 0.814 | 0.81 | 0.999 | 0.625 |
| ALA | 26 | A | 0.607 | 0.53 | 1 | 0.137 | LYS | 99 | A | 0.746 | 0.672 | 0.836 | 0.582 |
| GLU | 27 | A | 0.808 | 0.738 | 0.913 | 0.633 | LEU | 100 | A | 0.718 | 0.736 | 0.998 | 0.456 |
| ALA | 28 | A | 0.606 | 0.564 | 0.994 | 0.176 | LEU | 101 | A | 0.712 | 0.676 | 1 | 0.388 |
| LEU | 29 | A | 0.726 | 0.693 | 0.999 | 0.42 | SER | 102 | A | 0.719 | 0.705 | 1 | 0.424 |
| GLU | 30 | A | 0.786 | 0.763 | 1 | 0.549 | HIS | 103 | A | 0.806 | 0.801 | 0.999 | 0.608 |
| **ARG** | **31** | **A** | **0.852** | **0.872** | **1** | **0.724** | CYS | 104 | A | 0.627 | 0.628 | 1 | 0.255 |
| MET | 32 | A | 0.747 | 0.745 | 0.999 | 0.493 | LEU | 105 | A | 0.691 | 0.679 | 1 | 0.37 |
| PHE | 33 | A | 0.852 | 0.833 | 1 | 0.685 | LEU | 106 | A | 0.626 | 0.684 | 0.888 | 0.422 |
| LEU | 34 | A | 0.656 | 0.697 | 0.929 | 0.424 | VAL | 107 | A | 0.735 | 0.716 | 0.995 | 0.456 |
| SER | 35 | A | 0.634 | 0.646 | 1 | 0.28 | THR | 108 | A | 0.746 | 0.696 | 1 | 0.442 |
| PHE | 36 | A | 0.834 | 0.847 | 0.999 | 0.682 | LEU | 109 | A | 0.679 | 0.714 | 0.999 | 0.394 |
| PRO | 37 | A | 0.733 | 0.685 | 0.998 | 0.42 | ALA | 110 | A | 0.492 | 0.502 | 1 | -0.006 |
| THR | 38 | A | 0.653 | 0.638 | 0.913 | 0.378 | ALA | 111 | A | 0.43 | 0.491 | 1 | -0.079 |
| THR | 39 | A | 0.69 | 0.698 | 1 | 0.388 | HIS | 112 | A | 0.812 | 0.812 | 1 | 0.624 |
| LYS | 40 | A | 0.735 | 0.754 | 0.981 | 0.508 | LEU | 113 | A | 0.744 | 0.725 | 0.995 | 0.474 |
| THR | 41 | A | 0.759 | 0.681 | 0.915 | 0.525 | PRO | 114 | A | 0.761 | 0.745 | 0.999 | 0.507 |
| TYR | 42 | A | 0.838 | 0.842 | 1 | 0.68 | ALA | 115 | A | 0.572 | 0.61 | 0.999 | 0.183 |
| PHE | 43 | A | 0.835 | 0.848 | 0.999 | 0.684 | GLU | 116 | A | 0.765 | 0.78 | 0.844 | 0.701 |
| PRO | 44 | A | 0.768 | 0.693 | 0.973 | 0.488 | PHE | 117 | A | 0.798 | 0.835 | 0.999 | 0.634 |
| HIS | 45 | A | 0.812 | 0.809 | 0.997 | 0.624 | THR | 118 | A | 0.753 | 0.694 | 0.999 | 0.448 |
| PHE | 46 | A | 0.814 | 0.84 | 0.999 | 0.655 | PRO | 119 | A | 0.644 | 0.687 | 1 | 0.331 |
| ASP | 47 | A | 0.746 | 0.74 | 0.93 | 0.556 | ALA | 120 | A | 0.627 | 0.636 | 1 | 0.263 |
| LEU | 48 | A | 0.713 | 0.719 | 0.947 | 0.485 | VAL | 121 | A | 0.719 | 0.701 | 0.999 | 0.421 |
| SER | 49 | A | 0.736 | 0.749 | 1 | 0.485 | HIS | 122 | A | 0.788 | 0.775 | 1 | 0.563 |
| HIS | 50 | A | 0.749 | 0.725 | 0.896 | 0.578 | ALA | 123 | A | 0.573 | 0.579 | 1 | 0.152 |
| GLY | 51 | A | 0.652 | 0.656 | 0.977 | 0.331 | SER | 124 | A | 0.693 | 0.718 | 0.999 | 0.412 |
| SER | 52 | A | 0.726 | 0.751 | 0.997 | 0.48 | LEU | 125 | A | 0.75 | 0.722 | 0.853 | 0.619 |
| ALA | 53 | A | 0.642 | 0.626 | 1 | 0.268 | ASP | 126 | A | 0.731 | 0.773 | 0.886 | 0.618 |
| GLN | 54 | A | 0.749 | 0.76 | 0.998 | 0.511 | LYS | 127 | A | 0.753 | 0.814 | 0.922 | 0.645 |
| VAL | 55 | A | 0.646 | 0.662 | 0.999 | 0.309 | PHE | 128 | A | 0.816 | 0.816 | 0.999 | 0.633 |
| LYS | 56 | A | 0.772 | 0.725 | 0.961 | 0.536 | LEU | 129 | A | 0.664 | 0.704 | 1 | 0.368 |
| GLY | 57 | A | 0.64 | 0.647 | 0.992 | 0.295 | ALA | 130 | A | 0.599 | 0.617 | 1 | 0.216 |
| HIS | 58 | A | 0.785 | 0.807 | 1 | 0.592 | SER | 131 | A | 0.668 | 0.704 | 0.922 | 0.45 |
| GLY | 59 | A | 0.627 | 0.623 | 1 | 0.25 | VAL | 132 | A | 0.687 | 0.683 | 0.998 | 0.372 |
| LYS | 60 | A | 0.736 | 0.695 | 0.873 | 0.558 | SER | 133 | A | 0.671 | 0.725 | 0.996 | 0.4 |
| LYS | 61 | A | 0.648 | 0.754 | 0.971 | 0.431 | THR | 134 | A | 0.725 | 0.749 | 1 | 0.474 |
| VAL | 62 | A | 0.712 | 0.696 | 1 | 0.408 | VAL | 135 | A | 0.706 | 0.715 | 0.999 | 0.422 |
| ALA | 63 | A | 0.657 | 0.581 | 1 | 0.238 | LEU | 136 | A | 0.742 | 0.766 | 0.999 | 0.509 |
| ASP | 64 | A | 0.826 | 0.759 | 0.999 | 0.586 | THR | 137 | A | 0.704 | 0.679 | 0.948 | 0.435 |
| ALA | 65 | A | 0.621 | 0.579 | 0.999 | 0.201 | SER | 138 | A | 0.642 | 0.661 | 0.998 | 0.305 |
| LEU | 66 | A | 0.688 | 0.708 | 1 | 0.396 | **LYS** | **139** | **A** | **0.761** | **0.789** | **0.823** | **0.727** |
| THR | 67 | A | 0.731 | 0.704 | 1 | 0.435 | TYR | 140 | A | 0.811 | 0.829 | 0.963 | 0.677 |
| ASN | 68 | A | 0.739 | 0.665 | 0.963 | 0.441 | ARG | 141 | A | 0.669 | 0.832 | 0.924 | 0.577 |
| ALA | 69 | A | 0.561 | 0.571 | 0.999 | 0.133 | **VAL** | **1** | **B** | **0.779** | **0.756** | **0.814** | **0.721** |
| VAL | 70 | A | 0.688 | 0.679 | 1 | 0.367 | HIS | 2 | B | 0.778 | 0.723 | 0.91 | 0.591 |
| ALA | 71 | A | 0.487 | 0.534 | 0.958 | 0.063 | LEU | 3 | B | 0.704 | 0.696 | 0.979 | 0.421 |
| **HIS** | **72** | **A** | **0.826** | **0.873** | **0.925** | **0.774** | THR | 4 | B | 0.72 | 0.769 | 0.997 | 0.492 |
| VAL | 73 | A | 0.694 | 0.735 | 0.996 | 0.433 | PRO | 5 | B | 0.739 | 0.749 | 0.994 | 0.494 |

**Table S7(a): 1LFL (P), 1LFT (Q),** continued..

| Residue | | | S (P) | S (Q) | J (P,Q) | PRI |
|---|---|---|---|---|---|---|
| GLU | 6 | B | 0.772 | 0.692 | 0.949 | 0.515 |
| GLU | 7 | B | 0.814 | 0.799 | 0.993 | 0.62 |
| LYS | 8 | B | 0.715 | 0.758 | 0.959 | 0.514 |
| SER | 9 | B | 0.758 | 0.716 | 0.999 | 0.475 |
| ALA | 10 | B | 0.612 | 0.596 | 1 | 0.208 |
| VAL | 11 | B | 0.699 | 0.685 | 1 | 0.384 |
| THR | 12 | B | 0.682 | 0.811 | 0.997 | 0.496 |
| ALA | 13 | B | 0.58 | 0.651 | 1 | 0.231 |
| LEU | 14 | B | 0.672 | 0.668 | 0.997 | 0.343 |
| TRP | 15 | B | 0.845 | 0.846 | 0.999 | 0.692 |
| GLY | 16 | B | 0.647 | 0.637 | 1 | 0.284 |
| LYS | 17 | B | 0.73 | 0.725 | 0.978 | 0.477 |
| VAL | 18 | B | 0.704 | 0.683 | 0.996 | 0.391 |
| ASN | 19 | B | 0.753 | 0.77 | 0.99 | 0.533 |
| VAL | 20 | B | 0.608 | 0.718 | 0.818 | 0.508 |
| ASP | 21 | B | 0.824 | 0.685 | 0.996 | 0.513 |
| **GLU** | **22** | **B** | **0.798** | **0.738** | **0.709** | **0.827** |
| VAL | 23 | B | 0.727 | 0.703 | 1 | 0.43 |
| GLY | 24 | B | 0.623 | 0.624 | 0.999 | 0.248 |
| GLY | 25 | B | 0.629 | 0.626 | 0.998 | 0.257 |
| GLU | 26 | B | 0.803 | 0.761 | 0.982 | 0.582 |
| ALA | 27 | B | 0.621 | 0.615 | 0.999 | 0.237 |
| LEU | 28 | B | 0.659 | 0.675 | 0.844 | 0.49 |
| GLY | 29 | B | 0.631 | 0.628 | 1 | 0.259 |
| **ARG** | **30** | **B** | **0.838** | **0.849** | **0.927** | **0.76** |
| LEU | 31 | B | 0.721 | 0.703 | 0.999 | 0.425 |
| LEU | 32 | B | 0.722 | 0.772 | 0.886 | 0.608 |
| VAL | 33 | B | 0.704 | 0.707 | 1 | 0.411 |
| VAL | 34 | B | 0.682 | 0.666 | 0.999 | 0.349 |
| **TYR** | **35** | **B** | **0.861** | **0.873** | **1** | **0.734** |
| PRO | 36 | B | 0.693 | 0.685 | 0.999 | 0.379 |
| TRP | 37 | B | 0.844 | 0.858 | 0.997 | 0.705 |
| THR | 38 | B | 0.703 | 0.699 | 1 | 0.402 |
| GLN | 39 | B | 0.8 | 0.78 | 0.981 | 0.599 |
| ARG | 40 | B | 0.765 | 0.716 | 0.925 | 0.556 |
| PHE | 41 | B | 0.842 | 0.794 | 0.991 | 0.645 |
| PHE | 42 | B | 0.775 | 0.798 | 0.991 | 0.582 |
| GLU | 43 | B | 0.696 | 0.662 | 0.886 | 0.472 |
| SER | 44 | B | 0.597 | 0.663 | 0.93 | 0.33 |
| PHE | 45 | B | 0.83 | 0.836 | 0.998 | 0.668 |
| GLY | 46 | B | 0.649 | 0.652 | 0.999 | 0.302 |
| ASP | 47 | B | 0.779 | 0.708 | 0.997 | 0.49 |
| LEU | 48 | B | 0.759 | 0.724 | 0.998 | 0.485 |
| SER | 49 | B | 0.706 | 0.618 | 0.962 | 0.362 |
| THR | 50 | B | 0.777 | 0.786 | 0.999 | 0.564 |
| PRO | 51 | B | 0.666 | 0.733 | 0.997 | 0.402 |
| ASP | 52 | B | 0.678 | 0.65 | 0.99 | 0.338 |
| ALA | 53 | B | 0.629 | 0.584 | 1 | 0.213 |
| VAL | 54 | B | 0.658 | 0.698 | 1 | 0.356 |
| MET | 55 | B | 0.65 | 0.683 | 0.875 | 0.458 |
| GLY | 56 | B | 0.653 | 0.658 | 1 | 0.311 |
| ASN | 57 | B | 0.72 | 0.761 | 1 | 0.481 |
| PRO | 58 | B | 0.778 | 0.748 | 0.991 | 0.535 |
| LYS | 59 | B | 0.786 | 0.688 | 0.912 | 0.562 |
| VAL | 60 | B | 0.664 | 0.705 | 0.999 | 0.37 |
| LYS | 61 | B | 0.688 | 0.799 | 0.817 | 0.67 |
| ALA | 62 | B | 0.66 | 0.632 | 0.997 | 0.295 |
| HIS | 63 | B | 0.769 | 0.816 | 0.901 | 0.684 |
| GLY | 64 | B | 0.631 | 0.626 | 0.998 | 0.259 |
| LYS | 65 | B | 0.76 | 0.794 | 0.93 | 0.624 |
| LYS | 66 | B | 0.697 | 0.787 | 0.903 | 0.581 |
| VAL | 67 | B | 0.728 | 0.702 | 1 | 0.43 |
| LEU | 68 | B | 0.714 | 0.786 | 0.959 | 0.541 |
| GLY | 69 | B | 0.647 | 0.647 | 0.99 | 0.304 |
| ALA | 70 | B | 0.625 | 0.586 | 0.981 | 0.23 |
| **PHE** | **71** | **B** | **0.825** | **0.833** | **0.93** | **0.728** |
| SER | 72 | B | 0.697 | 0.707 | 0.97 | 0.434 |
| ASP | 73 | B | 0.747 | 0.629 | 0.964 | 0.412 |
| GLY | 74 | B | 0.63 | 0.633 | 0.999 | 0.264 |
| LEU | 75 | B | 0.688 | 0.707 | 0.995 | 0.4 |
| ALA | 76 | B | 0.462 | 0.459 | 0.999 | -0.078 |
| **HIS** | **77** | **B** | **0.826** | **0.855** | **0.943** | **0.738** |
| LEU | 78 | B | 0.681 | 0.771 | 0.927 | 0.525 |
| ASP | 79 | B | 0.683 | 0.678 | 0.804 | 0.557 |
| ASN | 80 | B | 0.776 | 0.76 | 0.988 | 0.548 |
| LEU | 81 | B | 0.677 | 0.652 | 0.915 | 0.414 |
| **LYS** | **82** | **B** | **0.673** | **0.722** | **0.678** | **0.717** |
| GLY | 83 | B | 0.646 | 0.651 | 0.999 | 0.298 |
| THR | 84 | B | 0.692 | 0.686 | 0.973 | 0.405 |
| PHE | 85 | B | 0.842 | 0.829 | 0.995 | 0.676 |
| ALA | 86 | B | 0.651 | 0.664 | 1 | 0.315 |
| THR | 87 | B | 0.626 | 0.686 | 0.919 | 0.393 |
| LEU | 88 | B | 0.689 | 0.705 | 0.999 | 0.395 |
| SER | 89 | B | 0.708 | 0.727 | 0.998 | 0.437 |
| GLU | 90 | B | 0.765 | 0.855 | 0.946 | 0.674 |
| LEU | 91 | B | 0.714 | 0.693 | 0.968 | 0.439 |
| HIS | 92 | B | 0.808 | 0.775 | 0.985 | 0.598 |
| CYS | 93 | B | 0.637 | 0.652 | 0.834 | 0.455 |
| ASP | 94 | B | 0.8 | 0.738 | 0.992 | 0.546 |
| LYS | 95 | B | 0.734 | 0.691 | 0.994 | 0.431 |
| LEU | 96 | B | 0.731 | 0.674 | 0.994 | 0.411 |
| HIS | 97 | B | 0.848 | 0.839 | 0.999 | 0.688 |
| VAL | 98 | B | 0.706 | 0.713 | 0.999 | 0.42 |
| ASP | 99 | B | 0.77 | 0.774 | 0.988 | 0.556 |
| PRO | 100 | B | 0.711 | 0.687 | 0.975 | 0.423 |
| GLU | 101 | B | 0.735 | 0.672 | 0.985 | 0.422 |
| **ASN** | **102** | **B** | **0.78** | **0.742** | **0.797** | **0.725** |
| PHE | 103 | B | 0.829 | 0.827 | 0.994 | 0.662 |
| **ARG** | **104** | **B** | **0.71** | **0.792** | **0.781** | **0.721** |
| LEU | 105 | B | 0.713 | 0.702 | 0.98 | 0.435 |
| LEU | 106 | B | 0.713 | 0.724 | 0.999 | 0.438 |
| GLY | 107 | B | 0.63 | 0.627 | 0.999 | 0.258 |
| ASN | 108 | B | 0.782 | 0.75 | 0.923 | 0.609 |
| VAL | 109 | B | 0.674 | 0.757 | 0.841 | 0.59 |
| LEU | 110 | B | 0.721 | 0.711 | 0.923 | 0.509 |
| VAL | 111 | B | 0.719 | 0.652 | 0.999 | 0.372 |
| CYS | 112 | B | 0.638 | 0.583 | 1 | 0.221 |
| VAL | 113 | B | 0.718 | 0.705 | 1 | 0.423 |
| LEU | 114 | B | 0.728 | 0.736 | 1 | 0.464 |
| ALA | 115 | B | 0.559 | 0.508 | 1 | 0.067 |
| HIS | 116 | B | 0.748 | 0.796 | 0.999 | 0.545 |
| **HIS** | **117** | **B** | **0.805** | **0.791** | **0.796** | **0.8** |
| PHE | 118 | B | 0.843 | 0.846 | 1 | 0.689 |
| GLY | 119 | B | 0.629 | 0.628 | 0.997 | 0.26 |
| LYS | 120 | B | 0.543 | 0.671 | 0.737 | 0.477 |
| GLU | 121 | B | 0.744 | 0.653 | 0.964 | 0.433 |
| PHE | 122 | B | 0.844 | 0.82 | 0.999 | 0.665 |
| THR | 123 | B | 0.757 | 0.773 | 1 | 0.53 |
| PRO | 124 | B | 0.656 | 0.657 | 1 | 0.313 |
| PRO | 125 | B | 0.71 | 0.703 | 1 | 0.413 |
| VAL | 126 | B | 0.714 | 0.662 | 1 | 0.376 |
| GLN | 127 | B | 0.777 | 0.766 | 1 | 0.543 |
| ALA | 128 | B | 0.583 | 0.514 | 0.999 | 0.098 |
| ALA | 129 | B | 0.557 | 0.566 | 1 | 0.123 |
| TYR | 130 | B | 0.844 | 0.822 | 1 | 0.666 |
| GLN | 131 | B | 0.736 | 0.764 | 1 | 0.5 |
| LYS | 132 | B | 0.775 | 0.772 | 0.999 | 0.548 |
| VAL | 133 | B | 0.695 | 0.699 | 1 | 0.394 |
| VAL | 134 | B | 0.698 | 0.713 | 0.897 | 0.514 |
| ALA | 135 | B | 0.62 | 0.636 | 0.999 | 0.257 |
| GLY | 136 | B | 0.635 | 0.639 | 1 | 0.274 |
| VAL | 137 | B | 0.678 | 0.693 | 0.998 | 0.373 |
| ALA | 138 | B | 0.564 | 0.516 | 0.996 | 0.084 |
| ASN | 139 | B | 0.58 | 0.817 | 0.967 | 0.43 |
| ALA | 140 | B | 0.563 | 0.545 | 0.998 | 0.11 |
| LEU | 141 | B | 0.701 | 0.71 | 0.999 | 0.412 |
| ALA | 142 | B | 0.465 | 0.462 | 0.999 | -0.072 |
| HIS | 143 | B | 0.818 | 0.816 | 0.97 | 0.664 |
| LYS | 144 | B | 0.78 | 0.823 | 0.98 | 0.623 |
| TYR | 145 | B | 0.849 | 0.804 | 0.989 | 0.664 |
| **HIS** | **146** | **B** | **0.847** | **0.803** | **0.856** | **0.794** |

| Table S7(b): 2DN1 (P), 2DN2 (Q) | | | | | | | | | | | | | | |
|---|---|---|---|---|---|---|---|---|---|---|---|---|---|---|
| Residue | | | S (P) | S (Q) | J (P,Q) | PRI | | Residue | | | S (P) | S (Q) | J (P,Q) | PRI |
| LEU | 2 | A | 0.765 | 0.753 | 0.997 | 0.521 | | ASP | 75 | A | 0.787 | 0.781 | 0.984 | 0.584 |
| SER | 3 | A | 0.743 | 0.73 | 0.999 | 0.474 | | MET | 76 | A | 0.738 | 0.732 | 0.991 | 0.479 |
| PRO | 4 | A | 0.757 | 0.749 | 0.999 | 0.507 | | PRO | 77 | A | 0.696 | 0.691 | 0.998 | 0.389 |
| ALA | 5 | A | 0.661 | 0.63 | 1 | 0.291 | | ASN | 78 | A | 0.677 | 0.622 | 0.996 | 0.303 |
| ASP | 6 | A | 0.815 | 0.77 | 0.999 | 0.586 | | ALA | 79 | A | 0.506 | 0.483 | 0.999 | -0.01 |
| LYS | 7 | A | 0.8 | 0.791 | 0.987 | 0.604 | | LEU | 80 | A | 0.716 | 0.708 | 0.998 | 0.426 |
| THR | 8 | A | 0.756 | 0.755 | 0.998 | 0.513 | | SER | 81 | A | 0.752 | 0.752 | 0.998 | 0.506 |
| ASN | 9 | A | 0.778 | 0.763 | 1 | 0.541 | | ALA | 82 | A | 0.667 | 0.679 | 0.999 | 0.347 |
| VAL | 10 | A | 0.727 | 0.684 | 1 | 0.411 | | LEU | 83 | A | 0.711 | 0.691 | 0.972 | 0.43 |
| LYS | 11 | A | 0.734 | 0.809 | 0.875 | 0.668 | | SER | 84 | A | 0.673 | 0.667 | 0.895 | 0.445 |
| ALA | 12 | A | 0.619 | 0.637 | 0.991 | 0.265 | | ASP | 85 | A | 0.784 | 0.785 | 1 | 0.569 |
| ALA | 13 | A | 0.568 | 0.584 | 0.998 | 0.154 | | LEU | 86 | A | 0.74 | 0.767 | 0.994 | 0.513 |
| **TRP** | **14** | **A** | **0.845** | **0.848** | **0.735** | **0.958** | | HIS | 87 | A | 0.787 | 0.815 | 0.987 | 0.615 |
| GLY | 15 | A | 0.653 | 0.656 | 0.997 | 0.312 | | ALA | 88 | A | 0.514 | 0.524 | 0.995 | 0.043 |
| **LYS** | **16** | **A** | **0.8** | **0.755** | **0.785** | **0.77** | | HIS | 89 | A | 0.847 | 0.803 | 0.971 | 0.679 |
| VAL | 17 | A | 0.711 | 0.688 | 0.997 | 0.402 | | LYS | 90 | A | 0.669 | 0.675 | 0.913 | 0.431 |
| GLY | 18 | A | 0.648 | 0.653 | 0.997 | 0.304 | | LEU | 91 | A | 0.714 | 0.686 | 0.999 | 0.401 |
| ALA | 19 | A | 0.524 | 0.544 | 0.999 | 0.069 | | ARG | 92 | A | 0.754 | 0.852 | 0.952 | 0.654 |
| HIS | 20 | A | 0.802 | 0.77 | 1 | 0.572 | | VAL | 93 | A | 0.682 | 0.677 | 1 | 0.359 |
| ALA | 21 | A | 0.596 | 0.536 | 0.999 | 0.133 | | ASP | 94 | A | 0.643 | 0.804 | 0.948 | 0.499 |
| GLY | 22 | A | 0.623 | 0.621 | 1 | 0.244 | | PRO | 95 | A | 0.699 | 0.682 | 0.997 | 0.384 |
| GLU | 23 | A | 0.79 | 0.656 | 0.936 | 0.51 | | VAL | 96 | A | 0.713 | 0.746 | 0.931 | 0.528 |
| **TYR** | **24** | **A** | **0.876** | **0.855** | **0.999** | **0.732** | | ASN | 97 | A | 0.739 | 0.745 | 0.996 | 0.488 |
| GLY | 25 | A | 0.629 | 0.635 | 0.999 | 0.265 | | PHE | 98 | A | 0.816 | 0.822 | 1 | 0.638 |
| ALA | 26 | A | 0.565 | 0.572 | 1 | 0.137 | | LYS | 99 | A | 0.748 | 0.746 | 0.972 | 0.522 |
| GLU | 27 | A | 0.775 | 0.772 | 0.927 | 0.62 | | LEU | 100 | A | 0.747 | 0.736 | 0.999 | 0.484 |
| ALA | 28 | A | 0.566 | 0.536 | 1 | 0.102 | | LEU | 101 | A | 0.678 | 0.66 | 1 | 0.338 |
| LEU | 29 | A | 0.703 | 0.715 | 1 | 0.418 | | SER | 102 | A | 0.713 | 0.72 | 1 | 0.433 |
| GLU | 30 | A | 0.786 | 0.783 | 0.965 | 0.604 | | HIS | 103 | A | 0.815 | 0.783 | 0.999 | 0.599 |
| **ARG** | **31** | **A** | **0.87** | **0.864** | **0.965** | **0.769** | | CYS | 104 | A | 0.613 | 0.609 | 1 | 0.222 |
| MET | 32 | A | 0.761 | 0.758 | 1 | 0.519 | | LEU | 105 | A | 0.764 | 0.746 | 0.997 | 0.513 |
| PHE | 33 | A | 0.851 | 0.841 | 1 | 0.692 | | LEU | 106 | A | 0.719 | 0.746 | 0.999 | 0.466 |
| LEU | 34 | A | 0.677 | 0.679 | 1 | 0.356 | | VAL | 107 | A | 0.709 | 0.702 | 1 | 0.411 |
| SER | 35 | A | 0.65 | 0.615 | 1 | 0.265 | | THR | 108 | A | 0.702 | 0.686 | 0.999 | 0.389 |
| PHE | 36 | A | 0.839 | 0.835 | 1 | 0.674 | | LEU | 109 | A | 0.759 | 0.74 | 0.999 | 0.5 |
| PRO | 37 | A | 0.7 | 0.693 | 1 | 0.393 | | ALA | 110 | A | 0.518 | 0.529 | 1 | 0.047 |
| THR | 38 | A | 0.64 | 0.686 | 1 | 0.326 | | ALA | 111 | A | 0.433 | 0.443 | 0.999 | -0.123 |
| THR | 39 | A | 0.692 | 0.695 | 1 | 0.387 | | HIS | 112 | A | 0.828 | 0.817 | 0.999 | 0.646 |
| LYS | 40 | A | 0.773 | 0.77 | 0.999 | 0.544 | | LEU | 113 | A | 0.73 | 0.728 | 0.999 | 0.459 |
| THR | 41 | A | 0.712 | 0.744 | 0.999 | 0.457 | | PRO | 114 | A | 0.677 | 0.734 | 0.995 | 0.416 |
| TYR | 42 | A | 0.832 | 0.853 | 1 | 0.685 | | ALA | 115 | A | 0.628 | 0.577 | 0.991 | 0.214 |
| PHE | 43 | A | 0.846 | 0.836 | 0.999 | 0.683 | | GLU | 116 | A | 0.778 | 0.772 | 0.999 | 0.551 |
| PRO | 44 | A | 0.727 | 0.713 | 0.998 | 0.442 | | PHE | 117 | A | 0.829 | 0.815 | 0.998 | 0.646 |
| HIS | 45 | A | 0.814 | 0.824 | 0.999 | 0.639 | | THR | 118 | A | 0.751 | 0.711 | 1 | 0.462 |
| PHE | 46 | A | 0.83 | 0.826 | 0.99 | 0.666 | | PRO | 119 | A | 0.663 | 0.659 | 1 | 0.322 |
| ASP | 47 | A | 0.779 | 0.807 | 0.997 | 0.589 | | ALA | 120 | A | 0.627 | 0.611 | 1 | 0.238 |
| LEU | 48 | A | 0.73 | 0.724 | 0.999 | 0.455 | | VAL | 121 | A | 0.743 | 0.712 | 1 | 0.455 |
| SER | 49 | A | 0.763 | 0.748 | 0.998 | 0.513 | | HIS | 122 | A | 0.779 | 0.788 | 0.999 | 0.568 |
| HIS | 50 | A | 0.764 | 0.758 | 0.999 | 0.523 | | ALA | 123 | A | 0.577 | 0.579 | 1 | 0.156 |
| GLY | 51 | A | 0.65 | 0.645 | 0.999 | 0.296 | | SER | 124 | A | 0.675 | 0.675 | 0.999 | 0.351 |
| SER | 52 | A | 0.707 | 0.662 | 0.987 | 0.382 | | LEU | 125 | A | 0.736 | 0.678 | 0.999 | 0.415 |
| ALA | 53 | A | 0.639 | 0.664 | 0.999 | 0.304 | | ASP | 126 | A | 0.752 | 0.758 | 0.888 | 0.622 |
| GLN | 54 | A | 0.729 | 0.727 | 0.998 | 0.458 | | **LYS** | **127** | **A** | **0.745** | **0.792** | **0.703** | **0.834** |
| VAL | 55 | A | 0.662 | 0.684 | 1 | 0.346 | | PHE | 128 | A | 0.816 | 0.828 | 0.999 | 0.645 |
| LYS | 56 | A | 0.775 | 0.787 | 0.998 | 0.564 | | LEU | 129 | A | 0.732 | 0.714 | 0.999 | 0.447 |
| GLY | 57 | A | 0.649 | 0.652 | 1 | 0.301 | | ALA | 130 | A | 0.585 | 0.592 | 0.999 | 0.178 |
| HIS | 58 | A | 0.788 | 0.778 | 1 | 0.566 | | SER | 131 | A | 0.715 | 0.721 | 0.998 | 0.438 |
| GLY | 59 | A | 0.628 | 0.629 | 0.999 | 0.258 | | VAL | 132 | A | 0.713 | 0.707 | 0.997 | 0.423 |
| LYS | 60 | A | 0.817 | 0.795 | 0.997 | 0.615 | | SER | 133 | A | 0.712 | 0.711 | 0.998 | 0.425 |
| LYS | 61 | A | 0.734 | 0.7 | 0.994 | 0.44 | | THR | 134 | A | 0.727 | 0.746 | 0.999 | 0.474 |
| VAL | 62 | A | 0.702 | 0.698 | 0.999 | 0.401 | | VAL | 135 | A | 0.716 | 0.695 | 0.994 | 0.417 |
| ALA | 63 | A | 0.586 | 0.587 | 0.999 | 0.174 | | LEU | 136 | A | 0.757 | 0.736 | 0.993 | 0.5 |
| ASP | 64 | A | 0.822 | 0.795 | 0.999 | 0.618 | | THR | 137 | A | 0.672 | 0.68 | 0.998 | 0.354 |
| ALA | 65 | A | 0.598 | 0.573 | 0.999 | 0.172 | | SER | 138 | A | 0.617 | 0.646 | 0.998 | 0.265 |
| LEU | 66 | A | 0.721 | 0.716 | 0.999 | 0.438 | | LYS | 139 | A | 0.774 | 0.783 | 0.99 | 0.567 |
| THR | 67 | A | 0.75 | 0.701 | 0.99 | 0.461 | | TYR | 140 | A | 0.821 | 0.797 | 0.97 | 0.648 |
| ASN | 68 | A | 0.761 | 0.733 | 0.997 | 0.497 | | **ARG** | **141** | **A** | **0.915** | **0.662** | **0.812** | **0.765** |
| ALA | 69 | A | 0.552 | 0.543 | 1 | 0.095 | | HIS | 2 | B | 0.685 | 0.647 | 0.993 | 0.339 |
| VAL | 70 | A | 0.726 | 0.676 | 0.889 | 0.513 | | LEU | 3 | B | 0.716 | 0.708 | 0.986 | 0.438 |
| ALA | 71 | A | 0.497 | 0.534 | 1 | 0.031 | | THR | 4 | B | 0.756 | 0.722 | 0.999 | 0.479 |
| HIS | 72 | A | 0.841 | 0.791 | 0.916 | 0.716 | | PRO | 5 | B | 0.757 | 0.749 | 1 | 0.506 |
| VAL | 73 | A | 0.695 | 0.674 | 0.998 | 0.371 | | GLU | 6 | B | 0.659 | 0.837 | 0.905 | 0.591 |
| ASP | 74 | A | 0.769 | 0.732 | 0.97 | 0.531 | | GLU | 7 | B | 0.837 | 0.831 | 0.998 | 0.67 |

**Table S7(b): 2DN1 (P), 2DN2 (Q),** continued..

| **Residue** | | | **S (P)** | **S (Q)** | **J (P,Q)** | **PRI** |
|---|---|---|---|---|---|---|
| LYS | 8 | B | 0.786 | 0.69 | 0.977 | 0.499 |
| SER | 9 | B | 0.678 | 0.68 | 0.992 | 0.366 |
| ALA | 10 | B | 0.557 | 0.591 | 0.992 | 0.156 |
| VAL | 11 | B | 0.723 | 0.723 | 0.998 | 0.448 |
| THR | 12 | B | 0.744 | 0.715 | 0.928 | 0.531 |
| ALA | 13 | B | 0.584 | 0.624 | 1 | 0.208 |
| LEU | 14 | B | 0.742 | 0.718 | 1 | 0.46 |
| **TRP** | **15** | **B** | **0.865** | **0.866** | **1** | **0.731** |
| GLY | 16 | B | 0.654 | 0.645 | 1 | 0.299 |
| LYS | 17 | B | 0.75 | 0.703 | 0.997 | 0.456 |
| VAL | 18 | B | 0.65 | 0.657 | 1 | 0.307 |
| ASN | 19 | B | 0.765 | 0.781 | 0.997 | 0.549 |
| VAL | 20 | B | 0.683 | 0.653 | 0.906 | 0.43 |
| ASP | 21 | B | 0.721 | 0.786 | 0.973 | 0.534 |
| **GLU** | **22** | **B** | **0.806** | **0.791** | **0.749** | **0.848** |
| VAL | 23 | B | 0.686 | 0.683 | 1 | 0.369 |
| GLY | 24 | B | 0.623 | 0.624 | 0.998 | 0.249 |
| GLY | 25 | B | 0.622 | 0.634 | 0.998 | 0.258 |
| GLU | 26 | B | 0.774 | 0.749 | 0.885 | 0.638 |
| ALA | 27 | B | 0.552 | 0.564 | 0.999 | 0.117 |
| LEU | 28 | B | 0.716 | 0.701 | 1 | 0.417 |
| GLY | 29 | B | 0.628 | 0.627 | 1 | 0.255 |
| ARG | 30 | B | 0.837 | 0.839 | 1 | 0.676 |
| LEU | 31 | B | 0.726 | 0.72 | 0.998 | 0.448 |
| LEU | 32 | B | 0.74 | 0.731 | 0.916 | 0.555 |
| VAL | 33 | B | 0.689 | 0.697 | 0.999 | 0.387 |
| VAL | 34 | B | 0.697 | 0.698 | 0.999 | 0.396 |
| **TYR** | **35** | **B** | **0.873** | **0.869** | **0.999** | **0.743** |
| PRO | 36 | B | 0.695 | 0.69 | 0.999 | 0.386 |
| **TRP** | **37** | **B** | **0.867** | **0.884** | **0.993** | **0.758** |
| THR | 38 | B | 0.687 | 0.712 | 1 | 0.399 |
| GLN | 39 | B | 0.777 | 0.787 | 0.999 | 0.565 |
| ARG | 40 | B | 0.765 | 0.693 | 0.931 | 0.527 |
| PHE | 41 | B | 0.827 | 0.834 | 0.997 | 0.664 |
| PHE | 42 | B | 0.831 | 0.847 | 1 | 0.678 |
| GLU | 43 | B | 0.837 | 0.686 | 0.822 | 0.701 |
| SER | 44 | B | 0.632 | 0.617 | 0.999 | 0.25 |
| PHE | 45 | B | 0.828 | 0.844 | 0.999 | 0.673 |
| GLY | 46 | B | 0.653 | 0.643 | 1 | 0.296 |
| ASP | 47 | B | 0.774 | 0.751 | 1 | 0.525 |
| LEU | 48 | B | 0.743 | 0.734 | 0.999 | 0.478 |
| SER | 49 | B | 0.699 | 0.656 | 0.999 | 0.356 |
| THR | 50 | B | 0.801 | 0.788 | 1 | 0.589 |
| PRO | 51 | B | 0.683 | 0.676 | 1 | 0.359 |
| ASP | 52 | B | 0.614 | 0.63 | 1 | 0.244 |
| ALA | 53 | B | 0.55 | 0.578 | 1 | 0.128 |
| VAL | 54 | B | 0.703 | 0.707 | 1 | 0.41 |
| MET | 55 | B | 0.739 | 0.709 | 0.999 | 0.449 |
| GLY | 56 | B | 0.639 | 0.642 | 1 | 0.281 |
| ASN | 57 | B | 0.743 | 0.745 | 1 | 0.488 |
| PRO | 58 | B | 0.735 | 0.727 | 0.998 | 0.464 |
| LYS | 59 | B | 0.747 | 0.704 | 0.968 | 0.483 |
| VAL | 60 | B | 0.668 | 0.681 | 1 | 0.349 |
| LYS | 61 | B | 0.708 | 0.757 | 0.866 | 0.599 |
| ALA | 62 | B | 0.651 | 0.597 | 1 | 0.248 |
| HIS | 63 | B | 0.773 | 0.785 | 0.999 | 0.559 |
| GLY | 64 | B | 0.631 | 0.627 | 1 | 0.258 |
| LYS | 65 | B | 0.755 | 0.78 | 0.847 | 0.688 |
| LYS | 66 | B | 0.759 | 0.722 | 0.988 | 0.493 |
| VAL | 67 | B | 0.701 | 0.73 | 0.998 | 0.433 |
| LEU | 68 | B | 0.731 | 0.726 | 0.825 | 0.632 |
| GLY | 69 | B | 0.647 | 0.639 | 0.995 | 0.291 |
| ALA | 70 | B | 0.557 | 0.575 | 0.988 | 0.144 |
| **PHE** | **71** | **B** | **0.827** | **0.824** | **0.918** | **0.733** |
| SER | 72 | B | 0.709 | 0.705 | 0.983 | 0.431 |
| ASP | 73 | B | 0.765 | 0.788 | 0.926 | 0.627 |
| GLY | 74 | B | 0.629 | 0.633 | 0.997 | 0.265 |
| LEU | 75 | B | 0.741 | 0.74 | 0.997 | 0.484 |
| ALA | 76 | B | 0.454 | 0.519 | 0.995 | -0.022 |
| HIS | 77 | B | 0.817 | 0.84 | 0.998 | 0.659 |
| LEU | 78 | B | 0.724 | 0.706 | 0.999 | 0.431 |
| ASP | 79 | B | 0.683 | 0.696 | 0.838 | 0.541 |
| ASN | 80 | B | 0.748 | 0.751 | 0.996 | 0.503 |
| LEU | 81 | B | 0.727 | 0.725 | 1 | 0.452 |
| LYS | 82 | B | 0.727 | 0.734 | 0.813 | 0.648 |
| GLY | 83 | B | 0.658 | 0.65 | 0.995 | 0.313 |
| THR | 84 | B | 0.63 | 0.564 | 0.996 | 0.198 |
| PHE | 85 | B | 0.831 | 0.842 | 0.993 | 0.68 |
| ALA | 86 | B | 0.63 | 0.626 | 0.999 | 0.257 |
| THR | 87 | B | 0.68 | 0.693 | 0.873 | 0.5 |
| LEU | 88 | B | 0.722 | 0.702 | 0.998 | 0.426 |
| SER | 89 | B | 0.693 | 0.72 | 0.999 | 0.414 |
| GLU | 90 | B | 0.793 | 0.651 | 0.994 | 0.45 |
| LEU | 91 | B | 0.755 | 0.752 | 0.974 | 0.533 |
| HIS | 92 | B | 0.751 | 0.794 | 0.991 | 0.554 |
| CYS | 93 | B | 0.661 | 0.642 | 0.852 | 0.451 |
| ASP | 94 | B | 0.689 | 0.768 | 0.981 | 0.476 |
| LYS | 95 | B | 0.716 | 0.741 | 0.972 | 0.485 |
| LEU | 96 | B | 0.694 | 0.742 | 0.997 | 0.439 |
| **HIS** | **97** | **B** | **0.858** | **0.866** | **0.997** | **0.727** |
| VAL | 98 | B | 0.691 | 0.682 | 1 | 0.373 |
| ASP | 99 | B | 0.79 | 0.739 | 0.997 | 0.532 |
| PRO | 100 | B | 0.682 | 0.671 | 0.982 | 0.371 |
| GLU | 101 | B | 0.731 | 0.752 | 0.953 | 0.53 |
| ASN | 102 | B | 0.741 | 0.777 | 0.97 | 0.548 |
| PHE | 103 | B | 0.84 | 0.838 | 0.992 | 0.686 |
| **ARG** | **104** | **B** | **0.783** | **0.788** | **0.731** | **0.84** |
| LEU | 105 | B | 0.75 | 0.722 | 0.989 | 0.483 |
| LEU | 106 | B | 0.721 | 0.714 | 0.998 | 0.437 |
| GLY | 107 | B | 0.626 | 0.634 | 1 | 0.26 |
| ASN | 108 | B | 0.78 | 0.716 | 0.994 | 0.502 |
| VAL | 109 | B | 0.727 | 0.703 | 0.999 | 0.431 |
| LEU | 110 | B | 0.738 | 0.728 | 1 | 0.466 |
| VAL | 111 | B | 0.714 | 0.697 | 0.999 | 0.412 |
| CYS | 112 | B | 0.606 | 0.599 | 0.999 | 0.206 |
| VAL | 113 | B | 0.698 | 0.703 | 1 | 0.401 |
| LEU | 114 | B | 0.74 | 0.707 | 1 | 0.447 |
| ALA | 115 | B | 0.538 | 0.538 | 1 | 0.076 |
| HIS | 116 | B | 0.775 | 0.785 | 0.999 | 0.561 |
| HIS | 117 | B | 0.751 | 0.757 | 0.984 | 0.524 |
| PHE | 118 | B | 0.849 | 0.846 | 1 | 0.695 |
| GLY | 119 | B | 0.627 | 0.628 | 1 | 0.255 |
| LYS | 120 | B | 0.593 | 0.649 | 0.724 | 0.518 |
| GLU | 121 | B | 0.74 | 0.638 | 0.962 | 0.416 |
| PHE | 122 | B | 0.843 | 0.842 | 1 | 0.685 |
| THR | 123 | B | 0.768 | 0.766 | 1 | 0.534 |
| PRO | 124 | B | 0.673 | 0.663 | 1 | 0.336 |
| PRO | 125 | B | 0.717 | 0.704 | 1 | 0.421 |
| VAL | 126 | B | 0.677 | 0.669 | 1 | 0.346 |
| GLN | 127 | B | 0.766 | 0.777 | 1 | 0.543 |
| ALA | 128 | B | 0.57 | 0.566 | 1 | 0.136 |
| ALA | 129 | B | 0.561 | 0.558 | 1 | 0.119 |
| TYR | 130 | B | 0.836 | 0.837 | 0.999 | 0.674 |
| GLN | 131 | B | 0.772 | 0.755 | 1 | 0.527 |
| LYS | 132 | B | 0.776 | 0.798 | 0.999 | 0.575 |
| VAL | 133 | B | 0.716 | 0.703 | 1 | 0.419 |
| VAL | 134 | B | 0.691 | 0.709 | 0.999 | 0.401 |
| ALA | 135 | B | 0.582 | 0.607 | 1 | 0.189 |
| GLY | 136 | B | 0.638 | 0.637 | 1 | 0.275 |
| VAL | 137 | B | 0.705 | 0.699 | 1 | 0.404 |
| ALA | 138 | B | 0.593 | 0.546 | 0.999 | 0.14 |
| ASN | 139 | B | 0.764 | 0.808 | 0.901 | 0.671 |
| ALA | 140 | B | 0.539 | 0.516 | 1 | 0.055 |
| LEU | 141 | B | 0.74 | 0.73 | 0.891 | 0.579 |
| ALA | 142 | B | 0.472 | 0.474 | 0.999 | -0.053 |
| HIS | 143 | B | 0.805 | 0.825 | 0.997 | 0.633 |
| LYS | 144 | B | 0.718 | 0.792 | 0.966 | 0.544 |
| TYR | 145 | B | 0.832 | 0.851 | 0.986 | 0.697 |
| **HIS** | **146** | **B** | **0.91** | **0.85** | **0.882** | **0.878** |

**Table S7(c): 2DN1 (P), 2DN3 (Q)**

| Residue | | | S (P) | S (Q) | J (P,Q) | PRI | Residue | | | S (P) | S (Q) | J (P,Q) | PRI |
|---|---|---|---|---|---|---|---|---|---|---|---|---|---|
| LEU | 2 | A | 0.765 | 0.761 | 0.998 | 0.528 | ASP | 75 | A | 0.787 | 0.799 | 0.997 | 0.589 |
| SER | 3 | A | 0.743 | 0.74 | 0.997 | 0.486 | MET | 76 | A | 0.738 | 0.723 | 0.843 | 0.618 |
| PRO | 4 | A | 0.757 | 0.755 | 0.998 | 0.514 | PRO | 77 | A | 0.696 | 0.709 | 0.999 | 0.406 |
| ALA | 5 | A | 0.661 | 0.641 | 0.998 | 0.304 | ASN | 78 | A | 0.677 | 0.699 | 0.991 | 0.385 |
| ASP | 6 | A | 0.815 | 0.807 | 0.999 | 0.623 | ALA | 79 | A | 0.506 | 0.481 | 1 | -0.013 |
| LYS | 7 | A | 0.8 | 0.764 | 0.884 | 0.68 | LEU | 80 | A | 0.716 | 0.731 | 0.977 | 0.47 |
| THR | 8 | A | 0.756 | 0.759 | 0.982 | 0.533 | SER | 81 | A | 0.752 | 0.722 | 0.998 | 0.476 |
| ASN | 9 | A | 0.778 | 0.806 | 0.986 | 0.598 | ALA | 82 | A | 0.667 | 0.678 | 0.999 | 0.346 |
| VAL | 10 | A | 0.727 | 0.71 | 0.994 | 0.443 | **LEU** | **83** | **A** | **0.711** | **0.773** | **0.615** | **0.869** |
| LYS | 11 | A | 0.734 | 0.758 | 0.785 | 0.707 | SER | 84 | A | 0.673 | 0.623 | 0.818 | 0.478 |
| ALA | 12 | A | 0.619 | 0.623 | 0.914 | 0.328 | ASP | 85 | A | 0.784 | 0.796 | 0.997 | 0.583 |
| ALA | 13 | A | 0.568 | 0.592 | 0.966 | 0.194 | LEU | 86 | A | 0.74 | 0.73 | 0.999 | 0.471 |
| **TRP** | **14** | **A** | **0.845** | **0.838** | **0.738** | **0.945** | HIS | 87 | A | 0.787 | 0.758 | 0.999 | 0.546 |
| GLY | 15 | A | 0.653 | 0.642 | 0.992 | 0.303 | ALA | 88 | A | 0.514 | 0.549 | 1 | 0.063 |
| LYS | 16 | A | 0.8 | 0.772 | 0.957 | 0.615 | HIS | 89 | A | 0.847 | 0.842 | 0.999 | 0.69 |
| VAL | 17 | A | 0.711 | 0.719 | 0.983 | 0.447 | LYS | 90 | A | 0.669 | 0.784 | 0.763 | 0.69 |
| GLY | 18 | A | 0.648 | 0.651 | 0.974 | 0.325 | LEU | 91 | A | 0.714 | 0.739 | 0.996 | 0.457 |
| ALA | 19 | A | 0.524 | 0.55 | 0.972 | 0.102 | ARG | 92 | A | 0.754 | 0.756 | 0.993 | 0.517 |
| HIS | 20 | A | 0.802 | 0.801 | 0.97 | 0.633 | VAL | 93 | A | 0.682 | 0.692 | 1 | 0.374 |
| ALA | 21 | A | 0.596 | 0.671 | 0.912 | 0.355 | ASP | 94 | A | 0.643 | 0.642 | 1 | 0.285 |
| GLY | 22 | A | 0.623 | 0.626 | 0.972 | 0.277 | PRO | 95 | A | 0.699 | 0.706 | 1 | 0.405 |
| GLU | 23 | A | 0.79 | 0.774 | 0.996 | 0.568 | VAL | 96 | A | 0.713 | 0.717 | 1 | 0.43 |
| **TYR** | **24** | **A** | **0.876** | **0.869** | **0.993** | **0.752** | ASN | 97 | A | 0.739 | 0.751 | 1 | 0.49 |
| GLY | 25 | A | 0.629 | 0.634 | 1 | 0.263 | PHE | 98 | A | 0.816 | 0.817 | 0.999 | 0.634 |
| ALA | 26 | A | 0.565 | 0.545 | 1 | 0.11 | LYS | 99 | A | 0.748 | 0.722 | 1 | 0.47 |
| GLU | 27 | A | 0.775 | 0.775 | 0.999 | 0.551 | LEU | 100 | A | 0.747 | 0.74 | 1 | 0.487 |
| ALA | 28 | A | 0.566 | 0.556 | 1 | 0.122 | LEU | 101 | A | 0.678 | 0.7 | 1 | 0.378 |
| LEU | 29 | A | 0.703 | 0.707 | 1 | 0.41 | SER | 102 | A | 0.713 | 0.704 | 1 | 0.417 |
| **GLU** | **30** | **A** | **0.786** | **0.782** | **0.766** | **0.802** | HIS | 103 | A | 0.815 | 0.804 | 1 | 0.619 |
| **ARG** | **31** | **A** | **0.87** | **0.867** | **0.79** | **0.947** | CYS | 104 | A | 0.613 | 0.613 | 1 | 0.226 |
| MET | 32 | A | 0.761 | 0.759 | 1 | 0.52 | LEU | 105 | A | 0.764 | 0.769 | 0.998 | 0.535 |
| PHE | 33 | A | 0.851 | 0.848 | 1 | 0.699 | LEU | 106 | A | 0.719 | 0.736 | 0.999 | 0.456 |
| LEU | 34 | A | 0.677 | 0.65 | 1 | 0.327 | VAL | 107 | A | 0.709 | 0.713 | 1 | 0.422 |
| SER | 35 | A | 0.65 | 0.627 | 1 | 0.277 | THR | 108 | A | 0.702 | 0.691 | 0.999 | 0.394 |
| PHE | 36 | A | 0.839 | 0.84 | 1 | 0.679 | LEU | 109 | A | 0.759 | 0.72 | 0.998 | 0.481 |
| PRO | 37 | A | 0.7 | 0.703 | 1 | 0.403 | ALA | 110 | A | 0.518 | 0.511 | 0.998 | 0.031 |
| THR | 38 | A | 0.64 | 0.645 | 1 | 0.285 | ALA | 111 | A | 0.433 | 0.459 | 0.999 | -0.107 |
| THR | 39 | A | 0.692 | 0.687 | 1 | 0.379 | HIS | 112 | A | 0.828 | 0.822 | 0.992 | 0.658 |
| LYS | 40 | A | 0.773 | 0.774 | 1 | 0.547 | LEU | 113 | A | 0.73 | 0.753 | 0.854 | 0.629 |
| THR | 41 | A | 0.712 | 0.73 | 0.997 | 0.445 | PRO | 114 | A | 0.677 | 0.72 | 0.898 | 0.499 |
| TYR | 42 | A | 0.832 | 0.825 | 0.999 | 0.658 | ALA | 115 | A | 0.628 | 0.569 | 0.98 | 0.217 |
| PHE | 43 | A | 0.846 | 0.83 | 0.998 | 0.678 | GLU | 116 | A | 0.778 | 0.783 | 0.955 | 0.606 |
| PRO | 44 | A | 0.727 | 0.725 | 0.995 | 0.457 | PHE | 117 | A | 0.829 | 0.834 | 0.997 | 0.666 |
| HIS | 45 | A | 0.814 | 0.809 | 0.987 | 0.636 | THR | 118 | A | 0.751 | 0.711 | 0.999 | 0.463 |
| PHE | 46 | A | 0.83 | 0.83 | 0.914 | 0.746 | PRO | 119 | A | 0.663 | 0.669 | 1 | 0.332 |
| ASP | 47 | A | 0.779 | 0.739 | 0.937 | 0.581 | ALA | 120 | A | 0.627 | 0.632 | 0.999 | 0.26 |
| LEU | 48 | A | 0.73 | 0.734 | 0.997 | 0.467 | VAL | 121 | A | 0.743 | 0.762 | 0.999 | 0.506 |
| SER | 49 | A | 0.763 | 0.748 | 0.996 | 0.515 | HIS | 122 | A | 0.779 | 0.778 | 0.998 | 0.559 |
| HIS | 50 | A | 0.764 | 0.771 | 0.998 | 0.537 | ALA | 123 | A | 0.577 | 0.592 | 0.999 | 0.17 |
| GLY | 51 | A | 0.65 | 0.653 | 0.999 | 0.304 | SER | 124 | A | 0.675 | 0.701 | 0.998 | 0.378 |
| SER | 52 | A | 0.707 | 0.676 | 0.993 | 0.39 | LEU | 125 | A | 0.736 | 0.701 | 0.998 | 0.439 |
| ALA | 53 | A | 0.639 | 0.634 | 0.999 | 0.274 | ASP | 126 | A | 0.752 | 0.746 | 0.996 | 0.502 |
| GLN | 54 | A | 0.729 | 0.738 | 0.927 | 0.54 | LYS | 127 | A | 0.745 | 0.724 | 0.998 | 0.471 |
| VAL | 55 | A | 0.662 | 0.675 | 0.998 | 0.339 | PHE | 128 | A | 0.816 | 0.827 | 0.999 | 0.644 |
| LYS | 56 | A | 0.775 | 0.802 | 0.998 | 0.579 | LEU | 129 | A | 0.732 | 0.736 | 0.998 | 0.47 |
| GLY | 57 | A | 0.649 | 0.652 | 1 | 0.301 | ALA | 130 | A | 0.585 | 0.584 | 1 | 0.169 |
| HIS | 58 | A | 0.788 | 0.803 | 0.925 | 0.666 | SER | 131 | A | 0.715 | 0.726 | 0.999 | 0.442 |
| GLY | 59 | A | 0.628 | 0.623 | 0.999 | 0.252 | VAL | 132 | A | 0.713 | 0.705 | 0.999 | 0.419 |
| **LYS** | **60** | **A** | **0.817** | **0.759** | **0.691** | **0.885** | SER | 133 | A | 0.712 | 0.712 | 0.998 | 0.426 |
| LYS | 61 | A | 0.734 | 0.767 | 0.994 | 0.507 | THR | 134 | A | 0.727 | 0.72 | 1 | 0.447 |
| VAL | 62 | A | 0.702 | 0.702 | 0.997 | 0.407 | VAL | 135 | A | 0.716 | 0.702 | 0.999 | 0.419 |
| ALA | 63 | A | 0.586 | 0.559 | 0.996 | 0.149 | LEU | 136 | A | 0.757 | 0.748 | 0.999 | 0.506 |
| ASP | 64 | A | 0.822 | 0.801 | 0.999 | 0.624 | THR | 137 | A | 0.672 | 0.684 | 0.999 | 0.357 |
| ALA | 65 | A | 0.598 | 0.598 | 0.997 | 0.199 | SER | 138 | A | 0.617 | 0.642 | 0.999 | 0.26 |
| LEU | 66 | A | 0.721 | 0.728 | 0.973 | 0.476 | LYS | 139 | A | 0.774 | 0.777 | 0.906 | 0.645 |
| **THR** | **67** | **A** | **0.75** | **0.776** | **0.501** | **1.025** | TYR | 140 | A | 0.821 | 0.825 | 0.998 | 0.648 |
| ASN | 68 | A | 0.761 | 0.744 | 0.964 | 0.541 | **ARG** | **141** | **A** | **0.915** | **0.928** | **0.761** | **1.082** |
| ALA | 69 | A | 0.552 | 0.521 | 0.978 | 0.095 | HIS | 2 | B | 0.685 | 0.623 | 0.993 | 0.315 |
| **VAL** | **70** | **A** | **0.726** | **0.706** | **0.598** | **0.834** | LEU | 3 | B | 0.716 | 0.691 | 0.999 | 0.408 |
| ALA | 71 | A | 0.497 | 0.535 | 0.986 | 0.046 | THR | 4 | B | 0.756 | 0.735 | 0.999 | 0.492 |
| **HIS** | **72** | **A** | **0.841** | **0.836** | **0.754** | **0.923** | PRO | 5 | B | 0.757 | 0.752 | 1 | 0.509 |
| VAL | 73 | A | 0.695 | 0.683 | 0.986 | 0.392 | GLU | 6 | B | 0.659 | 0.621 | 0.801 | 0.479 |
| ASP | 74 | A | 0.769 | 0.716 | 0.993 | 0.492 | GLU | 7 | B | 0.837 | 0.807 | 0.954 | 0.69 |

**Table S7(c): 2DN1 (P), 2DN3 (Q),** continued..

| Residue | | | S (P) | S (Q) | J (P,Q) | PRI | Residue | | | S (P) | S (Q) | J (P,Q) | PRI |
|---|---|---|---|---|---|---|---|---|---|---|---|---|---|
| LYS | 8 | B | 0.786 | 0.798 | 0.955 | 0.629 | LEU | 81 | B | 0.727 | 0.736 | 0.997 | 0.466 |
| SER | 9 | B | 0.678 | 0.703 | 0.992 | 0.389 | LYS | 82 | B | 0.727 | 0.784 | 0.989 | 0.522 |
| ALA | 10 | B | 0.557 | 0.593 | 0.976 | 0.174 | GLY | 83 | B | 0.658 | 0.662 | 0.998 | 0.322 |
| VAL | 11 | B | 0.723 | 0.727 | 0.996 | 0.454 | THR | 84 | B | 0.63 | 0.657 | 1 | 0.287 |
| THR | 12 | B | 0.744 | 0.759 | 0.998 | 0.505 | PHE | 85 | B | 0.831 | 0.833 | 0.999 | 0.665 |
| ALA | 13 | B | 0.584 | 0.622 | 1 | 0.206 | ALA | 86 | B | 0.63 | 0.664 | 1 | 0.294 |
| LEU | 14 | B | 0.742 | 0.719 | 0.984 | 0.477 | THR | 87 | B | 0.68 | 0.698 | 1 | 0.378 |
| TRP | 15 | B | 0.865 | 0.856 | 1 | 0.721 | LEU | 88 | B | 0.722 | 0.713 | 1 | 0.435 |
| GLY | 16 | B | 0.654 | 0.648 | 1 | 0.302 | SER | 89 | B | 0.693 | 0.712 | 1 | 0.405 |
| LYS | 17 | B | 0.75 | 0.74 | 0.986 | 0.504 | GLU | 90 | B | 0.793 | 0.801 | 0.999 | 0.595 |
| VAL | 18 | B | 0.65 | 0.658 | 1 | 0.308 | LEU | 91 | B | 0.755 | 0.737 | 1 | 0.492 |
| ASN | 19 | B | 0.765 | 0.754 | 0.999 | 0.52 | HIS | 92 | B | 0.751 | 0.725 | 1 | 0.476 |
| VAL | 20 | B | 0.683 | 0.694 | 0.993 | 0.384 | CYS | 93 | B | 0.661 | 0.65 | 1 | 0.311 |
| ASP | 21 | B | 0.721 | 0.741 | 0.999 | 0.463 | ASP | 94 | B | 0.689 | 0.705 | 0.988 | 0.406 |
| **GLU** | **22** | **B** | **0.806** | **0.825** | **0.618** | **1.013** | LYS | 95 | B | 0.716 | 0.708 | 0.997 | 0.427 |
| VAL | 23 | B | 0.686 | 0.674 | 0.998 | 0.362 | LEU | 96 | B | 0.694 | 0.693 | 0.999 | 0.388 |
| GLY | 24 | B | 0.623 | 0.625 | 1 | 0.248 | HIS | 97 | B | 0.858 | 0.865 | 0.999 | 0.724 |
| GLY | 25 | B | 0.622 | 0.628 | 0.999 | 0.251 | VAL | 98 | B | 0.691 | 0.699 | 1 | 0.39 |
| GLU | 26 | B | 0.774 | 0.773 | 1 | 0.547 | ASP | 99 | B | 0.79 | 0.802 | 1 | 0.592 |
| ALA | 27 | B | 0.552 | 0.566 | 1 | 0.118 | PRO | 100 | B | 0.682 | 0.679 | 1 | 0.361 |
| LEU | 28 | B | 0.716 | 0.707 | 0.999 | 0.424 | GLU | 101 | B | 0.731 | 0.767 | 1 | 0.498 |
| GLY | 29 | B | 0.628 | 0.626 | 1 | 0.254 | ASN | 102 | B | 0.741 | 0.753 | 1 | 0.494 |
| ARG | 30 | B | 0.837 | 0.84 | 1 | 0.677 | PHE | 103 | B | 0.84 | 0.841 | 1 | 0.681 |
| LEU | 31 | B | 0.726 | 0.725 | 0.999 | 0.452 | ARG | 104 | B | 0.783 | 0.777 | 1 | 0.56 |
| LEU | 32 | B | 0.74 | 0.744 | 1 | 0.484 | LEU | 105 | B | 0.75 | 0.756 | 1 | 0.506 |
| VAL | 33 | B | 0.689 | 0.697 | 1 | 0.386 | LEU | 106 | B | 0.721 | 0.715 | 1 | 0.436 |
| VAL | 34 | B | 0.697 | 0.702 | 0.999 | 0.4 | GLY | 107 | B | 0.626 | 0.632 | 1 | 0.258 |
| TYR | 35 | B | 0.873 | 0.871 | 0.998 | 0.746 | ASN | 108 | B | 0.78 | 0.765 | 0.998 | 0.547 |
| PRO | 36 | B | 0.695 | 0.681 | 0.998 | 0.378 | VAL | 109 | B | 0.727 | 0.728 | 0.999 | 0.456 |
| TRP | 37 | B | 0.867 | 0.87 | 0.999 | 0.738 | LEU | 110 | B | 0.738 | 0.731 | 1 | 0.469 |
| THR | 38 | B | 0.687 | 0.695 | 0.999 | 0.383 | VAL | 111 | B | 0.714 | 0.71 | 0.999 | 0.425 |
| GLN | 39 | B | 0.777 | 0.778 | 0.998 | 0.557 | CYS | 112 | B | 0.606 | 0.607 | 0.999 | 0.214 |
| ARG | 40 | B | 0.765 | 0.754 | 0.987 | 0.532 | VAL | 113 | B | 0.698 | 0.7 | 1 | 0.398 |
| PHE | 41 | B | 0.827 | 0.818 | 0.998 | 0.647 | LEU | 114 | B | 0.74 | 0.746 | 0.999 | 0.487 |
| PHE | 42 | B | 0.831 | 0.825 | 0.999 | 0.657 | ALA | 115 | B | 0.538 | 0.556 | 0.999 | 0.095 |
| GLU | 43 | B | 0.837 | 0.804 | 0.959 | 0.682 | HIS | 116 | B | 0.775 | 0.782 | 0.945 | 0.612 |
| SER | 44 | B | 0.632 | 0.633 | 0.99 | 0.275 | **HIS** | **117** | **B** | **0.751** | **0.818** | **0.76** | **0.809** |
| PHE | 45 | B | 0.828 | 0.833 | 0.998 | 0.663 | PHE | 118 | B | 0.849 | 0.849 | 1 | 0.698 |
| GLY | 46 | B | 0.653 | 0.657 | 1 | 0.31 | GLY | 119 | B | 0.627 | 0.634 | 0.999 | 0.262 |
| ASP | 47 | B | 0.774 | 0.763 | 0.999 | 0.538 | LYS | 120 | B | 0.593 | 0.63 | 0.936 | 0.287 |
| LEU | 48 | B | 0.743 | 0.735 | 0.999 | 0.479 | GLU | 121 | B | 0.74 | 0.716 | 0.899 | 0.557 |
| SER | 49 | B | 0.699 | 0.705 | 0.999 | 0.405 | PHE | 122 | B | 0.843 | 0.843 | 1 | 0.686 |
| THR | 50 | B | 0.801 | 0.806 | 1 | 0.607 | THR | 123 | B | 0.768 | 0.77 | 1 | 0.538 |
| PRO | 51 | B | 0.683 | 0.677 | 0.999 | 0.361 | PRO | 124 | B | 0.673 | 0.683 | 1 | 0.356 |
| ASP | 52 | B | 0.614 | 0.608 | 0.999 | 0.223 | PRO | 125 | B | 0.717 | 0.72 | 1 | 0.437 |
| ALA | 53 | B | 0.55 | 0.54 | 1 | 0.09 | VAL | 126 | B | 0.677 | 0.678 | 1 | 0.355 |
| VAL | 54 | B | 0.703 | 0.707 | 1 | 0.41 | GLN | 127 | B | 0.766 | 0.766 | 1 | 0.532 |
| MET | 55 | B | 0.739 | 0.735 | 0.999 | 0.475 | ALA | 128 | B | 0.57 | 0.55 | 1 | 0.12 |
| GLY | 56 | B | 0.639 | 0.661 | 1 | 0.3 | ALA | 129 | B | 0.561 | 0.543 | 1 | 0.104 |
| ASN | 57 | B | 0.743 | 0.731 | 0.999 | 0.475 | TYR | 130 | B | 0.836 | 0.835 | 1 | 0.671 |
| PRO | 58 | B | 0.735 | 0.727 | 0.999 | 0.463 | GLN | 131 | B | 0.772 | 0.774 | 1 | 0.546 |
| LYS | 59 | B | 0.747 | 0.742 | 1 | 0.489 | LYS | 132 | B | 0.776 | 0.791 | 0.999 | 0.568 |
| VAL | 60 | B | 0.668 | 0.672 | 1 | 0.34 | VAL | 133 | B | 0.716 | 0.701 | 1 | 0.417 |
| LYS | 61 | B | 0.708 | 0.753 | 0.992 | 0.469 | VAL | 134 | B | 0.691 | 0.688 | 1 | 0.379 |
| ALA | 62 | B | 0.651 | 0.656 | 0.999 | 0.308 | ALA | 135 | B | 0.582 | 0.619 | 1 | 0.201 |
| HIS | 63 | B | 0.773 | 0.785 | 0.984 | 0.574 | GLY | 136 | B | 0.638 | 0.643 | 1 | 0.281 |
| GLY | 64 | B | 0.631 | 0.619 | 1 | 0.25 | VAL | 137 | B | 0.705 | 0.704 | 1 | 0.409 |
| LYS | 65 | B | 0.755 | 0.769 | 0.999 | 0.525 | ALA | 138 | B | 0.593 | 0.572 | 1 | 0.165 |
| LYS | 66 | B | 0.759 | 0.749 | 0.953 | 0.555 | ASN | 139 | B | 0.764 | 0.754 | 0.805 | 0.713 |
| VAL | 67 | B | 0.701 | 0.693 | 0.999 | 0.395 | ALA | 140 | B | 0.539 | 0.541 | 1 | 0.08 |
| LEU | 68 | B | 0.731 | 0.731 | 0.999 | 0.463 | LEU | 141 | B | 0.74 | 0.738 | 1 | 0.478 |
| GLY | 69 | B | 0.647 | 0.648 | 0.999 | 0.296 | ALA | 142 | B | 0.472 | 0.485 | 1 | -0.043 |
| ALA | 70 | B | 0.557 | 0.569 | 0.998 | 0.128 | **HIS** | **143** | **B** | **0.805** | **0.804** | **0.846** | **0.763** |
| PHE | 71 | B | 0.827 | 0.826 | 0.999 | 0.654 | LYS | 144 | B | 0.718 | 0.743 | 0.989 | 0.472 |
| SER | 72 | B | 0.709 | 0.719 | 0.999 | 0.429 | TYR | 145 | B | 0.832 | 0.825 | 0.999 | 0.658 |
| ASP | 73 | B | 0.765 | 0.786 | 0.998 | 0.553 | **HIS** | **146** | **B** | **0.91** | **0.911** | **0.895** | **0.926** |
| GLY | 74 | B | 0.629 | 0.63 | 0.999 | 0.26 | | | | | | | |
| LEU | 75 | B | 0.741 | 0.754 | 0.999 | 0.496 | | | | | | | |
| ALA | 76 | B | 0.454 | 0.477 | 0.999 | -0.068 | | | | | | | |
| HIS | 77 | B | 0.817 | 0.81 | 1 | 0.627 | | | | | | | |
| LEU | 78 | B | 0.724 | 0.744 | 0.999 | 0.469 | | | | | | | |
| ASP | 79 | B | 0.683 | 0.706 | 0.933 | 0.456 | | | | | | | |
| ASN | 80 | B | 0.748 | 0.762 | 0.992 | 0.518 | | | | | | | |

| Table S7(d): 2DN2 (P), 2DN3 (Q) | | | | | | | | | | | | | | |
|---|---|---|---|---|---|---|---|---|---|---|---|---|---|---|
| Residue | | | S (P) | S (Q) | J (P,Q) | PRI | | Residue | | | S (P) | S (Q) | J (P,Q) | PRI |
| VAL | 1 | A | 0.725 | 0.696 | 0.973 | 0.448 | | ASP | 74 | A | 0.716 | 0.732 | 0.98 | 0.468 |
| LEU | 2 | A | 0.693 | 0.69 | 0.997 | 0.386 | | ASP | 75 | A | 0.799 | 0.781 | 0.98 | 0.6 |
| SER | 3 | A | 0.74 | 0.73 | 1 | 0.47 | | MET | 76 | A | 0.722 | 0.731 | 0.99 | 0.463 |
| PRO | 4 | A | 0.754 | 0.747 | 1 | 0.501 | | PRO | 77 | A | 0.708 | 0.69 | 0.998 | 0.4 |
| ALA | 5 | A | 0.64 | 0.629 | 1 | 0.269 | | ASN | 78 | A | 0.699 | 0.621 | 0.994 | 0.326 |
| ASP | 6 | A | 0.805 | 0.769 | 1 | 0.574 | | ALA | 79 | A | 0.481 | 0.483 | 1 | -0.036 |
| LYS | 7 | A | 0.763 | 0.79 | 0.999 | 0.554 | | LEU | 80 | A | 0.731 | 0.708 | 0.996 | 0.443 |
| THR | 8 | A | 0.758 | 0.754 | 1 | 0.512 | | SER | 81 | A | 0.722 | 0.752 | 0.997 | 0.477 |
| ASN | 9 | A | 0.805 | 0.763 | 0.999 | 0.569 | | ALA | 82 | A | 0.678 | 0.679 | 0.999 | 0.358 |
| VAL | 10 | A | 0.71 | 0.684 | 0.998 | 0.396 | | LEU | 83 | A | 0.773 | 0.691 | 0.995 | 0.469 |
| LYS | 11 | A | 0.758 | 0.809 | 0.922 | 0.645 | | SER | 84 | A | 0.623 | 0.667 | 0.972 | 0.318 |
| ALA | 12 | A | 0.623 | 0.637 | 0.999 | 0.261 | | ASP | 85 | A | 0.796 | 0.785 | 1 | 0.581 |
| ALA | 13 | A | 0.592 | 0.584 | 0.998 | 0.178 | | LEU | 86 | A | 0.73 | 0.767 | 0.991 | 0.506 |
| TRP | 14 | A | 0.838 | 0.848 | 0.995 | 0.691 | | HIS | 87 | A | 0.758 | 0.815 | 0.99 | 0.583 |
| GLY | 15 | A | 0.641 | 0.656 | 0.999 | 0.298 | | ALA | 88 | A | 0.548 | 0.524 | 0.992 | 0.08 |
| **LYS** | **16** | **A** | **0.772** | **0.755** | **0.768** | **0.759** | | HIS | 89 | A | 0.842 | 0.803 | 0.975 | 0.67 |
| VAL | 17 | A | 0.719 | 0.688 | 0.998 | 0.409 | | LYS | 90 | A | 0.784 | 0.675 | 0.813 | 0.646 |
| GLY | 18 | A | 0.651 | 0.653 | 0.999 | 0.305 | | LEU | 91 | A | 0.739 | 0.686 | 1 | 0.425 |
| ALA | 19 | A | 0.55 | 0.544 | 0.994 | 0.1 | | ARG | 92 | A | 0.756 | 0.852 | 0.947 | 0.661 |
| HIS | 20 | A | 0.801 | 0.77 | 0.994 | 0.577 | | VAL | 93 | A | 0.692 | 0.677 | 0.999 | 0.37 |
| ALA | 21 | A | 0.671 | 0.536 | 0.995 | 0.212 | | ASP | 94 | A | 0.642 | 0.804 | 0.941 | 0.505 |
| GLY | 22 | A | 0.626 | 0.62 | 0.999 | 0.247 | | PRO | 95 | A | 0.706 | 0.682 | 0.993 | 0.395 |
| GLU | 23 | A | 0.774 | 0.656 | 0.933 | 0.497 | | VAL | 96 | A | 0.717 | 0.746 | 0.924 | 0.539 |
| **TYR** | **24** | **A** | **0.869** | **0.855** | **0.999** | **0.725** | | ASN | 97 | A | 0.751 | 0.745 | 0.995 | 0.501 |
| GLY | 25 | A | 0.634 | 0.635 | 0.999 | 0.27 | | PHE | 98 | A | 0.817 | 0.822 | 1 | 0.639 |
| ALA | 26 | A | 0.545 | 0.571 | 1 | 0.116 | | LYS | 99 | A | 0.721 | 0.746 | 0.976 | 0.491 |
| GLU | 27 | A | 0.775 | 0.772 | 0.913 | 0.634 | | LEU | 100 | A | 0.74 | 0.736 | 0.999 | 0.477 |
| ALA | 28 | A | 0.556 | 0.536 | 1 | 0.092 | | LEU | 101 | A | 0.7 | 0.66 | 1 | 0.36 |
| LEU | 29 | A | 0.707 | 0.715 | 1 | 0.422 | | SER | 102 | A | 0.704 | 0.72 | 0.999 | 0.425 |
| GLU | 30 | A | 0.782 | 0.782 | 0.989 | 0.575 | | HIS | 103 | A | 0.804 | 0.783 | 0.999 | 0.588 |
| **ARG** | **31** | **A** | **0.867** | **0.864** | **0.998** | **0.733** | | CYS | 104 | A | 0.613 | 0.609 | 0.999 | 0.223 |
| MET | 32 | A | 0.759 | 0.758 | 0.999 | 0.518 | | LEU | 105 | A | 0.769 | 0.746 | 0.997 | 0.518 |
| PHE | 33 | A | 0.848 | 0.841 | 1 | 0.689 | | LEU | 106 | A | 0.736 | 0.746 | 0.999 | 0.483 |
| LEU | 34 | A | 0.65 | 0.679 | 1 | 0.329 | | VAL | 107 | A | 0.713 | 0.702 | 1 | 0.415 |
| SER | 35 | A | 0.627 | 0.615 | 1 | 0.242 | | THR | 108 | A | 0.691 | 0.686 | 0.999 | 0.378 |
| PHE | 36 | A | 0.84 | 0.835 | 1 | 0.675 | | LEU | 109 | A | 0.72 | 0.739 | 1 | 0.459 |
| PRO | 37 | A | 0.701 | 0.692 | 1 | 0.393 | | ALA | 110 | A | 0.511 | 0.528 | 1 | 0.039 |
| THR | 38 | A | 0.643 | 0.686 | 0.999 | 0.33 | | ALA | 111 | A | 0.459 | 0.443 | 1 | -0.098 |
| THR | 39 | A | 0.687 | 0.695 | 1 | 0.382 | | HIS | 112 | A | 0.822 | 0.817 | 0.999 | 0.64 |
| LYS | 40 | A | 0.773 | 0.769 | 1 | 0.542 | | LEU | 113 | A | 0.753 | 0.728 | 0.944 | 0.537 |
| THR | 41 | A | 0.73 | 0.744 | 0.999 | 0.475 | | PRO | 114 | A | 0.72 | 0.733 | 0.987 | 0.466 |
| TYR | 42 | A | 0.824 | 0.853 | 0.998 | 0.679 | | ALA | 115 | A | 0.57 | 0.577 | 0.999 | 0.148 |
| PHE | 43 | A | 0.83 | 0.836 | 1 | 0.666 | | GLU | 116 | A | 0.783 | 0.772 | 0.994 | 0.561 |
| PRO | 44 | A | 0.725 | 0.713 | 1 | 0.438 | | PHE | 117 | A | 0.834 | 0.815 | 0.998 | 0.651 |
| HIS | 45 | A | 0.809 | 0.824 | 1 | 0.633 | | THR | 118 | A | 0.711 | 0.711 | 1 | 0.422 |
| PHE | 46 | A | 0.83 | 0.825 | 0.995 | 0.66 | | PRO | 119 | A | 0.668 | 0.659 | 1 | 0.327 |
| ASP | 47 | A | 0.739 | 0.807 | 0.981 | 0.565 | | ALA | 120 | A | 0.632 | 0.611 | 0.999 | 0.244 |
| LEU | 48 | A | 0.734 | 0.724 | 0.995 | 0.463 | | VAL | 121 | A | 0.762 | 0.712 | 1 | 0.474 |
| SER | 49 | A | 0.748 | 0.748 | 0.994 | 0.502 | | HIS | 122 | A | 0.778 | 0.788 | 1 | 0.566 |
| HIS | 50 | A | 0.771 | 0.758 | 0.997 | 0.532 | | ALA | 123 | A | 0.592 | 0.579 | 0.999 | 0.172 |
| GLY | 51 | A | 0.653 | 0.645 | 1 | 0.298 | | SER | 124 | A | 0.701 | 0.675 | 0.999 | 0.377 |
| SER | 52 | A | 0.676 | 0.661 | 0.97 | 0.367 | | LEU | 125 | A | 0.701 | 0.678 | 0.999 | 0.38 |
| ALA | 53 | A | 0.634 | 0.664 | 0.997 | 0.301 | | ASP | 126 | A | 0.746 | 0.758 | 0.917 | 0.587 |
| GLN | 54 | A | 0.738 | 0.727 | 0.999 | 0.466 | | **LYS** | **127** | **A** | **0.721** | **0.795** | **0.716** | **0.8** |
| VAL | 55 | A | 0.675 | 0.684 | 0.999 | 0.36 | | PHE | 128 | A | 0.827 | 0.828 | 0.999 | 0.656 |
| LYS | 56 | A | 0.802 | 0.787 | 1 | 0.589 | | LEU | 129 | A | 0.736 | 0.714 | 0.998 | 0.452 |
| GLY | 57 | A | 0.652 | 0.653 | 1 | 0.305 | | ALA | 130 | A | 0.583 | 0.592 | 0.999 | 0.176 |
| HIS | 58 | A | 0.803 | 0.778 | 0.997 | 0.584 | | SER | 131 | A | 0.723 | 0.718 | 0.999 | 0.442 |
| GLY | 59 | A | 0.623 | 0.629 | 1 | 0.252 | | VAL | 132 | A | 0.705 | 0.707 | 0.998 | 0.414 |
| LYS | 60 | A | 0.759 | 0.794 | 0.965 | 0.588 | | SER | 133 | A | 0.712 | 0.711 | 0.999 | 0.424 |
| LYS | 61 | A | 0.767 | 0.7 | 0.995 | 0.472 | | THR | 134 | A | 0.718 | 0.745 | 0.999 | 0.464 |
| VAL | 62 | A | 0.702 | 0.698 | 0.999 | 0.401 | | VAL | 135 | A | 0.702 | 0.695 | 0.996 | 0.401 |
| ALA | 63 | A | 0.559 | 0.587 | 0.999 | 0.147 | | LEU | 136 | A | 0.748 | 0.737 | 0.994 | 0.491 |
| ASP | 64 | A | 0.801 | 0.794 | 0.999 | 0.596 | | THR | 137 | A | 0.684 | 0.68 | 0.998 | 0.366 |
| ALA | 65 | A | 0.598 | 0.574 | 0.998 | 0.174 | | SER | 138 | A | 0.642 | 0.646 | 0.998 | 0.29 |
| LEU | 66 | A | 0.728 | 0.716 | 0.995 | 0.449 | | LYS | 139 | A | 0.777 | 0.783 | 0.988 | 0.572 |
| THR | 67 | A | 0.776 | 0.701 | 0.893 | 0.584 | | TYR | 140 | A | 0.825 | 0.797 | 0.966 | 0.656 |
| ASN | 68 | A | 0.744 | 0.733 | 0.999 | 0.478 | | **ARG** | **141** | **A** | **0.928** | **0.661** | **0.809** | **0.78** |
| ALA | 69 | A | 0.521 | 0.543 | 1 | 0.064 | | VAL | 1 | B | 0.802 | 0.821 | 0.953 | 0.67 |
| VAL | 70 | A | 0.706 | 0.676 | 0.995 | 0.387 | | **HIS** | **2** | **B** | **0.82** | **0.792** | **0.818** | **0.794** |
| ALA | 71 | A | 0.535 | 0.534 | 1 | 0.069 | | LEU | 3 | B | 0.692 | 0.738 | 0.994 | 0.436 |
| HIS | 72 | A | 0.836 | 0.791 | 0.999 | 0.628 | | THR | 4 | B | 0.732 | 0.711 | 1 | 0.443 |
| VAL | 73 | A | 0.682 | 0.673 | 0.998 | 0.357 | | PRO | 5 | B | 0.75 | 0.746 | 0.999 | 0.497 |

**Table S7(d): 2DN2 (P), 2DN3 (Q),** continued..

| **Residue** | | | **S (P)** | **S (Q)** | **J (P,Q)** | **PRI** |
|---|---|---|---|---|---|---|
| GLU | 6 | B | 0.62 | 0.836 | 0.943 | 0.513 |
| GLU | 7 | B | 0.806 | 0.83 | 0.999 | 0.637 |
| LYS | 8 | B | 0.797 | 0.686 | 0.998 | 0.485 |
| SER | 9 | B | 0.703 | 0.68 | 0.998 | 0.385 |
| ALA | 10 | B | 0.592 | 0.591 | 0.999 | 0.184 |
| VAL | 11 | B | 0.726 | 0.722 | 0.999 | 0.449 |
| THR | 12 | B | 0.759 | 0.714 | 0.911 | 0.562 |
| ALA | 13 | B | 0.622 | 0.624 | 0.999 | 0.247 |
| LEU | 14 | B | 0.719 | 0.717 | 1 | 0.436 |
| **TRP** | **15** | **B** | **0.856** | **0.865** | **0.999** | **0.722** |
| GLY | 16 | B | 0.648 | 0.645 | 1 | 0.293 |
| LYS | 17 | B | 0.74 | 0.703 | 0.99 | 0.453 |
| VAL | 18 | B | 0.658 | 0.657 | 0.999 | 0.316 |
| ASN | 19 | B | 0.754 | 0.781 | 0.996 | 0.539 |
| VAL | 20 | B | 0.694 | 0.653 | 0.902 | 0.445 |
| ASP | 21 | B | 0.741 | 0.786 | 0.97 | 0.557 |
| **GLU** | **22** | **B** | **0.825** | **0.791** | **0.851** | **0.765** |
| VAL | 23 | B | 0.674 | 0.683 | 0.999 | 0.358 |
| GLY | 24 | B | 0.625 | 0.624 | 0.997 | 0.252 |
| GLY | 25 | B | 0.627 | 0.634 | 0.998 | 0.263 |
| GLU | 26 | B | 0.773 | 0.748 | 0.886 | 0.635 |
| ALA | 27 | B | 0.566 | 0.564 | 1 | 0.13 |
| LEU | 28 | B | 0.707 | 0.701 | 1 | 0.408 |
| GLY | 29 | B | 0.626 | 0.627 | 1 | 0.253 |
| ARG | 30 | B | 0.84 | 0.839 | 0.999 | 0.68 |
| LEU | 31 | B | 0.725 | 0.72 | 0.999 | 0.446 |
| LEU | 32 | B | 0.744 | 0.731 | 0.923 | 0.552 |
| VAL | 33 | B | 0.697 | 0.697 | 0.999 | 0.395 |
| VAL | 34 | B | 0.702 | 0.699 | 1 | 0.401 |
| **TYR** | **35** | **B** | **0.871** | **0.869** | **1** | **0.74** |
| PRO | 36 | B | 0.68 | 0.69 | 1 | 0.37 |
| **TRP** | **37** | **B** | **0.869** | **0.884** | **0.993** | **0.76** |
| THR | 38 | B | 0.696 | 0.712 | 0.999 | 0.409 |
| GLN | 39 | B | 0.778 | 0.786 | 0.998 | 0.566 |
| ARG | 40 | B | 0.754 | 0.693 | 0.916 | 0.531 |
| PHE | 41 | B | 0.818 | 0.834 | 0.995 | 0.657 |
| PHE | 42 | B | 0.825 | 0.846 | 0.999 | 0.672 |
| GLU | 43 | B | 0.804 | 0.686 | 0.838 | 0.652 |
| SER | 44 | B | 0.633 | 0.617 | 1 | 0.25 |
| PHE | 45 | B | 0.833 | 0.844 | 0.999 | 0.678 |
| GLY | 46 | B | 0.657 | 0.643 | 1 | 0.3 |
| ASP | 47 | B | 0.763 | 0.751 | 1 | 0.514 |
| LEU | 48 | B | 0.735 | 0.734 | 0.999 | 0.47 |
| SER | 49 | B | 0.705 | 0.656 | 1 | 0.361 |
| THR | 50 | B | 0.806 | 0.787 | 1 | 0.593 |
| PRO | 51 | B | 0.677 | 0.676 | 1 | 0.353 |
| ASP | 52 | B | 0.608 | 0.63 | 1 | 0.238 |
| ALA | 53 | B | 0.54 | 0.579 | 1 | 0.119 |
| VAL | 54 | B | 0.707 | 0.707 | 1 | 0.414 |
| MET | 55 | B | 0.735 | 0.709 | 0.999 | 0.445 |
| GLY | 56 | B | 0.66 | 0.642 | 1 | 0.302 |
| ASN | 57 | B | 0.731 | 0.745 | 1 | 0.476 |
| PRO | 58 | B | 0.727 | 0.727 | 1 | 0.454 |
| LYS | 59 | B | 0.742 | 0.704 | 0.971 | 0.475 |
| VAL | 60 | B | 0.672 | 0.681 | 0.999 | 0.354 |
| LYS | 61 | B | 0.753 | 0.757 | 0.866 | 0.644 |
| ALA | 62 | B | 0.656 | 0.597 | 1 | 0.253 |
| HIS | 63 | B | 0.785 | 0.785 | 0.992 | 0.578 |
| GLY | 64 | B | 0.619 | 0.627 | 1 | 0.246 |
| LYS | 65 | B | 0.769 | 0.78 | 0.841 | 0.708 |
| LYS | 66 | B | 0.749 | 0.722 | 0.996 | 0.475 |
| VAL | 67 | B | 0.693 | 0.73 | 0.996 | 0.427 |
| LEU | 68 | B | 0.731 | 0.726 | 0.8 | 0.657 |
| GLY | 69 | B | 0.648 | 0.64 | 0.992 | 0.296 |
| ALA | 70 | B | 0.569 | 0.575 | 0.985 | 0.159 |
| PHE | 71 | B | 0.826 | 0.824 | 0.934 | 0.716 |
| SER | 72 | B | 0.719 | 0.704 | 0.985 | 0.438 |
| ASP | 73 | B | 0.786 | 0.788 | 0.939 | 0.635 |
| GLY | 74 | B | 0.63 | 0.633 | 0.992 | 0.271 |
| LEU | 75 | B | 0.753 | 0.74 | 0.991 | 0.502 |
| ALA | 76 | B | 0.477 | 0.518 | 0.997 | -0.002 |
| HIS | 77 | B | 0.81 | 0.84 | 0.999 | 0.651 |
| LEU | 78 | B | 0.745 | 0.711 | 1 | 0.456 |
| ASP | 79 | B | 0.698 | 0.698 | 0.89 | 0.506 |
| ASN | 80 | B | 0.761 | 0.751 | 0.991 | 0.521 |
| LEU | 81 | B | 0.734 | 0.721 | 1 | 0.455 |
| **LYS** | **82** | **B** | **0.784** | **0.733** | **0.784** | **0.733** |
| GLY | 83 | B | 0.662 | 0.65 | 0.992 | 0.32 |
| THR | 84 | B | 0.657 | 0.564 | 0.994 | 0.227 |
| PHE | 85 | B | 0.832 | 0.842 | 0.985 | 0.689 |
| ALA | 86 | B | 0.664 | 0.626 | 0.998 | 0.292 |
| THR | 87 | B | 0.698 | 0.693 | 0.875 | 0.516 |
| LEU | 88 | B | 0.713 | 0.702 | 0.997 | 0.418 |
| SER | 89 | B | 0.712 | 0.72 | 0.997 | 0.435 |
| GLU | 90 | B | 0.801 | 0.651 | 0.993 | 0.459 |
| LEU | 91 | B | 0.737 | 0.752 | 0.972 | 0.517 |
| HIS | 92 | B | 0.725 | 0.794 | 0.994 | 0.525 |
| CYS | 93 | B | 0.65 | 0.642 | 0.855 | 0.437 |
| ASP | 94 | B | 0.705 | 0.768 | 0.97 | 0.503 |
| LYS | 95 | B | 0.707 | 0.741 | 0.98 | 0.468 |
| LEU | 96 | B | 0.693 | 0.742 | 0.996 | 0.439 |
| **HIS** | **97** | **B** | **0.864** | **0.866** | **0.999** | **0.731** |
| VAL | 98 | B | 0.699 | 0.682 | 1 | 0.381 |
| ASP | 99 | B | 0.802 | 0.738 | 0.997 | 0.543 |
| PRO | 100 | B | 0.678 | 0.671 | 0.982 | 0.367 |
| GLU | 101 | B | 0.767 | 0.752 | 0.953 | 0.566 |
| ASN | 102 | B | 0.753 | 0.777 | 0.968 | 0.562 |
| PHE | 103 | B | 0.841 | 0.838 | 0.99 | 0.689 |
| **ARG** | **104** | **B** | **0.777** | **0.788** | **0.726** | **0.839** |
| LEU | 105 | B | 0.756 | 0.722 | 0.987 | 0.491 |
| LEU | 106 | B | 0.715 | 0.714 | 0.997 | 0.432 |
| GLY | 107 | B | 0.632 | 0.634 | 1 | 0.266 |
| ASN | 108 | B | 0.765 | 0.716 | 0.988 | 0.493 |
| VAL | 109 | B | 0.728 | 0.703 | 0.999 | 0.432 |
| LEU | 110 | B | 0.731 | 0.728 | 1 | 0.459 |
| VAL | 111 | B | 0.71 | 0.697 | 0.999 | 0.408 |
| CYS | 112 | B | 0.607 | 0.599 | 0.999 | 0.207 |
| VAL | 113 | B | 0.7 | 0.703 | 1 | 0.403 |
| LEU | 114 | B | 0.745 | 0.707 | 1 | 0.452 |
| ALA | 115 | B | 0.556 | 0.537 | 1 | 0.093 |
| HIS | 116 | B | 0.782 | 0.785 | 0.999 | 0.568 |
| HIS | 117 | B | 0.818 | 0.757 | 0.873 | 0.702 |
| PHE | 118 | B | 0.849 | 0.846 | 1 | 0.695 |
| GLY | 119 | B | 0.634 | 0.628 | 1 | 0.262 |
| LYS | 120 | B | 0.63 | 0.649 | 0.712 | 0.567 |
| GLU | 121 | B | 0.716 | 0.638 | 0.955 | 0.399 |
| PHE | 122 | B | 0.843 | 0.842 | 1 | 0.685 |
| THR | 123 | B | 0.77 | 0.766 | 1 | 0.536 |
| PRO | 124 | B | 0.683 | 0.663 | 1 | 0.346 |
| PRO | 125 | B | 0.72 | 0.704 | 1 | 0.424 |
| VAL | 126 | B | 0.678 | 0.669 | 1 | 0.347 |
| GLN | 127 | B | 0.766 | 0.777 | 1 | 0.543 |
| ALA | 128 | B | 0.55 | 0.565 | 1 | 0.115 |
| ALA | 129 | B | 0.543 | 0.559 | 1 | 0.102 |
| TYR | 130 | B | 0.835 | 0.837 | 1 | 0.672 |
| GLN | 131 | B | 0.774 | 0.755 | 1 | 0.529 |
| LYS | 132 | B | 0.79 | 0.798 | 1 | 0.588 |
| VAL | 133 | B | 0.702 | 0.703 | 1 | 0.405 |
| VAL | 134 | B | 0.689 | 0.709 | 0.999 | 0.399 |
| ALA | 135 | B | 0.614 | 0.605 | 0.999 | 0.22 |
| GLY | 136 | B | 0.639 | 0.634 | 1 | 0.273 |
| VAL | 137 | B | 0.704 | 0.699 | 1 | 0.403 |
| ALA | 138 | B | 0.572 | 0.545 | 0.999 | 0.118 |
| ASN | 139 | B | 0.75 | 0.807 | 0.876 | 0.681 |
| ALA | 140 | B | 0.54 | 0.515 | 1 | 0.055 |
| LEU | 141 | B | 0.738 | 0.73 | 0.883 | 0.585 |
| ALA | 142 | B | 0.484 | 0.474 | 0.999 | -0.041 |
| HIS | 143 | B | 0.804 | 0.825 | 0.965 | 0.664 |
| LYS | 144 | B | 0.743 | 0.792 | 0.962 | 0.573 |
| TYR | 145 | B | 0.825 | 0.851 | 0.987 | 0.689 |
| **HIS** | **146** | **B** | **0.911** | **0.85** | **0.88** | **0.881** |

**Table S8(a): 1MDN (P), 1M6C (Q)**

| Residue | | | S (P) | S (Q) | J (P,Q) | PRI |
|---|---|---|---|---|---|---|
| GLY | 1 | A | 0.708 | 0.556 | 0.997 | 0.267 |
| LEU | 2 | A | 0.705 | 0.71 | 0.999 | 0.416 |
| SER | 3 | A | 0.735 | 0.743 | 0.999 | 0.479 |
| ASP | 4 | A | 0.603 | 0.615 | 1 | 0.218 |
| GLY | 5 | A | 0.632 | 0.625 | 1 | 0.257 |
| GLU | 6 | A | 0.82 | 0.823 | 0.999 | 0.644 |
| TRP | 7 | A | 0.886 | 0.885 | 0.999 | 0.772 |
| **GLN** | **8** | **A** | **0.831** | **0.857** | **0.805** | **0.883** |
| LEU | 9 | A | 0.675 | 0.662 | 0.999 | 0.338 |
| VAL | 10 | A | 0.713 | 0.72 | 1 | 0.433 |
| LEU | 11 | A | 0.706 | 0.717 | 0.999 | 0.424 |
| ASN | 12 | A | 0.829 | 0.808 | 0.999 | 0.638 |
| VAL | 13 | A | 0.488 | 0.495 | 1 | -0.017 |
| TRP | 14 | A | 0.884 | 0.877 | 0.999 | 0.762 |
| GLY | 15 | A | 0.637 | 0.636 | 0.998 | 0.275 |
| LYS | 16 | A | 0.762 | 0.753 | 0.795 | 0.72 |
| VAL | 17 | A | 0.681 | 0.679 | 0.999 | 0.361 |
| GLU | 18 | A | 0.761 | 0.77 | 0.733 | 0.798 |
| ALA | 19 | A | 0.499 | 0.428 | 0.988 | -0.061 |
| ASP | 20 | A | 0.73 | 0.727 | 1 | 0.457 |
| VAL | 21 | A | 0.726 | 0.728 | 0.999 | 0.455 |
| ALA | 22 | A | 0.575 | 0.571 | 1 | 0.146 |
| GLY | 23 | A | 0.635 | 0.637 | 1 | 0.272 |
| HIS | 24 | A | 0.78 | 0.779 | 1 | 0.559 |
| GLY | 25 | A | 0.632 | 0.632 | 0.999 | 0.265 |
| GLN | 26 | A | 0.7 | 0.737 | 0.987 | 0.45 |
| GLU | 27 | A | 0.778 | 0.795 | 0.999 | 0.574 |
| VAL | 28 | A | 0.713 | 0.725 | 0.999 | 0.439 |
| LEU | 29 | A | 0.679 | 0.714 | 0.995 | 0.398 |
| ILE | 30 | A | 0.752 | 0.752 | 1 | 0.504 |
| ARG | 31 | A | 0.794 | 0.813 | 0.998 | 0.609 |
| LEU | 32 | A | 0.715 | 0.705 | 0.999 | 0.421 |
| PHE | 33 | A | 0.836 | 0.845 | 0.999 | 0.682 |
| LYS | 34 | A | 0.705 | 0.732 | 0.658 | 0.779 |
| GLY | 35 | A | 0.634 | 0.641 | 1 | 0.275 |
| HIS | 36 | A | 0.849 | 0.847 | 0.999 | 0.697 |
| PRO | 37 | A | 0.716 | 0.707 | 1 | 0.423 |
| GLU | 38 | A | 0.765 | 0.764 | 1 | 0.529 |
| THR | 39 | A | 0.688 | 0.666 | 0.999 | 0.355 |
| LEU | 40 | A | 0.721 | 0.719 | 1 | 0.44 |
| **GLU** | **41** | **A** | **0.762** | **0.823** | **0.755** | **0.83** |
| LYS | 42 | A | 0.721 | 0.741 | 0.997 | 0.465 |
| PHE | 43 | A | 0.851 | 0.821 | 1 | 0.672 |
| ASP | 44 | A | 0.653 | 0.626 | 0.975 | 0.304 |
| LYS | 45 | A | 0.694 | 0.689 | 0.589 | 0.794 |
| PHE | 46 | A | 0.857 | 0.848 | 0.998 | 0.707 |
| **LYS** | **47** | **A** | **0.827** | **0.817** | **0.618** | **1.026** |
| HIS | 48 | A | 0.828 | 0.807 | 0.915 | 0.72 |
| LEU | 49 | A | 0.685 | 0.738 | 0.963 | 0.46 |
| LYS | 50 | A | 0.65 | 0.692 | 0.528 | 0.814 |
| SER | 51 | A | 0.703 | 0.739 | 1 | 0.442 |
| GLU | 52 | A | 0.793 | 0.787 | 0.991 | 0.589 |
| ASP | 53 | A | 0.714 | 0.709 | 1 | 0.423 |
| GLU | 54 | A | 0.79 | 0.8 | 0.998 | 0.592 |
| MET | 55 | A | 0.711 | 0.696 | 0.999 | 0.408 |
| LYS | 56 | A | 0.75 | 0.763 | 0.749 | 0.764 |
| ALA | 57 | A | 0.549 | 0.518 | 0.999 | 0.068 |
| SER | 58 | A | 0.628 | 0.601 | 1 | 0.229 |
| GLU | 59 | A | 0.727 | 0.754 | 0.996 | 0.485 |
| ASP | 60 | A | 0.675 | 0.786 | 0.651 | 0.81 |
| LEU | 61 | A | 0.713 | 0.716 | 0.998 | 0.431 |
| LYS | 62 | A | 0.774 | 0.773 | 0.997 | 0.55 |
| LYS | 63 | A | 0.792 | 0.806 | 0.995 | 0.603 |
| **HIS** | **64** | **A** | **0.755** | **0.743** | **0.637** | **0.861** |
| GLY | 65 | A | 0.619 | 0.631 | 1 | 0.25 |
| ASN | 66 | A | 0.764 | 0.773 | 1 | 0.537 |
| THR | 67 | A | 0.738 | 0.744 | 0.999 | 0.483 |
| ASN | 68 | A | 0.793 | 0.788 | 0.998 | 0.583 |
| LEU | 69 | A | 0.74 | 0.751 | 0.999 | 0.492 |
| THR | 70 | A | 0.712 | 0.707 | 0.999 | 0.42 |
| ALA | 71 | A | 0.625 | 0.606 | 0.998 | 0.233 |
| LEU | 72 | A | 0.676 | 0.693 | 1 | 0.369 |
| GLY | 73 | A | 0.61 | 0.617 | 0.999 | 0.228 |
| GLY | 74 | A | 0.641 | 0.645 | 1 | 0.286 |
| ILE | 75 | A | 0.755 | 0.761 | 0.999 | 0.517 |
| LEU | 76 | A | 0.721 | 0.72 | 0.999 | 0.442 |
| LYS | 77 | A | 0.767 | 0.767 | 0.987 | 0.547 |
| LYS | 78 | A | 0.8 | 0.801 | 0.998 | 0.603 |
| **ALYS** | **79** | **A** | **0.794** | **0.792** | **0.741** | **0.845** |
| AGLY | 80 | A | 0.791 | 0.78 | 0.841 | 0.73 |
| HIS | 81 | A | 0.847 | 0.835 | 0.997 | 0.685 |
| HIS | 82 | A | 0.806 | 0.812 | 0.998 | 0.62 |
| GLU | 83 | A | 0.801 | 0.8 | 0.998 | 0.603 |
| ALA | 84 | A | 0.599 | 0.635 | 1 | 0.234 |
| GLU | 85 | A | 0.753 | 0.748 | 0.999 | 0.502 |
| LEU | 86 | A | 0.718 | 0.716 | 0.999 | 0.435 |
| THR | 87 | A | 0.771 | 0.758 | 0.998 | 0.531 |
| PRO | 88 | A | 0.713 | 0.711 | 0.999 | 0.425 |
| LEU | 89 | A | 0.755 | 0.751 | 1 | 0.506 |
| ALA | 90 | A | 0.469 | 0.558 | 0.999 | 0.028 |
| GLN | 91 | A | 0.825 | 0.806 | 0.999 | 0.632 |
| SER | 92 | A | 0.708 | 0.708 | 1 | 0.416 |
| HIS | 93 | A | 0.813 | 0.825 | 0.999 | 0.639 |
| ALA | 94 | A | 0.492 | 0.568 | 0.998 | 0.062 |
| THR | 95 | A | 0.784 | 0.743 | 0.999 | 0.528 |
| LYS | 96 | A | 0.729 | 0.705 | 0.995 | 0.439 |
| HIS | 97 | A | 0.831 | 0.842 | 1 | 0.673 |
| **LYS** | **98** | **A** | **0.854** | **0.84** | **0.729** | **0.965** |
| ILE | 99 | A | 0.75 | 0.739 | 0.997 | 0.492 |
| PRO | 100 | A | 0.688 | 0.666 | 0.993 | 0.361 |
| VAL | 101 | A | 0.693 | 0.657 | 0.999 | 0.351 |
| LYS | 102 | A | 0.781 | 0.768 | 0.912 | 0.637 |
| TYR | 103 | A | 0.865 | 0.852 | 1 | 0.717 |
| LEU | 104 | A | 0.719 | 0.702 | 0.999 | 0.422 |
| GLU | 105 | A | 0.786 | 0.768 | 0.998 | 0.556 |
| PHE | 106 | A | 0.864 | 0.86 | 1 | 0.724 |
| ILE | 107 | A | 0.757 | 0.75 | 1 | 0.507 |
| SER | 108 | A | 0.684 | 0.695 | 0.999 | 0.38 |
| **GLU** | **109** | **A** | **0.872** | **0.835** | **0.801** | **0.906** |
| ALA | 110 | A | 0.521 | 0.531 | 1 | 0.052 |
| ILE | 111 | A | 0.761 | 0.77 | 1 | 0.531 |
| ILE | 112 | A | 0.613 | 0.642 | 1 | 0.255 |
| GLN | 113 | A | 0.834 | 0.826 | 0.999 | 0.661 |
| VAL | 114 | A | 0.7 | 0.704 | 1 | 0.404 |
| LEU | 115 | A | 0.745 | 0.751 | 1 | 0.496 |
| GLN | 116 | A | 0.751 | 0.745 | 0.999 | 0.497 |
| **SER** | **117** | **A** | **0.823** | **0.825** | **0.697** | **0.951** |
| LYS | 118 | A | 0.759 | 0.775 | 1 | 0.534 |
| HIS | 119 | A | 0.807 | 0.806 | 1 | 0.613 |
| PRO | 120 | A | 0.738 | 0.711 | 0.98 | 0.469 |
| GLY | 121 | A | 0.655 | 0.659 | 1 | 0.314 |
| ASP | 122 | A | 0.719 | 0.695 | 0.992 | 0.422 |
| PHE | 123 | A | 0.816 | 0.812 | 0.999 | 0.629 |
| GLY | 124 | A | 0.613 | 0.618 | 1 | 0.231 |
| ALA | 125 | A | 0.707 | 0.703 | 1 | 0.41 |
| ASP | 126 | A | 0.65 | 0.691 | 0.999 | 0.342 |
| ALA | 127 | A | 0.529 | 0.528 | 1 | 0.057 |
| **GLN** | **128** | **A** | **0.844** | **0.816** | **0.648** | **1.012** |
| GLY | 129 | A | 0.645 | 0.641 | 1 | 0.286 |
| ALA | 130 | A | 0.542 | 0.562 | 1 | 0.104 |
| MET | 131 | A | 0.71 | 0.707 | 0.999 | 0.418 |
| SER | 132 | A | 0.783 | 0.812 | 0.86 | 0.735 |
| LYS | 133 | A | 0.792 | 0.827 | 0.999 | 0.62 |
| ALA | 134 | A | 0.528 | 0.536 | 1 | 0.064 |
| LEU | 135 | A | 0.697 | 0.674 | 0.999 | 0.372 |
| GLU | 136 | A | 0.77 | 0.768 | 0.999 | 0.539 |
| LEU | 137 | A | 0.727 | 0.718 | 1 | 0.445 |
| PHE | 138 | A | 0.801 | 0.817 | 0.999 | 0.619 |
| ARG | 139 | A | 0.85 | 0.848 | 1 | 0.698 |
| ASN | 140 | A | 0.789 | 0.78 | 1 | 0.569 |
| ASP | 141 | A | 0.722 | 0.724 | 1 | 0.446 |
| MET | 142 | A | 0.74 | 0.74 | 0.999 | 0.481 |
| ALA | 143 | A | 0.579 | 0.583 | 1 | 0.162 |
| ALA | 144 | A | 0.616 | 0.611 | 0.999 | 0.228 |
| **LYS** | **145** | **A** | **0.902** | **0.878** | **0.756** | **1.024** |
| TYR | 146 | A | 0.856 | 0.861 | 0.999 | 0.718 |
| LYS | 147 | A | 0.744 | 0.675 | 0.966 | 0.453 |
| GLU | 148 | A | 0.792 | 0.736 | 0.918 | 0.61 |
| LEU | 149 | A | 0.703 | 0.702 | 0.985 | 0.42 |
| GLY | 150 | A | 0.652 | 0.652 | 1 | 0.304 |
| PHE | 151 | A | 0.829 | 0.825 | 1 | 0.654 |
| GLN | 152 | A | 0.764 | 0.704 | 0.774 | 0.694 |
| GLY | 153 | A | 0.766 | 0.791 | 0.998 | 0.559 |

**Table S8(b): 1MDN (P), 1MWC (Q)**

| Residue | | | S (P) | S (Q) | J (P,Q) | PRI |
|---|---|---|---|---|---|---|
| GLY | 1 | A | 0.708 | 0.727 | 0.999 | 0.436 |
| LEU | 2 | A | 0.705 | 0.701 | 0.998 | 0.408 |
| SER | 3 | A | 0.735 | 0.746 | 1 | 0.481 |
| ASP | 4 | A | 0.603 | 0.633 | 0.999 | 0.237 |
| GLY | 5 | A | 0.632 | 0.624 | 0.999 | 0.257 |
| GLU | 6 | A | 0.82 | 0.806 | 0.999 | 0.627 |
| TRP | 7 | A | 0.886 | 0.889 | 0.999 | 0.776 |
| **GLN** | **8** | **A** | **0.831** | **0.847** | **0.824** | **0.854** |
| LEU | 9 | A | 0.675 | 0.698 | 1 | 0.373 |
| VAL | 10 | A | 0.713 | 0.698 | 1 | 0.411 |
| LEU | 11 | A | 0.706 | 0.728 | 0.999 | 0.435 |
| ASN | 12 | A | 0.829 | 0.81 | 0.998 | 0.641 |
| VAL | 13 | A | 0.488 | 0.444 | 0.993 | -0.061 |
| TRP | 14 | A | 0.884 | 0.875 | 0.998 | 0.761 |
| GLY | 15 | A | 0.637 | 0.64 | 0.999 | 0.278 |
| **LYS** | **16** | **A** | **0.762** | **0.74** | **0.573** | **0.929** |
| VAL | 17 | A | 0.682 | 0.673 | 0.998 | 0.357 |
| GLU | 18 | A | 0.761 | 0.764 | 0.797 | **0.728** |
| ALA | 19 | A | 0.499 | 0.457 | 0.998 | -0.042 |
| ASP | 20 | A | 0.73 | 0.742 | 1 | 0.472 |
| VAL | 21 | A | 0.726 | 0.745 | 1 | **0.471** |
| ALA | 22 | A | 0.575 | 0.578 | 0.999 | 0.154 |
| GLY | 23 | A | 0.635 | 0.635 | 0.999 | 0.271 |
| HIS | 24 | A | 0.78 | 0.788 | 1 | 0.568 |
| GLY | 25 | A | 0.632 | 0.628 | 0.999 | 0.261 |
| GLN | 26 | A | 0.7 | 0.732 | 0.992 | 0.44 |
| GLU | 27 | A | 0.778 | 0.799 | 0.999 | 0.578 |
| VAL | 28 | A | 0.714 | 0.725 | 0.998 | 0.441 |
| LEU | 29 | A | 0.682 | 0.692 | 0.999 | 0.375 |
| ILE | 30 | A | 0.752 | 0.744 | 0.993 | 0.503 |
| ARG | 31 | A | 0.794 | 0.783 | 0.997 | 0.58 |
| LEU | 32 | A | 0.715 | 0.714 | 0.998 | 0.431 |
| PHE | 33 | A | 0.836 | 0.847 | 0.999 | 0.684 |
| LYS | 34 | A | 0.704 | 0.74 | 0.724 | 0.72 |
| GLY | 35 | A | 0.634 | 0.645 | 0.997 | 0.282 |
| HIS | 36 | A | 0.849 | 0.846 | 0.999 | 0.696 |
| PRO | 37 | A | 0.717 | 0.706 | 1 | 0.423 |
| GLU | 38 | A | 0.77 | 0.773 | 1 | 0.543 |
| THR | 39 | A | 0.689 | 0.677 | 0.998 | 0.368 |
| LEU | 40 | A | 0.723 | 0.723 | 0.992 | 0.454 |
| **GLU** | **41** | **A** | **0.841** | **0.885** | **0.814** | **0.912** |
| LYS | 42 | A | 0.721 | 0.747 | 0.986 | 0.482 |
| PHE | 43 | A | 0.855 | 0.844 | 0.999 | 0.7 |
| ASP | 44 | A | 0.664 | 0.633 | 0.995 | 0.302 |
| LYS | 45 | A | 0.691 | 0.636 | 0.99 | 0.337 |
| PHE | 46 | A | 0.858 | 0.852 | 0.995 | 0.715 |
| **LYS** | **47** | **A** | **0.882** | **0.909** | **0.797** | **0.994** |
| HIS | 48 | A | 0.736 | 0.702 | 0.988 | 0.45 |
| LEU | 49 | A | 0.73 | 0.74 | 1 | 0.47 |
| LYS | 50 | A | 0.646 | 0.686 | 0.756 | 0.576 |
| SER | 51 | A | 0.703 | 0.751 | 1 | 0.454 |
| GLU | 52 | A | 0.793 | 0.798 | 0.995 | 0.596 |
| ASP | 53 | A | 0.714 | 0.698 | 0.997 | 0.415 |
| GLU | 54 | A | 0.79 | 0.812 | 0.999 | 0.603 |
| MET | 55 | A | 0.71 | 0.687 | 0.999 | **0.398** |
| LYS | 56 | A | 0.75 | 0.767 | 0.84 | 0.677 |
| ALA | 57 | A | 0.549 | 0.517 | 1 | 0.066 |
| SER | 58 | A | 0.63 | 0.61 | 1 | 0.24 |
| GLU | 59 | A | 0.727 | 0.709 | 0.998 | 0.438 |
| ASP | 60 | A | 0.675 | 0.757 | 0.998 | 0.434 |
| LEU | 61 | A | 0.713 | 0.71 | 1 | **0.423** |
| LYS | 62 | A | 0.774 | 0.752 | 0.999 | 0.527 |
| LYS | 63 | A | 0.792 | 0.764 | 0.886 | 0.67 |
| HIS | 64 | A | 0.754 | 0.792 | 0.92 | 0.626 |
| GLY | 65 | A | 0.619 | 0.623 | 0.993 | 0.249 |
| ASN | 66 | A | 0.763 | 0.766 | 0.999 | 0.53 |
| THR | 67 | A | 0.738 | 0.739 | 0.999 | 0.478 |
| ASN | 68 | A | 0.624 | 0.612 | 0.999 | 0.237 |
| LEU | 69 | A | 0.711 | 0.716 | 0.998 | 0.429 |
| THR | 70 | A | 0.712 | 0.717 | 1 | 0.429 |
| ALA | 71 | A | 0.627 | 0.625 | 1 | 0.252 |
| LEU | 72 | A | 0.675 | 0.698 | 0.999 | 0.374 |
| GLY | 73 | A | 0.61 | 0.62 | 1 | 0.23 |
| GLY | 74 | A | 0.641 | 0.642 | 0.999 | 0.284 |
| ILE | 75 | A | 0.755 | 0.761 | 1 | 0.516 |
| LEU | 76 | A | 0.721 | 0.725 | 0.999 | 0.447 |
| LYS | 77 | A | 0.767 | 0.761 | 0.998 | 0.53 |
| LYS | 78 | A | 0.8 | 0.8 | 0.991 | 0.609 |
| ALYS | 79 | A | 0.794 | 0.78 | 0.754 | 0.82 |
| AGLY | 80 | A | 0.791 | 0.78 | 0.842 | 0.729 |
| HIS | 81 | A | 0.847 | 0.823 | 0.995 | 0.675 |
| HIS | 82 | A | 0.806 | 0.811 | 1 | 0.617 |
| **GLU** | **83** | **A** | **0.814** | **0.763** | **0.581** | **0.996** |
| ALA | 84 | A | 0.601 | 0.607 | 0.999 | 0.209 |
| GLU | 85 | A | 0.753 | 0.776 | 0.999 | 0.53 |
| LEU | 86 | A | 0.718 | 0.725 | 0.992 | 0.451 |
| THR | 87 | A | 0.771 | 0.742 | 0.998 | 0.515 |
| PRO | 88 | A | 0.713 | 0.713 | 0.999 | 0.427 |
| LEU | 89 | A | 0.755 | 0.753 | 0.997 | 0.511 |
| ALA | 90 | A | 0.469 | 0.546 | 0.998 | 0.017 |
| GLN | 91 | A | 0.826 | 0.816 | 0.993 | 0.649 |
| SER | 92 | A | 0.704 | 0.68 | 0.999 | 0.385 |
| HIS | 93 | A | 0.813 | 0.826 | 1 | 0.639 |
| ALA | 94 | A | 0.492 | 0.494 | 0.999 | -0.013 |
| THR | 95 | A | 0.786 | 0.756 | 0.999 | 0.543 |
| **LYS** | **96** | **A** | **0.842** | **0.812** | **0.674** | **0.98** |
| HIS | 97 | A | 0.828 | 0.84 | 1 | 0.668 |
| **LYS** | **98** | **A** | **0.852** | **0.823** | **0.781** | **0.894** |
| ILE | 99 | A | 0.75 | 0.752 | 0.98 | 0.522 |
| PRO | 100 | A | 0.687 | 0.68 | 0.994 | 0.373 |
| VAL | 101 | A | 0.693 | 0.637 | 0.965 | 0.365 |
| LYS | 102 | A | 0.78 | 0.808 | 0.96 | 0.628 |
| TYR | 103 | A | 0.865 | 0.859 | 0.998 | 0.726 |
| LEU | 104 | A | 0.72 | 0.654 | 0.999 | 0.375 |
| GLU | 105 | A | 0.786 | 0.754 | 0.996 | 0.544 |
| PHE | 106 | A | 0.863 | 0.869 | 0.998 | 0.734 |
| ILE | 107 | A | 0.713 | 0.703 | 1 | 0.416 |
| SER | 108 | A | 0.684 | 0.691 | 0.999 | 0.376 |
| **GLU** | **109** | **A** | **0.872** | **0.856** | **0.746** | **0.982** |
| ALA | 110 | A | 0.521 | 0.527 | 0.999 | 0.049 |
| ILE | 111 | A | 0.762 | 0.775 | 1 | 0.537 |
| ILE | 112 | A | 0.613 | 0.601 | 0.999 | 0.215 |
| GLN | 113 | A | 0.834 | 0.833 | 0.997 | 0.67 |
| VAL | 114 | A | 0.7 | 0.705 | 0.999 | 0.406 |
| LEU | 115 | A | 0.745 | 0.755 | 0.998 | 0.502 |
| GLN | 116 | A | 0.751 | 0.751 | 0.997 | 0.505 |
| **SER** | **117** | **A** | **0.823** | **0.792** | **0.724** | **0.891** |
| LYS | 118 | A | 0.759 | 0.768 | 0.998 | 0.529 |
| HIS | 119 | A | 0.807 | 0.807 | 0.998 | 0.616 |
| PRO | 120 | A | 0.738 | 0.703 | 0.968 | 0.473 |
| GLY | 121 | A | 0.655 | 0.664 | 0.999 | 0.32 |
| ASP | 122 | A | 0.719 | 0.656 | 0.948 | 0.427 |
| PHE | 123 | A | 0.816 | 0.816 | 0.992 | 0.64 |
| GLY | 124 | A | 0.613 | 0.621 | 0.996 | 0.238 |
| ALA | 125 | A | 0.707 | 0.717 | 0.996 | 0.428 |
| ASP | 126 | A | 0.65 | 0.659 | 0.996 | 0.313 |
| ALA | 127 | A | 0.529 | 0.552 | 0.999 | 0.082 |
| **GLN** | **128** | **A** | **0.844** | **0.828** | **0.615** | **1.057** |
| GLY | 129 | A | 0.645 | 0.647 | 0.999 | 0.293 |
| ALA | 130 | A | 0.542 | 0.549 | 0.999 | 0.092 |
| MET | 131 | A | 0.71 | 0.697 | 0.998 | 0.409 |
| SER | 132 | A | 0.783 | 0.822 | 0.788 | 0.817 |
| LYS | 133 | A | 0.792 | 0.801 | 0.999 | 0.594 |
| ALA | 134 | A | 0.528 | 0.532 | 1 | 0.06 |
| LEU | 135 | A | 0.697 | 0.705 | 0.999 | 0.403 |
| GLU | 136 | A | 0.77 | 0.764 | 0.992 | 0.542 |
| LEU | 137 | A | 0.727 | 0.732 | 0.999 | 0.46 |
| PHE | 138 | A | 0.802 | 0.805 | 0.998 | 0.609 |
| ARG | 139 | A | 0.85 | 0.85 | 1 | 0.7 |
| ASN | 140 | A | 0.789 | 0.794 | 1 | 0.583 |
| ASP | 141 | A | 0.722 | 0.728 | 0.999 | 0.451 |
| MET | 142 | A | 0.74 | 0.724 | 0.998 | 0.466 |
| ALA | 143 | A | 0.579 | 0.57 | 0.999 | 0.15 |
| ALA | 144 | A | 0.616 | 0.599 | 0.998 | 0.217 |
| **LYS** | **145** | **A** | **0.902** | **0.915** | **0.741** | **1.076** |
| TYR | 146 | A | 0.856 | 0.85 | 0.973 | 0.733 |
| LYS | 147 | A | 0.744 | 0.71 | 0.987 | 0.467 |
| GLU | 148 | A | 0.792 | 0.743 | 0.896 | 0.639 |
| LEU | 149 | A | 0.702 | 0.685 | 0.996 | 0.391 |
| GLY | 150 | A | 0.652 | 0.658 | 1 | 0.31 |
| PHE | 151 | A | 0.829 | 0.84 | 0.999 | 0.67 |
| GLN | 152 | A | 0.764 | 0.72 | 0.776 | 0.708 |
| GLY | 153 | A | 0.766 | 0.794 | 0.987 | 0.573 |

**Table S8(c): 1MWD (P), 1M6C (Q)**

| Residue | | | S (P) | S (Q) | J | PRI | Residue | | | S (P) | S (Q) | J | PRI |
|---|---|---|---|---|---|---|---|---|---|---|---|---|---|
| GLY | 1 | A | 0.557 | 0.731 | 0.962 | 0.326 | LYS | 78 | A | 0.801 | 0.792 | 0.999 | 0.594 |
| LEU | 2 | A | 0.71 | 0.707 | 0.999 | 0.418 | **ALYS** | **79** | **A** | **0.792** | **0.783** | **0.753** | **0.822** |
| SER | 3 | A | 0.743 | 0.744 | 0.999 | 0.488 | AGLY | 80 | A | 0.78 | 0.781 | 0.846 | 0.715 |
| ASP | 4 | A | 0.615 | 0.618 | 0.998 | 0.235 | HIS | 81 | A | 0.835 | 0.816 | 0.996 | 0.655 |
| GLY | 5 | A | 0.625 | 0.633 | 0.999 | 0.259 | HIS | 82 | A | 0.812 | 0.807 | 0.999 | 0.62 |
| GLU | 6 | A | 0.823 | 0.838 | 0.998 | 0.663 | **GLU** | **83** | **A** | **0.8** | **0.858** | **0.765** | **0.893** |
| TRP | 7 | A | 0.885 | 0.887 | 0.998 | 0.774 | ALA | 84 | A | 0.635 | 0.619 | 0.998 | 0.256 |
| **GLN** | **8** | **A** | **0.857** | **0.852** | **0.817** | **0.892** | GLU | 85 | A | 0.748 | 0.762 | 1 | 0.51 |
| LEU | 9 | A | 0.662 | 0.68 | 0.999 | 0.343 | LEU | 86 | A | 0.716 | 0.724 | 0.999 | 0.441 |
| VAL | 10 | A | 0.72 | 0.707 | 0.999 | 0.428 | THR | 87 | A | 0.758 | 0.76 | 0.995 | 0.523 |
| LEU | 11 | A | 0.717 | 0.703 | 0.999 | 0.421 | PRO | 88 | A | 0.711 | 0.71 | 1 | 0.421 |
| ASN | 12 | A | 0.808 | 0.814 | 0.999 | 0.623 | LEU | 89 | A | 0.752 | 0.742 | 0.999 | 0.495 |
| VAL | 13 | A | 0.495 | 0.458 | 0.998 | -0.045 | ALA | 90 | A | 0.559 | 0.549 | 0.997 | 0.111 |
| TRP | 14 | A | 0.877 | 0.874 | 0.999 | 0.752 | GLN | 91 | A | 0.806 | 0.817 | 0.999 | 0.624 |
| GLY | 15 | A | 0.636 | 0.645 | 1 | 0.281 | SER | 92 | A | 0.708 | 0.704 | 0.999 | 0.413 |
| LYS | 16 | A | 0.753 | 0.773 | 0.715 | 0.811 | HIS | 93 | A | 0.825 | 0.823 | 0.999 | 0.649 |
| VAL | 17 | A | 0.679 | 0.67 | 0.998 | 0.351 | ALA | 94 | A | 0.567 | 0.508 | 0.998 | 0.077 |
| GLU | 18 | A | 0.77 | 0.768 | 0.988 | 0.55 | THR | 95 | A | 0.743 | 0.797 | 0.998 | 0.542 |
| ALA | 19 | A | 0.428 | 0.452 | 0.996 | -0.116 | **LYS** | **96** | **A** | **0.706** | **0.76** | **0.644** | **0.822** |
| ASP | 20 | A | 0.727 | 0.748 | 0.997 | 0.478 | HIS | 97 | A | 0.842 | 0.845 | 0.999 | 0.688 |
| VAL | 21 | A | 0.728 | 0.735 | 0.999 | 0.464 | **LYS** | **98** | **A** | **0.84** | **0.853** | **0.724** | **0.969** |
| ALA | 22 | A | 0.571 | 0.562 | 0.999 | 0.134 | ILE | 99 | A | 0.739 | 0.751 | 0.999 | 0.491 |
| GLY | 23 | A | 0.637 | 0.638 | 1 | 0.275 | PRO | 100 | A | 0.666 | 0.663 | 0.999 | 0.33 |
| HIS | 24 | A | 0.779 | 0.785 | 0.999 | 0.565 | VAL | 101 | A | 0.657 | 0.683 | 1 | 0.34 |
| GLY | 25 | A | 0.632 | 0.63 | 0.997 | 0.265 | LYS | 102 | A | 0.768 | 0.79 | 0.878 | 0.68 |
| GLN | 26 | A | 0.737 | 0.714 | 0.995 | 0.456 | TYR | 103 | A | 0.852 | 0.854 | 1 | 0.706 |
| GLU | 27 | A | 0.795 | 0.798 | 1 | 0.593 | LEU | 104 | A | 0.702 | 0.713 | 0.999 | 0.416 |
| VAL | 28 | A | 0.727 | 0.722 | 0.999 | 0.45 | GLU | 105 | A | 0.768 | 0.797 | 0.998 | 0.567 |
| LEU | 29 | A | 0.716 | 0.696 | 0.995 | 0.417 | PHE | 106 | A | 0.859 | 0.867 | 1 | 0.726 |
| ILE | 30 | A | 0.752 | 0.754 | 0.996 | 0.51 | ILE | 107 | A | 0.716 | 0.718 | 0.999 | 0.435 |
| ARG | 31 | A | 0.813 | 0.788 | 0.993 | 0.608 | SER | 108 | A | 0.695 | 0.68 | 0.999 | 0.376 |
| LEU | 32 | A | 0.706 | 0.717 | 1 | 0.423 | **GLU** | **109** | **A** | **0.835** | **0.862** | **0.785** | **0.912** |
| PHE | 33 | A | 0.846 | 0.837 | 1 | 0.683 | ALA | 110 | A | 0.532 | 0.524 | 1 | 0.056 |
| LYS | 34 | A | 0.732 | 0.695 | 0.993 | 0.434 | ILE | 111 | A | 0.771 | 0.768 | 0.999 | 0.54 |
| GLY | 35 | A | 0.641 | 0.638 | 1 | 0.279 | ILE | 112 | A | 0.642 | 0.596 | 0.999 | 0.239 |
| HIS | 36 | A | 0.847 | 0.848 | 1 | 0.695 | GLN | 113 | A | 0.826 | 0.84 | 0.997 | 0.669 |
| PRO | 37 | A | 0.707 | 0.707 | 1 | 0.414 | VAL | 114 | A | 0.704 | 0.707 | 0.999 | 0.412 |
| GLU | 38 | A | 0.764 | 0.776 | 1 | 0.54 | LEU | 115 | A | 0.751 | 0.74 | 1 | 0.491 |
| THR | 39 | A | 0.666 | 0.671 | 1 | 0.337 | GLN | 116 | A | 0.745 | 0.75 | 0.999 | 0.496 |
| LEU | 40 | A | 0.719 | 0.711 | 0.997 | 0.433 | **SER** | **117** | **A** | **0.825** | **0.809** | **0.662** | **0.972** |
| GLU | 41 | A | 0.823 | 0.799 | 0.886 | 0.736 | LYS | 118 | A | 0.775 | 0.763 | 1 | 0.538 |
| LYS | 42 | A | 0.741 | 0.749 | 0.999 | 0.491 | HIS | 119 | A | 0.806 | 0.801 | 0.999 | 0.608 |
| PHE | 43 | A | 0.822 | 0.849 | 0.999 | 0.672 | PRO | 120 | A | 0.711 | 0.703 | 0.999 | 0.415 |
| ASP | 44 | A | 0.626 | 0.632 | 0.996 | 0.262 | GLY | 121 | A | 0.659 | 0.66 | 1 | 0.319 |
| LYS | 45 | A | 0.689 | 0.759 | 0.675 | 0.773 | ASP | 122 | A | 0.695 | 0.671 | 0.998 | 0.368 |
| PHE | 46 | A | 0.848 | 0.854 | 0.998 | 0.704 | PHE | 123 | A | 0.812 | 0.818 | 1 | 0.63 |
| **LYS** | **47** | **A** | **0.817** | **0.835** | **0.735** | **0.917** | GLY | 124 | A | 0.618 | 0.619 | 1 | 0.237 |
| HIS | 48 | A | 0.807 | 0.848 | 0.902 | 0.753 | ALA | 125 | A | 0.703 | 0.705 | 1 | 0.408 |
| LEU | 49 | A | 0.738 | 0.734 | 0.997 | 0.475 | ASP | 126 | A | 0.691 | 0.686 | 0.999 | 0.378 |
| LYS | 50 | A | 0.692 | 0.714 | 0.632 | 0.774 | ALA | 127 | A | 0.528 | 0.501 | 1 | 0.029 |
| SER | 51 | A | 0.739 | 0.707 | 0.999 | 0.447 | **GLN** | **128** | **A** | **0.816** | **0.818** | **0.627** | **1.007** |
| GLU | 52 | A | 0.787 | 0.76 | 0.996 | 0.551 | GLY | 129 | A | 0.641 | 0.648 | 0.999 | 0.29 |
| ASP | 53 | A | 0.709 | 0.711 | 0.998 | 0.422 | ALA | 130 | A | 0.562 | 0.579 | 1 | 0.141 |
| GLU | 54 | A | 0.8 | 0.814 | 0.998 | 0.616 | MET | 131 | A | 0.707 | 0.706 | 0.999 | 0.414 |
| MET | 55 | A | 0.696 | 0.701 | 0.982 | 0.415 | SER | 132 | A | 0.812 | 0.819 | 0.82 | 0.811 |
| LYS | 56 | A | 0.763 | 0.794 | 0.999 | 0.558 | LYS | 133 | A | 0.827 | 0.812 | 1 | 0.639 |
| ALA | 57 | A | 0.518 | 0.532 | 1 | 0.05 | ALA | 134 | A | 0.536 | 0.536 | 1 | 0.072 |
| SER | 58 | A | 0.601 | 0.634 | 1 | 0.235 | LEU | 135 | A | 0.674 | 0.707 | 0.999 | 0.382 |
| GLU | 59 | A | 0.754 | 0.712 | 0.997 | 0.469 | GLU | 136 | A | 0.768 | 0.763 | 0.996 | 0.535 |
| **ASP** | **60** | **A** | **0.786** | **0.743** | **0.652** | **0.877** | LEU | 137 | A | 0.718 | 0.729 | 0.999 | 0.448 |
| LEU | 61 | A | 0.716 | 0.707 | 0.998 | 0.425 | PHE | 138 | A | 0.818 | 0.812 | 0.999 | 0.631 |
| LYS | 62 | A | 0.773 | 0.77 | 0.999 | 0.544 | ARG | 139 | A | 0.848 | 0.852 | 1 | 0.7 |
| LYS | 63 | A | 0.806 | 0.812 | 0.984 | 0.634 | ASN | 140 | A | 0.78 | 0.782 | 1 | 0.562 |
| **HIS** | **64** | **A** | **0.743** | **0.767** | **0.633** | **0.877** | ASP | 141 | A | 0.724 | 0.724 | 1 | 0.448 |
| GLY | 65 | A | 0.631 | 0.618 | 0.999 | 0.25 | MET | 142 | A | 0.74 | 0.725 | 0.999 | 0.466 |
| ASN | 66 | A | 0.772 | 0.769 | 0.999 | 0.542 | ALA | 143 | A | 0.583 | 0.571 | 1 | 0.154 |
| THR | 67 | A | 0.744 | 0.73 | 0.997 | 0.477 | ALA | 144 | A | 0.611 | 0.586 | 0.999 | 0.198 |
| ASN | 68 | A | 0.626 | 0.608 | 0.998 | 0.236 | **LYS** | **145** | **A** | **0.878** | **0.92** | **0.77** | **1.028** |
| LEU | 69 | A | 0.722 | 0.711 | 0.999 | 0.434 | TYR | 146 | A | 0.861 | 0.863 | 0.999 | 0.725 |
| THR | 70 | A | 0.707 | 0.72 | 0.999 | 0.428 | LYS | 147 | A | 0.675 | 0.693 | 0.961 | 0.407 |
| ALA | 71 | A | 0.609 | 0.63 | 0.996 | 0.243 | GLU | 148 | A | 0.736 | 0.736 | 0.993 | 0.479 |
| LEU | 72 | A | 0.693 | 0.68 | 0.999 | 0.374 | LEU | 149 | A | 0.702 | 0.698 | 0.988 | 0.412 |
| GLY | 73 | A | 0.616 | 0.623 | 1 | 0.239 | GLY | 150 | A | 0.653 | 0.649 | 1 | 0.302 |
| GLY | 74 | A | 0.645 | 0.636 | 0.999 | 0.282 | PHE | 151 | A | 0.825 | 0.835 | 1 | 0.66 |
| ILE | 75 | A | 0.762 | 0.751 | 0.999 | 0.514 | GLN | 152 | A | 0.704 | 0.703 | 1 | 0.407 |
| LEU | 76 | A | 0.72 | 0.717 | 1 | 0.437 | GLY | 153 | A | 0.791 | 0.787 | 0.988 | 0.59 |
| LYS | 77 | A | 0.768 | 0.765 | 0.995 | 0.538 | | | | | | | |

**Table S8(d): 1MWD (P), 1MDN (Q)**

| Residue | | | S (P) | S (Q) | J (P,Q) | PRI |
|---|---|---|---|---|---|---|
| GLY | 1 | A | 0.708 | 0.731 | 0.958 | 0.481 |
| LEU | 2 | A | 0.705 | 0.707 | 0.999 | 0.413 |
| SER | 3 | A | 0.735 | 0.745 | 1 | 0.48 |
| ASP | 4 | A | 0.603 | 0.618 | 0.999 | 0.222 |
| GLY | 5 | A | 0.632 | 0.633 | 0.999 | 0.266 |
| GLU | 6 | A | 0.82 | 0.838 | 0.999 | 0.659 |
| TRP | 7 | A | 0.886 | 0.887 | 0.999 | 0.774 |
| **GLN** | **8** | **A** | **0.831** | **0.852** | **0.803** | **0.88** |
| LEU | 9 | A | 0.675 | 0.68 | 0.999 | 0.356 |
| VAL | 10 | A | 0.713 | 0.707 | 1 | 0.42 |
| LEU | 11 | A | 0.706 | 0.703 | 0.999 | 0.41 |
| ASN | 12 | A | 0.829 | 0.814 | 0.997 | 0.646 |
| VAL | 13 | A | 0.488 | 0.458 | 0.997 | -0.051 |
| TRP | 14 | A | 0.884 | 0.874 | 1 | 0.758 |
| GLY | 15 | A | 0.637 | 0.645 | 0.999 | 0.283 |
| **LYS** | **16** | **A** | **0.762** | **0.773** | **0.536** | **0.999** |
| VAL | 17 | A | 0.682 | 0.67 | 0.999 | 0.353 |
| GLU | 18 | A | 0.761 | 0.768 | 0.689 | 0.84 |
| ALA | 19 | A | 0.499 | 0.453 | 0.999 | -0.047 |
| ASP | 20 | A | 0.73 | 0.748 | 0.998 | 0.48 |
| VAL | 21 | A | 0.726 | 0.735 | 1 | 0.461 |
| ALA | 22 | A | 0.575 | 0.562 | 0.999 | 0.138 |
| GLY | 23 | A | 0.635 | 0.638 | 0.999 | 0.274 |
| HIS | 24 | A | 0.78 | 0.785 | 1 | 0.565 |
| GLY | 25 | A | 0.632 | 0.63 | 0.999 | 0.263 |
| GLN | 26 | A | 0.7 | 0.714 | 0.998 | 0.416 |
| GLU | 27 | A | 0.778 | 0.798 | 1 | 0.576 |
| VAL | 28 | A | 0.714 | 0.722 | 0.999 | 0.437 |
| LEU | 29 | A | 0.682 | 0.696 | 1 | 0.378 |
| ILE | 30 | A | 0.752 | 0.754 | 0.999 | 0.507 |
| ARG | 31 | A | 0.794 | 0.788 | 0.999 | 0.583 |
| LEU | 32 | A | 0.715 | 0.717 | 0.999 | 0.433 |
| PHE | 33 | A | 0.836 | 0.836 | 0.999 | 0.673 |
| LYS | 34 | A | 0.704 | 0.695 | 0.716 | 0.683 |
| GLY | 35 | A | 0.634 | 0.638 | 0.999 | 0.273 |
| HIS | 36 | A | 0.849 | 0.848 | 0.999 | 0.698 |
| PRO | 37 | A | 0.717 | 0.709 | 1 | 0.426 |
| GLU | 38 | A | 0.77 | 0.782 | 1 | 0.552 |
| THR | 39 | A | 0.689 | 0.673 | 1 | 0.362 |
| LEU | 40 | A | 0.722 | 0.712 | 0.991 | 0.443 |
| **GLU** | **41** | **A** | **0.841** | **0.883** | **0.791** | **0.933** |
| LYS | 42 | A | 0.722 | 0.749 | 0.999 | 0.472 |
| PHE | 43 | A | 0.855 | 0.853 | 0.999 | 0.709 |
| ASP | 44 | A | 0.665 | 0.648 | 0.993 | 0.32 |
| **LYS** | **45** | **A** | **0.695** | **0.759** | **0.567** | **0.887** |
| PHE | 46 | A | 0.859 | 0.855 | 1 | 0.714 |
| **LYS** | **47** | **A** | **0.882** | **0.892** | **0.717** | **1.057** |
| HIS | 48 | A | 0.825 | 0.847 | 0.997 | 0.675 |
| LEU | 49 | A | 0.686 | 0.734 | 0.999 | 0.421 |
| LYS | 50 | A | 0.645 | 0.71 | 0.741 | 0.614 |
| SER | 51 | A | 0.703 | 0.707 | 0.999 | 0.411 |
| GLU | 52 | A | 0.793 | 0.76 | 0.996 | 0.557 |
| ASP | 53 | A | 0.714 | 0.711 | 0.997 | 0.428 |
| GLU | 54 | A | 0.79 | 0.813 | 0.998 | 0.605 |
| MET | 55 | A | 0.71 | 0.701 | 0.997 | 0.414 |
| LYS | 56 | A | 0.75 | 0.794 | 0.79 | 0.754 |
| ALA | 57 | A | 0.549 | 0.532 | 1 | 0.081 |
| SER | 58 | A | 0.628 | 0.634 | 0.999 | 0.263 |
| GLU | 59 | A | 0.727 | 0.712 | 0.999 | 0.44 |
| ASP | 60 | A | 0.675 | 0.743 | 1 | 0.418 |
| LEU | 61 | A | 0.713 | 0.707 | 1 | 0.42 |
| LYS | 62 | A | 0.774 | 0.77 | 1 | 0.544 |
| LYS | 63 | A | 0.792 | 0.812 | 0.999 | 0.605 |
| HIS | 64 | A | 0.755 | 0.767 | 0.999 | 0.523 |
| GLY | 65 | A | 0.619 | 0.618 | 1 | 0.237 |
| ASN | 66 | A | 0.764 | 0.769 | 0.999 | 0.534 |
| THR | 67 | A | 0.738 | 0.73 | 0.998 | 0.47 |
| ASN | 68 | A | 0.624 | 0.607 | 0.999 | 0.232 |
| LEU | 69 | A | 0.711 | 0.711 | 0.999 | 0.423 |
| THR | 70 | A | 0.712 | 0.72 | 1 | 0.432 |
| ALA | 71 | A | 0.627 | 0.63 | 1 | 0.257 |
| LEU | 72 | A | 0.675 | 0.68 | 1 | 0.355 |
| GLY | 73 | A | 0.61 | 0.624 | 0.999 | 0.235 |
| GLY | 74 | A | 0.641 | 0.636 | 0.999 | 0.278 |
| ILE | 75 | A | 0.755 | 0.751 | 1 | 0.506 |
| LEU | 76 | A | 0.722 | 0.716 | 1 | 0.438 |
| LYS | 77 | A | 0.767 | 0.765 | 0.999 | 0.533 |
| LYS | 78 | A | 0.8 | 0.792 | 0.999 | 0.593 |
| **ALYS** | **79** | **A** | **0.794** | **0.783** | **0.716** | **0.861** |
| AGLY | 80 | A | 0.791 | 0.781 | 0.834 | 0.738 |
| HIS | 81 | A | 0.847 | 0.816 | 0.997 | 0.666 |
| HIS | 82 | A | 0.806 | 0.807 | 1 | 0.613 |
| **GLU** | **83** | **A** | **0.801** | **0.858** | **0.677** | **0.982** |
| ALA | 84 | A | 0.599 | 0.619 | 1 | 0.218 |
| GLU | 85 | A | 0.753 | 0.762 | 0.999 | 0.516 |
| LEU | 86 | A | 0.718 | 0.724 | 1 | 0.442 |
| THR | 87 | A | 0.771 | 0.76 | 0.996 | 0.535 |
| PRO | 88 | A | 0.713 | 0.71 | 1 | 0.423 |
| LEU | 89 | A | 0.755 | 0.743 | 0.998 | 0.5 |
| ALA | 90 | A | 0.469 | 0.549 | 0.989 | 0.029 |
| GLN | 91 | A | 0.825 | 0.817 | 0.998 | 0.644 |
| SER | 92 | A | 0.708 | 0.704 | 0.998 | 0.414 |
| HIS | 93 | A | 0.813 | 0.823 | 0.999 | 0.637 |
| ALA | 94 | A | 0.492 | 0.508 | 0.998 | 0.002 |
| THR | 95 | A | 0.784 | 0.797 | 0.998 | 0.583 |
| **LYS** | **96** | **A** | **0.729** | **0.759** | **0.63** | **0.858** |
| HIS | 97 | A | 0.831 | 0.844 | 1 | 0.675 |
| **LYS** | **98** | **A** | **0.851** | **0.851** | **0.697** | **1.005** |
| ILE | 99 | A | 0.75 | 0.751 | 0.999 | 0.502 |
| PRO | 100 | A | 0.687 | 0.663 | 0.996 | 0.354 |
| VAL | 101 | A | 0.693 | 0.683 | 1 | 0.376 |
| LYS | 102 | A | 0.78 | 0.789 | 0.981 | 0.588 |
| TYR | 103 | A | 0.865 | 0.854 | 0.999 | 0.72 |
| LEU | 104 | A | 0.72 | 0.713 | 0.999 | 0.434 |
| GLU | 105 | A | 0.786 | 0.797 | 0.998 | 0.585 |
| PHE | 106 | A | 0.863 | 0.866 | 0.999 | 0.73 |
| ILE | 107 | A | 0.713 | 0.718 | 1 | 0.431 |
| SER | 108 | A | 0.684 | 0.68 | 0.999 | 0.365 |
| **GLU** | **109** | **A** | **0.872** | **0.862** | **0.74** | **0.994** |
| ALA | 110 | A | 0.521 | 0.524 | 1 | 0.045 |
| ILE | 111 | A | 0.762 | 0.768 | 1 | 0.53 |
| ILE | 112 | A | 0.613 | 0.596 | 1 | 0.209 |
| GLN | 113 | A | 0.834 | 0.84 | 0.994 | 0.68 |
| VAL | 114 | A | 0.7 | 0.707 | 1 | 0.407 |
| LEU | 115 | A | 0.745 | 0.74 | 0.999 | 0.486 |
| GLN | 116 | A | 0.751 | 0.75 | 0.999 | 0.502 |
| **SER** | **117** | **A** | **0.823** | **0.809** | **0.62** | **1.012** |
| LYS | 118 | A | 0.759 | 0.763 | 1 | 0.522 |
| HIS | 119 | A | 0.807 | 0.801 | 0.999 | 0.609 |
| PRO | 120 | A | 0.738 | 0.703 | 0.985 | 0.456 |
| GLY | 121 | A | 0.655 | 0.66 | 1 | 0.315 |
| ASP | 122 | A | 0.719 | 0.671 | 0.981 | 0.409 |
| PHE | 123 | A | 0.816 | 0.818 | 0.997 | 0.637 |
| GLY | 124 | A | 0.613 | 0.619 | 0.998 | 0.234 |
| ALA | 125 | A | 0.707 | 0.705 | 0.998 | 0.414 |
| ASP | 126 | A | 0.65 | 0.686 | 0.999 | 0.337 |
| ALA | 127 | A | 0.529 | 0.502 | 1 | 0.031 |
| **GLN** | **128** | **A** | **0.844** | **0.818** | **0.598** | **1.064** |
| GLY | 129 | A | 0.645 | 0.648 | 1 | 0.293 |
| ALA | 130 | A | 0.542 | 0.578 | 1 | 0.12 |
| MET | 131 | A | 0.71 | 0.706 | 0.999 | 0.417 |
| SER | 132 | A | 0.783 | 0.819 | 0.777 | 0.825 |
| LYS | 133 | A | 0.792 | 0.812 | 0.998 | 0.606 |
| ALA | 134 | A | 0.528 | 0.536 | 0.999 | 0.065 |
| LEU | 135 | A | 0.697 | 0.707 | 0.998 | 0.406 |
| GLU | 136 | A | 0.77 | 0.763 | 0.981 | 0.552 |
| LEU | 137 | A | 0.727 | 0.729 | 1 | 0.456 |
| PHE | 138 | A | 0.802 | 0.812 | 0.998 | 0.616 |
| ARG | 139 | A | 0.85 | 0.852 | 1 | 0.702 |
| ASN | 140 | A | 0.789 | 0.782 | 1 | 0.571 |
| ASP | 141 | A | 0.722 | 0.724 | 0.999 | 0.447 |
| MET | 142 | A | 0.74 | 0.725 | 0.999 | 0.466 |
| ALA | 143 | A | 0.579 | 0.571 | 1 | 0.15 |
| ALA | 144 | A | 0.616 | 0.586 | 0.998 | 0.204 |
| **LYS** | **145** | **A** | **0.902** | **0.92** | **0.733** | **1.089** |
| TYR | 146 | A | 0.856 | 0.863 | 0.999 | 0.72 |
| LYS | 147 | A | 0.744 | 0.693 | 0.991 | 0.446 |
| GLU | 148 | A | 0.792 | 0.736 | 0.856 | 0.672 |
| LEU | 149 | A | 0.703 | 0.699 | 0.994 | 0.408 |
| GLY | 150 | A | 0.652 | 0.65 | 0.999 | 0.303 |
| PHE | 151 | A | 0.829 | 0.835 | 0.998 | 0.666 |
| GLN | 152 | A | 0.764 | 0.702 | 0.721 | 0.745 |
| GLY | 153 | A | 0.766 | 0.787 | 0.997 | 0.556 |

| Table S8(e): 1MWD (P), 1MWC (Q) | | | | | | |
|---|---|---|---|---|---|---|
| Residue | | | S (P) | S (Q) | J (P,Q) | PRI |
| GLY | 1 | A | 0.731 | 0.727 | 0.985 | 0.473 |
| LEU | 2 | A | 0.707 | 0.701 | 0.999 | 0.409 |
| SER | 3 | A | 0.745 | 0.746 | 0.999 | 0.492 |
| ASP | 4 | A | 0.618 | 0.633 | 0.999 | 0.252 |
| GLY | 5 | A | 0.633 | 0.624 | 0.999 | 0.258 |
| GLU | 6 | A | 0.839 | 0.806 | 0.999 | 0.646 |
| TRP | 7 | A | 0.887 | 0.889 | 0.998 | 0.778 |
| **GLN** | **8** | **A** | **0.852** | **0.847** | **0.792** | **0.907** |
| LEU | 9 | A | 0.68 | 0.698 | 0.999 | 0.379 |
| VAL | 10 | A | 0.707 | 0.697 | 0.999 | 0.405 |
| LEU | 11 | A | 0.703 | 0.728 | 0.999 | 0.432 |
| ASN | 12 | A | 0.814 | 0.81 | 0.996 | 0.628 |
| VAL | 13 | A | 0.458 | 0.444 | 0.998 | -0.096 |
| TRP | 14 | A | 0.874 | 0.875 | 1 | 0.749 |
| GLY | 15 | A | 0.645 | 0.64 | 1 | 0.285 |
| LYS | 16 | A | 0.773 | 0.74 | 0.803 | 0.71 |
| VAL | 17 | A | 0.67 | 0.673 | 1 | 0.343 |
| GLU | 18 | A | 0.768 | 0.764 | 0.969 | 0.563 |
| ALA | 19 | A | 0.452 | 0.458 | 0.999 | -0.089 |
| ASP | 20 | A | 0.748 | 0.742 | 0.999 | 0.491 |
| VAL | 21 | A | 0.735 | 0.745 | 1 | 0.48 |
| ALA | 22 | A | 0.562 | 0.578 | 0.999 | 0.141 |
| GLY | 23 | A | 0.638 | 0.636 | 1 | 0.274 |
| HIS | 24 | A | 0.785 | 0.788 | 1 | 0.573 |
| GLY | 25 | A | 0.63 | 0.628 | 0.998 | 0.26 |
| GLN | 26 | A | 0.714 | 0.732 | 0.999 | 0.447 |
| GLU | 27 | A | 0.798 | 0.799 | 1 | 0.597 |
| VAL | 28 | A | 0.721 | 0.725 | 0.999 | 0.447 |
| LEU | 29 | A | 0.694 | 0.69 | 0.999 | 0.385 |
| ILE | 30 | A | 0.754 | 0.744 | 0.999 | 0.499 |
| ARG | 31 | A | 0.788 | 0.783 | 0.998 | 0.573 |
| LEU | 32 | A | 0.716 | 0.713 | 0.998 | 0.431 |
| PHE | 33 | A | 0.836 | 0.847 | 1 | 0.683 |
| LYS | 34 | A | 0.695 | 0.74 | 0.998 | 0.437 |
| GLY | 35 | A | 0.637 | 0.644 | 1 | 0.281 |
| HIS | 36 | A | 0.848 | 0.846 | 0.999 | 0.695 |
| PRO | 37 | A | 0.709 | 0.706 | 1 | 0.415 |
| GLU | 38 | A | 0.782 | 0.774 | 0.998 | 0.558 |
| THR | 39 | A | 0.673 | 0.677 | 0.999 | 0.351 |
| LEU | 40 | A | 0.712 | 0.723 | 0.999 | 0.436 |
| **GLU** | **41** | **A** | **0.883** | **0.884** | **0.77** | **0.997** |
| LYS | 42 | A | 0.749 | 0.747 | 0.992 | 0.504 |
| PHE | 43 | A | 0.853 | 0.843 | 0.999 | 0.697 |
| ASP | 44 | A | 0.644 | 0.633 | 0.999 | 0.278 |
| LYS | 45 | A | 0.755 | 0.636 | 0.561 | 0.83 |
| PHE | 46 | A | 0.854 | 0.851 | 0.998 | 0.707 |
| **LYS** | **47** | **A** | **0.891** | **0.909** | **0.803** | **0.997** |
| HIS | 48 | A | 0.74 | 0.702 | 0.99 | 0.452 |
| LEU | 49 | A | 0.743 | 0.741 | 0.999 | 0.485 |
| LYS | 50 | A | 0.711 | 0.686 | 1 | 0.397 |
| SER | 51 | A | 0.707 | 0.751 | 1 | 0.458 |
| GLU | 52 | A | 0.76 | 0.798 | 0.998 | 0.56 |
| ASP | 53 | A | 0.711 | 0.698 | 0.991 | 0.418 |
| GLU | 54 | A | 0.814 | 0.812 | 1 | 0.626 |
| MET | 55 | A | 0.701 | 0.687 | 1 | 0.388 |
| LYS | 56 | A | 0.794 | 0.767 | 1 | 0.561 |
| ALA | 57 | A | 0.532 | 0.517 | 0.999 | 0.05 |
| SER | 58 | A | 0.635 | 0.61 | 0.997 | 0.248 |
| GLU | 59 | A | 0.713 | 0.709 | 0.998 | 0.424 |
| ASP | 60 | A | 0.744 | 0.757 | 0.996 | 0.505 |
| LEU | 61 | A | 0.707 | 0.71 | 0.999 | 0.418 |
| LYS | 62 | A | 0.77 | 0.753 | 0.999 | 0.524 |
| LYS | 63 | A | 0.812 | 0.764 | 0.931 | 0.645 |
| HIS | 64 | A | 0.776 | 0.788 | 0.946 | 0.618 |
| GLY | 65 | A | 0.617 | 0.622 | 0.994 | 0.245 |
| ASN | 66 | A | 0.769 | 0.766 | 0.999 | 0.536 |
| THR | 67 | A | 0.728 | 0.738 | 0.999 | 0.467 |
| VAL | 68 | A | 0.729 | 0.737 | 0.996 | 0.47 |
| LEU | 69 | A | 0.711 | 0.715 | 0.999 | 0.427 |
| THR | 70 | A | 0.72 | 0.717 | 1 | 0.437 |
| ALA | 71 | A | 0.628 | 0.624 | 1 | 0.252 |
| LEU | 72 | A | 0.68 | 0.698 | 1 | 0.378 |
| GLY | 73 | A | 0.623 | 0.62 | 1 | 0.243 |
| GLY | 74 | A | 0.636 | 0.642 | 0.999 | 0.279 |
| ILE | 75 | A | 0.751 | 0.761 | 0.998 | 0.514 |
| LEU | 76 | A | 0.716 | 0.725 | 0.999 | 0.442 |
| LYS | 77 | A | 0.765 | 0.761 | 0.998 | 0.528 |
| LYS | 78 | A | 0.792 | 0.8 | 0.987 | 0.605 |
| **ALYS** | **79** | **A** | **0.783** | **0.78** | **0.702** | **0.861** |
| AGLY | 80 | A | 0.781 | 0.78 | 0.833 | 0.728 |
| HIS | 81 | A | 0.816 | 0.823 | 0.996 | 0.643 |
| HIS | 82 | A | 0.807 | 0.811 | 0.999 | 0.619 |
| **GLU** | **83** | **A** | **0.852** | **0.763** | **0.527** | **1.088** |
| ALA | 84 | A | 0.619 | 0.607 | 0.999 | 0.227 |
| GLU | 85 | A | 0.762 | 0.776 | 0.997 | 0.541 |
| LEU | 86 | A | 0.724 | 0.725 | 0.995 | 0.454 |
| THR | 87 | A | 0.76 | 0.742 | 0.998 | 0.504 |
| PRO | 88 | A | 0.71 | 0.713 | 0.999 | 0.424 |
| LEU | 89 | A | 0.742 | 0.753 | 0.996 | 0.499 |
| ALA | 90 | A | 0.549 | 0.546 | 0.999 | 0.096 |
| GLN | 91 | A | 0.817 | 0.816 | 0.995 | 0.638 |
| SER | 92 | A | 0.703 | 0.68 | 0.997 | 0.386 |
| HIS | 93 | A | 0.822 | 0.825 | 0.998 | 0.649 |
| ALA | 94 | A | 0.508 | 0.494 | 0.998 | 0.004 |
| THR | 95 | A | 0.798 | 0.756 | 0.998 | 0.556 |
| **LYS** | **96** | **A** | **0.85** | **0.812** | **0.579** | **1.083** |
| HIS | 97 | A | 0.843 | 0.84 | 0.999 | 0.684 |
| **LYS** | **98** | **A** | **0.851** | **0.823** | **0.758** | **0.916** |
| ILE | 99 | A | 0.75 | 0.752 | 0.966 | 0.536 |
| PRO | 100 | A | 0.662 | 0.679 | 0.985 | 0.356 |
| VAL | 101 | A | 0.683 | 0.637 | 0.97 | 0.35 |
| LYS | 102 | A | 0.789 | 0.808 | 0.997 | 0.6 |
| TYR | 103 | A | 0.854 | 0.859 | 0.999 | 0.714 |
| LEU | 104 | A | 0.713 | 0.654 | 0.999 | 0.368 |
| GLU | 105 | A | 0.797 | 0.754 | 0.994 | 0.557 |
| PHE | 106 | A | 0.866 | 0.869 | 1 | 0.735 |
| ILE | 107 | A | 0.717 | 0.702 | 0.998 | 0.421 |
| SER | 108 | A | 0.68 | 0.691 | 0.999 | 0.372 |
| **GLU** | **109** | **A** | **0.862** | **0.856** | **0.719** | **0.999** |
| ALA | 110 | A | 0.524 | 0.527 | 0.999 | 0.052 |
| ILE | 111 | A | 0.768 | 0.775 | 1 | 0.543 |
| ILE | 112 | A | 0.596 | 0.601 | 1 | 0.197 |
| GLN | 113 | A | 0.84 | 0.833 | 0.999 | 0.674 |
| VAL | 114 | A | 0.707 | 0.705 | 0.999 | 0.413 |
| LEU | 115 | A | 0.74 | 0.755 | 0.999 | 0.496 |
| GLN | 116 | A | 0.75 | 0.751 | 0.998 | 0.503 |
| **SER** | **117** | **A** | **0.809** | **0.792** | **0.645** | **0.956** |
| LYS | 118 | A | 0.763 | 0.768 | 0.999 | 0.532 |
| HIS | 119 | A | 0.801 | 0.807 | 0.999 | 0.609 |
| PRO | 120 | A | 0.703 | 0.703 | 0.996 | 0.41 |
| GLY | 121 | A | 0.66 | 0.663 | 1 | 0.323 |
| ASP | 122 | A | 0.671 | 0.656 | 0.994 | 0.333 |
| PHE | 123 | A | 0.818 | 0.816 | 0.997 | 0.637 |
| GLY | 124 | A | 0.619 | 0.621 | 0.999 | 0.241 |
| ALA | 125 | A | 0.705 | 0.717 | 0.999 | 0.423 |
| ASP | 126 | A | 0.686 | 0.659 | 0.994 | 0.351 |
| ALA | 127 | A | 0.501 | 0.552 | 0.998 | 0.055 |
| **GLN** | **128** | **A** | **0.818** | **0.828** | **0.591** | **1.055** |
| GLY | 129 | A | 0.648 | 0.647 | 1 | 0.295 |
| ALA | 130 | A | 0.579 | 0.548 | 1 | 0.127 |
| MET | 131 | A | 0.706 | 0.697 | 0.998 | 0.405 |
| **SER** | **132** | **A** | **0.819** | **0.822** | **0.721** | **0.92** |
| LYS | 133 | A | 0.812 | 0.801 | 0.997 | 0.616 |
| ALA | 134 | A | 0.536 | 0.532 | 1 | 0.068 |
| LEU | 135 | A | 0.707 | 0.704 | 0.998 | 0.413 |
| GLU | 136 | A | 0.763 | 0.764 | 0.998 | 0.529 |
| LEU | 137 | A | 0.729 | 0.732 | 0.999 | 0.462 |
| PHE | 138 | A | 0.812 | 0.805 | 0.998 | 0.619 |
| ARG | 139 | A | 0.852 | 0.85 | 1 | 0.702 |
| ASN | 140 | A | 0.782 | 0.794 | 1 | 0.576 |
| ASP | 141 | A | 0.724 | 0.728 | 0.999 | 0.453 |
| MET | 142 | A | 0.725 | 0.724 | 0.998 | 0.451 |
| ALA | 143 | A | 0.571 | 0.57 | 1 | 0.141 |
| ALA | 144 | A | 0.586 | 0.599 | 0.998 | 0.187 |
| **LYS** | **145** | **A** | **0.92** | **0.914** | **0.739** | **1.095** |
| TYR | 146 | A | 0.863 | 0.85 | 0.955 | 0.758 |
| LYS | 147 | A | 0.693 | 0.71 | 0.992 | 0.411 |
| GLU | 148 | A | 0.736 | 0.743 | 0.997 | 0.482 |
| LEU | 149 | A | 0.698 | 0.685 | 0.995 | 0.388 |
| GLY | 150 | A | 0.649 | 0.659 | 0.998 | 0.31 |
| PHE | 151 | A | 0.835 | 0.84 | 0.998 | 0.677 |
| GLN | 152 | A | 0.702 | 0.72 | 0.992 | 0.43 |
| GLY | 153 | A | 0.787 | 0.794 | 0.96 | 0.621 |

| Table S9: 1RRS (P), 1RRQ (Q) | | | | | | | | | | | | | | |
|---|---|---|---|---|---|---|---|---|---|---|---|---|---|---|
| **Residue** | | | **S (P)** | **S (Q)** | **J (P,Q)** | **PRI** | | **Residue** | | | **S (P)** | **S (Q)** | **J (P,Q)** | **PRI** |
| PRO | 9 | A | 0.701 | 0.679 | 0.945 | 0.435 | | LEU | 80 | A | 0.729 | 0.711 | 1 | 0.44 |
| ALA | 10 | A | 0.618 | 0.609 | 0.805 | 0.422 | | LYS | 81 | A | 0.748 | 0.727 | 0.998 | 0.477 |
| ARG | 11 | A | 0.727 | 0.739 | 0.998 | 0.468 | | ALA | 82 | A | 0.471 | 0.461 | 1 | -0.068 |
| GLU | 12 | A | 0.684 | 0.66 | 0.992 | 0.352 | | **TRP** | **83** | **A** | **0.889** | **0.89** | **1** | **0.779** |
| PHE | 13 | A | 0.831 | 0.819 | 1 | 0.65 | | GLU | 84 | A | 0.776 | 0.758 | 0.998 | 0.536 |
| GLN | 14 | A | 0.85 | 0.761 | 0.866 | 0.745 | | GLY | 85 | A | 0.623 | 0.625 | 1 | 0.248 |
| ARG | 15 | A | 0.636 | 0.636 | 0.969 | 0.303 | | **LEU** | **86** | **A** | **0.726** | **0.708** | **0.655** | **0.779** |
| ASP | 16 | A | 0.677 | 0.673 | 0.991 | 0.359 | | GLY | 87 | A | 0.625 | 0.626 | 0.999 | 0.252 |
| LEU | 17 | A | 0.691 | 0.681 | 0.998 | 0.374 | | TYR | 88 | A | 0.8 | 0.79 | 1 | 0.59 |
| LEU | 18 | A | 0.775 | 0.743 | 0.999 | 0.519 | | TYR | 89 | A | 0.856 | 0.854 | 1 | 0.71 |
| ASP | 19 | A | 0.609 | 0.669 | 0.912 | 0.366 | | SER | 90 | A | 0.635 | 0.672 | 0.998 | 0.309 |
| **TRP** | **20** | **A** | **0.888** | **0.894** | **0.992** | **0.79** | | ARG | 91 | A | 0.848 | 0.838 | 0.961 | 0.725 |
| PHE | 21 | A | 0.835 | 0.834 | 0.993 | 0.676 | | VAL | 92 | A | 0.733 | 0.733 | 1 | 0.466 |
| ALA | 22 | A | 0.542 | 0.592 | 0.97 | 0.164 | | ARG | 93 | A | 0.812 | 0.834 | 0.994 | 0.652 |
| ARG | 23 | A | 0.495 | 0.507 | 0.993 | 0.009 | | ASN | 94 | A | 0.798 | 0.808 | 0.989 | 0.617 |
| GLU | 24 | A | 0.701 | 0.752 | 0.991 | 0.462 | | LEU | 95 | A | 0.682 | 0.673 | 1 | 0.355 |
| ARG | 25 | A | 0.81 | 0.842 | 0.99 | 0.662 | | HIS | 96 | A | 0.781 | 0.805 | 0.992 | 0.594 |
| ARG | 26 | A | 0.797 | 0.826 | 0.99 | 0.633 | | ALA | 97 | A | 0.594 | 0.634 | 0.999 | 0.229 |
| ASP | 27 | A | 0.451 | 0.451 | 0.997 | -0.095 | | ALA | 98 | A | 0.541 | 0.547 | 0.999 | 0.089 |
| LEU | 28 | A | 0.727 | 0.73 | 0.972 | 0.485 | | VAL | 99 | A | 0.684 | 0.735 | 0.992 | 0.427 |
| PRO | 29 | A | 0.685 | 0.647 | 0.999 | 0.333 | | LYS | 100 | A | 0.761 | 0.789 | 0.987 | 0.563 |
| **TRP** | **30** | **A** | **0.888** | **0.887** | **0.996** | **0.779** | | **GLU** | **101** | **A** | **0.77** | **0.803** | **0.517** | **1.056** |
| ARG | 31 | A | 0.827 | 0.823 | 0.992 | 0.658 | | VAL | 102 | A | 0.721 | 0.713 | 0.99 | 0.444 |
| LYS | 32 | A | 0.705 | 0.631 | 0.96 | 0.376 | | LYS | 103 | A | 0.712 | 0.685 | 0.907 | 0.49 |
| ASP | 33 | A | 0.766 | 0.722 | 1 | 0.488 | | THR | 104 | A | 0.782 | 0.762 | 0.98 | 0.564 |
| ARG | 34 | A | 0.831 | 0.844 | 0.997 | 0.678 | | ARG | 105 | A | 0.503 | 0.509 | 0.998 | 0.014 |
| ASP | 35 | A | 0.786 | 0.787 | 0.999 | 0.574 | | TYR | 106 | A | 0.845 | 0.844 | 0.997 | 0.692 |
| PRO | 36 | A | 0.679 | 0.674 | 0.996 | 0.357 | | GLY | 107 | A | 0.658 | 0.658 | 0.998 | 0.318 |
| TYR | 37 | A | 0.856 | 0.851 | 0.999 | 0.708 | | GLY | 108 | A | 0.622 | 0.622 | 1 | 0.244 |
| LYS | 38 | A | 0.762 | 0.755 | 0.877 | 0.64 | | LYS | 109 | A | 0.819 | 0.829 | 0.961 | 0.687 |
| VAL | 39 | A | 0.688 | 0.707 | 0.996 | 0.399 | | VAL | 110 | A | 0.687 | 0.678 | 0.996 | 0.369 |
| TRP | 40 | A | 0.872 | 0.875 | 1 | 0.747 | | PRO | 111 | A | 0.636 | 0.639 | 0.999 | 0.276 |
| VAL | 41 | A | 0.704 | 0.707 | 0.998 | 0.413 | | ASP | 112 | A | 0.716 | 0.69 | 0.997 | 0.409 |
| SER | 42 | A | 0.691 | 0.676 | 0.999 | 0.368 | | ASP | 113 | A | 0.789 | 0.8 | 0.993 | 0.596 |
| GLU | 43 | A | 0.761 | 0.777 | 0.999 | 0.539 | | PRO | 114 | A | 0.686 | 0.671 | 0.998 | 0.359 |
| VAL | 44 | A | 0.706 | 0.691 | 0.998 | 0.399 | | ASP | 115 | A | 0.677 | 0.693 | 0.944 | 0.426 |
| MET | 45 | A | 0.759 | 0.735 | 0.999 | 0.495 | | GLU | 116 | A | 0.645 | 0.643 | 0.997 | 0.291 |
| LEU | 46 | A | 0.753 | 0.746 | 0.997 | 0.502 | | PHE | 117 | A | 0.836 | 0.83 | 1 | 0.666 |
| GLN | 47 | A | 0.721 | 0.761 | 0.999 | 0.483 | | SER | 118 | A | 0.607 | 0.626 | 0.987 | 0.246 |
| **GLN** | **48** | **A** | **0.67** | **0.737** | **0.604** | **0.803** | | ARG | 119 | A | 0.481 | 0.524 | 0.999 | 0.006 |
| THR | 49 | A | 0.725 | 0.678 | 0.99 | 0.413 | | LEU | 120 | A | 0.752 | 0.755 | 0.999 | 0.508 |
| ARG | 50 | A | 0.58 | 0.568 | 0.91 | 0.238 | | LYS | 121 | A | 0.51 | 0.523 | 0.994 | 0.039 |
| VAL | 51 | A | 0.669 | 0.687 | 0.977 | 0.379 | | GLY | 122 | A | 0.628 | 0.631 | 1 | 0.259 |
| GLU | 52 | A | 0.652 | 0.654 | 0.997 | 0.309 | | VAL | 123 | A | 0.64 | 0.655 | 0.994 | 0.301 |
| THR | 53 | A | 0.688 | 0.661 | 0.965 | 0.384 | | GLY | 124 | A | 0.63 | 0.626 | 0.992 | 0.264 |
| VAL | 54 | A | 0.696 | 0.705 | 0.985 | 0.416 | | PRO | 125 | A | 0.689 | 0.691 | 0.999 | 0.381 |
| ILE | 55 | A | 0.764 | 0.784 | 0.998 | 0.55 | | TYR | 126 | A | 0.863 | 0.857 | 0.999 | 0.721 |
| PRO | 56 | A | 0.715 | 0.717 | 0.996 | 0.436 | | THR | 127 | A | 0.639 | 0.611 | 0.999 | 0.251 |
| TYR | 57 | A | 0.827 | 0.836 | 0.998 | 0.665 | | VAL | 128 | A | 0.715 | 0.717 | 0.995 | 0.437 |
| PHE | 58 | A | 0.826 | 0.839 | 0.981 | 0.684 | | GLY | 129 | A | 0.625 | 0.623 | 1 | 0.248 |
| **GLU** | **59** | **A** | **0.792** | **0.632** | **0.62** | **0.804** | | ALA | 130 | A | 0.597 | 0.597 | 1 | 0.194 |
| GLN | 60 | A | 0.786 | 0.599 | 0.702 | 0.683 | | VAL | 131 | A | 0.694 | 0.695 | 0.997 | 0.392 |
| PHE | 61 | A | 0.833 | 0.835 | 0.996 | 0.672 | | LEU | 132 | A | 0.786 | 0.78 | 0.999 | 0.567 |
| ILE | 62 | A | 0.732 | 0.709 | 0.989 | 0.452 | | SER | 133 | A | 0.685 | 0.706 | 0.999 | 0.392 |
| ASP | 63 | A | 0.816 | 0.813 | 0.992 | 0.637 | | LEU | 134 | A | 0.719 | 0.755 | 0.99 | 0.484 |
| ARG | 64 | A | 0.623 | 0.576 | 1 | 0.199 | | ALA | 135 | A | 0.49 | 0.53 | 0.995 | 0.025 |
| PHE | 65 | A | 0.854 | 0.836 | 0.999 | 0.691 | | TYR | 136 | A | 0.84 | 0.846 | 0.996 | 0.69 |
| PRO | 66 | A | 0.718 | 0.726 | 1 | 0.444 | | GLY | 137 | A | 0.64 | 0.637 | 0.998 | 0.279 |
| THR | 67 | A | 0.769 | 0.798 | 0.984 | 0.583 | | VAL | 138 | A | 0.716 | 0.71 | 0.993 | 0.433 |
| LEU | 68 | A | 0.741 | 0.775 | 0.998 | 0.518 | | PRO | 139 | A | 0.631 | 0.653 | 0.971 | 0.313 |
| GLU | 69 | A | 0.652 | 0.678 | 1 | 0.33 | | GLU | 140 | A | 0.611 | 0.548 | 0.957 | 0.202 |
| ALA | 70 | A | 0.574 | 0.579 | 1 | 0.153 | | PRO | 141 | A | 0.625 | 0.636 | 0.982 | 0.279 |
| **LEU** | **71** | **A** | **0.715** | **0.742** | **0.495** | **0.962** | | ALA | 142 | A | 0.401 | 0.42 | 0.995 | -0.174 |
| ALA | 72 | A | 0.542 | 0.578 | 0.999 | 0.121 | | VAL | 143 | A | 0.685 | 0.652 | 0.998 | 0.339 |
| ASP | 73 | A | 0.707 | 0.768 | 0.944 | 0.531 | | ASN | 144 | A | 0.769 | 0.708 | 0.94 | 0.537 |
| ALA | 74 | A | 0.535 | 0.526 | 0.978 | 0.083 | | GLY | 145 | A | 0.635 | 0.631 | 0.97 | 0.296 |
| ASP | 75 | A | 0.645 | 0.784 | 0.76 | 0.669 | | **ASN** | **146** | **A** | **0.756** | **0.687** | **0.621** | **0.822** |
| **GLU** | **76** | **A** | **0.719** | **0.746** | **0.648** | **0.817** | | VAL | 147 | A | 0.671 | 0.695 | 0.999 | 0.367 |
| ASP | 77 | A | 0.684 | 0.678 | 0.969 | 0.393 | | MET | 148 | A | 0.755 | 0.77 | 0.959 | 0.566 |
| GLU | 78 | A | 0.599 | 0.599 | 0.999 | 0.199 | | ARG | 149 | A | 0.836 | 0.821 | 0.986 | 0.671 |
| VAL | 79 | A | 0.692 | 0.745 | 0.997 | 0.44 | | VAL | 150 | A | 0.698 | 0.713 | 0.994 | 0.417 |

**Table S9: 1RRS (P), 1RRQ (Q)**, continued..

| Residue | | | S (P) | S (Q) | J | PRI |
|---|---|---|---|---|---|---|
| LEU | 151 | A | 0.695 | 0.714 | 0.991 | 0.418 |
| SER | 152 | A | 0.697 | 0.699 | 0.998 | 0.398 |
| ARG | 153 | A | 0.854 | 0.857 | 0.995 | 0.716 |
| LEU | 154 | A | 0.748 | 0.748 | 0.993 | 0.503 |
| PHE | 155 | A | 0.798 | 0.776 | 0.984 | 0.59 |
| LEU | 156 | A | 0.697 | 0.66 | 0.951 | 0.406 |
| VAL | 157 | A | 0.726 | 0.687 | 0.998 | 0.415 |
| THR | 158 | A | 0.704 | 0.705 | 0.998 | 0.411 |
| ASP | 159 | A | 0.756 | 0.737 | 0.998 | 0.495 |
| ASP | 160 | A | 0.788 | 0.76 | 0.804 | 0.744 |
| ILE | 161 | A | 0.738 | 0.744 | 0.99 | 0.492 |
| ALA | 162 | A | 0.559 | 0.621 | 0.87 | 0.31 |
| LYS | 163 | A | 0.649 | 0.707 | 0.976 | 0.38 |
| CYS | 164 | A | 0.696 | 0.683 | 0.98 | 0.399 |
| SER | 165 | A | 0.603 | 0.689 | 0.996 | 0.296 |
| THR | 166 | A | 0.675 | 0.657 | 0.985 | 0.347 |
| ARG | 167 | A | 0.829 | 0.771 | 0.993 | 0.607 |
| LYS | 168 | A | 0.774 | 0.752 | 0.994 | 0.532 |
| ARG | 169 | A | 0.602 | 0.626 | 0.999 | 0.229 |
| PHE | 170 | A | 0.839 | 0.853 | 0.987 | 0.705 |
| GLU | 171 | A | 0.816 | 0.813 | 0.996 | 0.633 |
| GLN | 172 | A | 0.658 | 0.641 | 0.997 | 0.302 |
| ILE | 173 | A | 0.711 | 0.715 | 1 | 0.426 |
| VAL | 174 | A | 0.706 | 0.701 | 1 | 0.407 |
| ARG | 175 | A | 0.837 | 0.848 | 0.995 | 0.69 |
| GLU | 176 | A | 0.707 | 0.734 | 0.835 | 0.606 |
| ILE | 177 | A | 0.738 | 0.71 | 0.994 | 0.454 |
| MET | 178 | A | 0.716 | 0.73 | 0.962 | 0.484 |
| ALA | 179 | A | 0.605 | 0.544 | 0.999 | 0.15 |
| TYR | 180 | A | 0.808 | 0.807 | 0.996 | 0.619 |
| GLU | 181 | A | 0.662 | 0.666 | 0.813 | 0.515 |
| ASN | 182 | A | 0.777 | 0.805 | 0.997 | 0.585 |
| PRO | 183 | A | 0.743 | 0.761 | 1 | 0.504 |
| GLY | 184 | A | 0.624 | 0.622 | 1 | 0.246 |
| ALA | 185 | A | 0.567 | 0.64 | 0.989 | 0.218 |
| PHE | 186 | A | 0.841 | 0.852 | 1 | 0.693 |
| ASN | 187 | A | 0.687 | 0.729 | 0.999 | 0.417 |
| GLU | 188 | A | 0.8 | 0.819 | 0.957 | 0.662 |
| ALA | 189 | A | 0.619 | 0.586 | 0.984 | 0.221 |
| LEU | 190 | A | 0.678 | 0.637 | 0.997 | 0.318 |
| ILE | 191 | A | 0.75 | 0.764 | 0.992 | 0.522 |
| GLU | 192 | A | 0.836 | 0.846 | 0.99 | 0.692 |
| LEU | 193 | A | 0.72 | 0.724 | 0.996 | 0.448 |
| GLY | 194 | A | 0.618 | 0.622 | 0.997 | 0.243 |
| ALA | 195 | A | 0.493 | 0.503 | 0.993 | 0.003 |
| LEU | 196 | A | 0.743 | 0.734 | 0.992 | 0.485 |
| VAL | 197 | A | 0.697 | 0.689 | 0.999 | 0.387 |
| CYS | 198 | A | 0.561 | 0.521 | 0.997 | 0.085 |
| THR | 199 | A | 0.691 | 0.682 | 0.992 | 0.381 |
| PRO | 200 | A | 0.734 | 0.682 | 0.999 | 0.417 |
| ARG | 201 | A | 0.669 | 0.668 | 1 | 0.337 |
| ARG | 202 | A | 0.65 | 0.652 | 0.993 | 0.309 |
| PRO | 203 | A | 0.625 | 0.624 | 0.983 | 0.266 |
| SER | 204 | A | 0.599 | 0.598 | 0.995 | 0.202 |
| CYS | 205 | A | 0.588 | 0.628 | 1 | 0.216 |
| LEU | 206 | A | 0.571 | 0.596 | 0.978 | 0.189 |
| LEU | 207 | A | 0.502 | 0.524 | 0.945 | 0.081 |
| CYS | 208 | A | 0.616 | 0.636 | 0.981 | 0.271 |
| PRO | 209 | A | 0.664 | 0.643 | 0.946 | 0.361 |
| VAL | 210 | A | 0.696 | 0.731 | 0.994 | 0.433 |
| **GLN** | **211** | **A** | **0.818** | **0.834** | **0.585** | **1.067** |
| ALA | 212 | A | 0.55 | 0.573 | 0.989 | 0.134 |
| TYR | 213 | A | 0.838 | 0.841 | 0.991 | 0.688 |
| CYS | 214 | A | 0.603 | 0.605 | 0.994 | 0.214 |
| GLN | 215 | A | 0.793 | 0.815 | 0.965 | 0.643 |
| ALA | 216 | A | 0.611 | 0.537 | 0.997 | 0.151 |
| PHE | 217 | A | 0.831 | 0.845 | 0.988 | 0.688 |
| ALA | 218 | A | 0.566 | 0.562 | 1 | 0.128 |

| Residue | | | S (P) | S (Q) | J | PRI |
|---|---|---|---|---|---|---|
| GLU | 219 | A | 0.619 | 0.627 | 0.995 | 0.251 |
| GLY | 220 | A | 0.646 | 0.649 | 1 | 0.295 |
| VAL | 221 | A | 0.722 | 0.725 | 1 | 0.447 |
| ALA | 222 | A | 0.542 | 0.592 | 0.995 | 0.139 |
| **GLU** | **223** | **A** | **0.753** | **0.78** | **0.674** | **0.859** |
| GLU | 224 | A | 0.762 | 0.748 | 0.985 | 0.525 |
| LEU | 225 | A | 0.731 | 0.728 | 0.998 | 0.461 |
| PRO | 226 | A | 0.676 | 0.618 | 0.994 | 0.3 |
| VAL | 227 | A | 0.661 | 0.743 | 0.946 | 0.458 |
| LYS | 228 | A | 0.604 | 0.606 | 0.998 | 0.212 |
| MET | 229 | A | 0.683 | 0.629 | 0.955 | 0.357 |
| VAL | 234 | A | 0.756 | 0.783 | 0.964 | 0.575 |
| LYS | 235 | A | 0.664 | 0.723 | 0.946 | 0.441 |
| GLN | 236 | A | 0.767 | 0.794 | 0.887 | 0.674 |
| VAL | 237 | A | 0.725 | 0.735 | 0.999 | 0.461 |
| PRO | 238 | A | 0.681 | 0.668 | 0.995 | 0.354 |
| LEU | 239 | A | 0.758 | 0.738 | 0.984 | 0.512 |
| ALA | 240 | A | 0.527 | 0.523 | 0.996 | 0.054 |
| VAL | 241 | A | 0.718 | 0.712 | 0.999 | 0.431 |
| ALA | 242 | A | 0.471 | 0.459 | 0.999 | -0.069 |
| VAL | 243 | A | 0.671 | 0.671 | 0.998 | 0.344 |
| **LEU** | **244** | **A** | **0.711** | **0.71** | **0.634** | **0.787** |
| ALA | 245 | A | 0.558 | 0.526 | 0.999 | 0.085 |
| ASP | 246 | A | 0.796 | 0.782 | 0.995 | 0.583 |
| ASP | 247 | A | 0.496 | 0.444 | 0.943 | -0.003 |
| GLU | 248 | A | 0.643 | 0.667 | 0.846 | 0.464 |
| GLY | 249 | A | 0.648 | 0.634 | 0.962 | 0.32 |
| ARG | 250 | A | 0.802 | 0.828 | 0.979 | 0.651 |
| VAL | 251 | A | 0.697 | 0.712 | 0.999 | 0.41 |
| LEU | 252 | A | 0.754 | 0.731 | 0.995 | 0.49 |
| ILE | 253 | A | 0.744 | 0.746 | 0.997 | 0.493 |
| ARG | 254 | A | 0.798 | 0.793 | 0.987 | 0.604 |
| LYS | 255 | A | 0.586 | 0.585 | 0.999 | 0.172 |
| ARG | 256 | A | 0.865 | 0.857 | 1 | 0.722 |
| ASP | 257 | A | 0.811 | 0.816 | 0.997 | 0.63 |
| SER | 258 | A | 0.637 | 0.635 | 0.99 | 0.282 |
| THR | 259 | A | 0.788 | 0.768 | 0.996 | 0.56 |
| GLY | 260 | A | 0.633 | 0.635 | 0.999 | 0.269 |
| LEU | 261 | A | 0.757 | 0.749 | 0.983 | 0.523 |
| LEU | 262 | A | 0.732 | 0.722 | 0.998 | 0.456 |
| ALA | 263 | A | 0.509 | 0.5 | 0.999 | 0.01 |
| ASN | 264 | A | 0.742 | 0.698 | 0.8 | 0.64 |
| LEU | 265 | A | 0.675 | 0.684 | 0.968 | 0.391 |
| **TRP** | **266** | **A** | **0.891** | **0.885** | **1** | **0.776** |
| GLU | 267 | A | 0.802 | 0.815 | 0.969 | 0.648 |
| PHE | 268 | A | 0.834 | 0.827 | 0.992 | 0.669 |
| PRO | 269 | A | 0.636 | 0.637 | 0.995 | 0.278 |
| SER | 270 | A | 0.677 | 0.616 | 0.997 | 0.296 |
| CYS | 271 | A | 0.655 | 0.682 | 0.984 | 0.353 |
| GLU | 272 | A | 0.428 | 0.421 | 0.964 | -0.115 |
| THR | 273 | A | 0.695 | 0.583 | 0.975 | 0.303 |
| ASP | 274 | A | 0.635 | 0.643 | 0.998 | 0.28 |
| GLY | 275 | A | 0.658 | 0.66 | 0.995 | 0.323 |
| ALA | 276 | A | 0.655 | 0.654 | 0.995 | 0.314 |
| ASP | 277 | A | 0.65 | 0.652 | 0.993 | 0.309 |
| GLY | 278 | A | 0.621 | 0.626 | 0.969 | 0.278 |
| LYS | 279 | A | 0.603 | 0.559 | 0.986 | 0.176 |
| GLU | 280 | A | 0.651 | 0.689 | 0.95 | 0.39 |
| LYS | 281 | A | 0.605 | 0.599 | 0.987 | 0.217 |
| **LEU** | **282** | **A** | **0.743** | **0.716** | **0.573** | **0.886** |
| GLU | 283 | A | 0.686 | 0.77 | 0.99 | 0.466 |
| GLN | 284 | A | 0.563 | 0.682 | 0.978 | 0.267 |
| MET | 285 | A | 0.726 | 0.686 | 0.992 | 0.42 |
| VAL | 286 | A | 0.752 | 0.704 | 0.949 | 0.507 |
| GLY | 287 | A | 0.556 | 0.755 | 0.918 | 0.393 |
| LEU | 292 | A | 0.58 | 0.709 | 0.722 | 0.567 |
| GLN | 293 | A | 0.633 | 0.635 | 0.961 | 0.307 |
| VAL | 294 | A | 0.62 | 0.628 | 1 | 0.248 |

| Residue | | | S (P) | S (Q) | J | PRI |
|---|---|---|---|---|---|---|
| GLU | 295 | A | 0.627 | 0.6 | 0.995 | 0.232 |
| LEU | 296 | A | 0.659 | 0.701 | 0.957 | 0.403 |
| THR | 297 | A | 0.722 | 0.709 | 0.998 | 0.433 |
| GLU | 298 | A | 0.639 | 0.637 | 1 | 0.276 |
| PRO | 299 | A | 0.648 | 0.731 | 0.906 | 0.473 |
| ILE | 300 | A | 0.627 | 0.624 | 0.999 | 0.252 |
| VAL | 301 | A | 0.678 | 0.674 | 0.999 | 0.353 |
| SER | 302 | A | 0.643 | 0.67 | 0.997 | 0.316 |
| PHE | 303 | A | 0.811 | 0.802 | 0.999 | 0.614 |
| GLU | 304 | A | 0.726 | 0.706 | 0.941 | 0.491 |
| HIS | 305 | A | 0.819 | 0.833 | 0.996 | 0.656 |
| ALA | 306 | A | 0.582 | 0.586 | 1 | 0.168 |
| PHE | 307 | A | 0.813 | 0.824 | 0.997 | 0.64 |
| SER | 308 | A | 0.693 | 0.686 | 1 | 0.379 |
| HIS | 309 | A | 0.815 | 0.813 | 0.992 | 0.636 |
| LEU | 310 | A | 0.732 | 0.739 | 0.923 | 0.548 |
| VAL | 311 | A | 0.72 | 0.739 | 0.998 | 0.461 |
| **TRP** | **312** | **A** | **0.897** | **0.889** | **0.998** | **0.788** |
| GLN | 313 | A | 0.792 | 0.77 | 0.832 | 0.73 |
| LEU | 314 | A | 0.662 | 0.67 | 0.994 | 0.338 |
| THR | 315 | A | 0.754 | 0.735 | 0.995 | 0.494 |
| VAL | 316 | A | 0.485 | 0.481 | 0.999 | -0.033 |
| PHE | 317 | A | 0.843 | 0.846 | 0.999 | 0.69 |
| PRO | 318 | A | 0.66 | 0.663 | 0.999 | 0.324 |
| GLY | 319 | A | 0.619 | 0.621 | 0.998 | 0.242 |
| ARG | 320 | A | 0.573 | 0.546 | 0.994 | 0.125 |
| LEU | 321 | A | 0.706 | 0.707 | 0.978 | 0.435 |
| VAL | 322 | A | 0.53 | 0.543 | 1 | 0.073 |
| HIS | 323 | A | 0.7 | 0.724 | 0.986 | 0.438 |
| GLY | 324 | A | 0.646 | 0.65 | 0.995 | 0.301 |
| GLY | 325 | A | 0.654 | 0.654 | 1 | 0.308 |
| PRO | 326 | A | 0.753 | 0.744 | 0.985 | 0.512 |
| VAL | 327 | A | 0.65 | 0.623 | 0.611 | 0.662 |
| GLU | 328 | A | 0.616 | 0.618 | 0.999 | 0.235 |
| GLU | 329 | A | 0.648 | 0.651 | 0.968 | 0.331 |
| PRO | 330 | A | 0.702 | 0.707 | 0.926 | 0.483 |
| TYR | 331 | A | 0.841 | 0.833 | 0.993 | 0.681 |
| **ARG** | **332** | **A** | **0.85** | **0.806** | **0.873** | **0.783** |
| LEU | 333 | A | 0.727 | 0.749 | 0.993 | 0.483 |
| ALA | 334 | A | 0.507 | 0.511 | 0.999 | 0.019 |
| PRO | 335 | A | 0.701 | 0.708 | 1 | 0.409 |
| GLU | 336 | A | 0.772 | 0.757 | 0.951 | 0.578 |
| ASP | 337 | A | 0.74 | 0.741 | 0.962 | 0.519 |
| GLU | 338 | A | 0.567 | 0.535 | 0.999 | 0.103 |
| LEU | 339 | A | 0.743 | 0.738 | 0.987 | 0.494 |
| LYS | 340 | A | 0.626 | 0.649 | 0.994 | 0.281 |
| ALA | 341 | A | 0.602 | 0.562 | 1 | 0.164 |
| TYR | 342 | A | 0.853 | 0.847 | 0.999 | 0.701 |
| ALA | 343 | A | 0.42 | 0.427 | 1 | -0.153 |
| PHE | 344 | A | 0.827 | 0.834 | 0.999 | 0.662 |
| PRO | 345 | A | 0.684 | 0.686 | 0.998 | 0.372 |
| VAL | 346 | A | 0.658 | 0.668 | 0.997 | 0.329 |
| SER | 347 | A | 0.682 | 0.701 | 0.99 | 0.393 |
| HIS | 348 | A | 0.809 | 0.774 | 0.989 | 0.594 |
| GLN | 349 | A | 0.665 | 0.733 | 0.998 | 0.4 |
| ARG | 350 | A | 0.879 | 0.86 | 0.99 | 0.749 |
| VAL | 351 | A | 0.732 | 0.696 | 0.986 | 0.442 |
| **TRP** | **352** | **A** | **0.892** | **0.877** | **0.993** | **0.776** |
| **ARG** | **353** | **A** | **0.832** | **0.821** | **0.82** | **0.833** |
| GLU | 354 | A | 0.745 | 0.789 | 0.995 | 0.539 |
| TYR | 355 | A | 0.855 | 0.845 | 0.983 | 0.717 |
| LYS | 356 | A | 0.563 | 0.566 | 0.997 | 0.132 |
| GLU | 357 | A | 0.816 | 0.78 | 0.924 | 0.672 |
| TRP | 358 | A | 0.833 | 0.823 | 0.999 | 0.657 |
| ALA | 359 | A | 0.456 | 0.373 | 0.689 | 0.14 |
| SER | 360 | A | 0.778 | 0.563 | 0.991 | 0.35 |

**Table S10(a): 6W0D (P), 6W0F (Q)**

| Residue | | | S (P) | S (Q) | J (P,Q) | PRI |
|---|---|---|---|---|---|---|
| ALA | 28 | C | 0.811 | 0.689 | 0.895 | 0.605 |
| ALA | 29 | C | 0.681 | 0.758 | 0.966 | 0.473 |
| GLY | 30 | C | 0.634 | 0.647 | 0.994 | 0.287 |
| ALA | 31 | C | 0.678 | 0.694 | 1 | 0.372 |
| ALA | 32 | C | 0.563 | 0.535 | 0.987 | 0.111 |
| THR | 33 | C | 0.755 | 0.714 | 0.993 | 0.476 |
| VAL | 34 | C | 0.77 | 0.785 | 0.998 | 0.557 |
| LEU | 35 | C | 0.664 | 0.725 | 0.99 | 0.399 |
| LEU | 36 | C | 0.728 | 0.745 | 0.996 | 0.477 |
| VAL | 37 | C | 0.76 | 0.647 | 0.992 | 0.415 |
| ILE | 38 | C | 0.792 | 0.756 | 0.997 | 0.551 |
| VAL | 39 | C | 0.685 | 0.673 | 0.997 | 0.361 |
| LEU | 40 | C | 0.742 | 0.75 | 0.998 | 0.494 |
| LEU | 41 | C | 0.728 | 0.782 | 0.999 | 0.511 |
| ALA | 42 | C | 0.632 | 0.604 | 0.997 | 0.239 |
| GLY | 43 | C | 0.624 | 0.632 | 1 | 0.256 |
| SER | 44 | C | 0.739 | 0.687 | 0.996 | 0.43 |
| TYR | 45 | C | 0.821 | 0.857 | 0.998 | 0.68 |
| LEU | 46 | C | 0.719 | 0.612 | 0.992 | 0.339 |
| ALA | 47 | C | 0.597 | 0.557 | 0.999 | 0.155 |
| VAL | 48 | C | 0.706 | 0.71 | 0.997 | 0.419 |
| LEU | 49 | C | 0.654 | 0.68 | 0.991 | 0.343 |
| ALA | 50 | C | 0.497 | 0.519 | 0.998 | 0.018 |
| GLU | 51 | C | 0.8 | 0.792 | 0.996 | 0.596 |
| ARG | 52 | C | 0.776 | 0.759 | 0.992 | 0.543 |
| GLY | 53 | C | 0.652 | 0.661 | 0.992 | 0.321 |
| ALA | 54 | C | 0.482 | 0.483 | 0.997 | -0.032 |
| PRO | 55 | C | 0.722 | 0.702 | 0.999 | 0.425 |
| GLY | 56 | C | 0.654 | 0.66 | 1 | 0.314 |
| ALA | 57 | C | 0.476 | 0.478 | 0.997 | -0.043 |
| GLN | 58 | C | 0.693 | 0.724 | 0.973 | 0.444 |
| LEU | 59 | C | 0.709 | 0.68 | 0.998 | 0.391 |
| ILE | 60 | C | 0.778 | 0.77 | 0.999 | 0.549 |
| THR | 61 | C | 0.77 | 0.75 | 0.996 | 0.524 |
| TYR | 62 | C | 0.81 | 0.832 | 0.999 | 0.643 |
| PRO | 63 | C | 0.745 | 0.72 | 1 | 0.465 |
| **ARG** | **64** | **C** | **0.818** | **0.879** | **0.999** | **0.698** |
| ALA | 65 | C | 0.547 | 0.54 | 0.997 | 0.09 |
| LEU | 66 | C | 0.713 | 0.683 | 0.997 | 0.399 |
| TRP | 67 | C | 0.787 | 0.819 | 0.996 | 0.61 |
| **TRP** | **68** | **C** | **0.887** | **0.891** | **0.99** | **0.788** |
| SER | 69 | C | 0.649 | 0.594 | 0.999 | 0.244 |
| VAL | 70 | C | 0.74 | 0.666 | 0.99 | 0.416 |
| GLU | 71 | C | 0.795 | 0.766 | 0.926 | 0.635 |
| THR | 72 | C | 0.712 | 0.717 | 0.996 | 0.433 |
| ALA | 73 | C | 0.549 | 0.602 | 0.994 | 0.157 |
| THR | 74 | C | 0.724 | 0.561 | 0.989 | 0.296 |
| THR | 75 | C | 0.703 | 0.685 | 0.977 | 0.411 |
| VAL | 76 | C | 0.715 | 0.603 | 0.993 | 0.325 |
| GLY | 77 | C | 0.657 | 0.66 | 0.996 | 0.321 |
| **TYR** | **78** | **C** | **0.844** | **0.848** | **0.989** | **0.703** |
| GLY | 79 | C | 0.645 | 0.662 | 0.997 | 0.31 |
| ASP | 80 | C | 0.753 | 0.723 | 0.985 | 0.491 |
| LEU | 81 | C | 0.753 | 0.724 | 0.95 | 0.527 |
| TYR | 82 | C | 0.737 | 0.77 | 0.998 | 0.509 |
| PRO | 83 | C | 0.634 | 0.631 | 0.997 | 0.268 |
| VAL | 84 | C | 0.703 | 0.728 | 0.991 | 0.44 |
| THR | 85 | C | 0.736 | 0.758 | 0.99 | 0.504 |
| LEU | 86 | C | 0.691 | 0.686 | 0.998 | 0.379 |
| **TRP** | **87** | **C** | **0.875** | **0.811** | **0.995** | **0.691** |
| GLY | 88 | C | 0.625 | 0.631 | 0.999 | 0.257 |
| **ARG** | **89** | **C** | **0.878** | **0.852** | **0.992** | **0.738** |
| CYS | 90 | C | 0.619 | 0.655 | 0.999 | 0.275 |
| VAL | 91 | C | 0.692 | 0.696 | 0.993 | 0.395 |
| ALA | 92 | C | 0.613 | 0.591 | 0.997 | 0.207 |
| VAL | 93 | C | 0.758 | 0.76 | 0.998 | 0.52 |
| VAL | 94 | C | 0.724 | 0.715 | 0.999 | 0.44 |
| VAL | 95 | C | 0.727 | 0.712 | 0.982 | 0.457 |
| MET | 96 | C | 0.758 | 0.792 | 0.996 | 0.554 |
| VAL | 97 | C | 0.76 | 0.768 | 0.988 | 0.54 |
| ALA | 98 | C | 0.564 | 0.578 | 0.991 | 0.151 |
| GLY | 99 | C | 0.619 | 0.628 | 0.992 | 0.255 |
| ILE | 100 | C | 0.789 | 0.798 | 0.997 | 0.59 |
| THR | 101 | C | 0.741 | 0.665 | 0.991 | 0.415 |
| SER | 102 | C | 0.746 | 0.709 | 0.986 | 0.469 |
| PHE | 103 | C | 0.726 | 0.812 | 0.968 | 0.57 |
| GLY | 104 | C | 0.64 | 0.649 | 0.999 | 0.29 |
| LEU | 105 | C | 0.72 | 0.784 | 0.948 | 0.556 |
| VAL | 106 | C | 0.713 | 0.716 | 0.986 | 0.443 |
| THR | 107 | C | 0.765 | 0.559 | 0.992 | 0.332 |
| ALA | 108 | C | 0.66 | 0.707 | 0.991 | 0.376 |
| ALA | 109 | C | 0.587 | 0.68 | 0.992 | 0.275 |
| LEU | 110 | C | 0.722 | 0.534 | 0.988 | 0.268 |
| ALA | 111 | C | 0.682 | 0.711 | 0.999 | 0.394 |
| THR | 112 | C | 0.702 | 0.811 | 0.982 | 0.531 |
| **TRP** | **113** | **C** | **0.833** | **0.868** | **0.925** | **0.776** |
| **PHE** | **114** | **C** | **0.852** | **0.819** | **0.983** | **0.688** |
| VAL | 115 | C | 0.782 | 0.687 | 0.99 | 0.479 |
| GLY | 116 | C | 0.638 | 0.639 | 0.998 | 0.279 |
| ARG | 117 | C | 0.493 | 0.604 | 0.997 | 0.1 |
| GLU | 118 | C | 0.704 | 0.744 | 0.969 | 0.479 |
| GLN | 119 | C | 0.706 | 0.748 | 0.99 | 0.464 |
| **GLU** | **120** | **C** | **0.871** | **0.819** | **0.933** | **0.757** |

| Table S10(b): 7MHR (P), 7MK6 (Q) | | | | | | | | | | | | | | |
|---|---|---|---|---|---|---|---|---|---|---|---|---|---|---|
| Residue | | | S (P) | S (Q) | J (P,Q) | PRI | | Residue | | | S (P) | S (Q) | J (P,Q) | PRI |
| **ALA** | **28** | **C** | **0.832** | **0.821** | **0.957** | **0.696** | | VAL | 91 | C | 0.704 | 0.702 | 0.991 | 0.415 |
| ALA | 29 | C | 0.679 | 0.659 | 0.997 | 0.341 | | ALA | 92 | C | 0.571 | 0.683 | 0.998 | 0.256 |
| GLY | 30 | C | 0.631 | 0.64 | 0.998 | 0.273 | | VAL | 93 | C | 0.748 | 0.79 | 0.997 | 0.541 |
| ALA | 31 | C | 0.671 | 0.699 | 0.999 | 0.371 | | VAL | 94 | C | 0.715 | 0.745 | 0.997 | 0.463 |
| ALA | 32 | C | 0.543 | 0.647 | 0.992 | 0.198 | | VAL | 95 | C | 0.704 | 0.641 | 0.99 | 0.355 |
| THR | 33 | C | 0.752 | 0.756 | 0.997 | 0.511 | | MET | 96 | C | 0.762 | 0.647 | 0.982 | 0.427 |
| VAL | 34 | C | 0.756 | 0.822 | 0.997 | 0.581 | | VAL | 97 | C | 0.748 | 0.815 | 0.987 | 0.576 |
| LEU | 35 | C | 0.675 | 0.704 | 0.993 | 0.386 | | ALA | 98 | C | 0.575 | 0.65 | 0.998 | 0.227 |
| LEU | 36 | C | 0.735 | 0.749 | 0.985 | 0.499 | | GLY | 99 | C | 0.616 | 0.619 | 0.969 | 0.266 |
| VAL | 37 | C | 0.734 | 0.748 | 0.998 | 0.484 | | ILE | 100 | C | 0.791 | 0.842 | 0.99 | 0.643 |
| ILE | 38 | C | 0.778 | 0.837 | 0.997 | 0.618 | | THR | 101 | C | 0.704 | 0.728 | 0.973 | 0.459 |
| VAL | 39 | C | 0.71 | 0.697 | 0.997 | 0.41 | | SER | 102 | C | 0.734 | 0.663 | 0.958 | 0.439 |
| LEU | 40 | C | 0.701 | 0.789 | 0.994 | 0.496 | | PHE | 103 | C | 0.754 | 0.842 | 0.975 | 0.621 |
| LEU | 41 | C | 0.745 | 0.652 | 0.996 | 0.401 | | GLY | 104 | C | 0.647 | 0.64 | 0.998 | 0.289 |
| ALA | 42 | C | 0.606 | 0.694 | 0.998 | 0.302 | | LEU | 105 | C | 0.668 | 0.764 | 0.986 | 0.446 |
| GLY | 43 | C | 0.63 | 0.621 | 0.998 | 0.253 | | VAL | 106 | C | 0.722 | 0.683 | 0.987 | 0.418 |
| SER | 44 | C | 0.708 | 0.723 | 0.994 | 0.437 | | THR | 107 | C | 0.765 | 0.653 | 0.989 | 0.429 |
| **TYR** | **45** | **C** | **0.868** | **0.834** | **0.996** | **0.706** | | ALA | 108 | C | 0.605 | 0.656 | 0.989 | 0.272 |
| LEU | 46 | C | 0.737 | 0.797 | 0.998 | 0.536 | | ALA | 109 | C | 0.585 | 0.651 | 0.998 | 0.238 |
| ALA | 47 | C | 0.606 | 0.588 | 0.999 | 0.195 | | LEU | 110 | C | 0.753 | 0.749 | 0.994 | 0.508 |
| VAL | 48 | C | 0.705 | 0.764 | 0.998 | 0.471 | | ALA | 111 | C | 0.603 | 0.621 | 0.999 | 0.225 |
| LEU | 49 | C | 0.64 | 0.669 | 0.99 | 0.319 | | THR | 112 | C | 0.694 | 0.799 | 0.985 | 0.508 |
| ALA | 50 | C | 0.51 | 0.541 | 0.999 | 0.052 | | **TRP** | **113** | **C** | **0.839** | **0.834** | **0.994** | **0.679** |
| GLU | 51 | C | 0.823 | 0.787 | 0.992 | 0.618 | | **PHE** | **114** | **C** | **0.847** | **0.815** | **0.964** | **0.698** |
| ARG | 52 | C | 0.78 | 0.784 | 0.981 | 0.583 | | VAL | 115 | C | 0.63 | 0.683 | 1 | 0.313 |
| GLY | 53 | C | 0.642 | 0.656 | 0.99 | 0.308 | | GLY | 116 | C | 0.668 | 0.658 | 0.999 | 0.327 |
| ALA | 54 | C | 0.511 | 0.535 | 0.998 | 0.048 | | ARG | 117 | C | 0.591 | 0.655 | 0.999 | 0.247 |
| PRO | 55 | C | 0.716 | 0.733 | 0.998 | 0.451 | | GLU | 118 | C | 0.77 | 0.649 | 0.998 | 0.421 |
| GLY | 56 | C | 0.656 | 0.654 | 1 | 0.31 | | | | | | | | |
| ALA | 57 | C | 0.553 | 0.48 | 0.996 | 0.037 | | | | | | | | |
| GLN | 58 | C | 0.712 | 0.754 | 0.957 | 0.509 | | | | | | | | |
| LEU | 59 | C | 0.702 | 0.682 | 0.997 | 0.387 | | | | | | | | |
| ILE | 60 | C | 0.772 | 0.785 | 0.999 | 0.558 | | | | | | | | |
| THR | 61 | C | 0.764 | 0.772 | 0.997 | 0.539 | | | | | | | | |
| TYR | 62 | C | 0.81 | 0.812 | 0.968 | 0.654 | | | | | | | | |
| PRO | 63 | C | 0.735 | 0.749 | 0.999 | 0.485 | | | | | | | | |
| ARG | 64 | C | 0.829 | 0.842 | 0.997 | 0.674 | | | | | | | | |
| ALA | 65 | C | 0.541 | 0.553 | 0.998 | 0.096 | | | | | | | | |
| LEU | 66 | C | 0.728 | 0.697 | 0.999 | 0.426 | | | | | | | | |
| **TRP** | **67** | **C** | **0.895** | **0.83** | **0.965** | **0.76** | | | | | | | | |
| **TRP** | **68** | **C** | **0.875** | **0.899** | **0.991** | **0.783** | | | | | | | | |
| SER | 69 | C | 0.639 | 0.625 | 0.999 | 0.265 | | | | | | | | |
| VAL | 70 | C | 0.657 | 0.711 | 0.989 | 0.379 | | | | | | | | |
| VAL | 71 | C | 0.731 | 0.713 | 0.997 | 0.447 | | | | | | | | |
| THR | 72 | C | 0.706 | 0.562 | 0.991 | 0.277 | | | | | | | | |
| ALA | 73 | C | 0.561 | 0.494 | 1 | 0.055 | | | | | | | | |
| THR | 74 | C | 0.731 | 0.699 | 0.993 | 0.437 | | | | | | | | |
| THR | 75 | C | 0.686 | 0.695 | 0.999 | 0.382 | | | | | | | | |
| VAL | 76 | C | 0.718 | 0.707 | 0.996 | 0.429 | | | | | | | | |
| GLY | 77 | C | 0.632 | 0.641 | 0.994 | 0.279 | | | | | | | | |
| **TYR** | **78** | **C** | **0.836** | **0.845** | **0.975** | **0.706** | | | | | | | | |
| GLY | 79 | C | 0.633 | 0.638 | 1 | 0.271 | | | | | | | | |
| ASP | 80 | C | 0.755 | 0.759 | 0.986 | 0.528 | | | | | | | | |
| LEU | 81 | C | 0.735 | 0.743 | 0.99 | 0.488 | | | | | | | | |
| TYR | 82 | C | 0.792 | 0.743 | 0.999 | 0.536 | | | | | | | | |
| PRO | 83 | C | 0.636 | 0.624 | 0.997 | 0.263 | | | | | | | | |
| VAL | 84 | C | 0.707 | 0.664 | 0.997 | 0.374 | | | | | | | | |
| THR | 85 | C | 0.751 | 0.74 | 0.985 | 0.506 | | | | | | | | |
| LEU | 86 | C | 0.732 | 0.751 | 0.994 | 0.489 | | | | | | | | |
| **TRP** | **87** | **C** | **0.874** | **0.824** | **0.99** | **0.708** | | | | | | | | |
| GLY | 88 | C | 0.624 | 0.624 | 0.999 | 0.249 | | | | | | | | |
| **ARG** | **89** | **C** | **0.866** | **0.862** | **0.998** | **0.73** | | | | | | | | |
| CYS | 90 | C | 0.675 | 0.666 | 0.985 | 0.356 | | | | | | | | |

**Table S10(c): 1K4C (P), 1K4D (Q)**

| Residue | | | S (P) | S (Q) | J (P,Q) | PRI |
|---|---|---|---|---|---|---|
| SER | 22 | C | 0.72 | 0.714 | 0.689 | 0.745 |
| ALA | 23 | C | 0.599 | 0.588 | 0.996 | 0.191 |
| LEU | 24 | C | 0.683 | 0.694 | 0.997 | 0.38 |
| HIS | 25 | C | 0.846 | 0.856 | 0.995 | 0.707 |
| **TRP** | **26** | **C** | **0.871** | **0.877** | **0.948** | **0.8** |
| ARG | 27 | C | 0.646 | 0.673 | 0.999 | 0.32 |
| ALA | 28 | C | 0.592 | 0.58 | 0.997 | 0.175 |
| ALA | 29 | C | 0.606 | 0.646 | 0.995 | 0.257 |
| GLY | 30 | C | 0.646 | 0.641 | 0.999 | 0.288 |
| ALA | 31 | C | 0.651 | 0.663 | 0.997 | 0.317 |
| ALA | 32 | C | 0.564 | 0.596 | 0.994 | 0.166 |
| THR | 33 | C | 0.75 | 0.741 | 0.994 | 0.497 |
| VAL | 34 | C | 0.753 | 0.761 | 0.995 | 0.519 |
| **LEU** | **35** | **C** | **0.646** | **0.697** | **0.576** | **0.767** |
| LEU | 36 | C | 0.713 | 0.694 | 0.997 | 0.41 |
| VAL | 37 | C | 0.737 | 0.76 | 0.997 | 0.5 |
| ILE | 38 | C | 0.797 | 0.796 | 0.999 | 0.594 |
| VAL | 39 | C | 0.686 | 0.697 | 0.999 | 0.384 |
| LEU | 40 | C | 0.753 | 0.731 | 0.999 | 0.485 |
| LEU | 41 | C | 0.724 | 0.711 | 0.997 | 0.438 |
| ALA | 42 | C | 0.634 | 0.637 | 0.999 | 0.272 |
| GLY | 43 | C | 0.631 | 0.631 | 0.999 | 0.263 |
| SER | 44 | C | 0.707 | 0.705 | 0.998 | 0.414 |
| TYR | 45 | C | 0.818 | 0.806 | 0.995 | 0.629 |
| LEU | 46 | C | 0.764 | 0.726 | 0.998 | 0.492 |
| ALA | 47 | C | 0.569 | 0.565 | 1 | 0.134 |
| VAL | 48 | C | 0.704 | 0.693 | 0.999 | 0.398 |
| LEU | 49 | C | 0.661 | 0.656 | 1 | 0.317 |
| ALA | 50 | C | 0.488 | 0.496 | 1 | -0.016 |
| GLU | 51 | C | 0.794 | 0.788 | 1 | 0.582 |
| ARG | 52 | C | 0.774 | 0.774 | 0.998 | 0.55 |
| GLY | 53 | C | 0.667 | 0.665 | 1 | 0.332 |
| ALA | 54 | C | 0.501 | 0.485 | 0.998 | -0.012 |
| PRO | 55 | C | 0.727 | 0.74 | 0.943 | 0.524 |
| GLY | 56 | C | 0.662 | 0.662 | 0.996 | 0.328 |
| ALA | 57 | C | 0.468 | 0.496 | 0.999 | -0.035 |
| GLN | 58 | C | 0.713 | 0.7 | 0.994 | 0.419 |
| LEU | 59 | C | 0.711 | 0.706 | 0.999 | 0.418 |
| ILE | 60 | C | 0.77 | 0.763 | 1 | 0.533 |
| THR | 61 | C | 0.766 | 0.767 | 0.996 | 0.537 |
| TYR | 62 | C | 0.849 | 0.837 | 0.996 | 0.69 |
| PRO | 63 | C | 0.749 | 0.751 | 1 | 0.5 |
| ARG | 64 | C | 0.803 | 0.785 | 0.998 | 0.59 |
| ALA | 65 | C | 0.524 | 0.539 | 0.999 | 0.064 |
| LEU | 66 | C | 0.685 | 0.69 | 0.995 | 0.38 |
| TRP | 67 | C | 0.796 | 0.85 | 0.982 | 0.664 |
| **TRP** | **68** | **C** | **0.884** | **0.893** | **0.993** | **0.784** |
| **SER** | **69** | **C** | **0.66** | **0.664** | **0.431** | **0.893** |
| VAL | 70 | C | 0.745 | 0.722 | 0.986 | 0.481 |
| GLU | 71 | C | 0.792 | 0.807 | 0.907 | 0.692 |
| THR | 72 | C | 0.686 | 0.714 | 0.987 | 0.413 |
| ALA | 73 | C | 0.495 | 0.517 | 0.983 | 0.029 |
| THR | 74 | C | 0.726 | 0.722 | 0.974 | 0.474 |
| THR | 75 | C | 0.659 | 0.646 | 0.944 | 0.361 |
| VAL | 76 | C | 0.715 | 0.672 | 0.834 | **0.553** |
| GLY | 77 | C | 0.64 | 0.653 | 0.968 | 0.325 |
| TYR | 78 | C | 0.863 | 0.856 | 0.976 | 0.743 |
| GLY | 79 | C | 0.657 | 0.656 | 0.996 | 0.317 |
| ASP | 80 | C | 0.776 | 0.747 | 0.968 | 0.555 |
| LEU | 81 | C | 0.768 | 0.743 | 0.997 | 0.514 |
| TYR | 82 | C | 0.787 | 0.73 | 0.998 | **0.519** |
| PRO | 83 | C | 0.629 | 0.634 | 0.999 | 0.264 |
| VAL | 84 | C | 0.71 | 0.703 | 0.999 | 0.414 |
| THR | 85 | C | 0.715 | 0.736 | 0.999 | 0.452 |
| LEU | 86 | C | 0.741 | 0.697 | 1 | 0.438 |
| TRP | 87 | C | 0.841 | 0.87 | 0.991 | 0.72 |
| GLY | 88 | C | 0.629 | 0.63 | 0.998 | 0.261 |
| **ARG** | **89** | **C** | **0.886** | **0.869** | **0.999** | **0.756** |
| CYS | 90 | C | 0.696 | 0.671 | 0.998 | 0.369 |
| VAL | 91 | C | 0.705 | 0.703 | 1 | 0.408 |
| ALA | 92 | C | 0.577 | 0.596 | 1 | 0.173 |
| VAL | 93 | C | 0.743 | 0.75 | 0.997 | 0.496 |
| VAL | 94 | C | 0.721 | 0.713 | 0.999 | 0.435 |
| VAL | 95 | C | 0.723 | 0.707 | 0.984 | 0.446 |
| MET | 96 | C | 0.724 | 0.745 | 0.982 | 0.487 |
| VAL | 97 | C | 0.758 | 0.762 | 0.999 | 0.521 |
| ALA | 98 | C | 0.576 | 0.58 | 0.998 | 0.158 |
| GLY | 99 | C | 0.628 | 0.627 | 0.999 | 0.256 |
| ILE | 100 | C | 0.788 | 0.785 | 0.996 | 0.577 |
| THR | 101 | C | 0.712 | 0.744 | 0.994 | 0.462 |
| SER | 102 | C | 0.706 | 0.715 | 0.999 | 0.422 |
| PHE | 103 | C | 0.722 | 0.726 | 0.999 | 0.449 |
| GLY | 104 | C | 0.649 | 0.651 | 1 | 0.3 |
| LEU | 105 | C | 0.713 | 0.728 | 0.924 | 0.517 |
| VAL | 106 | C | 0.733 | 0.732 | 0.999 | 0.466 |
| THR | 107 | C | 0.771 | 0.756 | 1 | 0.527 |
| ALA | 108 | C | 0.609 | 0.611 | 0.998 | 0.222 |
| ALA | 109 | C | 0.555 | 0.573 | 0.988 | 0.14 |
| LEU | 110 | C | 0.719 | 0.751 | 0.997 | 0.473 |
| ALA | 111 | C | 0.646 | 0.653 | 1 | 0.299 |
| THR | 112 | C | 0.744 | 0.761 | 0.995 | 0.51 |
| TRP | 113 | C | 0.855 | 0.848 | 0.976 | 0.727 |
| PHE | 114 | C | 0.846 | 0.854 | 0.994 | 0.706 |
| VAL | 115 | C | 0.771 | 0.755 | 0.999 | 0.527 |
| GLY | 116 | C | 0.648 | 0.646 | 0.999 | 0.295 |
| ARG | 117 | C | 0.62 | 0.641 | 1 | 0.261 |
| **GLU** | **118** | **C** | **0.801** | **0.726** | **0.651** | **0.876** |
| **GLN** | **119** | **C** | **0.795** | **0.782** | **0.792** | **0.785** |
| GLU | 120 | C | 0.714 | 0.742 | 0.873 | 0.583 |
| **ARG** | **121** | **C** | **0.819** | **0.828** | **0.881** | **0.766** |
| ARG | 122 | C | 0.47 | 0.489 | 0.995 | -0.036 |
| GLY | 123 | C | 0.66 | 0.667 | 0.787 | 0.54 |
| **HIS** | **124** | **C** | **0.844** | **0.827** | **0.841** | **0.83** |
